# Computer Simulations and Analyses of Coupling Among Reproductive Barriers in Late-Stage Sympatric Speciation

**DOI:** 10.1101/2025.04.21.649836

**Authors:** John Lin, Zakia Sultana, Curtis Heyda, Lauren Gieck, James Lin

## Abstract

The mechanisms driving sympatric speciation remain an unresolved challenge in evolutionary biology. In nature, closely related “good” species are observed to possess multiple different barriers in their genomes that collectively generate strong and irreversible reproductive isolation (RI). Theorists hypothesize that early-stage mechanisms responsible for establishing an initial reproductive barrier differ from those driving the coupling of barriers in later stages of sympatric speciation. In a prior study, we developed a two-allele mathematical model of mating-bias traits to demonstrate how initial premating RI can arise in a sympatric population under disruptive ecological selection. Here, we extend this model to investigate how different pre- and post-mating barriers can couple with such an initial barrier and with one another during late-stage sympatric speciation to establish strong and irreversible RI. We developed computer applications to examine the properties of various barrier mechanisms and the conditions required for their invasion and coupling. Early-stage, adaptive premating barriers, driven by maladaptive hybrid loss, are effective in establishing initial RI and coupling with other barriers but are easily reversed if disruptive ecological selection weakens. Late-stage barriers, by contrast, often rely on earlier barriers to create an environment of reduced gene flow to facilitate their invasion and coupling. Mutations reducing hybrid viability are underdominant in inter-niche matings and can only invade by hitchhiking with barriers conferring a fitness advantage. Chromosomal inversions, an adaptive late-stage mechanism, can combine different barrier properties into a supergene and gain a net fitness advantage to invade and couple with other barriers to create strong and less reversible RI. Late-stage nonadaptive Bateson–Dobzhansky–Muller (BDM) barriers evolve more slowly but confer the strongest and least reversible RI. Our findings reveal a positive feedback loop in which early-stage barriers facilitate the establishment of late-stage barriers, while late-stage barriers strengthen and secure early-stage barriers. This positive reinforcement progressively strengthens overall RI until it becomes irreversible. By examining the properties and invasion dynamics of various barrier mechanisms, this study complements our previous study to propose a comprehensive process of sympatric speciation that explains how barriers emerge and couple to complete the speciation process.

## I. INTRODUCTION

In modern evolutionary biology, sympatric speciation remains an unresolved mystery [1, 2]. In sympatric populations, disruptive ecological selection followed by assortative mating is thought to be the sequence of steps leading to sympatric speciation [3]. However, researchers struggle to understand how differential assortment of mating traits can arise between niche ecotypes in the presence of gene flow. Additionally, it appears that most of the few seemingly incontrovertible cases of sympatric speciation found in nature can always be explained away by a model of brief allopatric divergence followed by secondary contact [4]. These theoretical difficulties have led many modern-day evolutionary biologists to reject the plausibility of sympatric speciation and its prevalence [1].

In the past three decades, there has been a resurgence of interest in studying the mechanisms of sympatric speciation, leading to the development of computer simulations based on population dynamic models that demonstrate how an initial premating reproductive barrier can be established between sympatric ecotypes under disruptive ecological selection [5-8]. However, these models have often been criticized as biologically unrealistic, with the conditions required to establish reproductive isolation (RI) deemed too stringent to be feasible [9-13]. Furthermore, there appears to be a fundamental lack of understanding of how the various forces in these models interact to produce the simulation results [1, 14].

In current evolutionary thinking, speciation is considered a process rather than a single event. Incipient sister species initially experience significant gene flow and introgression between them. They then gradually develop increasingly strong reproductive isolation (RI) along a continuum until the speciation process is complete, with no gene flow or introgression [9].

In nature, closely-related “good” species that have completed the speciation process are observed to have many different barrier mechanisms in their genomes that interact to produce strong and irreversible RI [15-19]. It remains unclear how an initial reproductive barrier becomes coupled with additional barrier mechanisms to achieve this final level of strong RI. Current hypotheses suggest that the mechanisms responsible for producing the initial RI—whether through geographical isolation in allopatric speciation or premating RI in sympatric speciation—are likely different from those that subsequently recruit additional barriers to reinforce RI and make speciation irreversible [20, 21].

Many researchers have studied the coupling of different reproductive barriers in late-stage speciation [17, 19, 22-24]. These include adaptive and nonadaptive premating and post-mating barrier mechanisms, chromosomal rearrangements, and the Bateson-Dobzhansky-Muller (BDM) mechanism of divergence and incompatibilities [25, 26]. However, few of these barrier mechanisms have been studied specifically in sympatric speciation, chiefly due to the lack of a convincing model that could establish an initial barrier in sympatric speciation.

Sympatric barrier mechanisms can be described as either a one-allele model or a two-allele model [14, 27]. In a one-allele model, a mutant allele produces the same effect in different niche ecotypes. Only the substitution of a single allele is needed to produce RI. As a result, the one-allele model is unaffected by recombination caused by gene flow. In contrast, a two-allele model requires distinct mutant alleles in different niche ecotypes to achieve RI. This model is more challenging to establish because recombination can disrupt the linkage disequilibrium between mutant alleles and their associated ecotypes and hinder differential assortment [3].

In a previous study, we developed a two-allele mathematical model to demonstrate how premating RI can be established in a sympatric population under disruptive ecological selection [28]. In the model, hybrid loss creates selection pressure that drives adaptive changes in population dynamics, mediated by interactions between ecological and sexual selection, to create differential assortment of mating-bias alleles between niche ecotypes and establish premating RI. Based on our findings, a five-stage mechanism of sympatric speciation was formulated: Initially, disruptive ecological selection creates selection pressure for high-mating-bias alleles to invade and rapidly establish premating RI. This paves the way for the recruitment of late-stage mechanisms, such as adaptive coupling and post-zygotic isolation, to complete speciation.

In this study, we extend the mathematical model from our previous work to investigate how an initial premating RI can couple with various other barrier mechanisms in late-stage sympatric speciation to complete the speciation process. Specifically, we examine the one- and two-allele models of mating-bias barriers, mutations that reduce hybrid viability, chromosomal inversions, and the BDM mechanism of divergence and incompatibilities. These barrier mechanisms are modeled and analyzed using MATLAB applications to determine their invasion dynamics and their ability to couple with the initial barrier and with one another.

We aim to achieve the following objectives in our study: (1) demonstrate that an initial premating barrier can facilitate the coupling of additional adaptive and nonadaptive prezygotic and postzygotic barrier mechanisms to strengthen overall RI and make it irreversible, ultimately completing sympatric speciation; and (2) assess the parameters, ease, and reversibility of how these various barrier mechanisms may be coupled.

The coupling barrier mechanisms investigated in this study are summarized in Fig 26 and Table I. These mechanisms can be categorized as early-stage or late-stage and as adaptive or nonadaptive. Early-stage barriers include the one- and two-allele models of mating-bias and habitat-preference premating barriers, both of which can establish initial premating RI in sympatric populations. Late-stage mechanisms focus on barriers that reduce hybrid viability, chromosomal inversions, and the BDM mechanism of divergence and genetic incompatibilities. Unlike early-stage barriers, these late-stage mechanisms are capable of producing post-mating isolation. The emergence of adaptive barriers is driven by maladaptive hybrid loss. Nonadaptive barriers, such as the BDM mechanism, generate hybrid and mating-bias incompatibilities as chance byproducts of neutral and ecological divergence. More detailed studies of habitat preference barriers and the role of multi-locus ecological and mating-bias genotypes are presented in our separate papers [29, 30].

**Table I.** Summary of different barrier mechanisms.

|  | Early stage | Late stage |  |
| --- | --- | --- | --- |
|  | Mating-bias/<br>habitat preference | Chromosomal Inversion | BDM Incompatibility |
| Mechanism | Adaptive. Driven by selective pressures created by hybrid loss. | Adaptive. Driven by selective pressures created by hybrid loss. | Nonadaptive. Drift and ecological adaptations create hybrid and mating-bias incompatibilities as chance byproducts. |
| Stage of speciation | Early stage. Establish initial pre-mating reproductive barriers | Late stage. CIs are more likely to capture adaptive ecological and mating-bias alleles when there are already niche-specific assortments of those alleles. | Late stage. Ecological adaptations and the development of BDM incompatibilities are more likely without the homogenizing effects of gene flow. |
| Type of RI | Premating. | Pre-mating and post-mating. CIs can capture favorable mating-bias alleles to reduce mating-bias hybrids and increase pre-mating RI. CIs can capture adaptive ecological alleles and mating-bias alleles to increase overall pre-mating RI. CIs can also capture adaptive ecological alleles with alleles that reduce hybrid viability to increase post-mating RI. | Pre-mating and post-mating. The BDM mechanism can produce both pre-mating RI (mating-bias incompatibility) and post-mating RI (reduced ecological hybrid viability) as byproducts of genomic divergence. |
| Speed | Fast. | Fast in late-stage sympatric speciation when a CI is able to capture favorable alleles. But as hybrid loss is reduced, selection pressure for advantageous CIs to invade also decreases. | Slow, especially when incompatibilities are developed through drift. The process is accelerated when mutational incompatibilities are developed through differential adaptations to different niche environments. |
| Reversibility | Easily reversible when disruptive ecological selection against hybrids is reduced. | Reversible when overall RI is weak or intermediate and selection against hybrids is reduced. However, in very late stages of speciation, when RI is very strong, the RI produced by CIs can become irreversible even when disruptive ecological selection is removed. Susceptible to replacement by hybrid swamp. Capturing an $f_{(-)}$ allele that reduce hybrid viability with favorable ecological or mating-bias alleles can create partial irreversibility. | Generally irreversible. It is almost impossible to retrace and reverse the adaptive mutational steps. Therefore, a new species created by the BDM incompatibilities acts like a foreign species that can only be replaced by another fitter species. |
| Reduction of gene flow | Genome-wide. | CIs that capture only locally adaptive ecological alleles can reduce genome-wide RI through negative coupling. However, when CIs capture both locally adaptive ecological alleles and favorable mating bias alleles, they can create genome-wide RI. | Genome-wide. |
| Effect on hybrid viability | None. | CIs capturing an $f_{(-)}$ allele that reduces hybrid viability with favorable ecological alleles or mating-bias alleles can induce decreased hybrid viability. | BDM mechanism produces increased hybrid incompatibility over time. |
| Further divergence | Can be repeatedly invaded by high-mating-bias alleles as long as there is hybrid loss to generate selection pressure for them to invade. | Adaptive alleles within inverted regions are free to further adapt and diverge through the BDM mechanism ("capture and gain"). | Divergence increases with time through drift and differential adaptations. |

In general, our modeling and simulation results confirm that the establishment of an initial premating reproductive barrier can facilitate the coupling of additional barrier mechanisms to create ever stronger and irreversible RI. However, the invasion and coupling dynamics, as well as the conditions under which coupling is feasible, differ according to the distinct properties of the mechanisms involved. The invasion fitness of a specific barrier mechanism is determined by the sum of its inter-niche and intra-niche fitness advantages and disadvantages. Adaptive barriers are readily reversed when disruptive ecological selection is reduced or eliminated. Less reversible mutational changes, such as those in chromosomal inversions and BDM divergence and incompatibilities, are required to achieve stable and lasting RI. Therefore, the role of the adaptive barriers is to provide a rapid and temporary RI for those more permanent late-stage barriers to take hold and mature. Eventually, a positive feedback loop is created, in which early-stage adaptive premating barriers facilitate the coupling of late-stage adaptive and nonadaptive prezygotic and postzygotic barriers. These late-stage barriers, in turn, strengthen and secure the early-stage adaptive barriers, further promoting the establishment of late-stage barriers. This reinforcing cycle ultimately completes the speciation process.

Our previous study proposed a mechanism for how an initial premating RI can be established in a sympatric species undergoing disruptive ecological selection. The current study demonstrates how this initial premating RI can be coupled with additional barrier mechanisms to make the overall RI stronger and irreversible. Together, the results of our two studies present a mechanism of sympatric speciation, from the inception of an initial premating barrier to the completion of the speciation process, in which multiple barriers are coupled in the genomes of sister species to finalize speciation. We hope that our findings will forge a new path forward in the intricate field of speciation research and serve as a framework for further refinement and extension to help solve the mystery of sympatric speciation and restore its role as an important mode of speciation in nature.

## II. METHODOLOGY

Our study uses MATLAB (version R2021a) software. Utilizing the App Designer tool in MATLAB, we designed user-friendly graphical user interface (GUI) applications to solve various mathematical models that depict the coupling of different barrier mechanisms. The results are displayed as phase portraits.

For the initial premating barrier, we use the 2-mating-bias-allele, 2-ecological-niche mathematical model without viable hybrids, which was derived in our previous study (see Fig M3) [28]. We then investigate how other one-allele and two-allele models of pre- and post-zygotic barrier mechanisms can be coupled to this initial barrier.

Fig M1 shows the life cycle of a sympatric population used in our computer simulations. In each generation, individuals in the population undergo ecological selection and sexual selection to produce offspring for the next generation. Two separate niche resources, *A* and *B*, created by disruptive ecological selection, exist in the sympatric ecosystem. In the ecological selection phase of the life cycle, only individuals with genotypes adaptive to the two niches (i.e., ecotype *A* or ecotype *B*) are able to extract niche resources and survive. The flowchart in Fig M2 describes the sexual selection phase of the life cycle. Eligible mating individuals encounter each other randomly in a matching round. In each encounter, their matching success is determined by two mating-bias alleles, *X* or *Y*, that exist at a single gene locus. Fig M4 shows a matching compatibility table for the mating-bias alleles. When two individuals possess the same mating-bias alleles, they are 100% compatible and always match. When they possess different mating-bias alleles, their matching success is determined by a mating-bias value, *α*, that varies between 0 and 1. Unmatched individuals after a matching round can try to match again in the next round, up to a total of *n* matching rounds. After *n* matching rounds, all unmatched individuals die without leaving offspring, while all matched individuals reproduce offspring by random assortment of their alleles at each gene locus. The value of *n* is therefore a measure of the cost of assortative mating.

**Fig M1.**
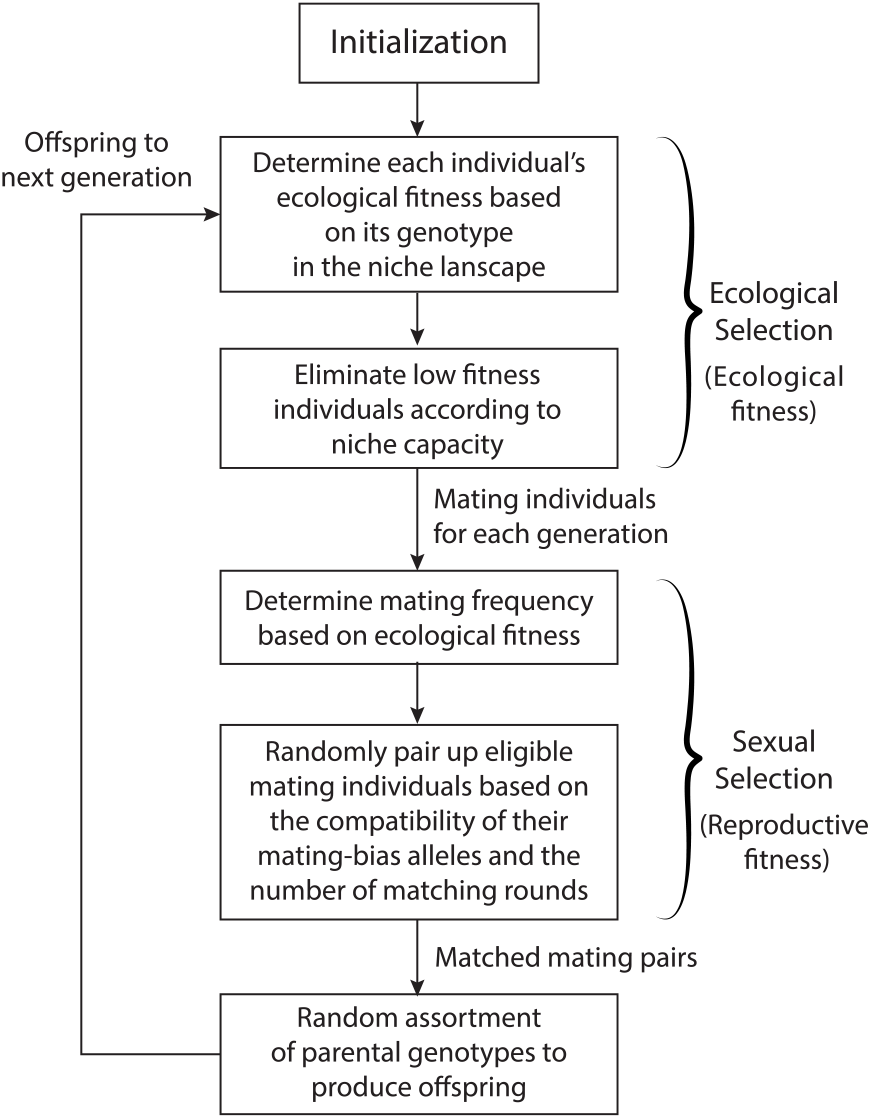
A flowchart showing the life cycle of a sympatric population used in the computer simulations.

**Fig M2.**
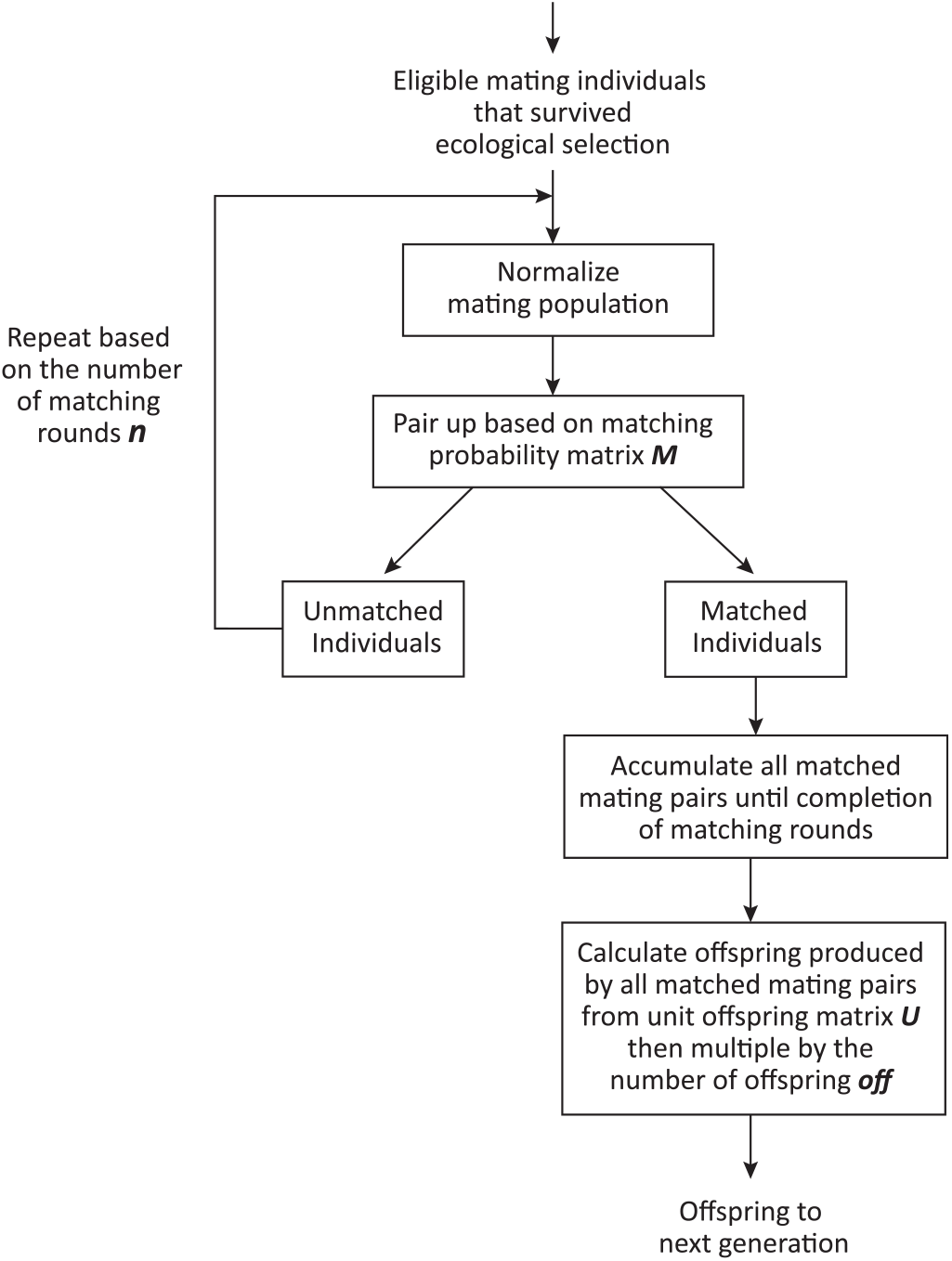
A flowchart showing the algorithm used to match eligible mating individuals and reproduce offspring.

**Fig M3.**
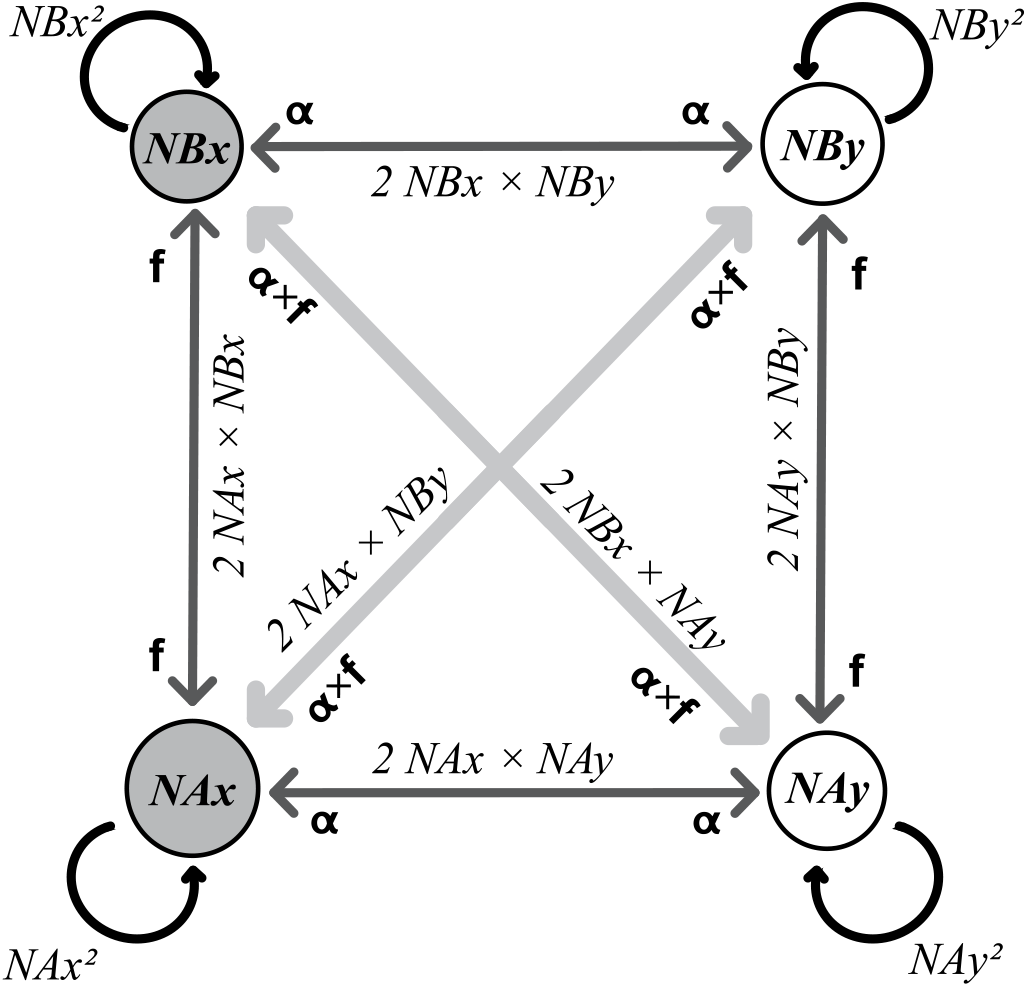
Mathematical model of a two-mating-bias-allele, two-ecological-niche sympatric ecosystem.

**Fig M4.**
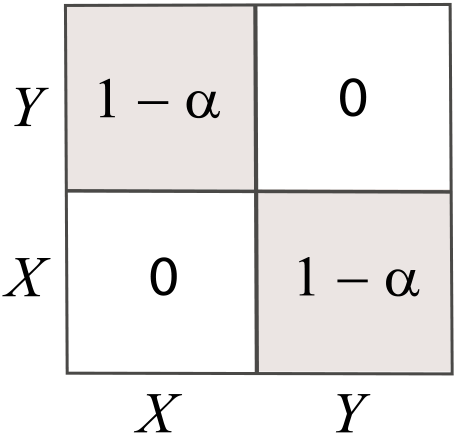
A matching compatibility table for mating-bias alleles.

**Fig M5.**
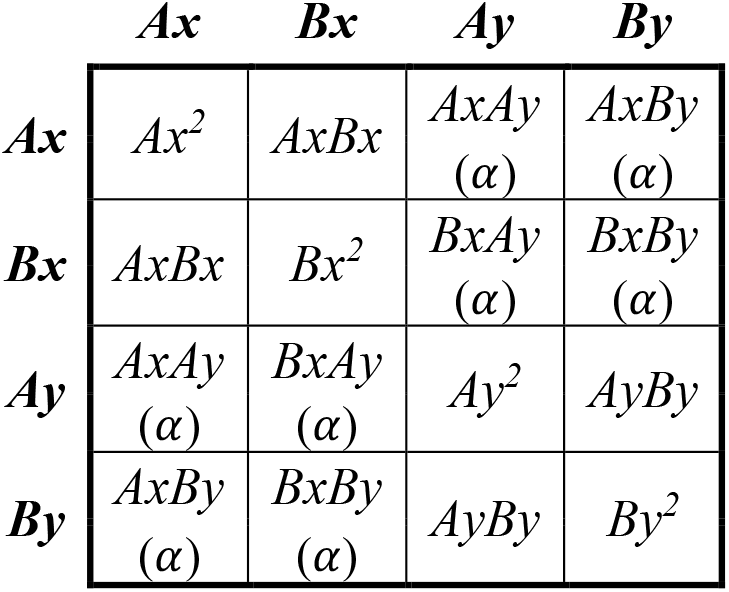
A matching probability matrix *M* for a sympatric model with no viable hybrids.

**Fig M6.**
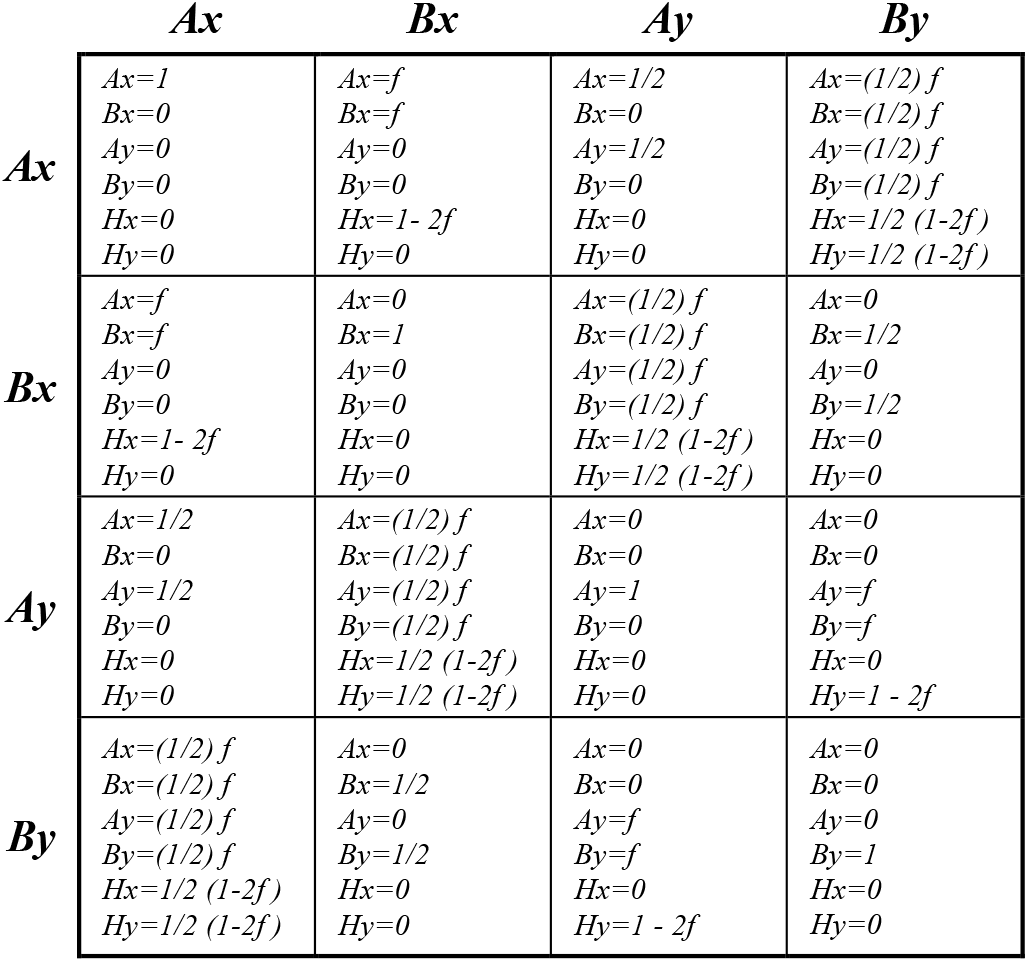
Unit offspring matrix *U*.

Fig M3 shows the 2-mating-bias-allele, 2-niche mathematical model that describes the population dynamics of the sympatric population during a matching round. It consists of four genotype groups: *NAx, NAy, NBx*, and *NBy*, which represent the population ratios of the different ecotypes (*A* or *B*) and their associated mating-bias alleles (*X* or *Y*). All the parametric values in the model are normalized (i.e., *NAx*+*NAy*+*NBx*+*NBy* = 1).

The four genotype groups in Fig M3 encounter one another randomly. Fig M5 shows a matching probability matrix that specifies the probabilities of encounters between the different groups. Note that the elements of the matrix add up to 1. The variable *α* is also displayed to remind us that the matching success between individuals possessing different mating-bias alleles is determined by the value of *α*. The values of *α* and *n* determine the strength of sexual selection.

After *n* matching rounds, all successfully matched genotype pairs are collected. These parental pairs reproduce offspring by random assortment of alleles at their gene loci. The Unit Offspring Matrix in Fig M6 shows the ratios of the different offspring genotypes produced by the different parental genotypes, assuming that each parent can produce only one offspring to replace itself (i.e., reproducing a “unit offspring”). The variable *f* is the *“*offspring return ratio*”* that specifies the proportion of offspring with the same genotypes as their parents. The value of *f* ranges from 0 to 0.5. For instance, a genotype with three gene loci will have an *f* value of 1/8, while 6/8 of the offspring from inter-niche matings are hybrids.

Although our model assumes no viable hybrid offspring, the presence of viable hybrids due to hybrid niche resources can still be accounted for by increasing the value of *f*. In the model, the value of *f* represents the strength of ecological selection. Note that all the offspring ratios in each cell of the Unit Offspring Matrix add up to 1. To obtain the final offspring populations, we can simply multiply the unit offspring ratios by a fertility ratio, *off*, which specifies the number of offspring each parent actually produces.

To counter the effect of “incumbent selection” [28], our mathematical model assumes that in each mating generation, the ecotype in each niche reproduces enough offspring to completely fill the niche’s carrying capacity. Therefore, the normalized niche population ratios, *NA* and *NB*, are always fixed at the start of the sexual selection phase in each generation.

Our GUI computer application then plots the *Ax*/*Bx* phase portrait solutions of the model based on user-specified parametric values. Here, *Ax* is the normalized ratio of the niche-*A* ecotypes that have the *X* allele, and *Ay* is similarly defined, so that *Ax* + *Ay* = 1. The same definitions apply for *Bx* and *By*. The phase portrait then displays the change in normalized genotype ratios over *g* generations.

A system’s phase-portrait solutions are either “convergent,” meaning its vector field converges to fixed points, or “divergent,” meaning it contains no fixed points. If we let *NAx* be the largest genotype group in the model, then to have maximum RI, we want *NAx* and *NBy* to be large and *NAy* and *NBx* to be small. This is achieved by having a fixed point on the *Ax*/*Bx* phase portrait as close to the lower right corner (where *Ax* = 1 and *Bx* = 0) as possible.

All the GUI applications in this study follow the same algorithms as outlined in Fig M1 and Fig M2. Coupling additional barrier mechanisms to the initial premating RI involves simply modifying or expanding the matrices in Fig M4, M5, and M6. The modified matrices for each GUI application are listed in the Appendix.

To examine the invasion and coupling dynamics of various barrier mechanisms, our general modeling approach involves specifying a mutant allele with properties characteristic of different types of barriers. We then vary the parametric values in the model to identify the conditions under which the mutant allele can successfully invade and proliferate in the presence of other barrier systems. Successful invasion corresponds to successful coupling, and the resulting changes in population dynamics and overall RI are calculated and displayed in the model’s phase portraits.

## III. RESULTS

### 1. Coupling of Early -Stage, Premating Barriers

MATLAB Graphical User Interface (GUI) applications were developed to investigate how an initial premating mating-bias barrier may be coupled with other one- and two-allele models of mating-bias barriers to increase the overall premating RI.

#### 1.1. Coupling with one-allele mechanisms of mating-bias barriers

Figs 1a and 1b show that a one-allele mechanism of mating-bias barrier can be coupled to a pre-existing 2-allele mechanism of premating RI to further strengthen the overall RI between niche ecotypes. Here, a mutant allele *K* at a separate gene locus decreases the mating-bias value of *α*, changing *α* to *αk*, and increases the mating barrier between genotypes carrying different mating-bias alleles, *X* and *Y*. As shown in Fig 1a, a mutant allele *K* with an initial population ratio of 0.01 is able to invade and become fixed in niche-*A* and niche-*B* ecotypes after 3000 generations, reduce the hybrid offspring ratio from 0.22128 to 0.16200, and strengthen the overall RI. Because the allele *K* operates by a one-allele mechanism—i.e., a single allele has effect in both niche ecotypes—it is immune to the homogenizing effect of recombination due to gene flow. In a convergent system, the mutant allele *K* can always invade if *n* > 1 and *αk* < *α*.

**Fig 1a.**
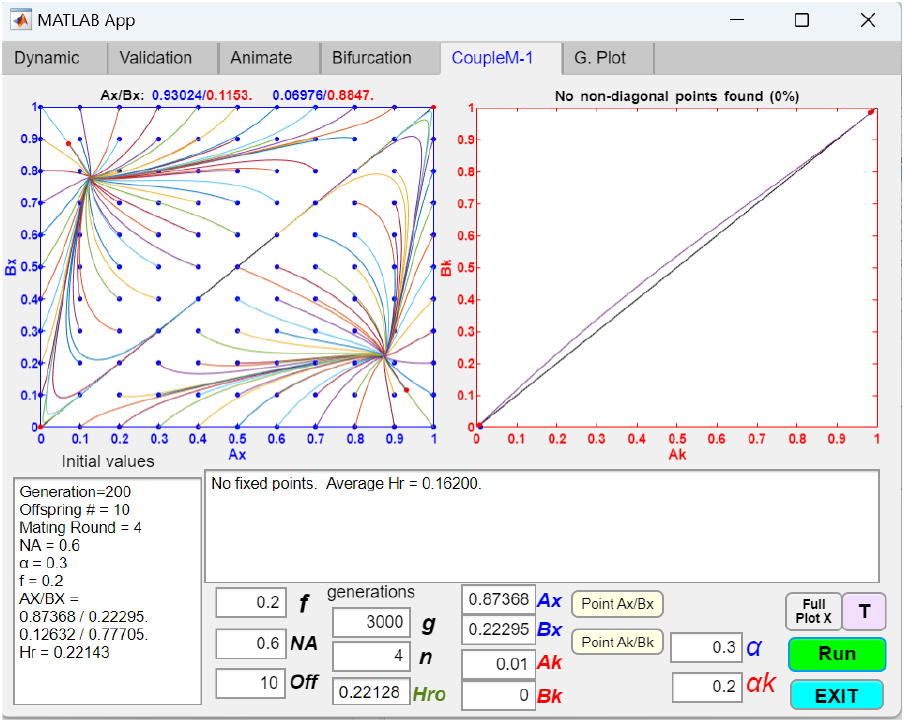
Coupling of an initial two-allele premating barrier with a one-allele barrier mechanism that increases its mating bias. A computer screenshot of a MATLAB GUI application displays the phase portrait solutions of a two-allele barrier system (*Ax*/*Bx*) coupled with a one-allele barrier system (*Ak*/*Bk*). An initial two-allele, single-gene-locus premating barrier mechanism has attained a fixed-point polymorphism at *Ax* = 0.87368 and *Bx* = 0.22295, given the initial parametric values shown in the table insert. The mating bias value of the *Ax*/*Bx* barrier system is *α* = 0.3, while *αk* = 0.2 is the new mating bias value introduced by the presence of a mutant *k* allele. *Ak* represents the ratio of the *k* allele in niche *A*, and *Bk* represents the ratio of the *k* allele in niche *B*. The rest of the parametric variables are the same as those described in the Methodology section. The phase portraits show that a starting mutant genotype population of *Ak* = 0.01 can invade any starting populations of *Ax*/*Bx* (shown as blue dots) and become fixed in niche *A* and niche *B* (i.e., *Ak* = *Bk* = 1). Consequently, the coupling of these two barrier systems shifts the initial *Ax*/*Bx* fixed point to a new fixed point, *Ax* = 0.93024 and *Bx* = 0.1153 (shown as a red dot), which is closer to the lower right corner in the phase portrait, resulting in stronger overall premating RI. *Hro* = 0.22128 is the hybrid offspring ratio before the invasion of the *k* allele, and *Hr* = 0.162 is the hybrid offspring ratio after the *k* allele is fixed. The reduction in the overall hybrid offspring ratio reflects the increase in premating RI as a result of the coupling of the two barriers. The matching probability table and the unit-offspring table used for this GUI application are included in the Appendix (Tables 1a and 1b).

**Fig 1b.**
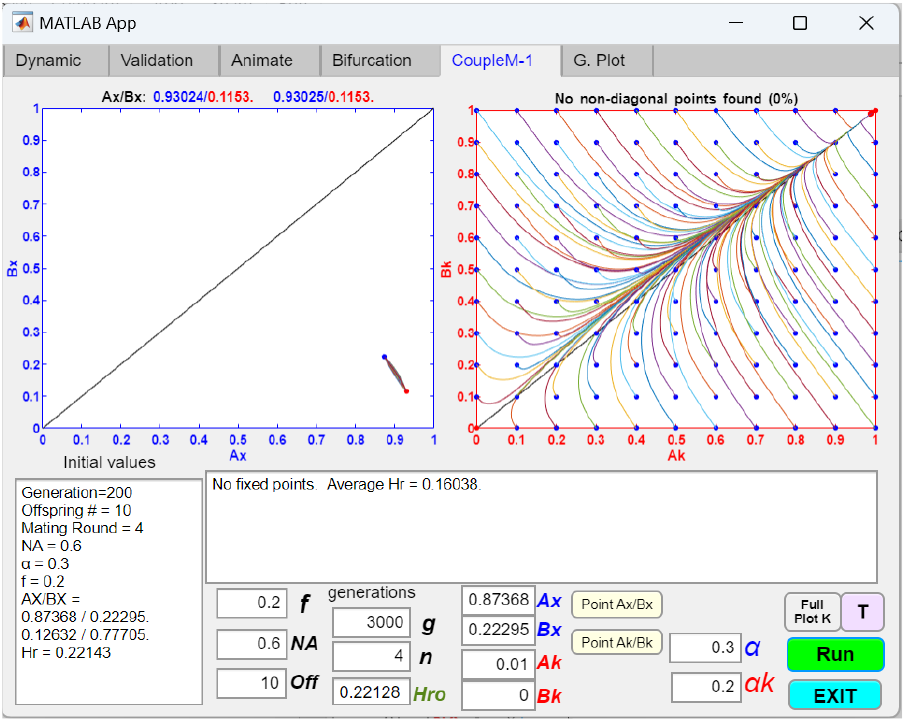
Coupling of an initial two-allele barrier with a one-allele barrier, showing the phase portrait of the invading allele. Using the same parametric values as in Fig 1a, a computer screenshot shows the phase portrait solution of *Ak*/*Bk*, given the initial *Ax*/*Bx* fixed point. As shown, all initial values of *Ak*/*Bk* converge to fixation, which shifts the final *Ax*/*Bx* fixed point closer to the lower right corner coordinate of *Ax* = 1/*Bx* = 0, creating stronger premating RI.

Fig 1b describes the same population dynamics in Fig 1a but viewed using different plots. It shows the phase portrait of *Ak*/*Bk* given the initial premating *Ax*/*Bx* fixed-point polymorphism (shown as the blue dot in the *Ax*/*Bx* phase portrait). The phase portrait of *Ak*/*Bk* shows a globally convergent pattern. Any mutation arising from the origin will be drawn to the fixed point at *Ak* = 1 and *Bk* = 1. In the process, the fixed point in the *Ax*/*Bx* phase portrait is moved closer toward the lower right corner where *Ax* = 1 and *Bx* = 0 (the new fixed point is shown as the red dot in the phase portrait), creating greater premating RI as a result of the coupling.

**Table 1a.** Matching table for coupling to a one-allele barrier mechanism that increases mating bias. The table shows the encounter probabilities of the different genotypes, *h*1 to *h*4, in a sympatric population. An individual possessing the *XK* genotype has the *X* allele and the *K* allele in its genotype. When genotypes with different mating-bias alleles, *X* or *Y*, meet, their matching success is determined by mating bias *α*, except when the *K* allele is present in the genotypes. In that case, the mating bias is changed from *α* to *αk* by the *K* allele. The presence of the *J* allele, denoting the absence of the K allele, has no effect.

VIII. APPENDIX
| | XJ $h1$ | YJ $h2$ | XK $h3$ | YK $h4$ |
| --- | --- | --- | --- | --- |
| XJ $h1$ | X J<br>X J<br>1 | X J<br>Y J<br>$\alpha$ | X J<br>X K<br>1 | X J<br>Y K<br>$\alpha k$ |
| YJ $h2$ | Y J<br>X J<br>$\alpha$ | Y J<br>Y J<br>1 | Y J<br>X K<br>$\alpha k$ | Y J<br>Y K<br>1 |
| XK $h3$ | X K<br>X J<br>1 | X K<br>Y J<br>$\alpha k$ | X K<br>X K<br>1 | X K<br>Y K<br>$\alpha k$ |
| YK $h4$ | Y K<br>X J<br>$\alpha k$ | Y K<br>Y J<br>1 | Y K<br>X K<br>$\alpha k$ | Y K<br>Y K<br>1 |

**Table 1b.** Unit offspring table for coupling to a one-allele barrier mechanism that increases mating bias. Each cell of the table shows the offspring genotype ratios produced by the different parental genotypes. The parental genotypes are shown in the vertical and horizontal headings. The offspring genotypes are displayed in each cell along with their fractional ratios in parentheses. Note that all the offspring ratios in a cell add up to 1.

| | XJ $h1$ | YJ $h2$ | XK $h3$ | YK $h4$ |
| --- | --- | --- | --- | --- |
| XJ $h1$ | XJ $h1$ (1) | XJ $h1$ (1/2)<br>YJ $h2$ (1/2) | XJ $h1$ (1/2)<br>XK $h3$ (1/2) | XJ $h1$ (1/4)<br>YJ $h2$ (1/4)<br>XK $h3$ (1/4)<br>YK $h4$ (1/4) |
| YJ $h2$ | XJ $h1$ (1/2)<br>YJ $h2$ (1/2) | YJ $h2$ (1) | XJ $h1$ (1/4)<br>YJ $h2$ (1/4)<br>XK $h3$ (1/4)<br>YK $h4$ (1/4) | YJ $h2$ (1/2)<br>YK $h4$ (1/2) |
| XK $h3$ | XJ $h1$ (1/2)<br>XK $h3$ (1/2) | XJ $h1$ (1/4)<br>YJ $h2$ (1/4)<br>XK $h3$ (1/4)<br>YK $h4$ (1/4) | XK $h3$ (1) | XK $h3$ (1/2)<br>YK $h4$ (1/2) |
| YK $h4$ | XJ $h1$ (1/4)<br>YJ $h2$ (1/4)<br>XK $h3$ (1/4)<br>YK $h4$ (1/4) | YJ $h2$ (1/2)<br>YK $h4$ (1/2) | XK $h3$ (1/2)<br>YK $h4$ (1/2) | YK $h4$ (1) |

In the Fig 2a example, the system is initially divergent, i.e., no *Ax*/*Bx* fixed-point polymorphism is possible in the phase portrait because the value of *f* = 0.3 is too high for system convergence. Nevertheless, the phase portrait solution of *Ak*/*Bk* revealed that it is still possible for a high mating-bias allele *K* to convert such an initially divergent system into a convergent one through coupling and create premating RI. However, to achieve this typically requires lower mating-bias values of *αk*, larger starting *Ak* and *Bk* populations, or greater numbers of matching rounds *n* than are required for comparable convergent systems.

**Fig 2a.**
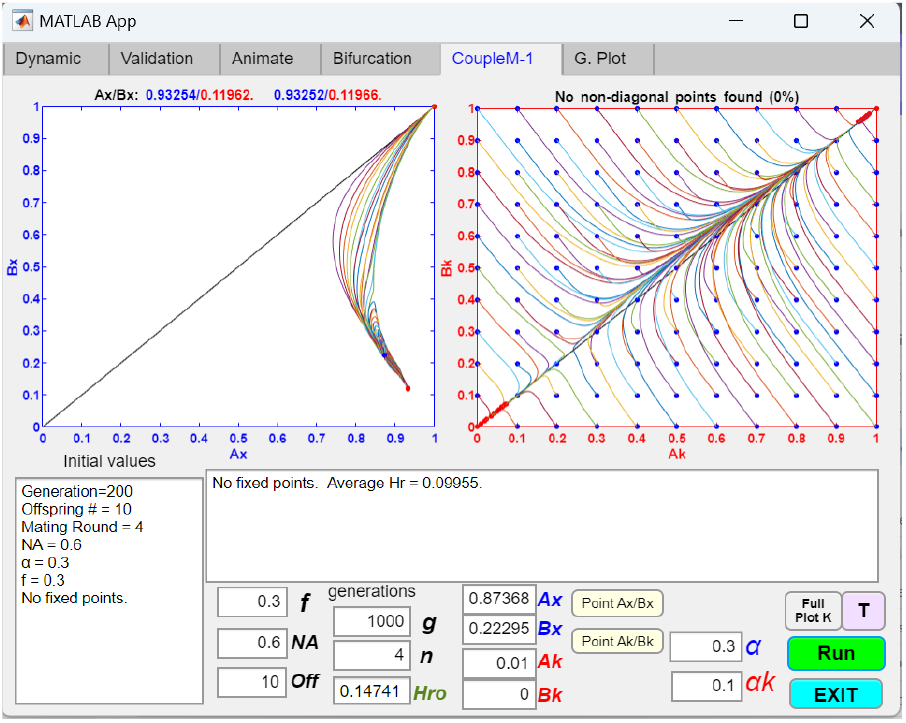
Coupling of an initially divergent two-allele barrier system with a one-allele barrier mechanism to achieve premating RI. The *Ak*/*Bk* phase portrait shows an invasion resistant pattern. Starting *Ak*/*Bk* populations (e.g., *Ak* = 0.01/ *Bk* = 0) near the origin cannot invade. Higher starting population ratios of *Ak* and *Bk* are needed for the *K* allele to invade and become fixed in both niches.

**Fig 2b.**
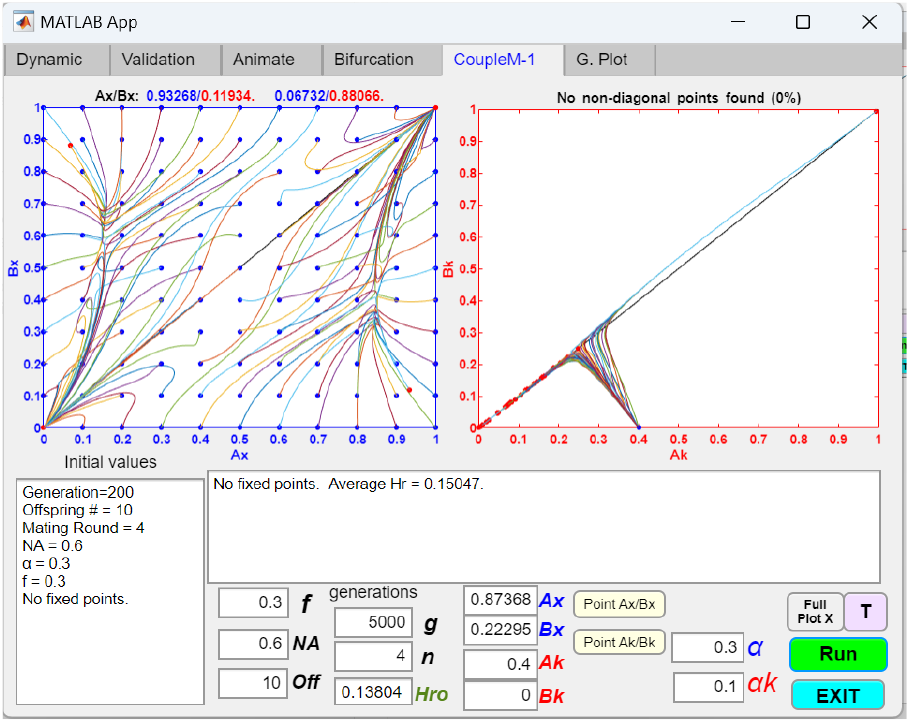
Larger *Ak* values can cause *Ax*/*Bx* populations near the lower right corner of the Ax/Bx phase portrait to converge to a fixed point. The same parametric values as in Fig 2b are used, except *Ak* = 0.4.

For instance, in the Fig 2a example, starting with the *Ax*/*Bx* fixed-point values in Fig 1a and *f* = 0.3, an *Ak* mutant can invade only if the initial *Ak* ≥ 0.352. If *αk* is reduced from 0.1 to 0.05, invasion occurs if the initial *Ak* ≥ 0.254. If *n* is increased from 4 to 10, invasion occurs if the initial *Ak* ≥ 0.09. Fig 2b shows that with an initial *Ak* value of 0.4, the *Ax*/*Bx* phase portrait is partially convergent. Fixed point polymorphism only occurs when the difference between *Ax* and *Bx* starting populations is large—i.e., with starting populations near the lower right corner in the phase portrait. If the initial *Ak* = 0.4 and *n* = 10, the *Ax*/*Bx* phase portrait becomes globally convergent.

In general, it is easier to strengthen the premating RI in a convergent system through coupling than to convert a divergent system without fixed points into a convergent system with fixed points. These results are consistent with our previous findings [28].

#### 1.2. Coupling with two-allele mechanisms of mating-bias barriers

Fig 3a shows that a pre-existing 2-allele mechanism of premating RI can be coupled with another two-allele mechanism of mating-bias barrier to further strengthen the overall RI. Here, mating-bias alleles *Z* and *W* exist at a separate gene locus. Similar to the 2-allele model for the *X* and *Y* alleles, the matching success of genotype pairs possessing different *Z* and *W* alleles (e.g., *Ax* and *Bw, Aw* and *Ax*, etc.) is determined by the mating-bias variable *β*. The overall mating bias between genotypes possessing different mating bias alleles at the two gene loci, e.g., *Axx* and *Byw*, is the product of *α* and *β*, i.e., *α* × *β* (refer to the mating-bias table, Table 2a, in the Appendix).

**Table 2a.** Matching table for coupling to a two-allele barrier mechanism with mating bias *β*. The table shows the encounter probabilities of the different genotypes, *h*1 to *h*4, in a sympatric population. The first two-allele system has mating-bias alleles, *X* and *Y*, and mating bias *α*. The second two-allele system has matingbias alleles, *Z* and *W*, and mating bias *β*. When two genotypes have different alleles from the two systems, their matching success is determined by the product of the mating biases *α* × *β*.

| | XZ $h1$ | XW $h2$ | YZ $h3$ | YW $h4$ |
| --- | --- | --- | --- | --- |
| XZ $h1$ | X Z<br>X Z<br>1 | X Z<br>X W<br>$\beta$ | X Z<br>Y Z<br>$\alpha$ | X Z<br>Y W<br>$\alpha\beta$ |
| XW $h2$ | X W<br>X Z<br>$\beta$ | X W<br>X W<br>1 | X W<br>Y Z<br>$\alpha\beta$ | X W<br>Y W<br>$\alpha$ |
| YZ $h3$ | Y Z<br>X Z<br>$\alpha$ | Y Z<br>X W<br>$\alpha\beta$ | Y Z<br>Y Z<br>1 | Y Z<br>Y W<br>$\beta$ |
| YW $h4$ | Y W<br>X Z<br>$\alpha\beta$ | Y W<br>X W<br>$\alpha$ | Y W<br>Y Z<br>$\beta$ | Y W<br>Y W<br>1 |

**Table 2b.**
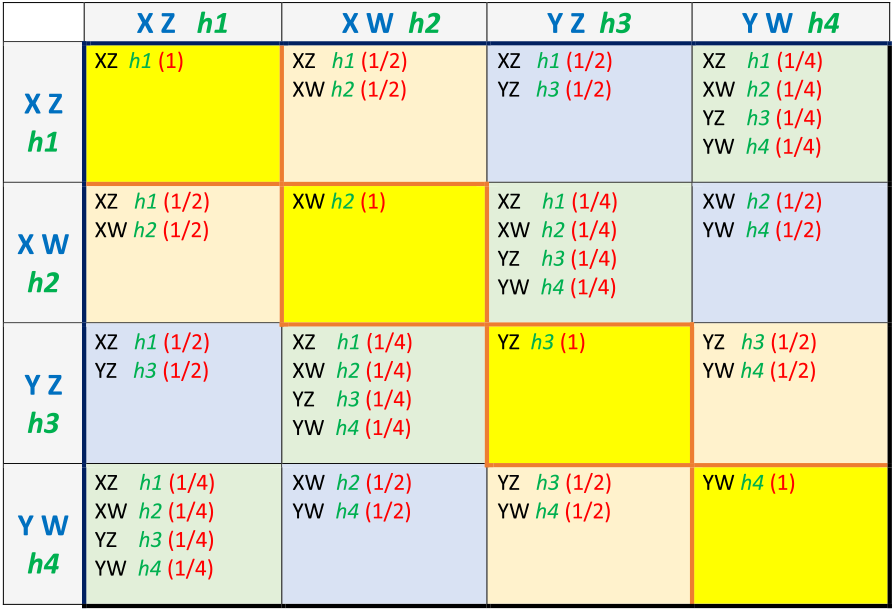
Unit offspring table for coupling to a two-allele barrier mechanism with mating bias *β*.

| | XZ $h1$ | XW $h2$ | YZ $h3$ | YW $h4$ |
| --- | --- | --- | --- | --- |
| XZ $h1$ | XZ $h1$ (1) | XZ $h1$ (1/2)<br>XW $h2$ (1/2) | XZ $h1$ (1/2)<br>YZ $h3$ (1/2) | XZ $h1$ (1/4)<br>XW $h2$ (1/4)<br>YZ $h3$ (1/4)<br>YW $h4$ (1/4) |
| XW $h2$ | XZ $h1$ (1/2)<br>XW $h2$ (1/2) | XW $h2$ (1) | XZ $h1$ (1/4)<br>XW $h2$ (1/4)<br>YZ $h3$ (1/4)<br>YW $h4$ (1/4) | XW $h2$ (1/2)<br>YW $h4$ (1/2) |
| YZ $h3$ | XZ $h1$ (1/2)<br>YZ $h3$ (1/2) | XZ $h1$ (1/4)<br>XW $h2$ (1/4)<br>YZ $h3$ (1/4)<br>YW $h4$ (1/4) | YZ $h3$ (1) | YZ $h3$ (1/2)<br>YW $h4$ (1/2) |
| YW $h4$ | XZ $h1$ (1/4)<br>XW $h2$ (1/4)<br>YZ $h3$ (1/4)<br>YW $h4$ (1/4) | XW $h2$ (1/2)<br>YW $h4$ (1/2) | YZ $h3$ (1/2)<br>YW $h4$ (1/2) | YW $h4$ (1) |

**Fig 3a.**
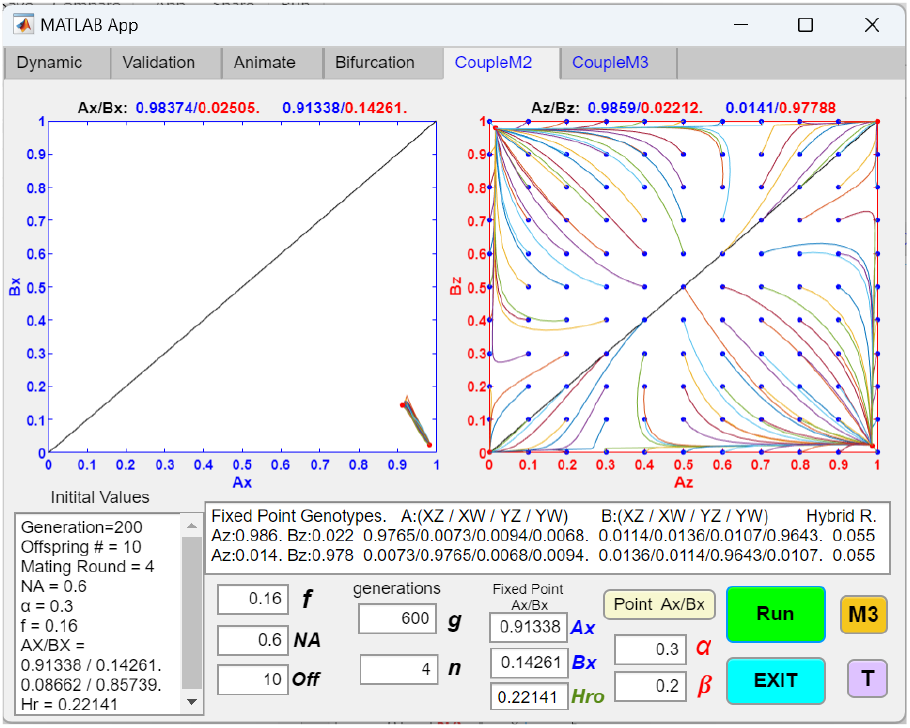
Coupling of an initially convergent two-allele barrier system with another two-allele barrier system. An initial two-allele, single-gene-locus premating barrier mechanism has attained a fixed-point polymorphism at *Ax* = 0.91338 and *Bx* = 0.14261, given the initial parametric values shown in the table insert. This first barrier system is then coupled to another two-allele, single-gene-locus barrier system with mating-bias alleles, *Z* and *W*, and a mating-bias value *β*. The computer screenshot shows the *Ax*/*Bx* phase portrait solution given the *Ax*/*Bx* fixed-point values. As a result of the coupling, the Initial *Ax*/*Bx* fixed point is shifted to *Ax*/*Bx* = 0.98374/0.02505, and the hybrid offspring ratio is decreased from 0.22141 to 0.055, signifying increased overall premating RI. Note that the vector field in the *Ak*/*Bk* phase portrait exhibits an “invasion resistant” pattern, meaning that mutants with low initial *Ax*/*Bx* population ratios around the origin are drawn back toward the origin and cannot invade. The matching and unit-offspring tables associated with this GUI application are included in the Appendix (Tables 2a and 2b).

**Fig 3b.**
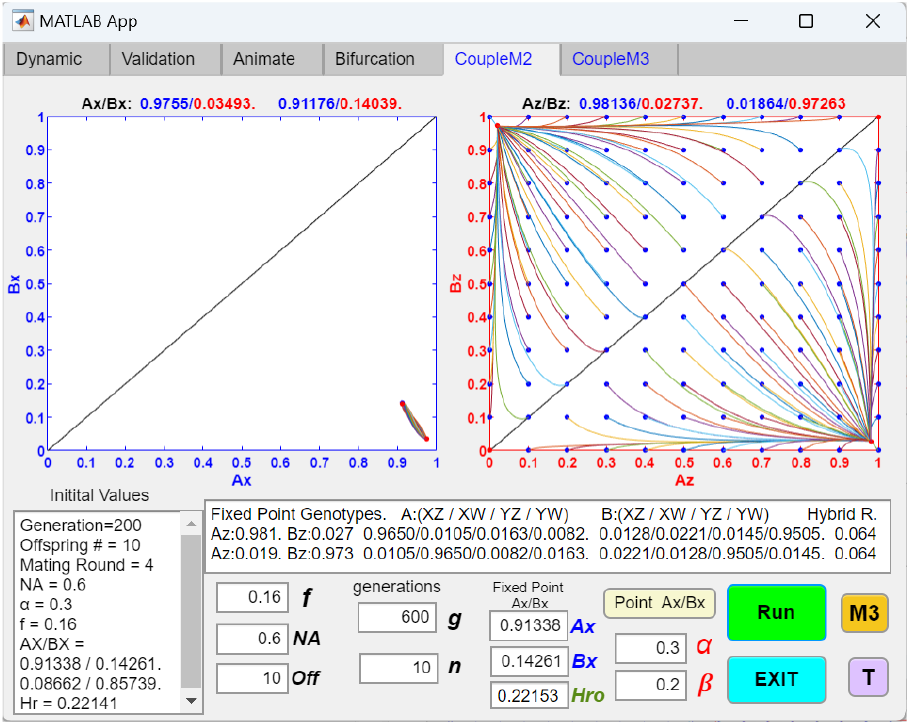
Disappearance of the “invasion resistance” patten when the number of matching rounds, *n*, is increased. Increasing the number of matching rounds *n* from 4 to 10 in Fig 3a, while keeping the rest of the parametric values the same, eliminates the “invasion-resistant” pattern in the *Ax*/*Bx* phase portrait. The system becomes globally convergent, allowing rare mutants to invade and achieve fixed-point polymorphism.

**Fig 3c.**
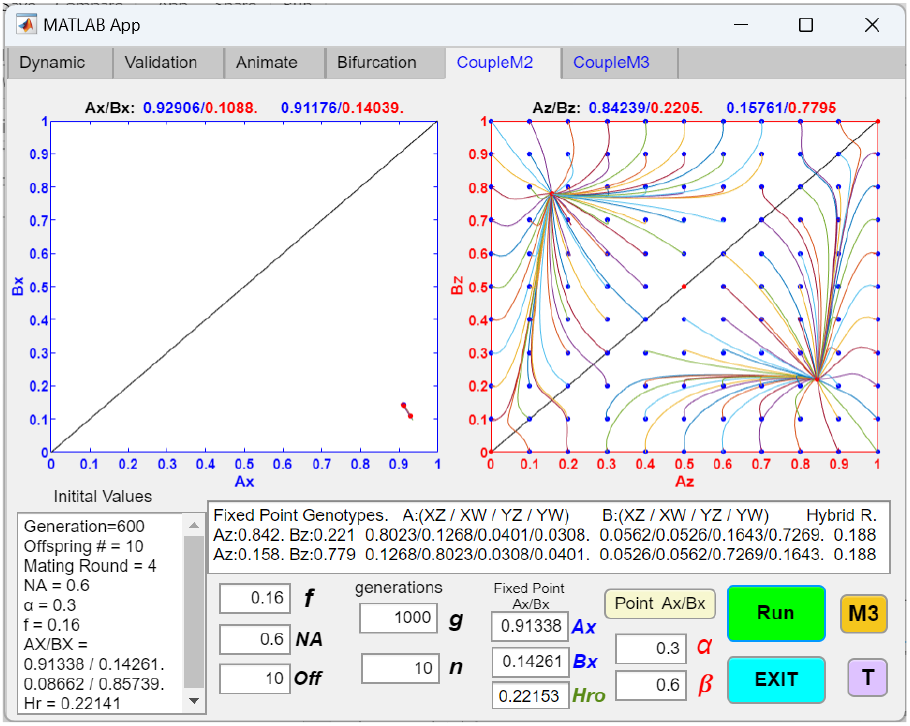
Coupling to a two-allele barrier system can still occur with a high value of mating bias *β*. When the value of *β* in Fig 3b is increased from 0.2 to 0.6 while keeping the rest of the parametric values the same, the *Ax*/*Bx* phase portrait remains globally convergent. However, this coupling with a weaker mating-bias system results in a smaller increase in premating RI.

Fig 3a shows that, after the two-allele mechanism at the *X*/*Y* gene locus established an initial fixed-point polymorphism (*Ax* = 0.91338 and *Bx* = 0.14261), it can be coupled to another two-allele barrier mechanism at the *Z*/*W* gene locus to increase the overall premating RI. As a result, fixed-point polymorphism emerges in the *Ax*/*Bx* phase portrait, the initial *Ax*/*Bx* fixed point is shifted toward the lower right corner (*Ax*/*Bx* = 0.98374/0.02505), and the hybrid offspring ratio is reduced from 0.22141 to 0.055, signifying greater overall RI.

However, the *Ax*/*Bx* phase portrait in Fig 3a cannot be invaded by an initially small population of *Ax* or *Bx* mutants. This is because the origin (where *Ax* = 0 and *Bx* = 0) does not lie in the basins of attraction of the fixed points. Consequently, starting *Ax*/*Bx* populations around the origin converge to the origin rather than to the fixed points. (Similar observations can be made about populations near the corner opposite the origin, i.e., where *Ax* = 1 and *Bx* = 1 or where *Aw* = 0 and *Bw* = 0). We can describe such a phase portrait as exhibiting an *“*invasion resistant pattern.” We seem to observe such a pattern commonly in our simulations. It tends to occur when gene flow and hybrid loss are already significantly reduced before coupling with the additional two-allele model of mating-bias system. In most instances, if we increase the number of matching rounds *n*, thereby reducing the assortative mating cost, such a pattern can be eliminated. Fig 3b shows that if *n* is increased from 4 to 10, while keeping the rest of the parametric values the same as in Fig 3a, the invasion-resistant pattern is eliminated, and the *Ax*/*Bx* phase portrait becomes globally convergent, allowing the invasion of rare mutants. Such a change typically incurs only a minimal reduction in the strength of the barriers. In the Fig 3b example, the hybrid ratio is only slightly increased from 0.055 to 0.064 as a result of the change.

Alternatively, in the presence of an invasion-resistant pattern, reducing the ratio of *NA* facilitates the invasion of a mutant *Z* allele arising in niche *A* (e.g., *Ax* = 0.01), while the presence of the *Z* allele in niche *B* (i.e., *Bx* > 0) reduces the invasion fitness of the *Ax* mutant genotype. In general, decreasing the niche size *NA* increases the fitness advantage of *Ax* due to increased inter-niche mating, allowing the *Ax* mutant to invade more quickly (i.e., requiring fewer generations to achieve fixed-point polymorphism), but this tends to result in weaker RI (with the fixed point located further away from the bottom-right corner of the phase portrait). Conversely, when the sizes of niches *A* and *B* are roughly equal, it takes longer for the *Ax* mutant to reach fixed-point polymorphism, but the fixed point tends to lie closer to the bottom-right corner, resulting in stronger RI.

Fig 3c shows the effect of changing the value of *β* in Fig 3b from 0.2 to 0.6. Coupling still occurs, and the overall hybrid ratio decreases from 0.22153 to 0.188, signifying a stronger overall premating RI. This is despite the fact that, on its own, with the parametric values *f* = 0.16, *NA* = 0.6 and *n* = 10, a mating bias value of 0.6 is not low enough to cause the system to converge and achieve fixed-point polymorphism. Therefore, by coupling or hitchhiking with other barriers, a weak barrier mechanism can nonetheless gain a disproportionately greater selective advantage to invade and multiply.

In Fig 4, the two coupled systems in Fig 3c are again coupled with a third two-allele premating barrier with a mating-bias value *γ* = 0.2, further reducing the overall hybrid ratio to 0.042. This occurs because genotypes from different niches, possessing different mating-bias alleles (e.g., genotypes *Axzj* and *Bywk*), experience the maximum mating bias between them, calculated by multiplying all three mating-bias values *α* × *β* × *γ*. As a result, such genotype pairs have the greatest fitness compared to other genotype pairs with allele combinations that result in lower overall mating biases.

**Fig 4.**
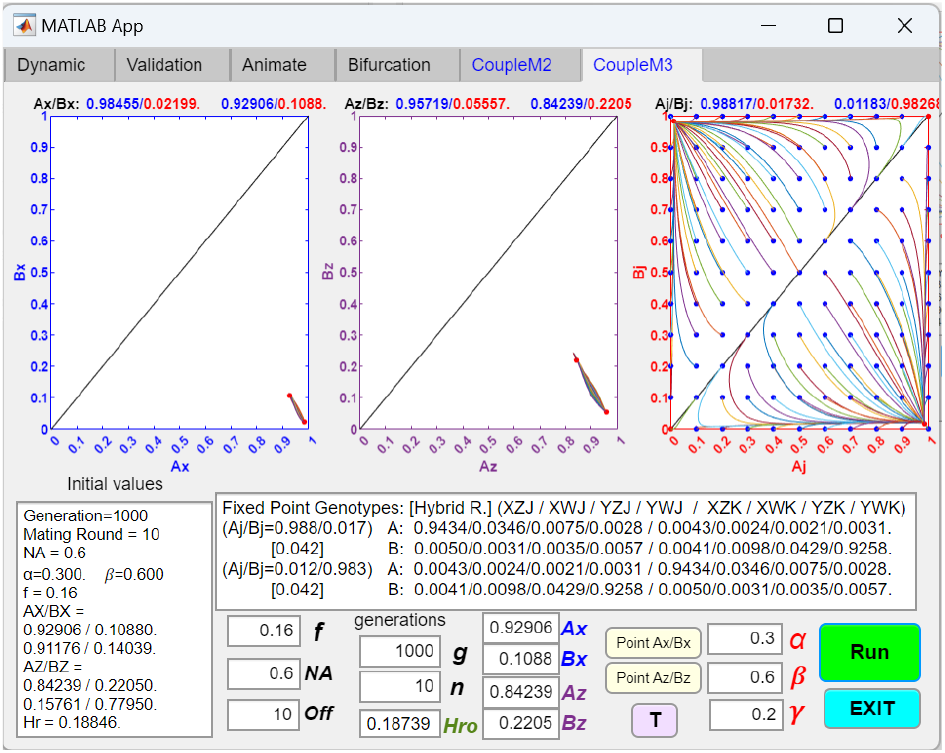
Coupling of three one-gene-locus, two-allele mating-bias barrier systems. The two-allele mating-bias systems in Fig 3c are coupled to a third two-allele system. The third system has two mating-bias alleles, J and K, with a mating bias *γ*. The phase portrait solutions of the three systems show that coupling to a third system further increases the premating RI of the two coupled systems in Fig 3c. The matching and unit-offspring tables for the GUI application is included in the Appendix (Tables 3a and 3b).

Fig 5 shows what happens if we change the order of coupling. In contrast to the scenario in Fig 4, in Fig 5, the initial premating barrier is coupled to a second two-allele barrier with a strong mating bias *β* = 0.2 and then to a third two-allele barrier with a weak mating bias *γ* = 0.6. Even though the maximum overall mating-bias possessed by the genotypes in Fig 4 and Fig 5 is the same, i.e., *α* × *β* × *γ* = 0.036, the phase portrait of the third barrier displays an invasion-resistant pattern that makes invasion by a small population of mutants impossible. This invasion-resistant pattern is not eliminated despite increasing *n* to 100. Thus, it appears that changing the order of how different barrier mechanisms are coupled makes a difference in the final outcome. The result in Fig 5 can be explained by the fact that adaptive coupling is driven by selection pressures arising from hybrid loss. When two coupled strong barriers have already established strong RI and drastically reduced hybrid loss, there is not much driving force or positive selection pressure left for a third, weak barrier to couple effectively and further reduce overall RI.

**Table 3a.**
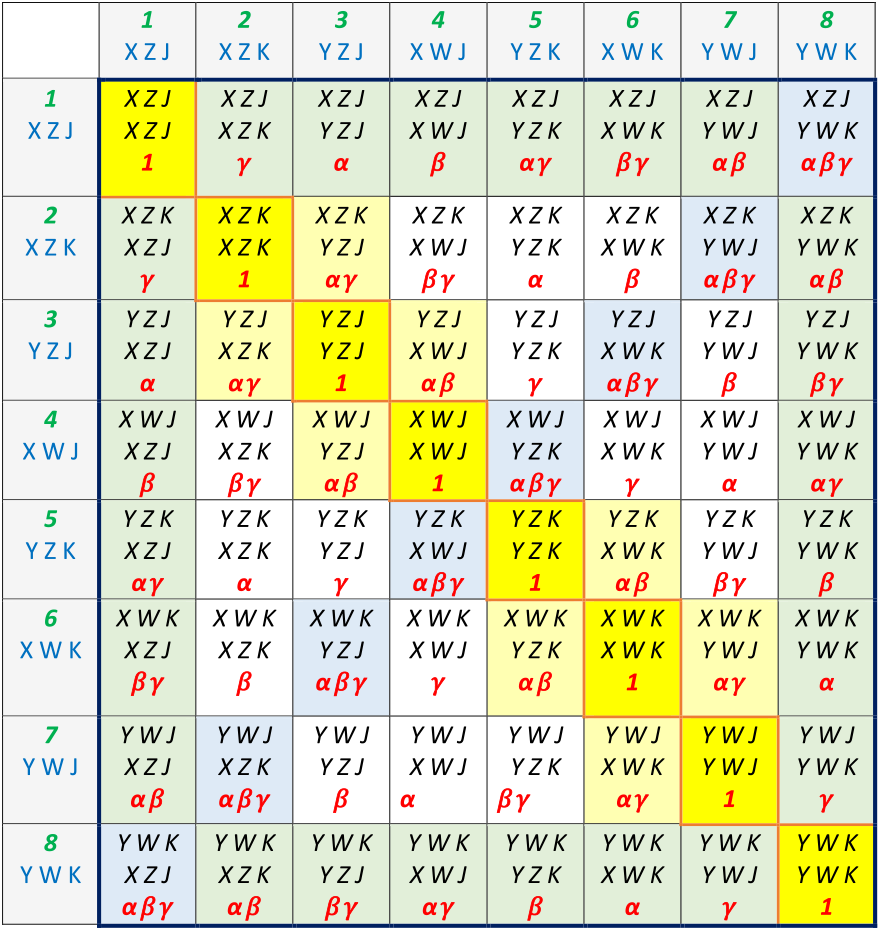
Matching table for the coupling of three one-genelocus, two-allele premating barrier systems. The first system has mating-bias alleles, *X* and *Y*, with mating bias *α*. The second system has mating-bias alleles, *Z* and *W*, with mating bias *β*. The third system has mating-bias alleles, *J* and *K*, with mating bias *γ*. When two genotypes possess different mating-bias alleles at the barrier gene loci, their matching success is determined by the product of the mating biases at the mismatched gene loci.

**Table 3b.**
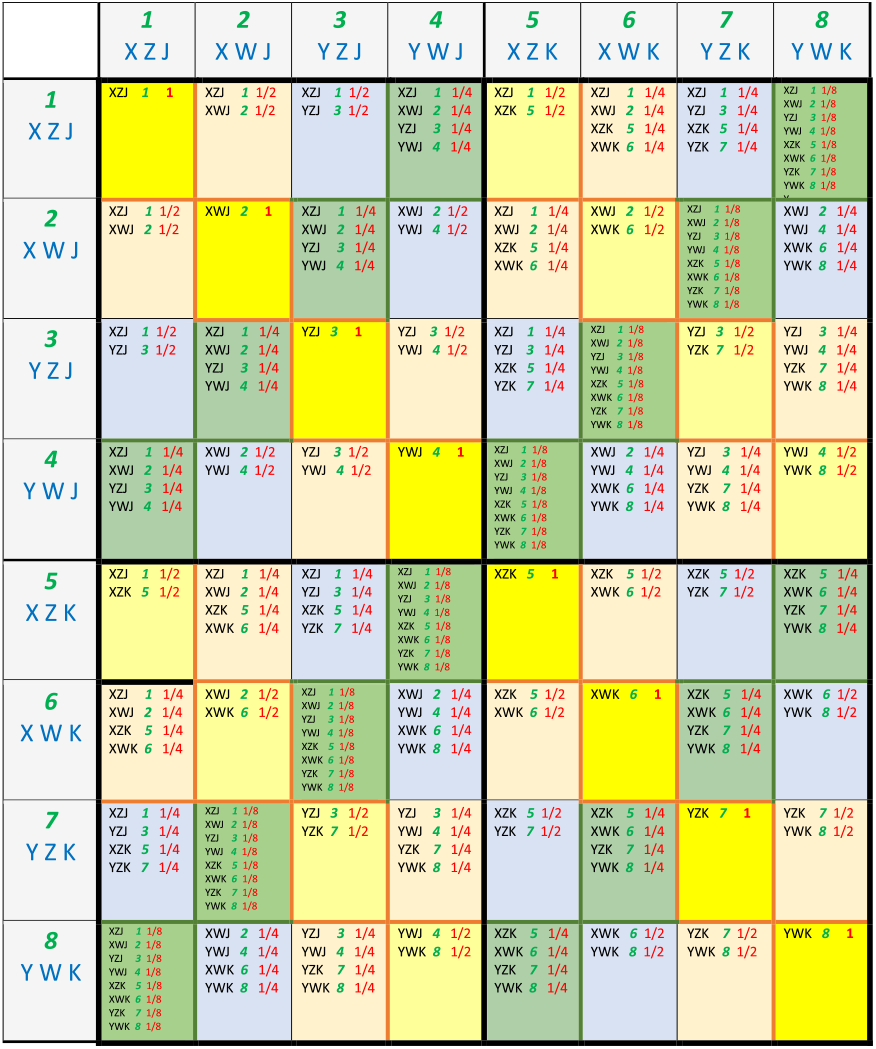
Unit offspring table for the coupling of three one-gene-locus, two-allele premating barrier systems.

**Fig 5.**
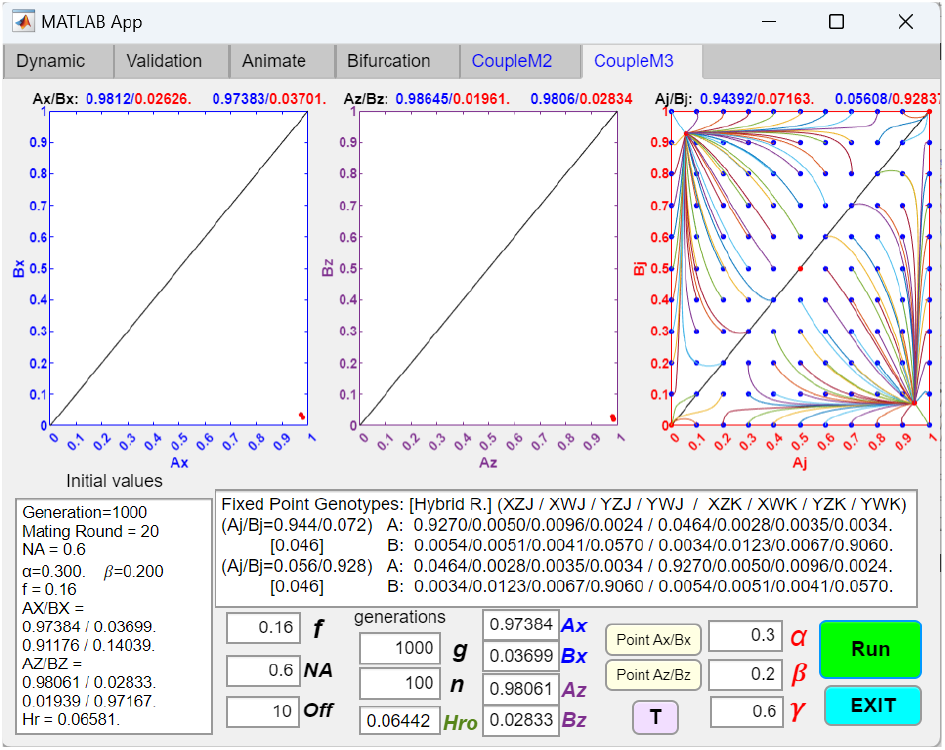
Changing the order of coupling affects the final outcome. Here, the order of coupling of the three two-allele systems in Fig 4 is changed. The values of the mating biases, *β* and *γ*, are reversed while keeping the other parameter values the same. This results in an invasion-resistant pattern appearing in the *AAA*/*BAA* phase portrait.

In Fig 6, the third two-allele barrier in Fig 5 is replaced by a one-allele barrier mechanism while keeping the overall maximum mating bias the same— i.e., the presence of the *K* allele changes *β* from 0.2 to 0.12 so that the overall maximum mating bias remains 0.036. With such a replacement, the third one-allele barrier can invade and further increase the overall premating RI. In general, it is easier to couple a one-allele mechanism than a two-allele mechanism. Less stringent parameters are required. This is because, unlike the two-allele mechanism, the one-allele mechanism is immune to the homogenizing effects of recombination.

**Fig 6.**
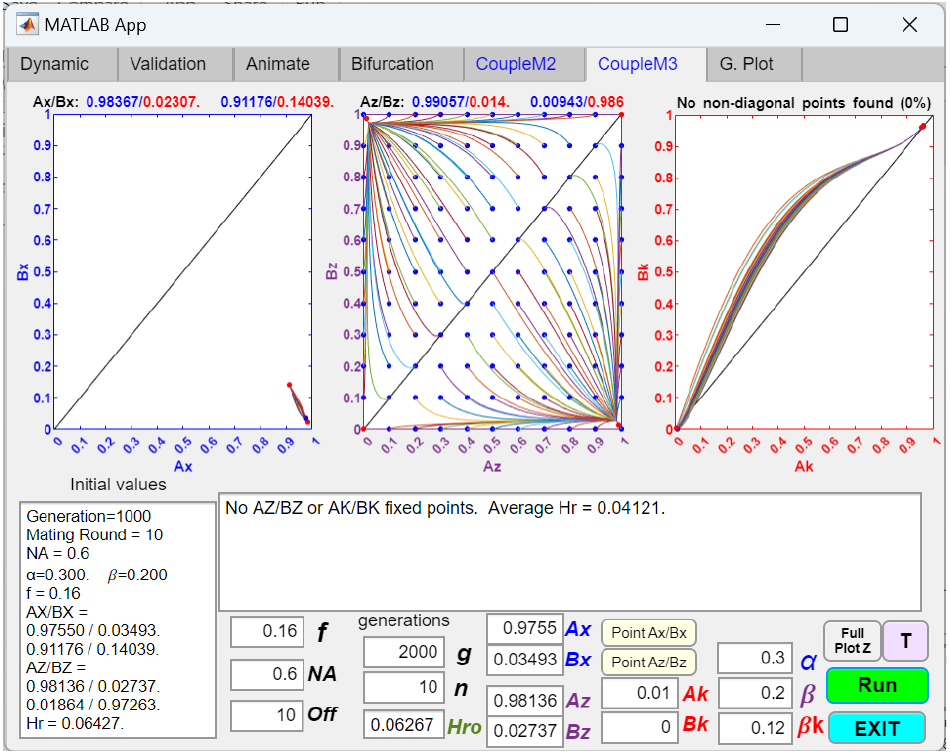
Coupling of two one-gene-locus, two-allele mating-bias barrier systems with a third, one-allele mating-bias system. The third mating-bias system in Fig 5 is changed to a one-allele system that modifies *β* into *βk* in the presence of a *K* allele in the genotypes (see Fig 1a). The phase portraits show that a mutant *K* allele with a population ratio *Ak* = 0.01 and mating bias *βk* = 0.12 can invade and increase the overall premating RI. The associated matching tables is included in the Appendix (Table 4).

Fig 7 demonstrates that an initially divergent two-allele system (*f* = 0.4, *α* = 0.3, *NA* = 0.6, *n* = 10) can be converted into a convergent system with fixed-point polymorphism through coupling with a strong two-allele barrier system (*β* = 0.08). In contrast to coupling with an already convergent system, converting a divergent system into a convergent system usually requires coupling with other barriers that have stronger mating biases and larger initial invading populations.

**Table 4.**
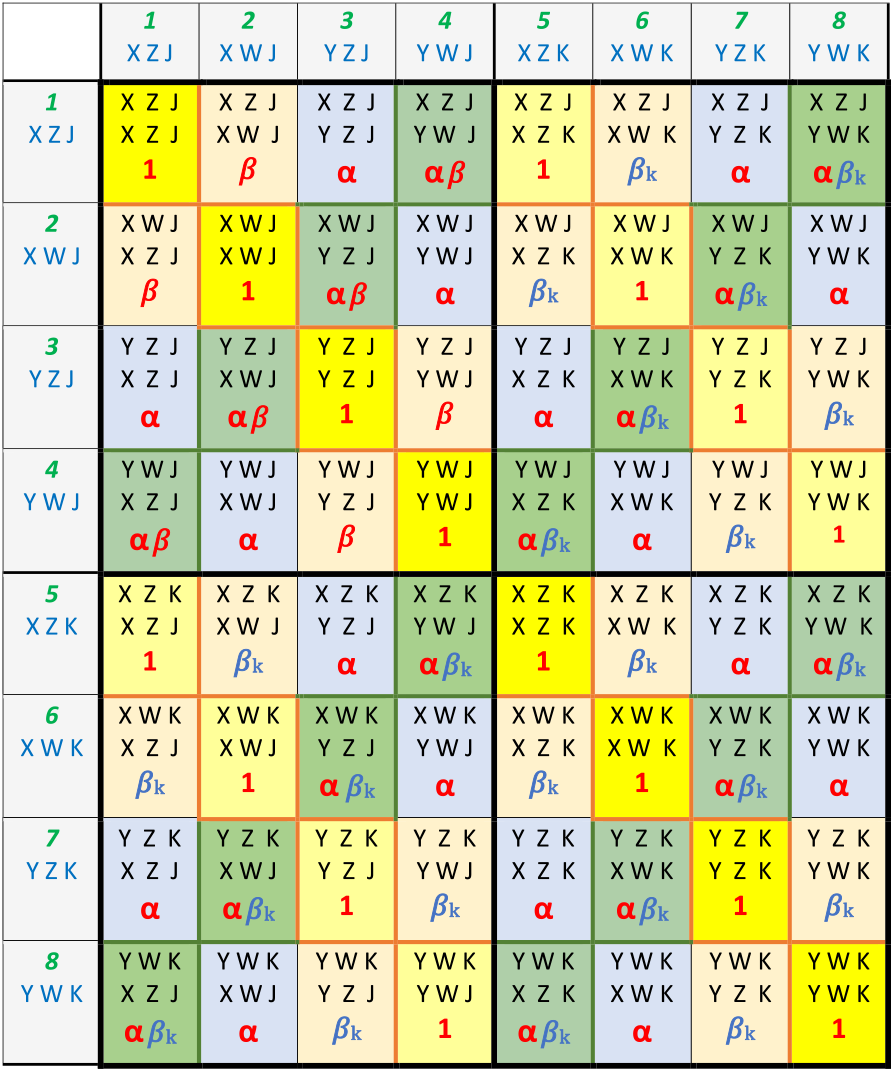
Matching table for the coupling of two one-gene-locus, two-allele premating barrier systems with a third, one-allele premating barrier system.

**Fig 7.**
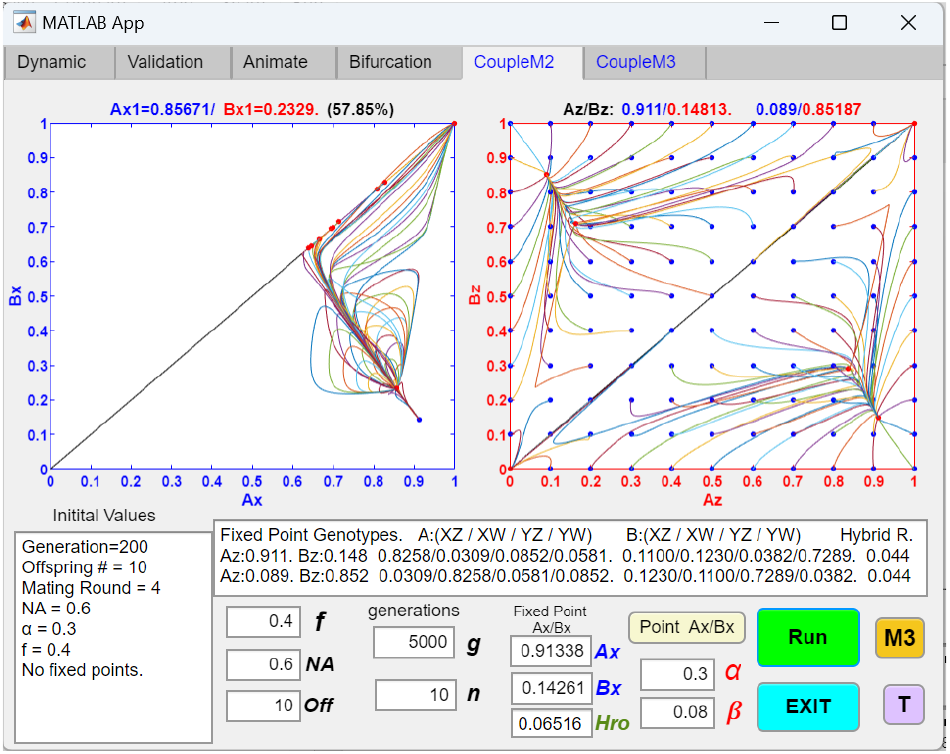
An initially divergent two-allele mating-bias barrier system can be converted into a convergent system through coupling with another two-allele system with a strong mating bias. Here, the second two-allele system has mating-bias alleles *Z* and *W*, and mating bias *β*. Notice that the *Ax*/*Bx* phase portrait shows an invasion resistant pattern.

Even though Fig 4 shows that the coupling of three different two-allele barrier loci can produce very strong overall premating RI, the premating RI can be reversed if the disruptive ecological selection is reduced or eliminated. As shown in Fig 8, if the value of *f* in Fig 4 is increased from 0.16 to 0.4, all three barrier systems become divergent. The increase in *f* required to reverse convergence is greater only because the overall mating-bias is stronger, as the effects of ecological selection and sexual selection are complementary in our mathematical model (Fig M3). These results seem to hold true whether the coupling is with one-allele mechanisms or two-allele mechanisms. They attest to the nonpermanent and reversible nature of premating RI created through population dynamics. Coupling with other late-stage post-zygotic mechanisms that can create hybrid incompatibility is therefore needed to make the premating RI irreversible [31].

**Fig 8.**
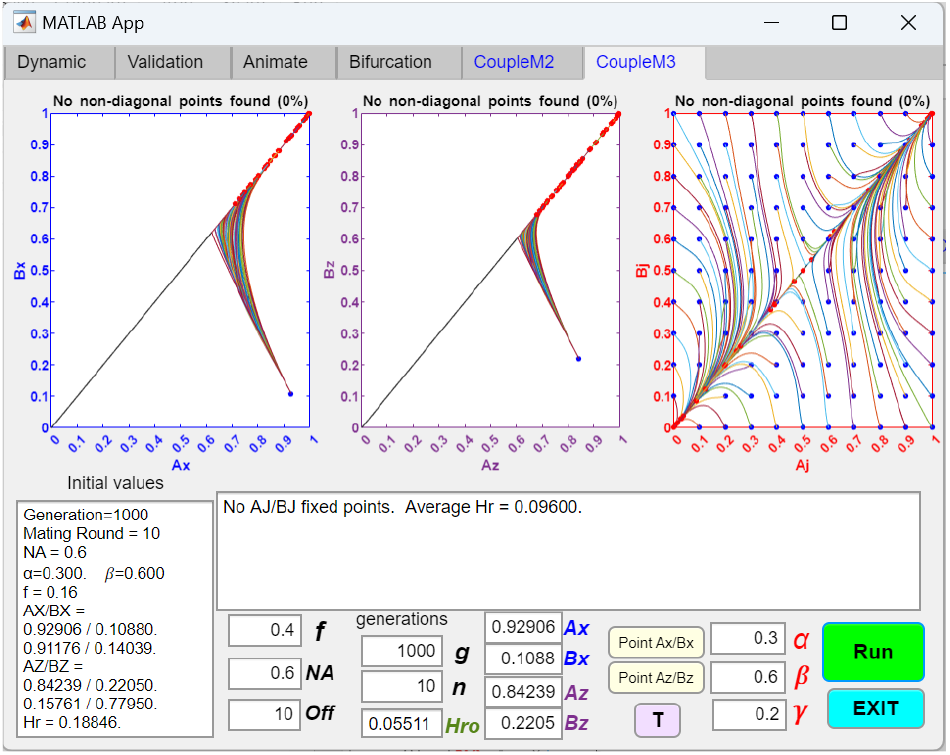
All coupled premating RI can be reversed by a reduction in disruptive ecological selection. If the value of *f* in Fig 4 is increased from 0.16 to 0.4, signifying a reduction in disruptive ecological selection, the phase portraits of all three coupled premating systems become divergent, and premating RI no longer exists.

### 2. Coupling of Late -Stage Premating and Post-mating Barriers

Next, we developed MATLAB GUI applications to investigate how initial, early-stage premating barriers can be coupled with late-stage barrier mechanisms to strengthen overall RI and make it irreversible. We studied three such late-stage barrier mechanisms: (1) alleles that affect hybrid viability, (2) chromosomal inversions, and (3) the BDM mechanism of incompatibility.

#### 2.1. Coupling with a mutant allele f^(−)^ or f_(+)_ that alters hybrid viability

Figs 9a and 9b show the phase portrait solutions of an initially convergent two-allele premating barrier system coupling with a one-allele barrier system that alters hybrid viability. Here, the presence of a *K* allele in an individual’s genotype signals the existence of an *f*_**(−)**_ or *f*_**(+)**_ mutation that alters hybrid viability. We use *f*_**(−)**_ to represent a mutation that decreases hybrid viability and *f*_**(+)**_ to represent a mutation that increases hybrid viability. When present in either parental genotype, the *K* allele alters the hybrid unit offspring ratios by changing the value of *f* to *fk* in the unit offspring table (see Table 5 in the Appendix). In general, a mutant allele *K* changing *f* to *fk* cannot invade if *fk* < *f*, but it can invade if *fk* > *f* (See Fig 9a and 9b). This is because when *fk* < *f*, a niche-*A* individual with genotype *AXK* will suffer more offspring loss when it mates with an individual from niche *B* (because it has more nonviable hybrid offspring) than its cohorts in niche *A* that do not have the mutant *K* allele. The end result is that the *AXK* individuals have a fitness disadvantage when compared to the *AX* individuals in niche *A*, so the *K* allele tends to get eliminated.

**Table 5.**
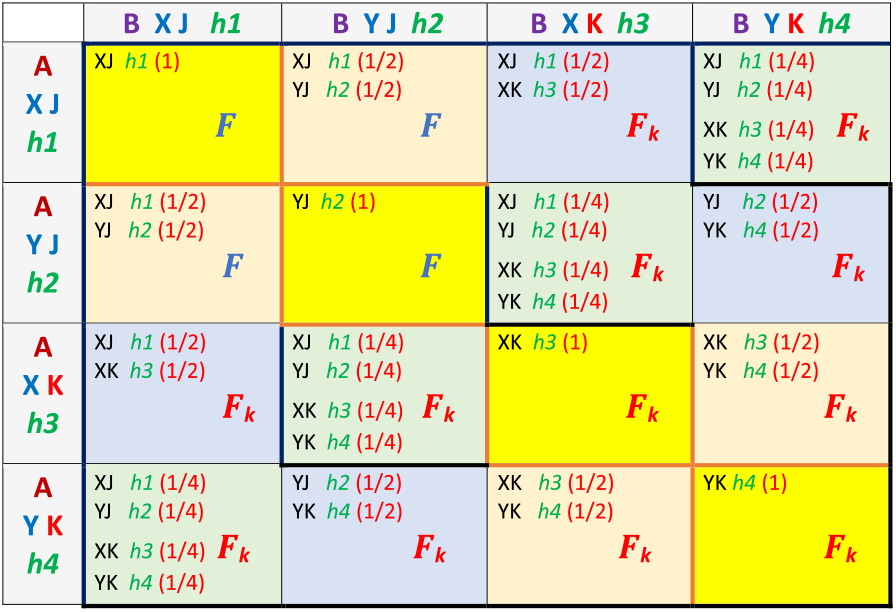
Unit offspring table for the coupling of a one-allele barrier system that alters hybrid viability. In the presence of a *K* allele, the value of *f* is changed to *F*_*k*_ to produce unit offspring in inter-niche mating. The *J* allele simply signals the absence of the *K* allele and has no effect.

**Fig 9a.**
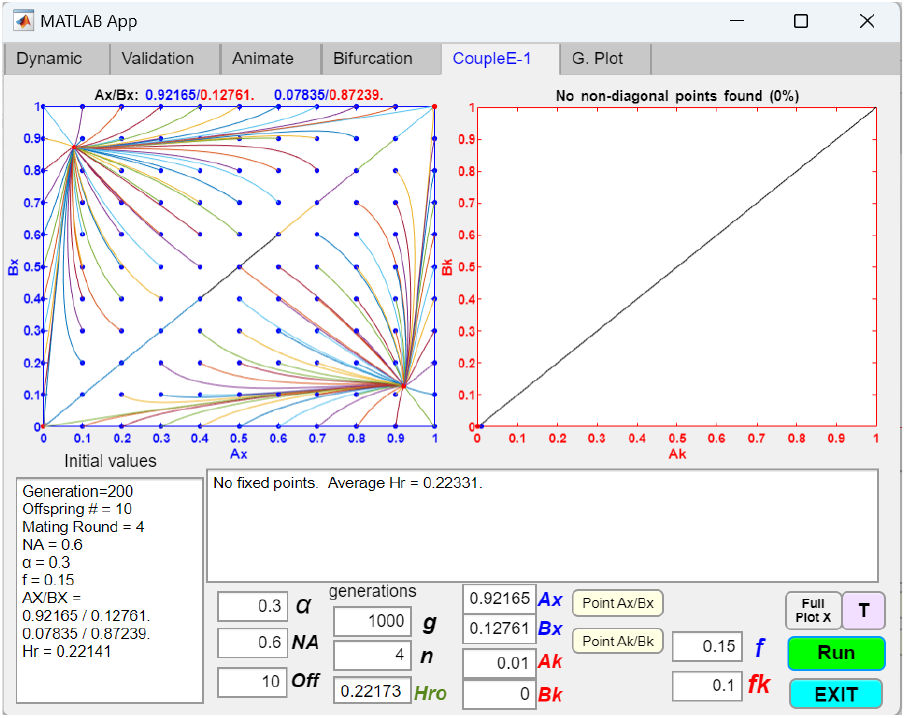
Coupling of an initial two-allele premating barrier with a one-allele barrier mechanism that decreases hybrid viability. An initial two-allele premating barrier mechanism establishes a fixed-point polymorphism at *Ax*/*Bx* = 0.92165/0.12761. It then attempts to couple with a one-allele barrier mechanism that decreases hybrid viability. The presence of a *K* allele at a separate gene locus in an individual’s genotype signals the presence of an *f*_**(-)**_ mutation that decreases the viability of its hybrid offspring from *f* to *fk*. As shown, a mutant allele *K* with *fk* < *f* and an initial population ratio of 0.01 cannot invade because it suffers a fitness disadvantage in inter-niche mating compared to its cohorts in the same niche, which do not suffer such increased hybrid offspring loss. The presence of the *K* allele only alters the hybrid unit-offspring ratios. The modified unit offspring table for the GUI application is included in the Appendix (Table 5).

**Fig 9b.**
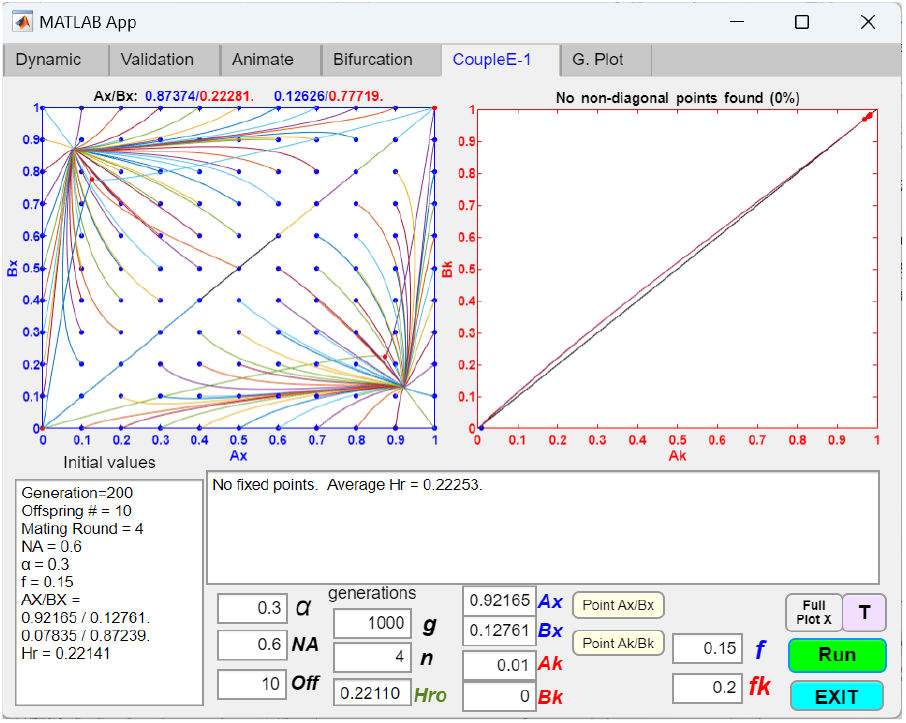
Coupling of an initial two-allele premating barrier with a one-allele barrier mechanism that increases hybrid viability. If the value of *fk* in Fig 9a is changed to 0.2 (so that *fk* > *f*) while keeping the rest of the parametric values the same, a small mutant population of *K* alleles can invade and become fixed. However, the fixed point in the *Ax*/*Bx* phase portrait is shifted to *Ax*/*Bx* = 0.87374/0.22281 (show as a red dot), signifying reduced overall premating RI as a result of the coupling.

**Fig 9c.**
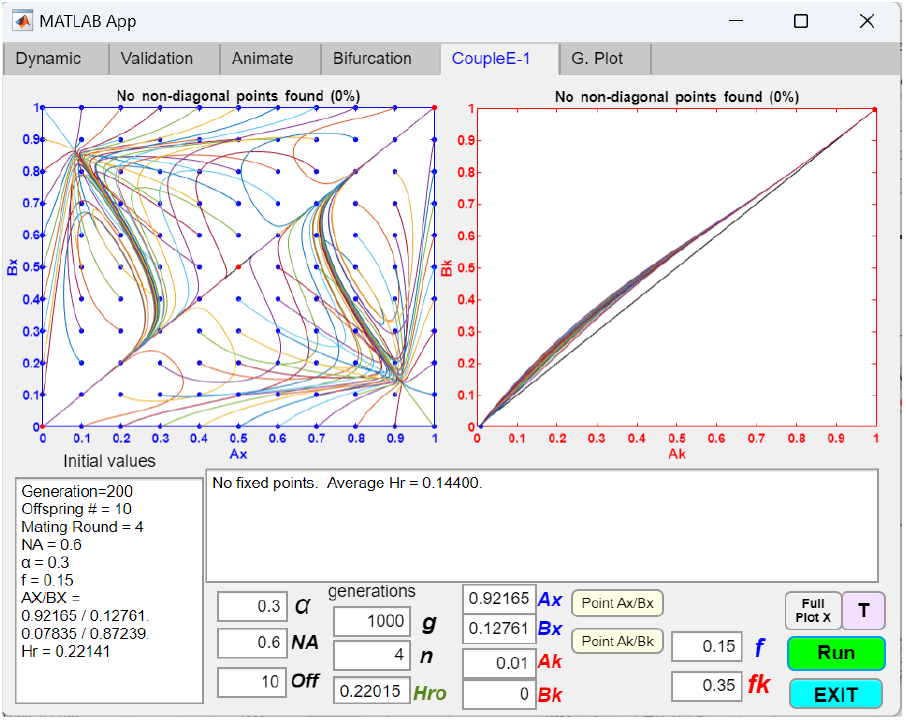
A convergent two-allele premating barrier system is converted into a divergent system after coupling with a one-allele barrier mechanism that increases hybrid viability. If the value of *fk* in Fig 9b is further increased to 0.35, then the coupling can cause divergence of the initially convergent two-allele barrier system.

Fig 9b shows that when a mutant *K* allele with a higher value *fk* invades, it can cause the fixed point of the existing mating-bias system to shift away from the lower right corner and reduce the overall premating RI (i.e., negative coupling [32]). Fig 9c shows that if the value of *fk* is high enough, the *K* allele can invade and cause an existing convergent mating-bias system to become divergent.

Fig 10 shows an initial premating barrier coupling with a two-allele barrier mechanism that alters hybrid viability. Two alleles, *J* and *K*, exist at a gene locus that controls hybrid viability. The value of *f* is changed to *Fjk* only when two mating parents possess different alleles, *K* or *J*. The two-allele model yields similar results as the one-allele model: only mutants with *Fjk* > *f* can invade. Except in this case, the *Ak* and *Bk* ratios converge to the fixed-point values *Ak* = 0.5 and *Bk* = 0.5 (see Fig 10)

**Fig 10.**
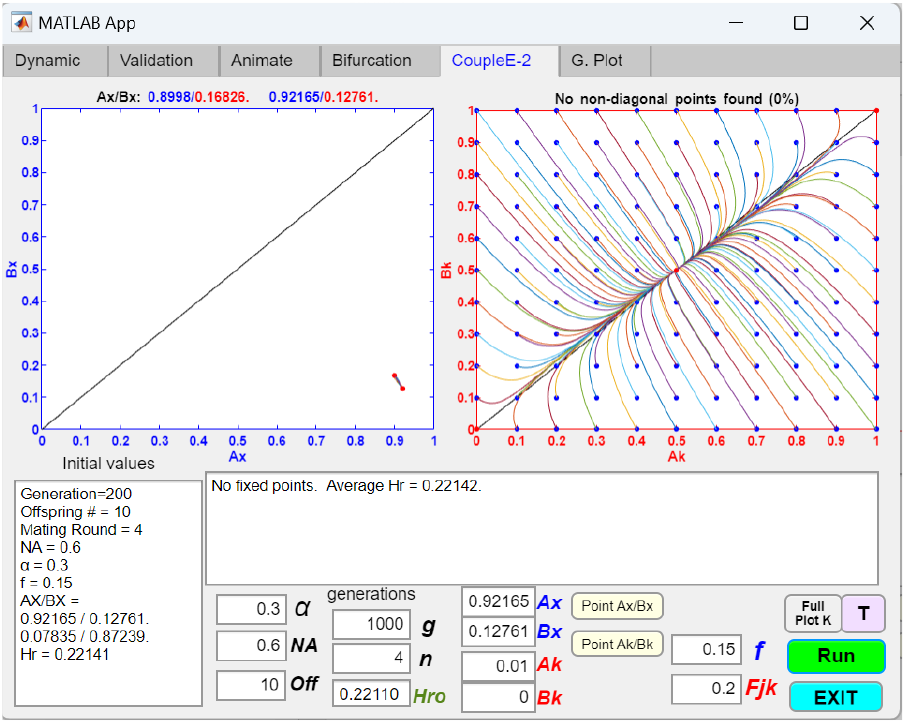
Coupling of an initial two-allele premating barrier with a two-allele barrier mechanism that alters hybrid viability. In this two-allele model that alters hybrid viability, two alleles, *J* and *K*, exist at a single gene locus. In inter-niche mating, if parental genotypes possess different alleles, *J* or *K*, the value of *f* is changed to *FAAk* in the unit offspring table, thereby altering the selection against hybrids and the resulting offspring ratios. The *Ak*/*Bk* phase portrait is plotted against the *Ax*/*Bx* fixed point established by the initial premating barrier. In this example, the value of *FAAk* (0.2) is greater than the value of *f* (0.15), and the *Ak*/*Bk* phase portrait is globally convergent. Consequently, small *Ak*/*Bk* mutant populations near the origin can invade and converge to a fixed-point polymorphism at *Ak* = *Bk* = 0.5. This causes the *Ax*/*Bx* fixed point to shift away from the lower right corner of the phase portrait, resulting in reduced premating RI. The unit offspring table used by the GUI application is included in the Appendix (Table 6).

While a mutation, *f*_**(+)**_, that increases hybrid viability could invade and weaken overall reproductive isolation, this is unlikely if disruptive ecological selection fails to provide sufficient food resources to sustain the hybrid genotypes. Increased hybrid viability would only occur if the mutation, *f*_**(+)**_, enhances the hybrids’ ability to more efficiently extract existing resources. Thus, the availability of niche resources imposes an upper limit on how much intrinsic genetic changes can increase hybrid viability and population size.

**Table 6.** Unit offspring table for the coupling of a two-allele barrier system that alters hybrid viability. In inter-niche mating, if the parental genotypes possess different alleles, *J* or *K*, the value of *f* is changed to *F*_*jk*_ when determining the offspring ratios.

|  | B XJ h1 | B YJ h2 | B XK h3 | B YK h4 |
| --- | --- | --- | --- | --- |
| A XJ h1 | XJ h1 (1)<br>$F$ | XJ h1 (1/2)<br>YJ h2 (1/2)<br>$F$ | XJ h1 (1/2)<br>XK h3 (1/2)<br>$F_{jk}$ | XJ h1 (1/4)<br>YJ h2 (1/4)<br>XK h3 (1/4) $F_{jk}$<br>YK h4 (1/4) |
| A YJ h2 | XJ h1 (1/2)<br>YJ h2 (1/2)<br>$F$ | YJ h2 (1)<br>$F$ | XJ h1 (1/4)<br>YJ h2 (1/4)<br>XK h3 (1/4) $F_{jk}$<br>YK h4 (1/4) | YJ h2 (1/2)<br>YK h4 (1/2)<br>$F_{jk}$ |
| A XK h3 | XJ h1 (1/2)<br>XK h3 (1/2)<br>$F_{jk}$ | XJ h1 (1/4)<br>YJ h2 (1/4)<br>XK h3 (1/4) $F_{jk}$<br>YK h4 (1/4) | XK h3 (1)<br>$F$ | XK h3 (1/2)<br>YK h4 (1/2)<br>$F$ |
| A YK h4 | XJ h1 (1/4)<br>YJ h2 (1/4)<br>XK h3 (1/4) $F_{jk}$<br>YK h4 (1/4) | YJ h2 (1/2)<br>YK h4 (1/2)<br>$F_{jk}$ | XK h3 (1/2)<br>YK h4 (1/2)<br>$F$ | YK h4 (1)<br>$F$ |

Our mathematical model (Fig M3) predicts that a lower value of *f* can increase ecological selection and bring about greater assortative mating (by shifting the *Ax*/*Bx* fixed point toward the lower right corner in the phase portrait) and create stronger premating RI. However, by themselves, *f*_**(−)**_ mutant alleles that decrease *f* cannot invade because the increased hybrid loss from inter-niche mating gives them a fitness disadvantage when compared to their cohorts. The only way such negative-fitness alleles can invade and further increase hybrid loss is through linkage to other mechanisms with sufficient fitness advantage to offset their fitness disadvantage.

Fig 11a shows the coupling between an initial two-allele premating barrier and a second two-allele barrier system that can alter the overall mating bias and hybrid viability. The example demonstrates that, given favorable parametric conditions, if a mutant allele *Z* can produce both a strong mating bias and a lower value of *f*, such an allele can invade and creating an overall stronger premating RI. In this case, the fitness advantage of having a low *β* overcomes the fitness disadvantage of having a low *f*2. After coupling, the lower overall *f* value conveyed by the Z allele augments the premating RI due to the stronger mating bias in a synergistic fashion.

**Fig 11a.**
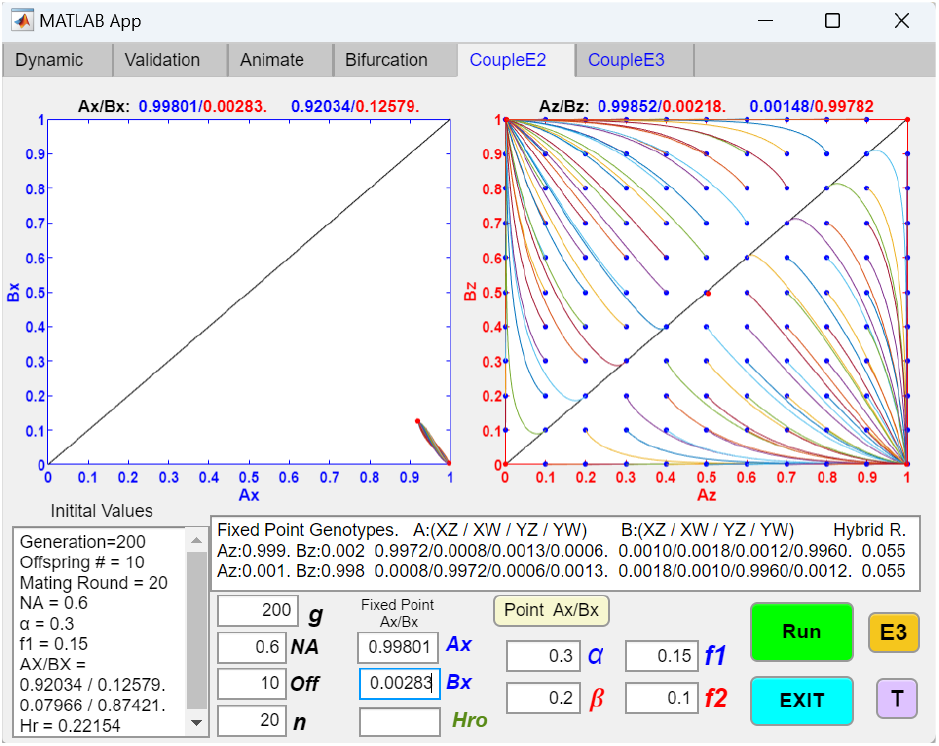
Coupling of an initial two-allele premating barrier with a two-allele barrier mechanism that alters both mating bias and hybrid viability. The phase portraits show the results of coupling between an initial two-allele premating barrier and another two-allele barrier system that affects both mating bias and hybrid viability. The second two-allele barrier system has two alleles, *Z* and *W*, which are associated with a mating bias value, *β*, and an offspring return ratio, *f*2. When genotypes carrying different alleles, *Z* or *W*, encounter each other, their matching success is determined by multiplying the mating bias between them by *β*. Similarly, in inter-niche mating, if the matched parental pairs have different *Z* or *W* alleles, the value of *f* (denoted as *f*1) is multiplied by *f*2 in the unit offspring table to determine the unit offspring ratios. In this example, the *Ax*/*Bx* phase portrait, plotted against the initial *Ax*/*Bx* fixed point, is globally convergent. A mutant *Z* allele with a low associated *β* value gains a fitness advantage, which allows it to invade despite the fitness cost of having a low associated *f*2 value. Consequently, a negatively selected low *f*2 value is able to hitchhike with a positively selected low *β* value to invade and exert its effects. This close linkage between high mating bias (low *β* value) and low hybrid viability (low *f*2 value) provided by alleles in the second barrier system can result in strong overall RI. The matching and unit-offspring tables used in the GUI application are included in the Appendix (Tables 7a and 7b).

**Fig 11b.**
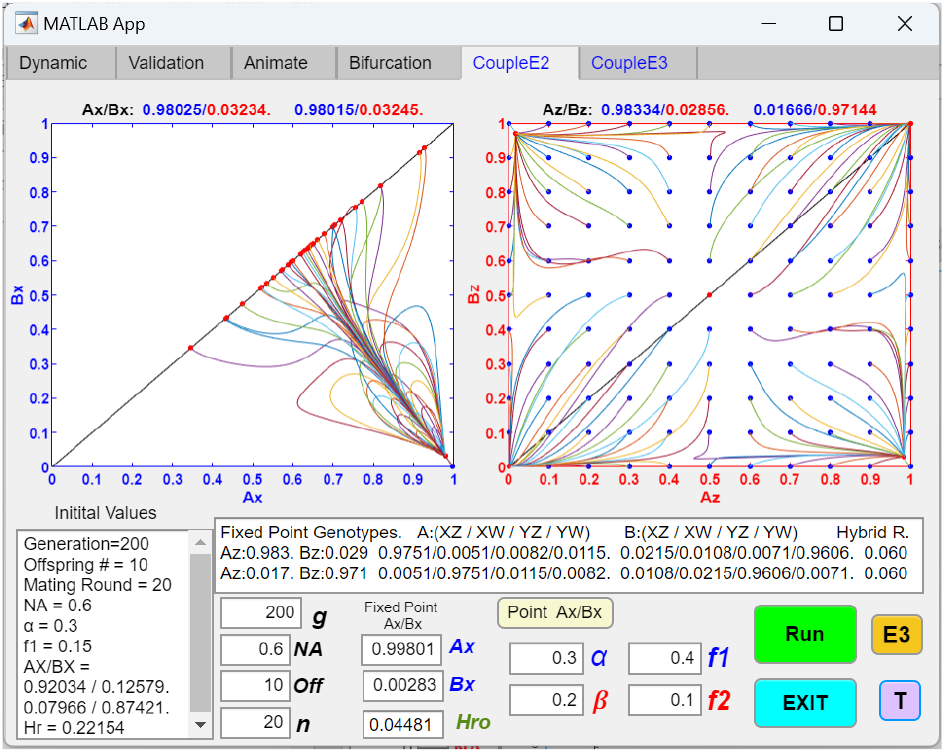
Coupling with a two-allele barrier mechanism that alters both mating bias and hybrid viability can produce RI that is resistant to reversal. After the population genotype ratios of the two coupled systems have stabilized at the fixed points shown in Fig 11a (i.e., at *Ax*/*Bx* = 0.99801/0.00283 and *Ax*/*Bx* = 0.99852/ 0.00218), if the original disruptive ecological selection is reduced by increasing the value of *f*1 from 0.15 to 0.4, the *f*2 value (0.1) associated with the nearly fixed *Ax*/*Bw* genotypes can still take over to ensure continued hybrid loss and maintain RI.

Fig 11b shows that if we let the population start at the system’s fixed-point polymorphisms in Fig 11a after the invasion of the *Z* allele (i.e., starting at *Ax*/*Bx* = 0.99801/0.00283 and *Ax*/*Bx* = 0.99852 /0.00218) and if we then reduce the original disruptive ecological selection by increasing the value of *f*1 from 0.15 to 0.4, the new *f*2 value of 0.1, conveyed by the now nearly fixed *Ax* and *Bw* genotypes, can still take over to ensure continued high hybrid loss and maintain fixed-point polymorphism and premating RI. However, if we completely remove the disruptive ecological selection by setting *f*1 = 0.5, both the *Ax*/*Bx* and *Ax*/*Bx* fixed points disappear, the system becomes divergent, and no premating RI exists. Therefore, the RI generated by our coupled system appears to have a certain degree of resilience against reductions in disruptive ecological selection, although this resistance to reversibility is not absolute.

**Table 7a.** Matching table for coupling to a two-allele barrier mechanism that alters both mating bias and hybrid viability. When two genotypes possess different allele, *Z* and *W*, the mating bias between them is multiplied by *β*.

|  | XZ h1 | XW h2 | YZ h3 | YW h4 |
| --- | --- | --- | --- | --- |
| XZ h1 | X Z<br>X Z<br>1 | X Z<br>X W<br>$\beta$ | X Z<br>Y Z<br>$\alpha$ | X Z<br>Y W<br>$\alpha\beta$ |
| XW h2 | X W<br>X Z<br>$\beta$ | X W<br>X W<br>1 | X W<br>Y Z<br>$\alpha\beta$ | X W<br>Y W<br>$\alpha$ |
| YZ h3 | Y Z<br>X Z<br>$\alpha$ | Y Z<br>X W<br>$\alpha\beta$ | Y Z<br>Y Z<br>1 | Y Z<br>Y W<br>$\beta$ |
| YW h4 | Y W<br>X Z<br>$\alpha\beta$ | Y W<br>X W<br>$\alpha$ | Y W<br>Y Z<br>$\beta$ | Y W<br>Y W<br>1 |

**Table 7b.** Unit offspring table for the coupling of a two-allele barrier system that alters both mating bias and hybrid viability. In inter-niche mating, when two parental genotypes possess different alleles, *Z* and *W*, the overall *f* value used to determine offspring genotype ratios is the product of *f*1 × *f*2.

|  | B XZ h1 | B XW h2 | B YZ h3 | B YW h4 |
| --- | --- | --- | --- | --- |
| A XZ h1 | XZ h1 (1) $f1$ | XZ h1 (1/2) $f1f2$<br>XW h2 (1/2) $f1f2$ | XZ h1 (1/2) $f1$<br>YZ h3 (1/2) $f1$ | XZ h1 (1/4) $f1f2$<br>XW h2 (1/4) $f1f2$<br>YZ h3 (1/4) $f1f2$<br>YW h4 (1/4) $f1f2$ |
| A XW h2 | XZ h1 (1/2) $f1f2$<br>XW h2 (1/2) $f1f2$ | XW h2 (1) $f1$ | XZ h1 (1/4) $f1f2$<br>XW h2 (1/4) $f1f2$<br>YZ h3 (1/4) $f1f2$<br>YW h4 (1/4) $f1f2$ | XW h2 (1/2) $f1$<br>YW h4 (1/2) $f1$ |
| A YZ h3 | XZ h1 (1/2) $f1$<br>YZ h3 (1/2) $f1$ | XZ h1 (1/4) $f1f2$<br>XW h2 (1/4) $f1f2$<br>YZ h3 (1/4) $f1f2$<br>YW h4 (1/4) $f1f2$ | YZ h3 (1) $f1$ | YZ h3 (1/2) $f1f2$<br>YW h4 (1/2) $f1f2$ |
| A YW h4 | XZ h1 (1/4) $f1f2$<br>XW h2 (1/4) $f1f2$<br>YZ h3 (1/4) $f1f2$<br>YW h4 (1/4) $f1f2$ | XW h2 (1/2) $f1$<br>YW h4 (1/2) $f1$ | YZ h3 (1/2) $f1f2$<br>YW h4 (1/2) $f1f2$ | YW h4 (1) $f1$ |

#### 2.2. Coupling with chromosomal inversions

Chromosomal inversions (CIs) have long been implicated in playing a role in sympatric speciation [33-36]. Locally adaptive alleles captured in the inverted regions of CIs are resistant to recombination during heterokaryotype mating, effectively turning them into “super-alleles” that link and preserve favorable allele combinations [37-40]. This gives them a fitness advantage to invade and accelerates the accumulation of adaptive ecological and mating-bias alleles in the different niches undergoing disruptive selection [39, 41]. Chromosomal inversions are considered a late-stage mechanism because a CI is more likely to capture locally adaptive and advantageous ecological and mating-bias allele combinations when there is already a certain degree of nonrandom assortment of those alleles in different niches due to prior barriers [42]. Chromosomal inversion is considered an adaptive barrier mechanism [35, 42]. This means that the invasion of a CI mutant is driven by the selection pressure created by hybrid loss during disruptive ecological selection.

Chromosomal inversions have certain peculiar properties that we must consider when developing computer applications to model their behavior. Single crossovers between inverted and noninverted regions in heterokaryotype mating can produce nonviable hybrids [38, 43]. Similarly, the rarer events of double crossovers and gene conversion in heterokaryotype mating can cause recombination within the inverted regions [44, 45]. For the most part, the effects of these uncommon events can be ignored in our modeling, but they may become important in special circumstances.

In our study, we have developed MATLAB GUI applications to investigate how an initial premating barrier, established using our 2-allele mathematical model without viable hybrids (Fig M3), could be coupled with other CI barrier mechanisms. We first investigate CIs that capture only locally adaptive ecological alleles before turning our attention to CIs that capture both ecological and mating-bias alleles.

##### 2.2.1. Chromosomal inversions that capture only ecological alleles

Figs 12a and 12b show the phase portrait solutions of an initial two-allele premating barrier coupled with a chromosomal inversion that captures locally adaptive ecological alleles. *Ak* signifies the presence of a chromosomal inversion in niche *A* that captures the niche-*A* ecological alleles. *Bk* is always zero. *Fk*1 denotes the offspring ratio that is returned to the parental genotypes in intra-niche (e.g., between niche *A* individuals) heterokaryotype mating. *Fk*2 denotes the offspring ratio that is returned to the parental genotypes in inter-niche (between niche *A* and niche *B* individuals) heterokaryotype mating. The values of *Fk*1 and *Fk*2 range from 0 to 0.5.

**Fig 12a.**
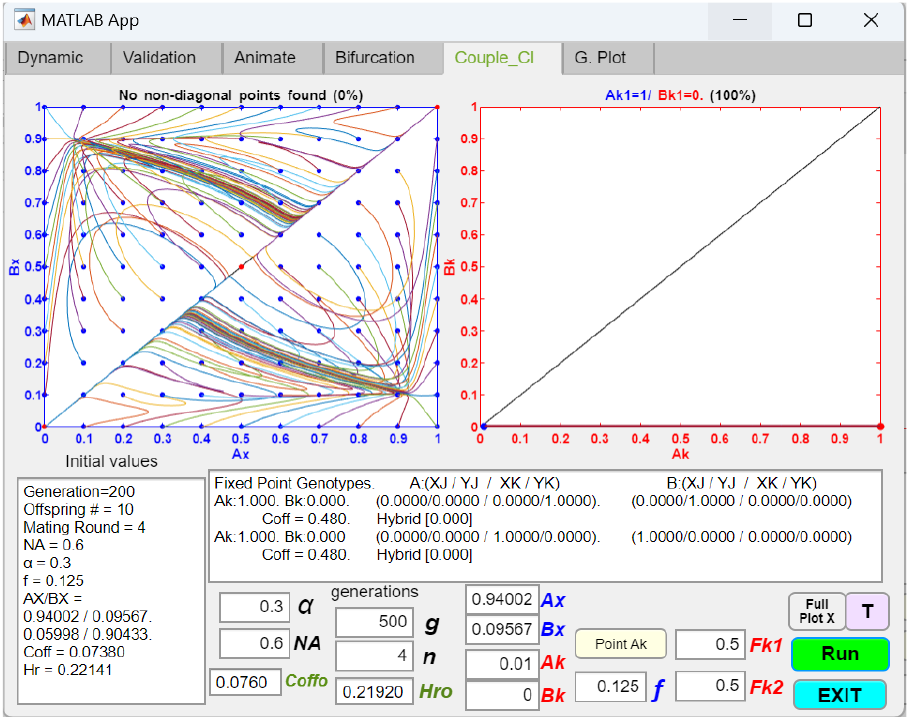
Coupling of an initial two-allele premating barrier with a chromosomal inversion that captures locally adaptive ecological alleles. The phase portraits show the results of coupling an initial two-allele premating barrier with a chromosomal inversion (CI) that captures locally adaptive ecological alleles in niche A. The table insert shows the fixed-point polymorphism (*Ax*/*Bx* = 0.94002/0.09567) achieved by the initial two-allele premating barrier and the associated system parameters. We use allele *K* to denote the presence of a CI in a genotype. The *Ak*/*Bk* phase portrait shows that a CI capturing locally adaptive ecological alleles in niche *A*, with a starting population ratio of *Ak* = 0.01 successfully invades and becomes fixed in niche *A. Bk*, by definition, is always zero because the niche-*B* ecotype does not have any alleles of the niche-*A* ecotype. *Fk*1 is the offspring return ratio in intra-niche heterokaryotype mating. *Fk*2 is the offspring return ratio in inter-niche heterokaryotype mating. An *Fk*2 = 0.5 means that the CI captures all ecologically adaptive alleles in niche *A*, without producing any hybrid offspring or suffering any offspring loss in inter-niche heterokaryotype mating. *Hro* = 0.21920 is the initial hybrid ratio from inter-niche mating, and *Coffo* = 0.0760 is the initial non-hybrid offspring ratio from inter-niche mating. *Hro* + *Coffo* = 0.29520 is the initial total offspring ratio from inter-niche mating. The phase portraits show that a CI with *Fk*2 = 0.5 is able to invade and cause the initially convergent *Ax*/*Bx* phase portrait to become divergent. The final hybrid ratio after 500 generations is zero. The final non-hybrid offspring ratio from inter-niche mating, *Coff*, is 0.48. Even though no ecological hybrid offspring are produced, the total offspring ratio from inter-niche mating (*Coff* + *Hr* = 0.48) is increased, indicating reduced RI between the niches. The matching and unit-offspring tables used in the GUI application are included in the Appendix (Tables 8 and 9).

**Fig 12b.**
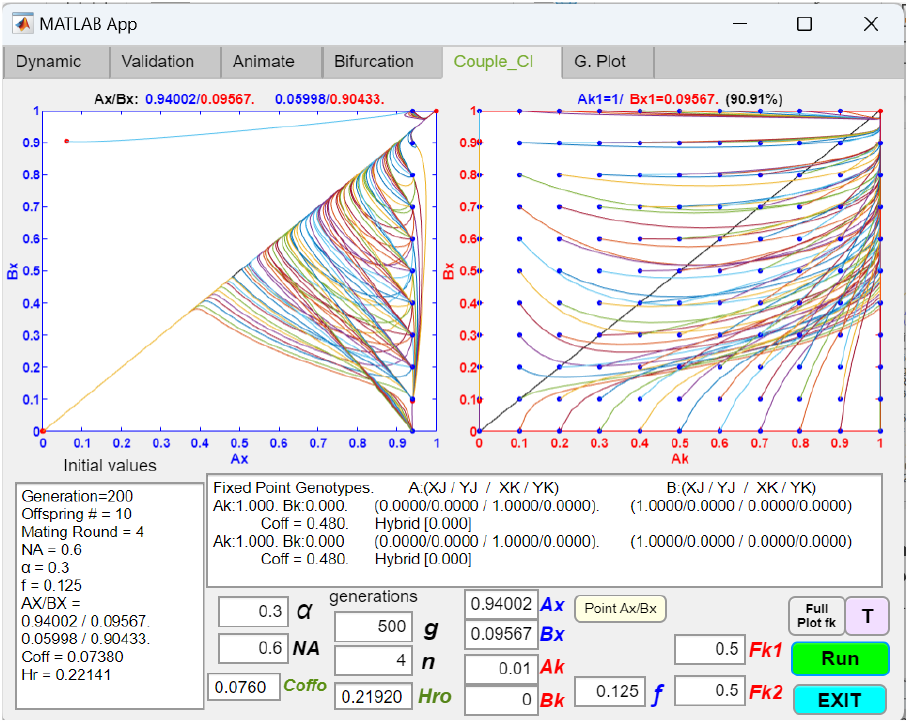
Coupling of an initial two-allele premating barrier with a chromosomal inversion that captures locally adaptive ecological alleles, showing an *Ak*/*Bx* phase portrait. Using the same parametric values as in Fig 12a, an *Ak*/*Bx* phase portrait, plotted against the initial *Ax*/*Bx* fixed point, is shown.

In inter-niche heterokaryotype mating, an *Fk*2 value of 0.5 is possible if a CI can capture all of the locally adaptive ecological alleles so that the linked alleles cannot be broken up by recombination. Consequently, 50% of the offspring have the same ecological genotype as the niche-*A* parent, and 50% have the same genotype as the niche-*B* parent, with no hybrid offspring produced.

**Table 8.** Matching table for coupling to a chromosomal inversion.

|  | A XJ h1 | A YJ h2 | A XK h3 | A YK h4 | B XJ h1 | B YJ h2 | B XK h3 | B YK h4 |
| --- | --- | --- | --- | --- | --- | --- | --- | --- |
| A XJ h1 | X J<br>X J<br>1 | X J<br>Y J<br>$\alpha$ | X J<br>X K<br>1 | X J<br>Y K<br>$\alpha$ | X J<br>X J<br>1 | X J<br>Y J<br>$\alpha$ | X J<br>X K<br>1 | X J<br>Y K<br>$\alpha$ |
| A YJ h2 | X J<br>Y J<br>$\alpha$ | Y J<br>Y J<br>1 | X K<br>Y J<br>$\alpha$ | Y J<br>Y K<br>1 | X J<br>Y J<br>$\alpha$ | Y J<br>Y J<br>1 | X K<br>Y J<br>$\alpha$ | Y J<br>Y K<br>1 |
| A XK h3 | X J<br>X K<br>1 | X K<br>Y J<br>$\alpha$ | X K<br>X K<br>1 | X K<br>Y K<br>$\alpha$ | X J<br>X K<br>1 | X K<br>Y J<br>$\alpha$ | X K<br>X K<br>1 | X K<br>Y K<br>$\alpha$ |
| A YK h4 | X J<br>Y K<br>$\alpha$ | Y J<br>Y K<br>1 | X K<br>Y K<br>$\alpha$ | Y K<br>Y K<br>1 | X J<br>Y K<br>$\alpha$ | Y J<br>Y K<br>1 | X K<br>Y K<br>$\alpha$ | Y K<br>Y K<br>1 |
| B XJ h1 | X J<br>X J<br>1 | X J<br>Y J<br>$\alpha$ | X J<br>X K<br>1 | X J<br>Y K<br>$\alpha$ | X J<br>X J<br>1 | X J<br>Y J<br>$\alpha$ | X J<br>X K<br>1 | X J<br>Y K<br>$\alpha$ |
| B YJ h2 | X J<br>Y J<br>$\alpha$ | Y J<br>Y J<br>1 | X K<br>Y J<br>$\alpha$ | Y J<br>Y K<br>1 | X J<br>Y J<br>$\alpha$ | Y J<br>Y J<br>1 | X K<br>Y J<br>$\alpha$ | Y J<br>Y K<br>1 |
| B XK h3 | X J<br>X K<br>1 | X K<br>Y J<br>$\alpha$ | X K<br>X K<br>1 | X K<br>Y K<br>$\alpha$ | X J<br>X K<br>1 | X K<br>Y J<br>$\alpha$ | X K<br>X K<br>1 | X K<br>Y K<br>$\alpha$ |
| B YK h4 | X J<br>Y K<br>$\alpha$ | Y J<br>Y K<br>1 | X K<br>Y K<br>$\alpha$ | Y K<br>Y K<br>1 | X J<br>Y K<br>$\alpha$ | Y J<br>Y K<br>1 | X K<br>Y K<br>$\alpha$ | Y K<br>Y K<br>1 |

**Table 9.**
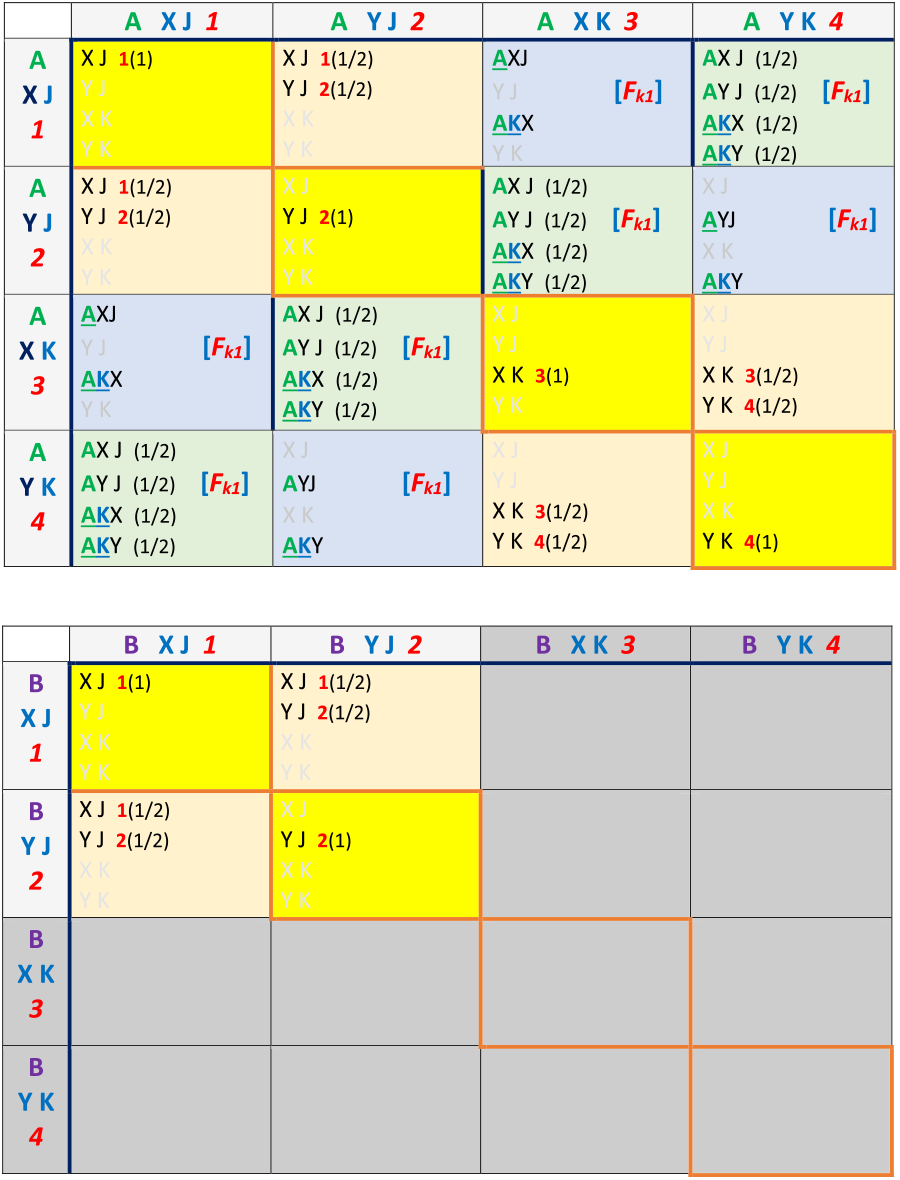
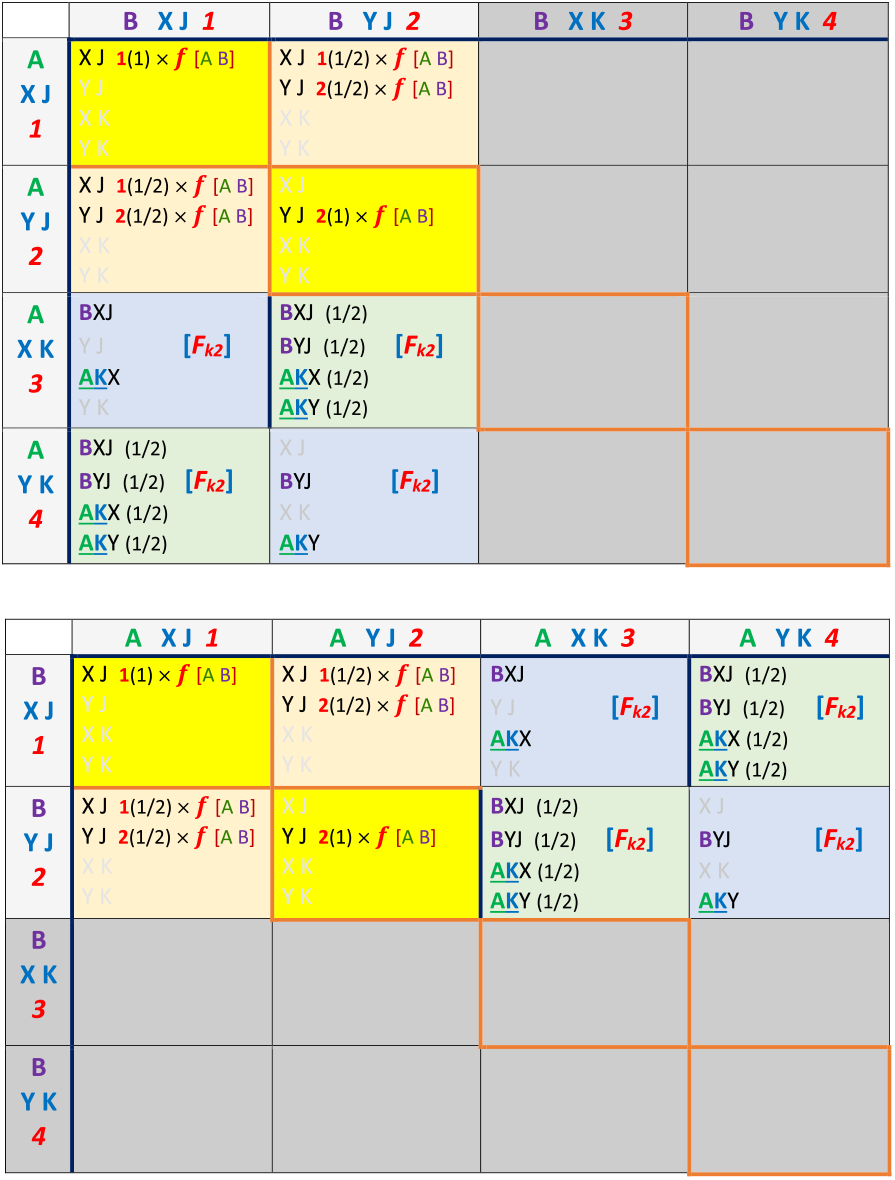
Unit offspring tables for the coupling of a chromosomal inversion that captures locally adaptive ecological alleles in niche *A*. The *K* allele denotes the presence of the CI in the genotype, while the *J* allele denotes the absence of the *K* allele. *Fk*1 is the offspring return ratio in intra-niche heterokaryotype mating. *Fk*2 is the offspring return ratio in inter-niche heterokaryotype mating. *f* is the offspring return ratio in inter-niche homokaryotype mating. By definition, it is impossible to have the *BK* genotype. Therefore, all cells associated with the *BK* genotype are grayed out.

The value of *Fk*2 decreases if the CI captures only a partial number of all the locally adaptive alleles. In this case, hybrid offspring are produced in inter-niche heterokaryotype matings. If the CI also happens to capture an allele *f*_**(−)**_ that reduces hybrid viability, this can further lower the value of *Fk*2. Less common events, such as single and double crossovers and gene conversion in the inverted regions, can also decrease the values of *Fk*1 and *Fk*2.

In the figures, the variable *Coff* represents the non-hybrid offspring ratio from inter-niche mating, and *Hr* represents the hybrid offspring ratio from inter-niche mating. Therefore, *Coff* + *Hr* is the total offspring ratio produced from inter-niche mating and can serve as a measure of gene flow or RI between the niches. In our models, *Hr* represents the hybrid offspring loss that drives adaptive coupling. Coupling of additional adaptive barriers is impossible when *Hr* is zero. For any given value of *f*, the ratio between *Hr* and *Coff* is fixed. Therefore, *Hr* can be used as a relative measure of *Coff* + *Hr*, indicating the amount of gene flow as measured by the total ratio of offspring produced through inter-niche mating. However, as the value of *f* increases toward 0.5, the value of *Hr* gradually decreases, reaching zero when *f* = 0.5. In such a case, *Hr* is no longer a reliable indicator of gene flow between the two niches. Instead, when the value of *f* is high, the total ratio of inter-niche offspring, *Coff* + *Hr*, provides a better gauge of the degree of gene flow.

Fig 12a shows a small population of chromosomal inversion (*Ak* = 0.01) with *Fk*1 = *Fk*2 = 0.5 successfully invades and becomes fixed in niche *A*. This makes sense because CI is a recombination suppressor. The *Ak* ecotype does not suffer any hybrid loss in its inter-niche mating with the *B* ecotype. Therefore the *Ak* genotype has a fitness advantage over its noninverted niche-*A* cohorts that suffer hybrid loss in their inter-niche mating.

However, as *Ak* becomes fixed in niche *A*, no more hybrids are produced between the mating of *A* and *B* ecotypes. The effective value of *f* in our mathematical model in Fig M3 becomes 0.5, and the *Ax*/*Bx* mating-bias phase portrait becomes divergent. Sexual selection drives the most prevalent mating-bias allele in the population to fixation and completely eliminates the less prevalent mating-bias allele. Even though there is no gene flow between the ecological gene loci in the chromosomally inverted regions, the rest of the genome acts like a panmictic population without any RI to impede gene flow. Because only the locally adaptive gene loci are reproductively isolated, they may be detected as islands of divergence when comparing genomes between niche populations.

Next, let us consider what happens when a CI captures only a partial number of the ecological gene loci. For heterokaryotype mating within the same niche—e.g., a niche-*A* ecotype with CI mating with a niche-*A* ecotype without CI—*Fk*1 is always 0.5 (discounting the rare occurrence of single crossovers in the inverted region) because both parents have the same ecological genotypes. However, for inter-niche heterokaryotype mating, *Fk*2 will be less than 0.5 because the niche-*A* and niche-*B* ecotypes are different. For instance, suppose a 3-gene-locus niche-*A* ecotype has the genotype [*a*1, *a*2, *a*3] and a niche-*B* ecotype has the genotype [*b*1, *b*2, *b*3]. If a CI captures only the first two loci, then the parents will produce offspring with the following genotype permutations: 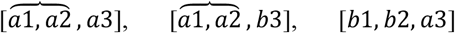,and [*b*1, *b*2, *b*3], where the curly brackets indicate the alleles that are linked by the CI. Assuming random assortment of the alleles at each gene locus, inter-niche heterokaryotype mating will produce 50% hybrids, and therefore *Fk*2 is reduced to 0.25.

Figs 13a and 13b show how the *Ax*/*Bx* phase portrait gradually moves from divergence to convergence as we keep *Fk*1 = 0.5 and gradually decrease *Fk*2 towards *f*. When *Fk*2 = *f*, the *Ax*/*Bx* fixed point is the same as that before the appearance of the CI. When *Fk*2 is less than *f* (see Fig 13c), the CI fails to invade, and the *Ax*/*Bx* phase portrait remains unchanged.

**Fig 13a.**
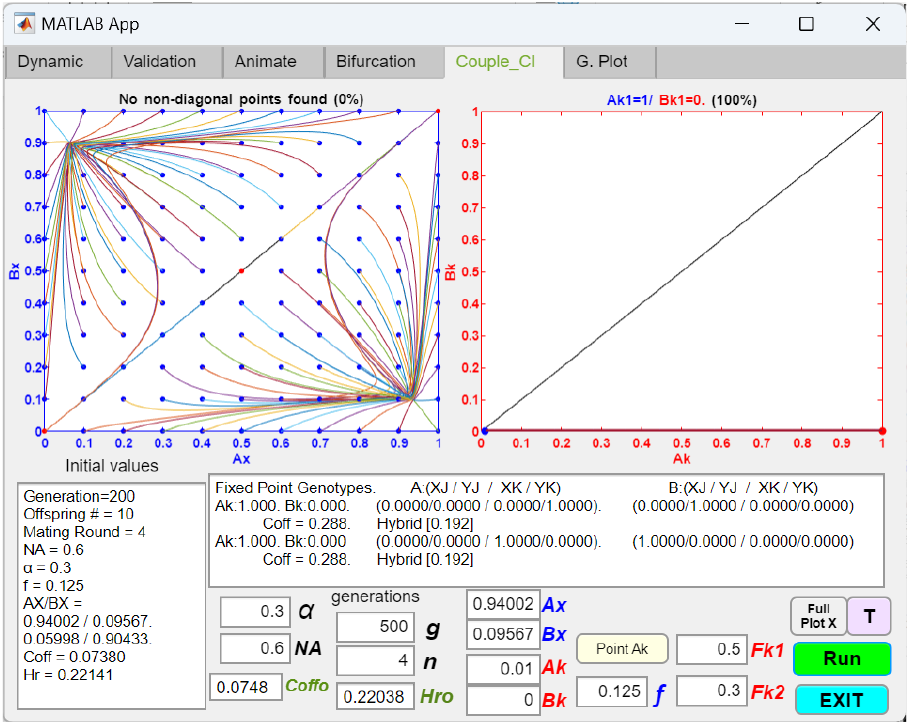
Coupling of an initial two-allele premating barrier with a chromosomal inversion that captures a partial number of the locally adaptive ecological alleles, causing divergence. Changing the value of *Fk*2 in Figure 12a from 0.5 to 0.3, while keeping the rest of the parametric values the same, still allows the CI to invade and cause the *Ax*/*Bx* phase portrait to become divergent.

**Fig 13b.**
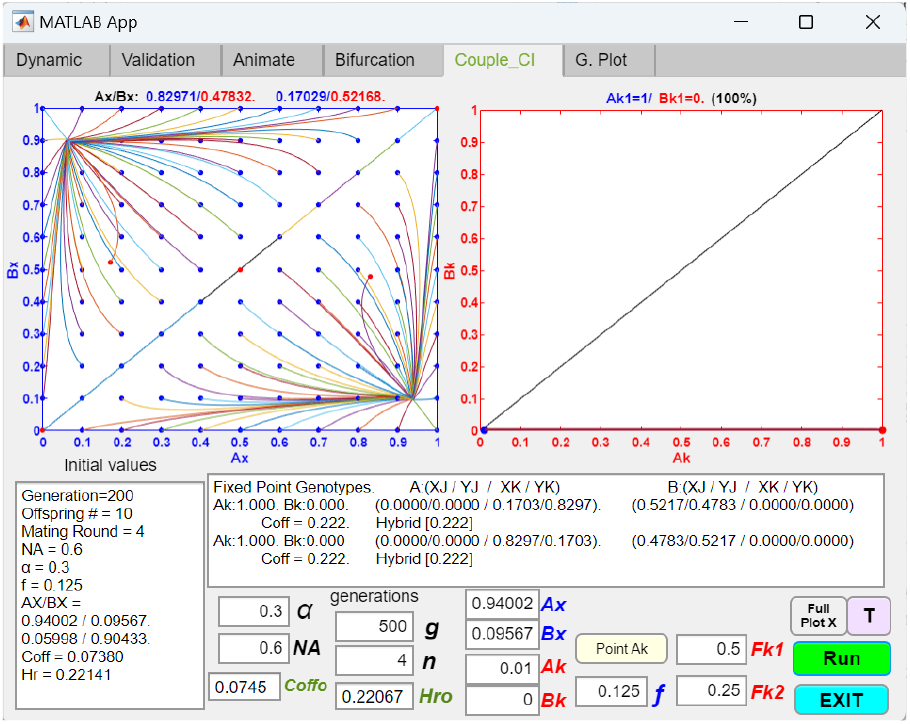
Coupling of an initial two-allele premating barrier with a chromosomal inversion that captures a partial number of the locally adaptive ecological alleles, producing reduced RI. Further reducing the value of *Fk*2 in Fig 13a to 0.25 allows the CI to invade and shift the fixed points in the *Ax*/*Bx* phase portrait away from the upper left and lower right corners, resulting in reduced overall RI. The new fixed points are shown as red dots in the phase portraits.

**Fig 13c.**
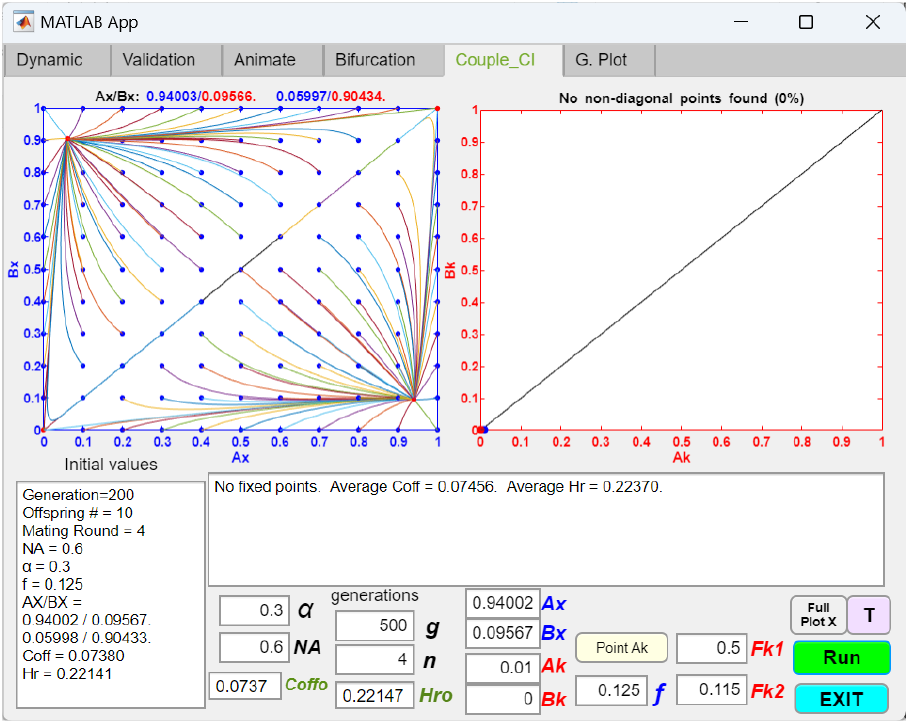
Coupling of an initial two-allele premating barrier with a chromosomal inversion that captures a partial number of the locally adaptive ecological alleles, showing the CI failing to invade. If the value of *Fk*2 in Figure 13a is less than the value of *f*, the CI fails to invade, and the fixed points in the *Ax*/*Bx* phase portrait remain unchanged.

Fig 14a shows that in a single, homogeneous population (*NA* = 1, *NB* = 0, and *Ax* = 1), a CI capturing locally adaptive ecological alleles cannot invade without having additional fitness advantage. This is because the rare occurrence of a single crossover in its inverted region during heterokaryotype mating produces nonviable offspring, which reduces the fitness of the CI mutant relative to other individuals in the population without the CI. This effect is modeled in Fig 14a by setting *Fk*1 = 0.49. Any initial *Ak* mutant population less than 0.5 will be driven to extinction (In the Fig 14a example, *Ak* = 0.4). However, if *Ak* is greater than 0.5, it will be driven to fixation. Once *Ak* becomes the predominant genotype in the population, it becomes the norm, and individuals from the original population (i.e., the previous noninverted genotypes) are now considered mutants with a CI attempting to invade.

**Fig 14a.**
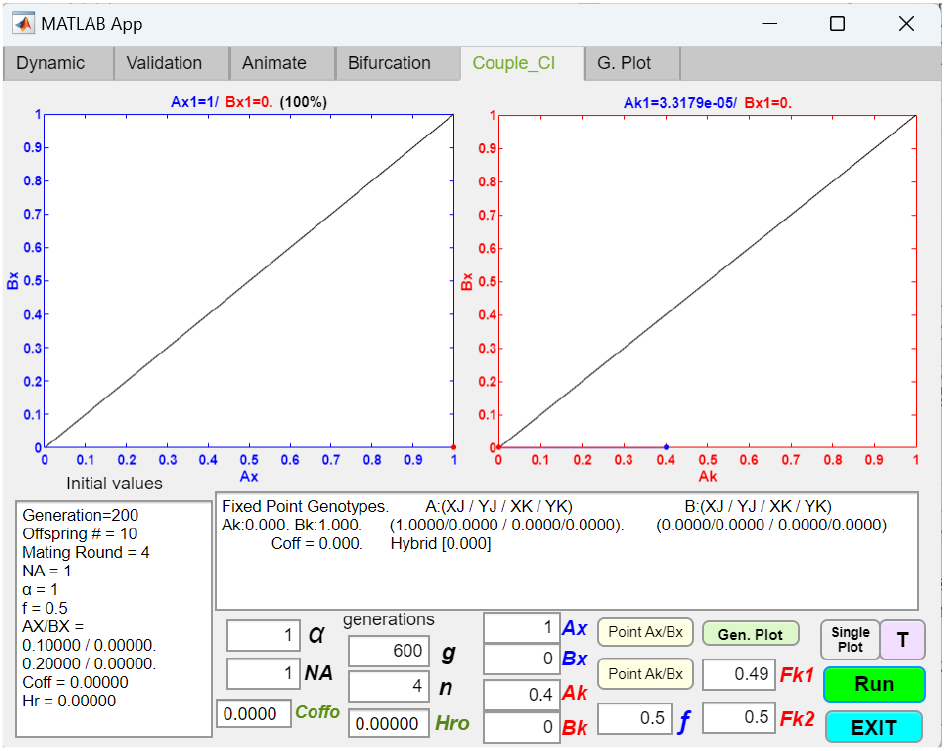
In a single-niche ecosystem, a mutant chromosomal inversion capturing locally adaptive ecological alleles may be at a fitness disadvantage and cannot invade due to offspring loss from single-crossover events in heterokaryotype mating. *Fk*1 is set to 0.49 to model the effect of nonviable offspring loss caused by single crossover events in heterokaryotype mating. With a starting population ratio of *Ak* = 0.4, the CI fails to invade.

**Fig 14b.**
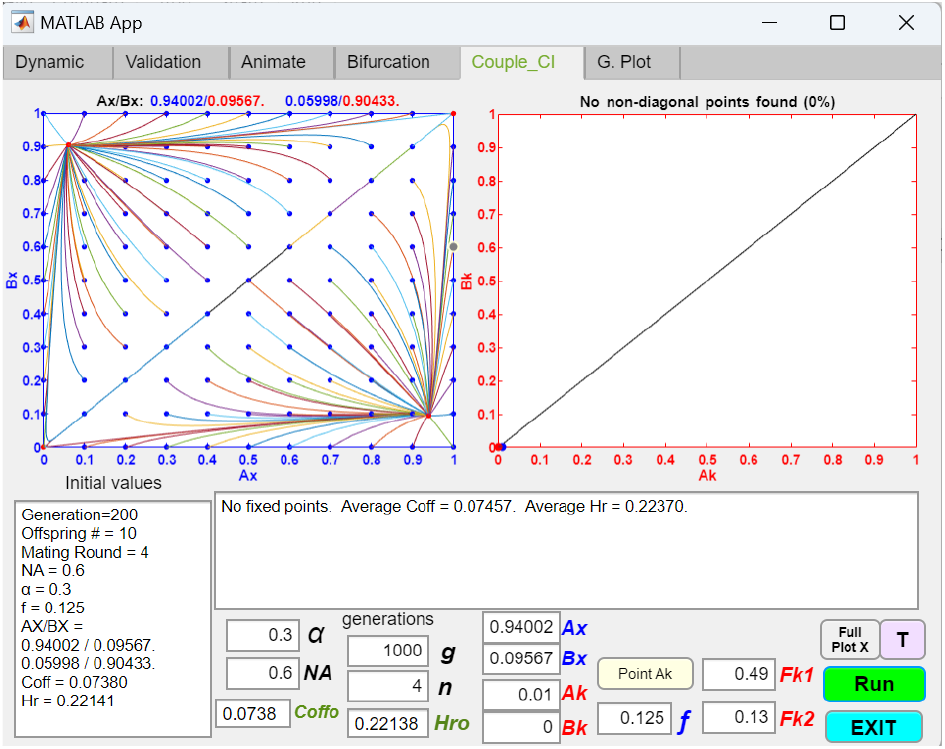
A mutant chromosomal inversion capturing locally adaptive ecological alleles may have a net fitness disadvantage and cannot invade due to offspring loss from single-crossover events in heterokaryotype mating. The value of *Fk*2 in Fig 14a is reduced from 0.5 to 0.13, signifying that fewer locally-adaptive alleles in niche *A* are captured by the invading CI. The fitness advantage of capturing locally adaptive alleles (*Fk*2 = 0.13) is not great enough to offset the fitness disadvantage of offspring loss in intra-niche heterokaryotype mating (*Fk*1 = 0.49). Therefore, the mutant CI cannot invade.

**Fig 14c.**
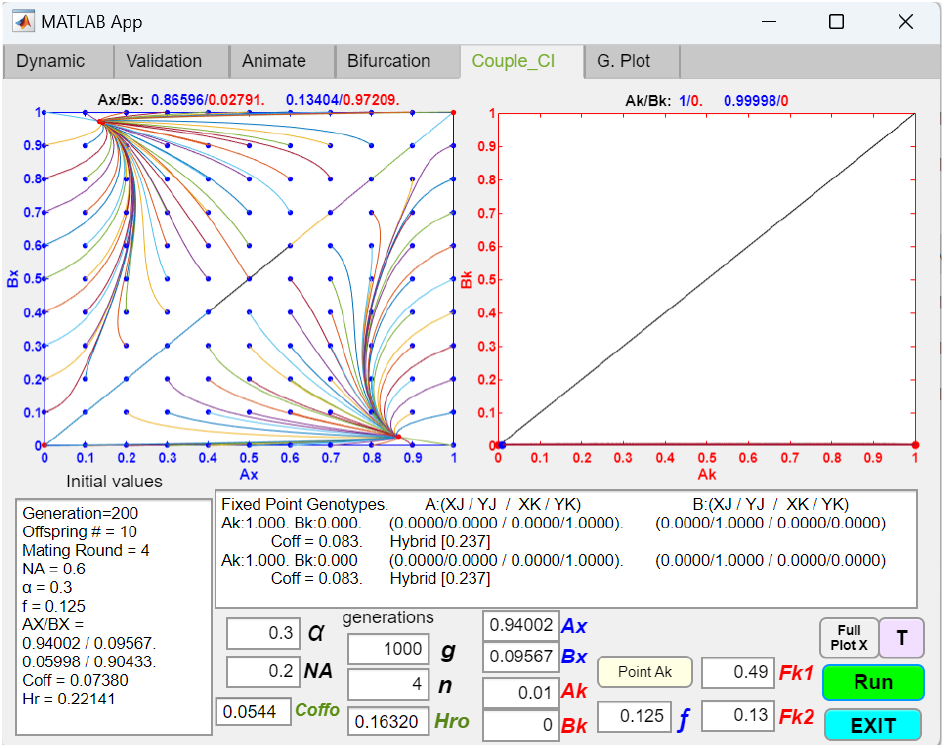
A mutant chromosomal inversion capturing locally adaptive ecological alleles may achieve a net fitness advantage and invade, despite offspring loss from intra-niche heterokaryotype mating, if the niche size *NA* is small. The value of *NA* in Fig 14b is reduced from 0.6 to 0.2, enabling the mutant CI to invade. When *NA* is small, the mutant CI in niche *A* has mostly inter-niche mating rather than intra-niche mating. Therefore, the fitness advantage of reduced offspring loss from inter-niche mating (compared to same-niche, noninverted cohorts) offsets the fitness disadvantage of offspring loss from intra-niche heterokaryotype mating.

The examples in Figs 14b and 14c illustrate that a reduction in the niche size *NA* may help a mutant CI overcome the invasion resistance arising from offspring loss in intra-niche heterokaryotype mating. When *NA* is small relative to *NB*, a mutant CI capturing locally adaptive ecological alleles in niche *A* primarily engages in inter-niche mating rather than intra-niche mating. Consequently, the CI‘s fitness advantage, gained through reduced offspring loss in inter-niche heterokaryotype mating (compared to the offspring loss experienced by its noninverted same-niche counterparts) may counterbalance its fitness disadvantage from offspring loss in intra-niche heterokaryotype mating, giving the CI mutant a net fitness advantage to invade and proliferate.

##### 2.2.2. Chromosomal inversions that capture both ecological alleles and mating-bias alleles

Fig 15 shows the phase portrait solutions from a GUI application investigating the coupling of an initial two-allele premating barrier with a chromosomal inversion that captures both adaptive ecological alleles and a mating-bias allele. *Ak* denotes the presence of a CI that includes both locally adaptive ecological alleles in niche *A* and the mating-bias allele, *X*, in the genotype of a niche-*A* individual. By definition, *Bk* is always zero.

**Fig 15.**
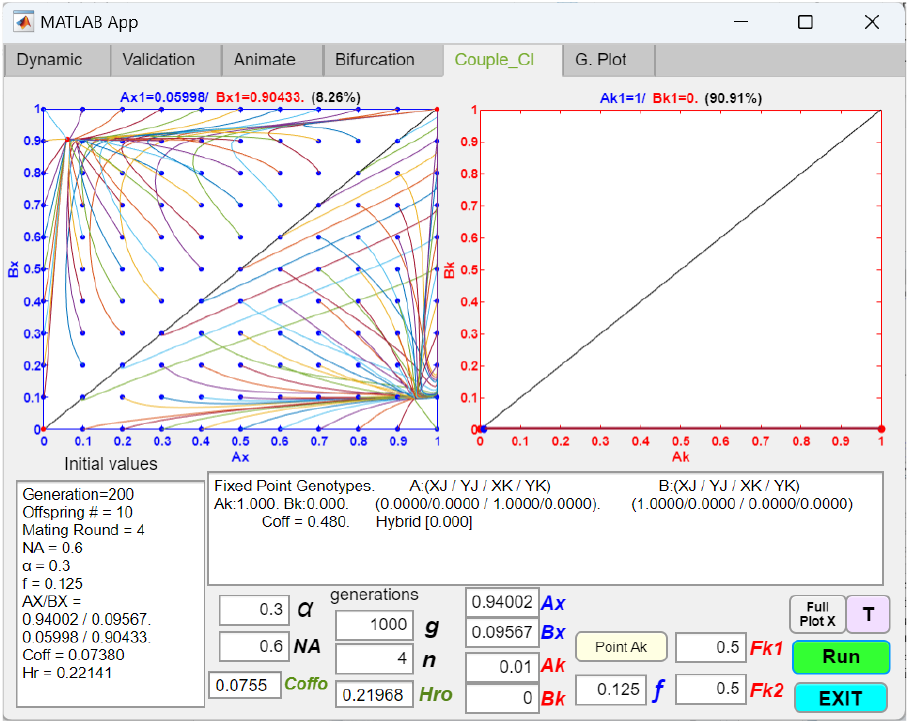
Coupling of an initial two-allele premating barrier with a chromosomal inversion that captures all locally-adaptive ecological alleles and a mating-bias allele, resulting in the elimination of premating RI. The *Ak* genotype signifies the presence of a CI that captures both the locally adaptive ecological alleles and the prevalent mating-bias allele, *X*, in niche *A* after the establishment of an initial premating barrier. The remaining parameters are defined in the same way as in Fig 12a. As shown, a CI with an initial population ratio of *Ak* = 0.01 and *Fk*1 = *Fk*2 = 0.5 can invade and become fixed in niche *A*. Because the population ratio *NA* is greater than the population ratio *NB*, the phase portrait becomes divergent, and sexual selection eventually drives the *X* allele to fixation and eliminates the *Y* allele in the sympatric population. The matching and unit-offspring tables used in the GUI application are included in the Appendix (Tables 8 and 10).

As shown in Fig 15, when *Fk*1 = 0.5 and *Fk*2 = 0.5, no ecological hybrids are produced. The effective value of *f* in our mathematical model in Fig M3 is 0.5, and all individuals in the sympatric population can freely mate without any barriers. Because *Ak* is able to invade and become fixed in niche *A*, the niche-*A* ecotypes can only have the *X* mating bias allele in their genotypes. In Fig 15, because *NA* (0.6) is larger than *NB* (0.4), the phase portrait becomes divergent, and sexual selection drives the more prevalent mating bias allele *X* to fixation and eliminates the less prevalent *Y* allele in the population.

**Table 10.** Unit offspring tables for the coupling of a chromosomal inversion that captures locally adaptive ecological alleles in niche *A* and the mating-bias allele, *X*. The *K* allele denotes the presence of a CI that captures locally adaptive alleles in niche *A* and the mating-bias allele, *X*, in the genotype of a niche-*A* individual. The *J* allele denotes the absence of the *K* allele. The rest of the parametric variables are the same as those in Fig 9. By definition, it is impossible to have the *BK* and *AYK* genotypes; therefore, all cells associated with the *BK* and *AYK* genotypes are grayed out.

|  | A XJ 1 | A YJ 2 | A XK 3 | A YK 4 |
| --- | --- | --- | --- | --- |
| A | XJ $1(1)$ | XJ $1(1/2)$ | AXJ | |
| XJ | YJ | YJ $2(1/2)$ | YJ $[F_{k1}]$ | |
| 1 | XK | XK | AKX |  |
|  | YK | YK | YK |  |
| A | XJ $1(1/2)$ | XJ | XJ | |
| YJ | YJ $2(1/2)$ | YJ $2(1)$ | AYJ $[F_{k1}]$ | |
| 2 | XK | XK | AKX |  |
|  | YK | YK | YK |  |
| A | AXJ | XJ | XJ |  |
| XK | YJ $[F_{k1}]$ | AYJ $[F_{k1}]$ | YJ | |
| 3 | AKX | AKX | XK $3(1)$ | |
|  | YK | YK | YK |  |
| A |  |  |  |  |
| YK |  |  |  |  |
| 4 |  |  |  |  |

|  | B XJ 1 | B YJ 2 | B XK 3 | B YK 4 |
| --- | --- | --- | --- | --- |
| B | XJ $1(1)$ | XJ $1(1/2)$ | | |
| XJ | YJ | YJ $2(1/2)$ | | |
| 1 | XK | XK |  |  |
|  | YK | YK |  |  |
| B | XJ $1(1/2)$ | XJ | | |
| YJ | YJ $2(1/2)$ | YJ $2(1)$ | | |
| 2 | XK | XK |  |  |
|  | YK | YK |  |  |
| B |  |  |  |  |
| XK |  |  |  |  |
| 3 |  |  |  |  |
| B |  |  |  |  |
| YK |  |  |  |  |
| 4 |  |  |  |  |
| A | XJ $1(1) \times f[A B]$ | XJ $1(1/2) \times f[A B]$ | | |
| XJ | YJ | YJ $2(1/2) \times f[A B]$ | | |
| 1 | XK | XK |  |  |
|  | YK | YK |  |  |
| A | XJ $1(1/2) \times f[A B]$ | XJ | | |
| YJ | YJ $2(1/2) \times f[A B]$ | YJ $2(1) \times f[A B]$ | | |
| 2 | XK | XK |  |  |
|  | YK | YK |  |  |
| A | BXJ | XJ |  |  |
| XK | YJ $[F_{k2}]$ | BYJ $[F_{k2}]$ | | |
| 3 | AKX | AKX |  |  |
|  | YK | YK |  |  |
| A |  |  |  |  |
| YK |  |  |  |  |
| 4 |  |  |  |  |

|  | A XJ 1 | A YJ 2 | A XK 3 | A YK 4 |
| --- | --- | --- | --- | --- |
| B | XJ $1(1) \times f[A B]$ | XJ $1(1/2) \times f[A B]$ | BXJ | |
| XJ | YJ | YJ $2(1/2) \times f[A B]$ | YJ $[F_{k2}]$ | |
| 1 | XK | XK | AKX |  |
|  | YK | YK | YK |  |
| B | XJ $1(1/2) \times f[A B]$ | XJ | XJ | |
| YJ | YJ $2(1/2) \times f[A B]$ | YJ $2(1) \times f[A B]$ | BYJ $[F_{k2}]$ | |
| 2 | XK | XK | AKX |  |
|  | YK | YK | YK |  |
| B |  |  |  |  |
| XK |  |  |  |  |
| 3 |  |  |  |  |
| B |  |  |  |  |
| YK |  |  |  |  |
| 4 |  |  |  |  |

Conversely, in Fig 16a and Fig 16b, *NB* (0.6) is larger than *NA* (0.4). Normally, in such situations, the more numerous *Y* mating-bias allele in niche *B* would drive the less numerous *X* allele in the panmictic population to extinction. However, because the chromosomal inversion *Ak* is fixed in niche *A*, the niche-*A* ecotypes can only be associated with the *X* allele. Consequently, the *Y* alleles from niche *B* are unable to invade the niche-*A* ecotypes, and strong mating-bias allele polymorphism (*Ax* = 1 and *Bx* = 0) emerges in the phase portrait. This creates premating RI between the two niche ecotypes and results in reduced gene flow between the entire genomes of the niche-*A* and niche-*B* individuals.

**Fig 16a.**
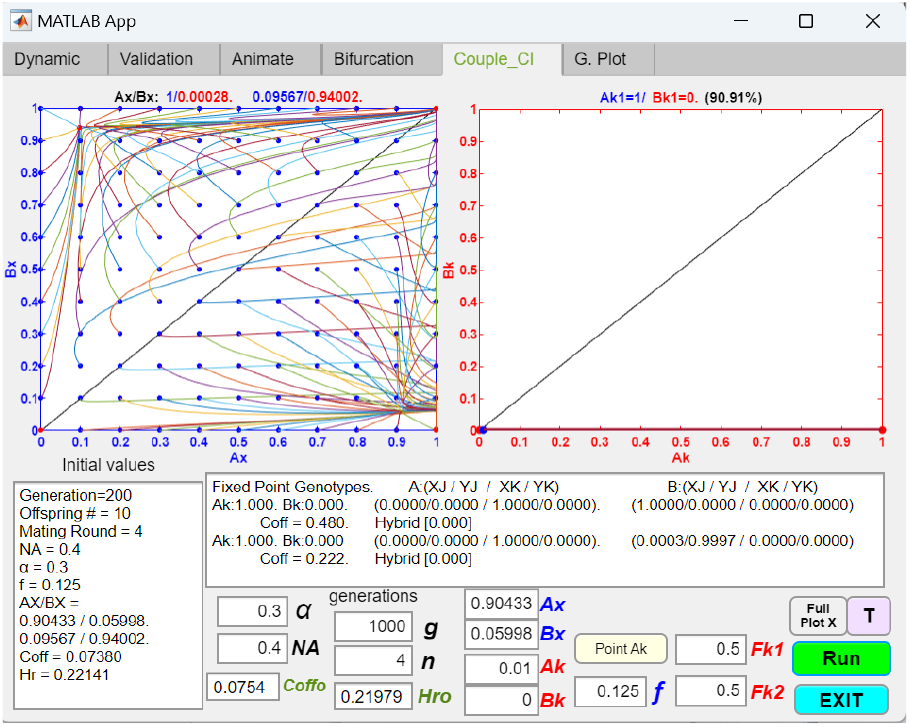
Coupling of an initial two-allele premating barrier with a chromosomal inversion that captures all locally-adaptive ecological alleles and a mating-bias allele, resulting in strong premating RI. If the value of *NA* in Fig 15 is changed to 0.4 (*NB* = 0.6) while keeping the rest of the parametric values the same, the *Ax*/*Bx* phase portrait converges to a fixed-point polymorphism at *Ax* = 1 and *Bx* = 0 (shown as a red dot).

**Fig 16b.**
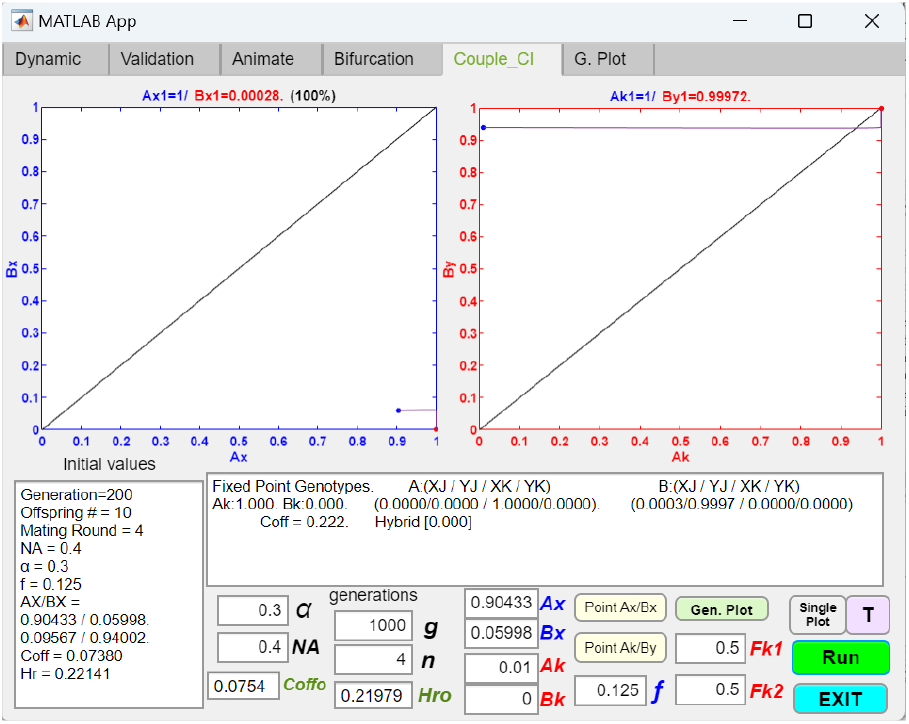
Coupling of an initial two-allele premating barrier with a chromosomal inversion that captures all locally-adaptive ecological alleles and a mating-bias allele, resulting in strong premating RI (single line plot). Using the same parametric values as in Fig 16a, the single-line trajectories in the *Ax*/*Bx* and *Ak*/*By* phase portraits show that the initial *Ax*/*Bx* fixed point shifts to the lower right corner of the phase portrait following the successful invasion of the CI.

However, even when *NB* > *NA*, the fixed-point polymorphisms in Fig 16a and 16b can still be destroyed if the mating bias is greatly reduced (i.e., large values of *α*) and *NA* is close to 0.5, leaving only the *X* allele in the system (Fig 17a). This occurs because large values of *α* and *NA* increase inter-niche matings and reduce intra-niche matings for niche-*B* individuals. Consequently, sexual selection against the *X* alleles in niche *B* is reduced, while sexual selection against the *Y* alleles in niche *B* is increased, which could result in the elimination of the *Y* alleles. Eliminating sexual selection by increasing the number of matching rounds, *n*, causes a stable polymorphism of *X* and *Y* alleles to exist in niche *B* (Fig 17b).

**Fig 17a.**
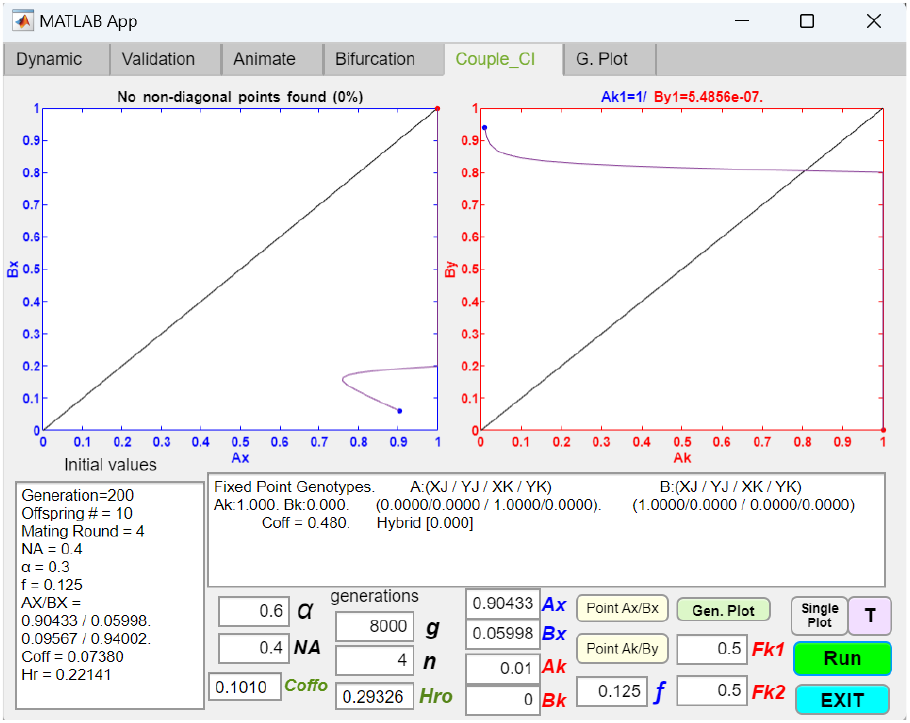
When coupling a chromosomal inversion that captures all locally adaptive ecological alleles and a mating-bias allele, a large *α* value can still eliminate the premating RI even with *NB* > *NA*. Single-line vector trajectories in the phase portraits show that when the value of *α* in Fig 16b is increased from 0.3 to 0.6, the initial *Ax*/*Bx* fixed point is eliminated following the invasion of the CI.

**Fig 17b.**
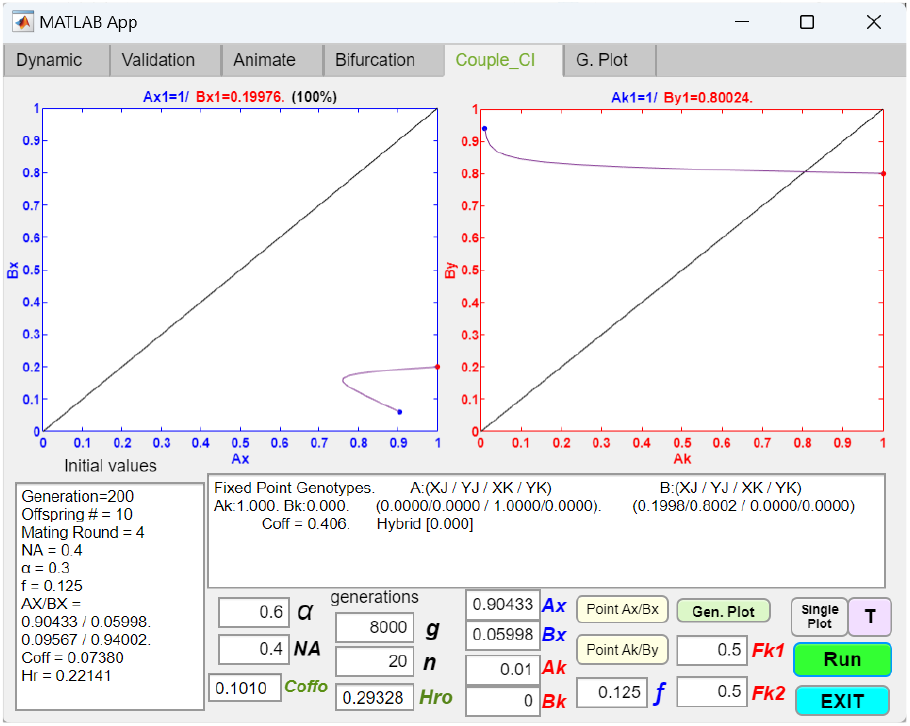
When coupling a chromosomal inversion that captures all locally adaptive ecological alleles and a mating-bias allele, increasing the number of matching rounds, *n*, can counter the effect of a large *α* and restore *Ax*/*Bx* fixed-point polymorphism. Reducing the assortative mating cost by increasing the number of matching rounds, *n*, to 20 in Fig 17a restores the *Ax*/*Bx* fixed-point polymorphism and premating RI.

If the chromosomal inversion *Ak* captures only a portion of the total number of locally adaptive ecological alleles in niche *A*, then *Fk*2 will be less than 0.5. Once again, this results in the production of less-fit, ecologically nonviable hybrids through inter-niche heterokaryotype mating. The effective value of *f* in the mathematical model in Fig M3 is now less than 0.5, restoring the effect of disruptive ecological selection in the system. Because all niche-*A* individuals have the same ecological genotypes, *Fk*1 remains close to 0.5.

In such a situation, when *Fk*1 = 0.5 and *Fk*2 < 0.5, coupling with a CI that also captures a favorable mating-bias allele can strengthen premating RI regardless of the difference in niche sizes. This is illustrated in Fig 18a, where *NA* = 0.6, *NB* = 0.4, and in Fig 18b, where *NA* = 0.4 and *NB* = 0.6. In both cases, a small mutant population *Ak* = 0.01 is able to invade and shift the *Ax*/*Bx* fixed point toward the lower right corner, where *Ax* = 1 and *Bx* = 0, thereby increasing the overall premating RI between the two niches.

**Fig 18a.**
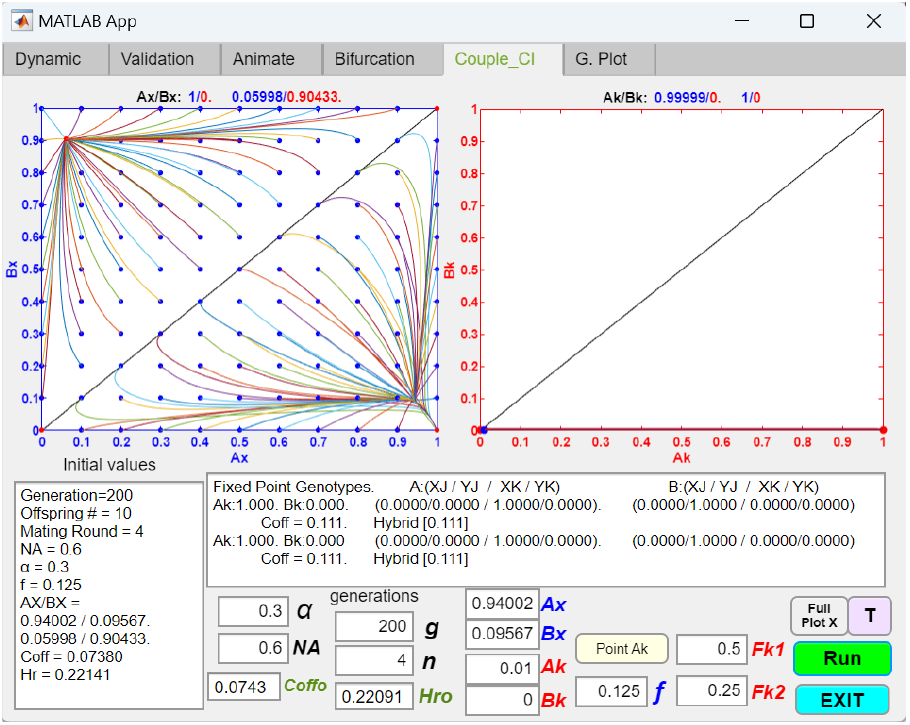
Coupling an initial two-allele premating barrier with a chromosomal inversion that captures a partial set of the locally adaptive ecological alleles in niche *A*, along with the mating-bias allele *X*, results in strong premating RI even when *NA* > *NB*. The value of *Fk*2 in Fig 15 is changed from 0.5 to 0.25 to indicate that the CI captures only a partial number of the adaptive alleles in niche *A*, while keeping the rest of the parametric values the same. The invasion of the CI causes the *Ax*/*Bx* fixed point to shift to the lower right corner, where *Ax* = 1 and *Bx* = 0.

**Fig 18b.**
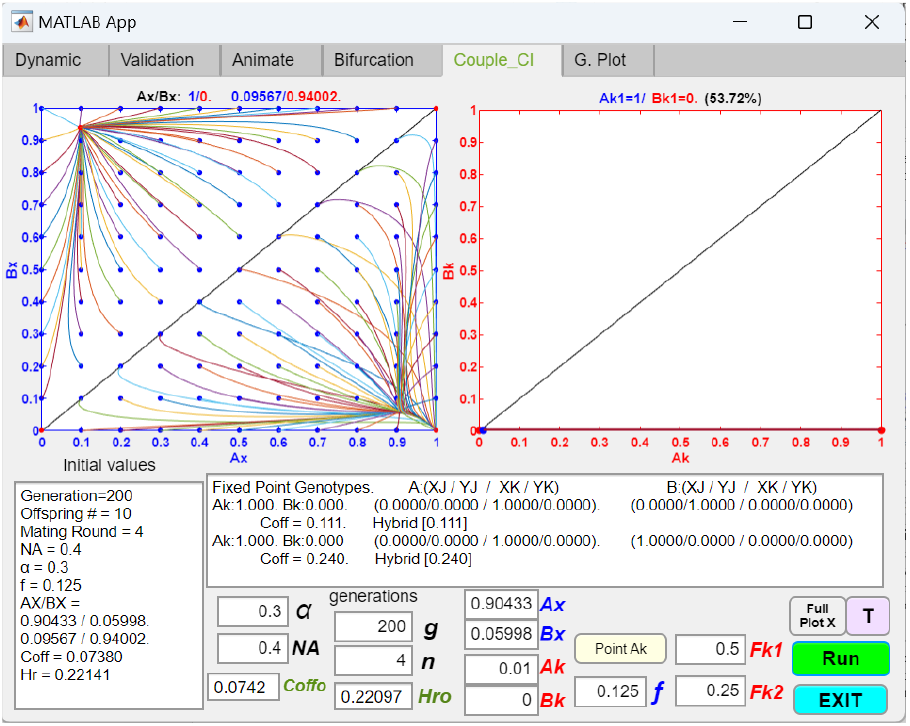
Coupling an initial two-allele premating barrier with a chromosomal inversion that captures a partial set of the locally adaptive ecological alleles in niche *A*, along with the mating-bias allele *X*, results in strong premating RI when *NB* > *NA*. The value of *NA* in Fig 18a is changed from 0.6 to 0.4, while keeping the rest of the parametric values the same. The invasion of the CI causes the *Ax*/*Bx* fixed point to shift to the lower right corner, where *Ax* = 1 and *Bx* = 0.

Fig 19 shows that even if the preexisting mating-bias system is divergent, i.e., when *α* = 0.4, *f* = 0.3, and no *Ax*/*Bx* fixed point exists, an *Ak* with a value of *Fk*2 greater than *f* can invade and create a fixed-point mating-bias allele polymorphism. However, when the rest of the parameters in Fig 19 are kept the same, *Ak* fails to invade if *Fk*2 is less than *f*. This result suggests that if a CI mutation can fortuitously capture a set of locally advantageous ecological alleles and the appropriate mating-bias allele, it could invade and convert a divergent mating-bias system into a convergent one. The challenge is that when the system is divergent, the likelihood of capturing this advantageous set of alleles is much lower than when the system is already convergent with a concentrated assortment of locally adaptive alleles in the niches.

**Fig 19.**
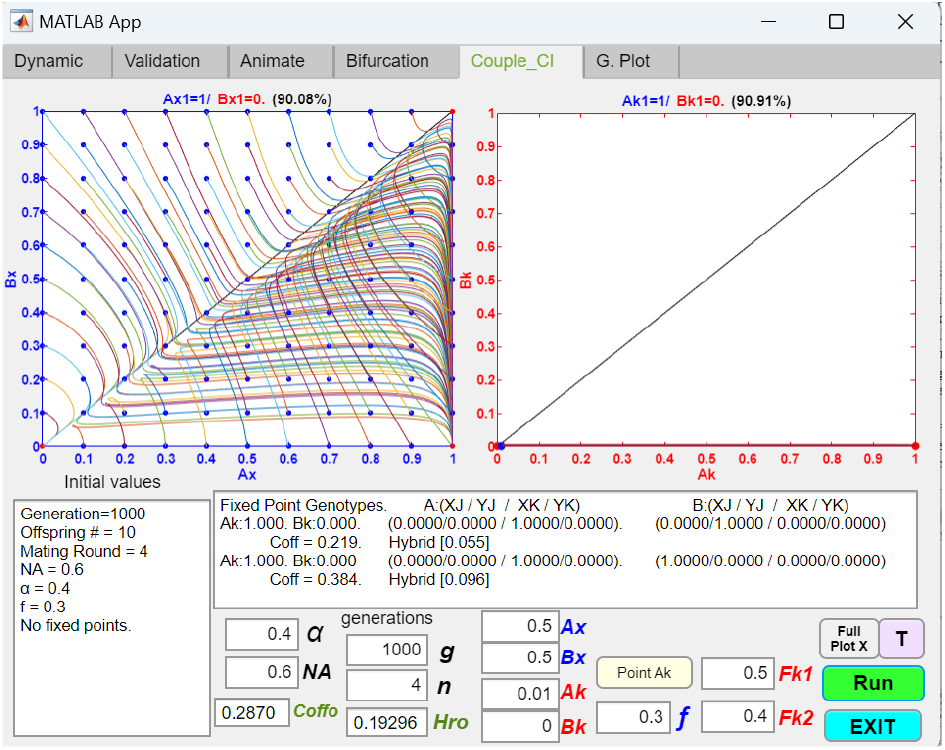
Coupling an initially divergent two-allele premating system with a chromosomal inversion that captures a partial set of the locally adaptive ecological alleles in niche *A*, along with the mating-bias allele *X*, results in system convergence and premating RI. The initial two-allele mating-bias system is divergent with parametric values *f* = 0.3 and *α* = 0.4. However, a CI that captures a partial number of the locally adaptive alleles in niche *A* (*Fk*2 = 0.4), along with the mating-bias allele *X*, can invade and convert the initially divergent mating-bias system into a convergent one, with a fixed point at *Ax* = 1 and *Bx* = 0.

So far, we have established that if *Fk*2 is greater than *f*, then a CI mutant has a fitness advantage over its same-niche noninverted cohorts in inter-niche mating and can invade. Interestingly, linkage of locally adaptive ecological alleles with mating-bias alleles also enables a CI to invade even when *Fk*2 is less than *f*. In Figs 18a and 18b, if the parametric values are kept the same except for *Fk*2, the mutant CI, *Ak*, can invade if *Fk*2 > 0.08 but not if *Fk*2 < 0.07, despite having an *f* value of 0.125.

These findings can be explained by the fact that, in order to invade successfully, a mutant CI must have a net selection advantage over the rest of the native, noninverted population in the same niche. A CI can link adaptive ecological alleles in its inverted region and prevent their breakdown by recombination. Consequently, when *Fk*2 > *f*, such a mutant CI has a fitness advantage in inter-niche mating (because it can recover a higher ratio of offspring) over its noninverted cohorts in the same niche. Because of this fitness advantage, it can invade successfully. As shown by the examples in Figs 18a and 18b, when a mutant CI is able to link adaptive ecological alleles in niche *A* with the mating-bias allele *X* and when *Fk*2 is less than 0.5, the advantageous mutant CI can carry the mating-bias allele *X* with it to fixation and establish premating RI between niche ecotypes. Without capturing a mating-bias allele, a mutant CI of just ecologically alleles cannot invade for *Fk*2 < *f*. However, linkage to a mating-bias allele allows a mutant CI to derive additional selection advantage even if *Fk*2 < *f*. This is because, in late-stage sympatric speciation, when there is already assortment of mating-bias alleles in the two niches, a mutant CI that can only have the niche-predominant mating-bias allele associated with it (i.e., the *X* allele in niche *A* in the Figs 18a and 18b examples) has reduced inter-niche mating compared to its noninverted cohorts that still possess a polymorphism of mating-bias (*X* and *Y*) alleles. This reduced inter-niche mating can mitigate the hybrid loss of the mutant CI caused by its lower *Fk*2 value. Eventually, as the value of *Fk*2 is further reduced, the hybrid loss becomes too significant even with the reduced inter-niche mating, the mutant CI suffers a fitness disadvantage compared to its same-niche noninverted cohorts, and invasion becomes impossible.

Therefore, a CI that captures favorable ecological and mating-bias alleles can produce not only premating RI of the entire genome but also increase the degree of hybrid inviability it can generate and still invade—i.e., by making invasion possible even when *Fk*2 < *f*. We can call this lower limit of *Fk*2 the *“*lower invasion threshold,*”* and the difference between *f* and this lower limit of *Fk*2 the *“*invasion buffer range.*”*

As expected, reducing the mating-bias value *α* (i.e., increasing the mating-bias) extends the lower limit to which *Fk*2 can be reduced below *f* and still permit invasion, i.e., it increases the invasion buffer range. However, as *α* tends toward zero, invasion becomes more difficult because there is not enough hybrid loss to drive the invasion of the mutant CI. On the other hand, increasing the value of *f* decreases the invasion buffer range because the noninverted genotypes suffer less hybrid loss from inter-niche mating. In the limit, when *f* = 0.5, the invasion buffer range is zero because there is no hybrid loss from inter-niche mating. Therefore, the invasion of a beneficial CI is most favored in late-stage sympatric speciation when the values of *α* and *f* are low, indicating strong mating-bias and hybrid loss.

Having established the lower limit of *Fk*2 for a mutant CI to invade, we next examine if there is an upper limit of *Fk*2 beyond which the CI is no longer able to establish premating RI. In order for a CI that captures favorable ecological and mating-bias alleles to effectively produce mating-bias allele polymorphism and premating RI, its *Fk*2 should be less than 0.5 so that disruptive ecological selection, driven by hybrid loss, can be reestablished in the system. This hybrid loss can then help increase the assortment of mating-bias alleles by eliminating the less prevalent mating-bias allele in each niche. However, there appears to be an upper limit of *Fk*2 above which the CI mutant can invade but cannot establish mating-bias allele polymorphism, i.e., the system remains divergent. This likely happens because a very high value of *FK*2 decreases the effective value of *f* and hybrid loss that are necessary for the *Ax*/*Bx* phase portrait to converge. Given the parametric values shown in Fig 18a, this upper limit is *Fk*2 > 0.492. If *α* is increased from 0.3 to 0.5, this upper limit of *Fk*2 rises to 0.495. If *α* is decreased to 0.1, this upper limit of *Fk*2 decreases to 0.489.

The value of *FK*1 also affects the invasion dynamics of a mutant CI. In intra-niche mating, if *Fk*1 is less than 0.5—e.g., due to offspring loss caused by the rare occurrence of single crossover in the inverted region—this could create a fitness disadvantage for a mutant CI and hinder its invasion (see Fig 14a example).

Fig 20 shows the results of a GUI application that investigates the reversibility of a coupled CI when external, environmental disruptive ecological selection is relaxed. In Fig 18a, a mutant CI linking ecological alleles in niche *A* and mating-bias allele *X* is able to invade, become fixed, and couple with an initial premating barrier to increase the overall RI. We can determine the reversibility of the coupled CI by checking whether mutant populations in the vicinities of the established fixed points can invade or go extinct.

**Fig 20.**
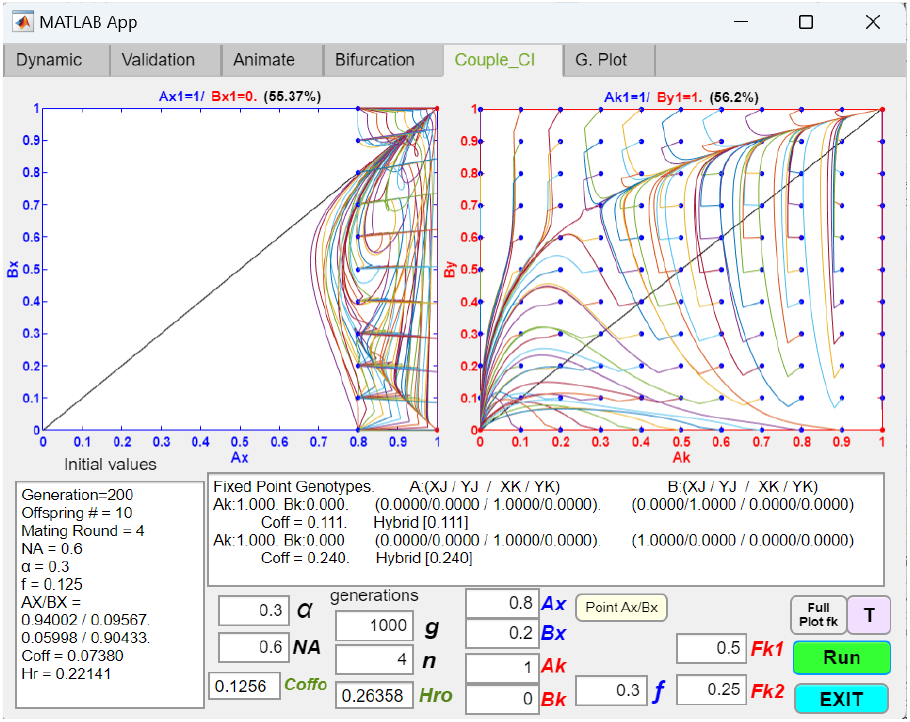
Fixation of a coupled chromosomal inversion that captures a partial set of the locally adaptive ecological alleles in niche *A*, along with the mating-bias allele *X*, showing resistance to invasion by noninverted genotypes. In the Fig 18a example, once the CI became fixed (*Ak* = 1) and the *Ax*/*Bx* fixed point moved to *Ax* = 1 and *Bx* = 0, a small population of noninverted genotypes cannot invade after *f* is increased from 0.125 to 0.3 and disruptive ecological selection is reduced. The *Ak*/*By* phase portrait (plotted against Ax=0.8 and Bx=0.2) shows that populations in the vicinity of the fixed point, *Ak* = 1 and *By* = 1, converge back to the fixed point, indicating that the fixed CI is resistant to invasion by noninverted mutants. Similarly, a protective zone exists around the fixed point *Ax* = 1/*Bx* = 0 that prevents invasion due to fluctuations in Ax/Bx population ratios when disruptive ecological selection is reduced.

Fig 20 adopts the example in Fig 18a, except that here the value of *f* in Fig 18a is increased from 0.125 to 0.3, signifying a decrease in the strength of disruptive ecological selection. The phase portraits in Fig 20 show that, given the post-invasion system parameters with fixed points at *Ax*/*Bx* = 1/0 and *Ak* = 1, the coupled systems cannot be invaded by a small population of noninverted mutants in the vicinities of the fixed points. As shown in the *Ak*/*By* phase portrait, all the polymorphic starting populations surrounding the coordinate point *Ak* = 1, *By* = 1 converge to that point. In fact, the results in Fig 20 demonstrate that for starting *Ax*/*Bx* values that meet the criteria *Ax* > 0.8 and *Bx* < 0.2 (or *By* > 0.8), the *Ak*/*By* phase portrait can be partitioned into two regions. One region converges to the upper right corner where *Ak* = *By* = 1, signifying invasion resistance and the persistence of premating RI. The other region converges to the origin where *Ak* = *By* = 0, signifying the elimination of the *Ak* genotype and its CI by the invasion of noninverted mutants and the elimination of premating RI.

In Fig 20, the larger the region that converges to the upper right corner of the *Ak*/*By* phase portrait, the more resistant the CI population is against invasion by noninverted mutants. In general, decreasing the value of *α* (increasing the mating bias) and increasing the value of *n* (decreasing the assortative mating cost) can expand this invasion-resistant region. Because a noninverted mutant is treated as a new CI by the existing *Ak* population, a lower value of *Fk*1 will hinder its invasion.

Consistent with the findings regarding how changes in *Fk*2 values may affect the invasion of a mutant CI, similar logic and dynamics apply to how changes in *Fk*2 values may affect the invasion of a noninverted mutant into a population with a fixed CI. In this case, however, the fixed CI population is linked to the prevalent mating-bias allele, while the invading native ecotype, now treated like an invading mutant CI, is not physically linked to the mating-bias alleles.

Given the parametric values in Fig 20, with *f* = 0.3 and initial *Ax*/*Bx* population ratios in the range *Ax* ≥ 0.9, *Bx* ≤ 0.1 (or *By* ≥ 0. 9), if we let *Ak* = 0.9, signifying the presence of a noninverted mutant population ratio of 1 − *Ak* = 0.1, the noninverted mutant population can invade if *Fk*2 ≤ 0.16, which then drives *Ak* to extinction and destroys the premating RI. Even though the noninverted mutant population cannot invade for values of *Fk*2 > 0.16, when *Fk*2 is increased to 0.491 or above, *Ak* still fixes, but the *Ax*/*Bx* phase portrait now diverges and no premating RI is possible. Therefore, only values of *Fk*2 between 0.16 and 0.491 will drive *Ak* to fixation, create *Ax*/*Bx* mating-bias-allele polymorphism, and establish premating RI between the niches. In this way, the invasion dynamics of a noninverted mutant genotype mirror those of an inverted mutant CI invading a noninverted population—i.e., the noninverted native mutant cannot invade above a lower limit of Fk2 (which is 0.16 in the Fig 20 example).

Fig 21a shows that, after coupling, if disruptive ecological selection is completely removed by changing the value of *f* in Fig 20 from 0.3 to 0.5, with *Ax* = 0.9, *By* = 0.9, *Ak* = 0.95, *Fk*1 = 0.49 and *Fk*2 = 0.48, a small population of noninverted genotypes, with a population ratio 1 − *Ak* = 0.05, cannot invade and is eliminated. Therefore, the fixed CI and premating RI are resistant to invasion by noninverted mutants and cannot be reversed. Fig 21b shows the region in the *Ax*/*Bx* phase portrait that is resistant to invasion. However, if *Fk*1 is increased to 0.5, the noninverted mutant population can invade and eliminate the CI and its associated premating RI.

**Fig 21a.**
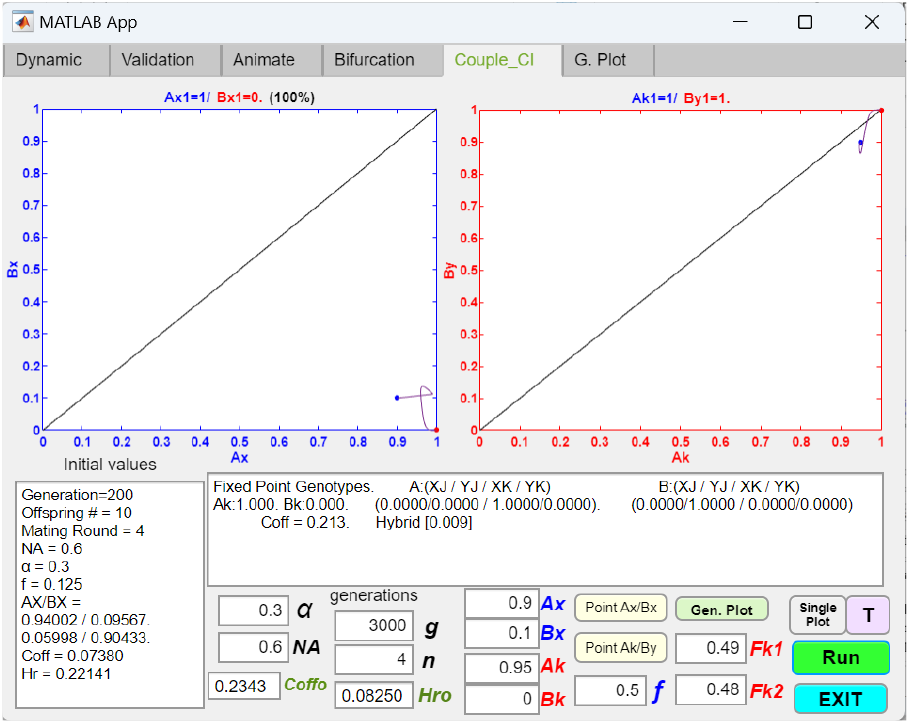
shows the fixation of a coupled chromosomal inversion that captures a partial set of locally adaptive ecological alleles in niche *A*, along with the mating-bias allele *X*, demonstrating resistance to invasion even when disruptive ecological selection is removed. An initial *Ax*/*Bx* mating-bias barrier has coupled with a CI with *Fk*1 = 0.49, *Fk*2 = 0.48, and their phase portraits have fixed points at *Ax*/*Bx* = 1/0, and *Ak* = 1. If disruptive ecological selection is removed by changing *f* from 0.3 to 0.5, mutant populations in the vicinities of the fixed points—i.e., at *Ax*/*Bx* = 0.9/0.1 and a noninverted population ratio of 1 − *Ak* = 0.05—cannot invade, as their vector trajectories converge back to the fixed points in the phase portraits. Therefore, the coupled barrier systems and RI are resistant to reversal.

**Fig 21b.**
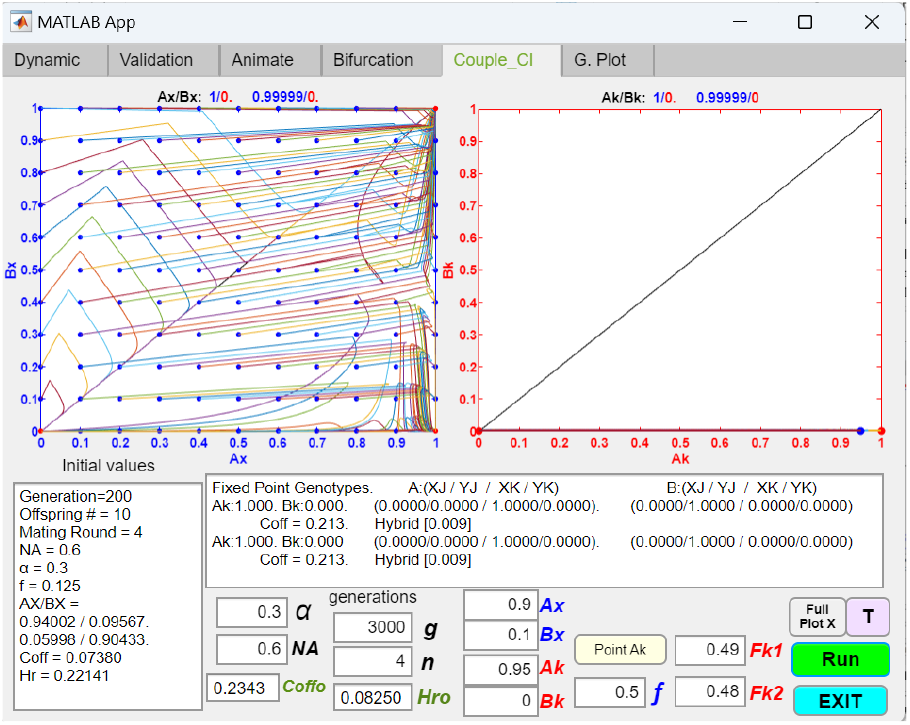
Fixation of a coupled chromosomal inversion that captures a partial set of the locally adaptive ecological alleles in niche *A*, along with the mating-bias allele *X*, showing the region in the *Ax*/*Bx* phase portrait that is resistant to invasion. Using the same parametric values as in Fig 21a, the *Ax*/*Bx* phase portrait (plotting against an invading noninverted population of 1 − *Ak* = 0.05) shows the region around the fixed point *Ax*/*Bx* = 1/0 that is resistant to invasion.

**Fig 21c.**
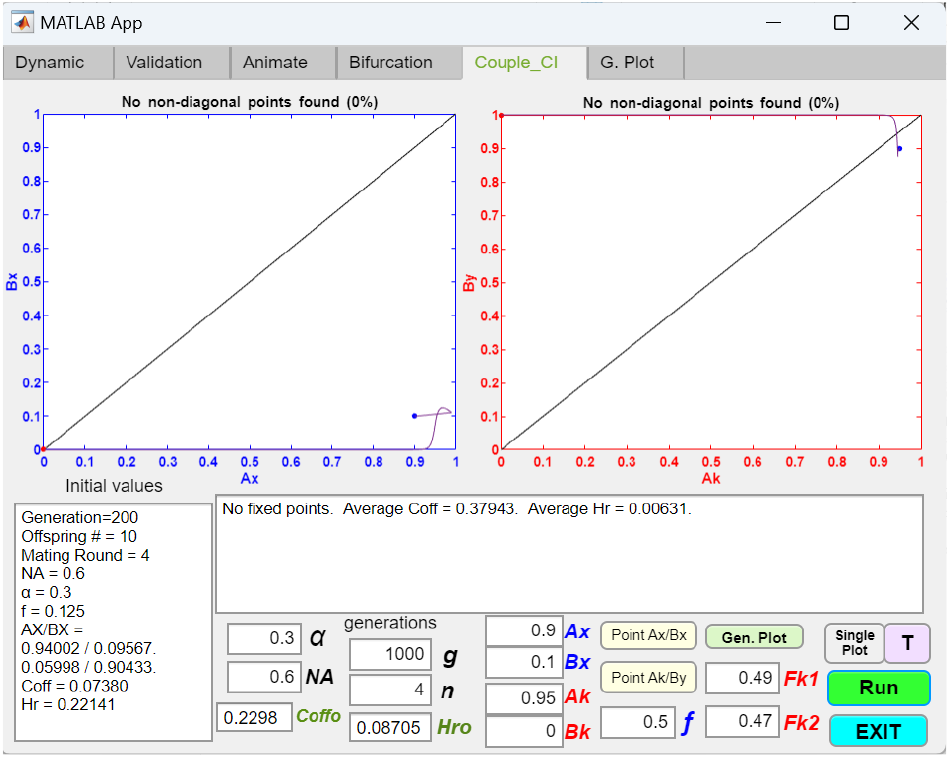
Fixation of a coupled chromosomal inversion that captures a partial set of the locally adaptive ecological alleles in niche *A*, along with the mating-bias allele *X*, showing invasion by noninverted genotypes when disruptive ecological selection is removed. In Figure 21a, if *Fk*2 of the CI is changed from 0.48 to 0.47, the noninverted population is able to invade and eliminate the CI genotype *Ak*. Consequently, the *Ax*/*Bx* phase portrait becomes divergent, and premating RI no longer exists.

**Fig 21d.**
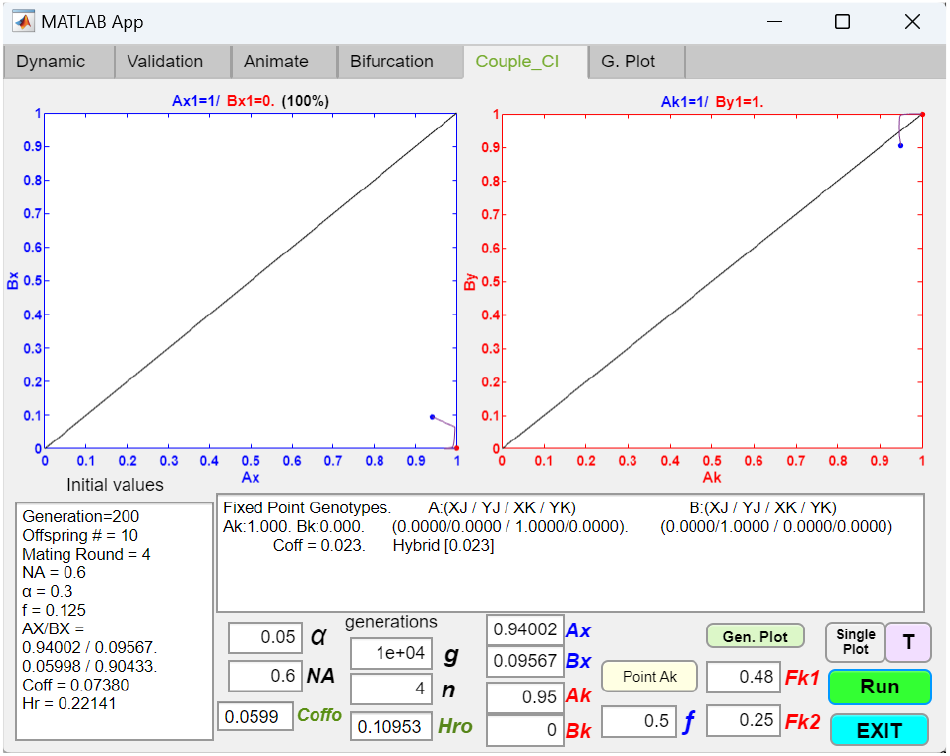
Fixation of a coupled chromosomal inversion that captures a partial set of the locally adaptive ecological alleles in niche *A*, along with the mating-bias allele *X*, showing increased invasion resistance when mating bias is increased (lower value of *α*). Decreasing the value of *α* in Figure 21c from 0.3 to 0.05 lowers the minimal value of *Fk*2 required to prevent the invasion of noninverted genotypes. Here, the established fixed points are resistant to invasion with an *Fk*2 value as low as 0.25.

**Fig 21e.**
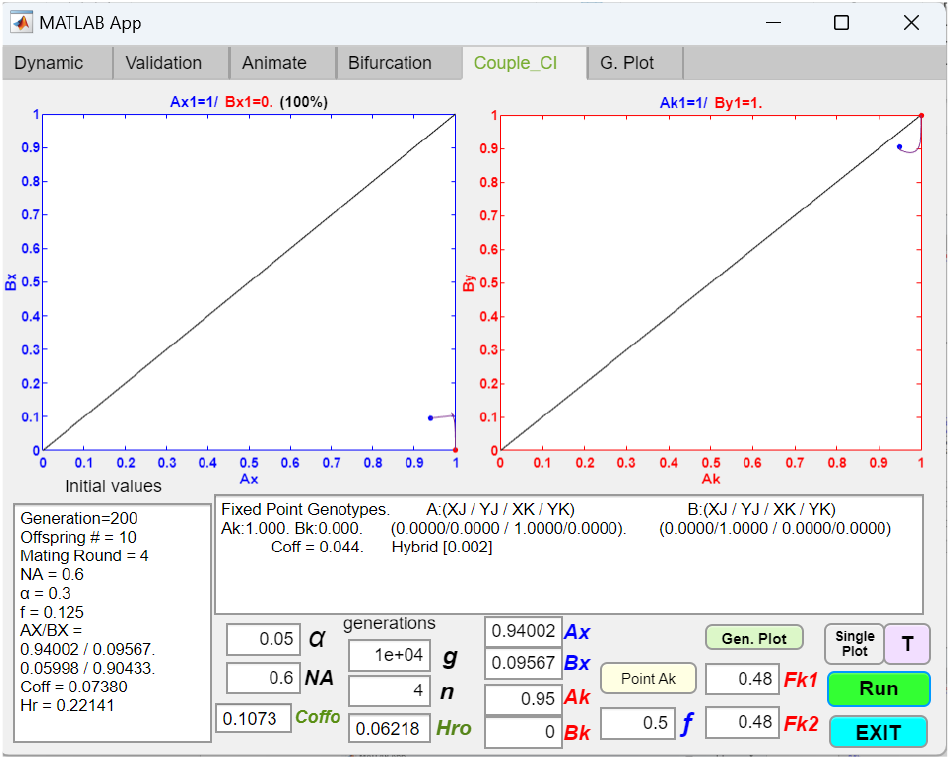
Fixation of a coupled chromosomal inversion that captures a partial set of the locally adaptive ecological alleles in niche *A*, along with the mating-bias allele *X*, showing the maximum value of *Fk*2 that can maintain premating RI after the invasion of a CI. If the value of *Fk*2 in Figure 21d is increased to 0.48, the *Ax*/*Bx* phase portrait remains convergent. However, if *Fk*2 is increased beyond 0.48, *Ak* remains fixed at 1, but the *Ax*/*Bx* phase portrait becomes divergent and premating RI is eliminated.

Fig 21c shows that if the value of *Fk*2 in Fig 20a is decreased from 0.48 to 0.47, both the CI and the premating RI are eliminated. However, in Fig 21d, if *α* is reduced to 0.05, then the fixed CI is resistant to invasion for *Fk*2 value as low as 0.25. When *α* = 0, a noninverted mutant cannot invade for any value of *Fk*2. We can call this *f* − *Fk*2 difference that protects the fixed CI from invasion the “invasion resistant range.” Fig 21e shows that the value of *FK*2 in Fig 21d can be increased to 0.48 and still maintain *Ax*/*Bx* premating RI. However, if *Fk*2 is increased above 0.48, the *Ax*/*Bx* phase portrait becomes divergent, and no premating RI exists. Therefore, based on the results shown in Figs 21d and 21e, when *f* is increased to 0.5 and *α* is reduced to 0.05, the range 0.25 < *Fk*2 < 0.48 defines where the fixed CI is resistant to invasion and can still maintain premating RI.

In general, decreasing the values of *α* and *Fk*1 increases the invasion resistant range for a fixed CI. A lower value of *Fk*1 makes it more difficult for a small population of noninverted mutants to invade because it suffers more offspring loss in intra-niche mating (Fig 14a). Similarly, a lower value of *α* reduces the offspring that the noninverted mutant population can receive from inter-niche mating. These findings support the expectation that, in late-stage sympatric speciation, when premating RI is already strong, a fixed CI linking favorable ecological and mating-bias alleles is more resistant to invasion.

#### 2.3. Coupling with Bateson–Dobzhansky– Muller (BDM) Incompatibilities

BDM incompatibilities are considered a late-stage mechanism of reproductive isolation (RI) that occurs only after a sufficient barrier to gene flow has developed between isolated populations. Without the homogenizing effects of gene flow, two isolated populations are free to accumulate ecological and mating incompatibilities as their genomes diverge through genetic drift, differential adaptations to different environments, or mutation-order adaptations to similar environments [25, 26]. In our study, we have developed GUI applications to study the coupling of BDM barriers after an initial two-allele mating-bias barrier has been established. Since differential adaptations can lead to very rapid divergence compared to other BDM mechanisms, it is the focus of our analyses, even though our GUI application is equally suited to analyzing mutation-order and genetic drift mechanisms as well.

Fig 22a shows the interface of a GUI application that models the coupling of a one-gene-locus BDM barrier with an initial two-allele mating-bias barrier. In this model, a *Z* allele represents evolved adaptations to the niche-*A* environment, mediated by the BDM mechanism. Consequently, individuals with the *Ax* genotype carry adaptive genetic changes that make them more fit than the native niche-*A* population at extracting niche-*A* resources. The value *Sax* specifies the fitness advantage that the *Ax* genotype has over its niche-*A* ecotype counterparts that lack BDM adaptations. For example, an *Sax* ratio of 1.1 indicates that at the end of each generation, the *Ax* offspring ratio receives an extra 10% boost because of the increased ecological fitness of the *Ax* genotype in niche *A*. Because the *Z* mutation is specifically evolved to adapt to the niche-*A* environment, its presence in the niche-*B* ecotype will likely suffer a fitness disadvantage. The value *Sbx* specifies the fitness disadvantage that a genotype *Bx* may suffer when compared to the native niche-*B* ecotype. For instance, an *Sbx* value of 0.9 indicates that the *Bx* offspring ratio will be reduced by an extra 10% in the niche-*B* environment.

**Fig 22a.**
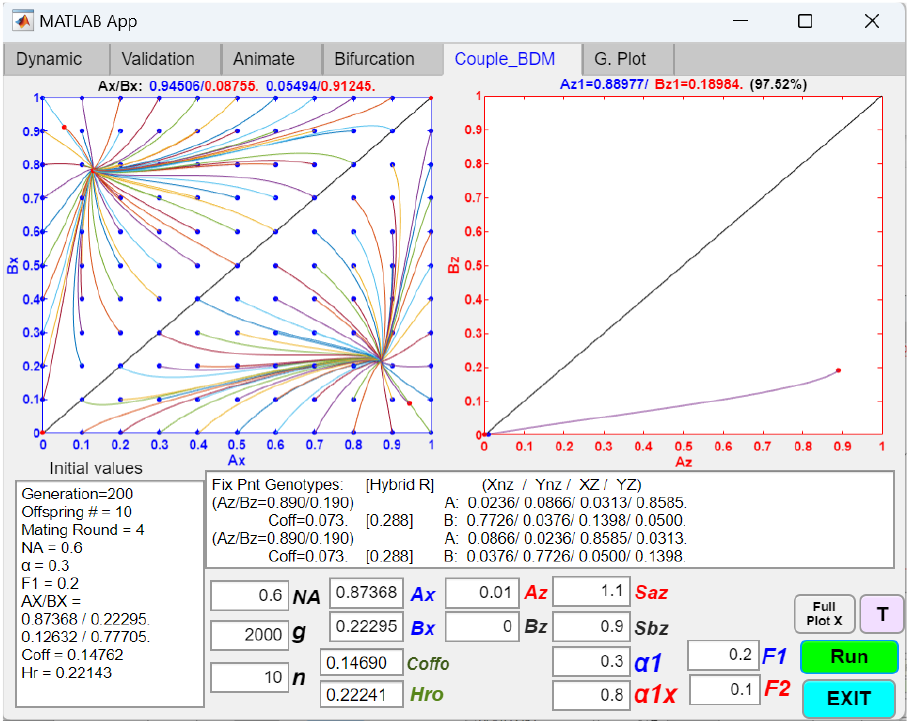
Coupling an initial two-allele premating barrier with a one-gene-locus BDM barrier mechanism, resulting in increased RI. The *Z* allele indicates the presence of evolved adaptations to the niche-*A* environment through the BDM mechanism, along with the associated mating-bias and hybrid incompatibilities. *Sax* represents the fitness advantage that an *Ax* genotype has over native niche-*A* ecotypes that lack the *Z* allele. An *Sax* value of 1.1 means that the *Ax* offspring produced in each generation is increased by 10% due to their increased adaptive fitness in the niche-*A* environment. Conversely, *Sbx* represents the fitness disadvantage that a niche-*B* ecotype has if it acquires the *Z* allele, which is maladaptive in the niche-*B* environment. An *Sbx* value of 0.9 means that the *Bx* offspring ratio is reduced by 10% after each mating generation. *F*1 is the same as *f*, the offspring return ratio for the original ecotypes. *F*2 is the new offspring return ratio due to the hybrid incompatibility created by the *Z* allele. Similarly, *α*1 = *α* is the original mating bias, and *α*1x is the mating bias due to the evolved BDM incompatibilities. When an *Ax* genotype encounters a native ecotype and they carry the same *X* or *Y* mating-bias allele, their matching success is determined by the mating-bias value 1 × *α*1x. When they carry different mating-bias alleles, their matching success is determined by the mating-bias value *α* × *α*1x. In this example, a small population of mutant genotype, *Ax*, has evolved better adaptive traits (indicated by the presence of the *Z* allele) in the niche-*A* environment, with a fitness advantage of *Sax* = 1.1 in niche *A*. However, when the same traits are transferred to the niche-*B* ecotype through the *Z* allele, the *Bx* genotypes suffer a fitness disadvantage of *Sbx* = 0.9 in the niche-*B* environment. *F*2 = 0.1 and *α*1x = 0.8 reflect the degrees of hybrid and mating-bias incompatibilities associated with the evolved BDM traits. The phase portraits show that a small mutant population, *Ax* = 0.01, is able to invade, reach a polymorphic fixed point at *Ak*/*Bk* = 0.88977/ 0.18984 after 2000 generations, and shift the initial *Ax*/*Bx* fixed point toward the lower right corner of the *Ax*/*Bx* phase portrait (from *Ax*/*Bx* = 0.87368/0.22295 to *Ax*/*Bx* = 0.94506/ 0.08755), while *Coff* decreases from 0.14690 to 0.073, resulting in greater overall RI. The matching and unit-offspring tables used in the GUI application are included in the Appendix (Tables 11a and 11b).

**Fig 22b.**
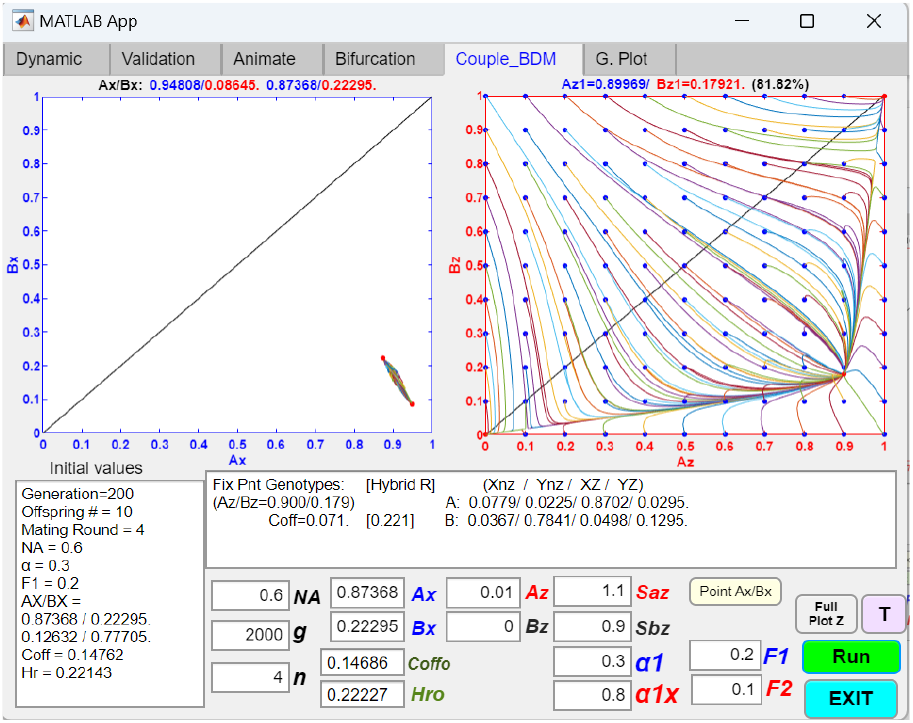
Coupling an initial two-allele premating barrier with a one-gene-locus BDM barrier mechanism, demonstrating an invasion resistance pattern in the *Ax*/*Bx* phase portrait as the value of *n* decreases (indicating increased assortative mating cost). Using the same parametric values as in Fig 22a, except that the number of matching rounds, *n*, is decreased from 10 to 4, the *Ax*/*Bx* phase portrait (plotted against the initial *Ax*/*Bx* fixed point: *Ax*/*Bx* = 0.87368/0.22295) shows an invasion resistance pattern, indicating that small mutant populations around the origin (where *Ak* = 0 and *Bk* = 0) cannot invade.

According to the BDM mechanism, hybrid incompatibility tends to arise as the niche-*A* and niche-*B* ecotypes diverge. This incompatibility is quantified by an offspring return ratio, *F*2, which measures the loss of hybrid offspring from matings between *Ax* individuals and niche-*B* individuals without the *Z* mutation. In Fig 22a, *F*1 = *f* represents the initial offspring return ratio for niche ecotypes lacking the *Z* mutation. As the niche populations diverge, their increased hybrid incompatibility will produce a *F*2 value that is lower than *F*1.

**Table 11a.**
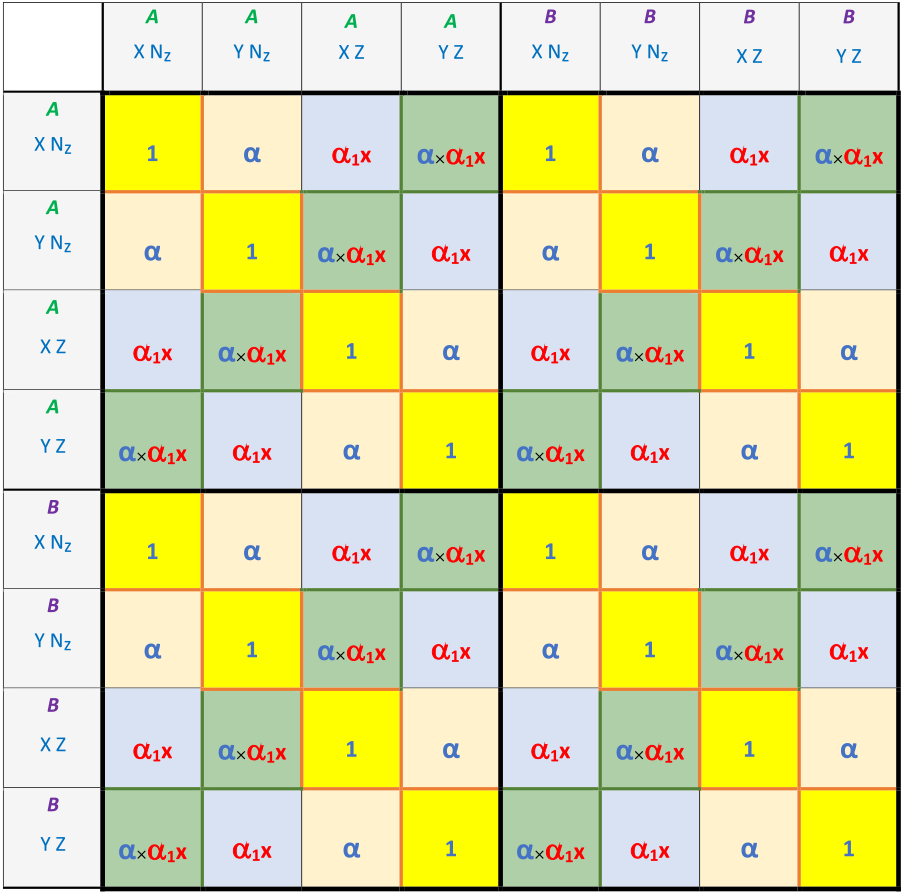
Matching table for coupling to a one-gene-locus BDM barrier mechanism. The presence of a *Z* allele in an individual‘s genotype indicates that adaptive changes to the niche-*A* environment due to the BDM mechanism have occurred. *Nx* indicates the absence of the *Z* allele in a genotype. *α*1x represents the evolved mating bias associated with the *Z* allele. When an individual carrying the *Z* allele encounters an individual lacking it, their matching success is determined by multiplying their pre-existing mating bias (1 or *α*) by. *α*1x.

**Table 11b.**
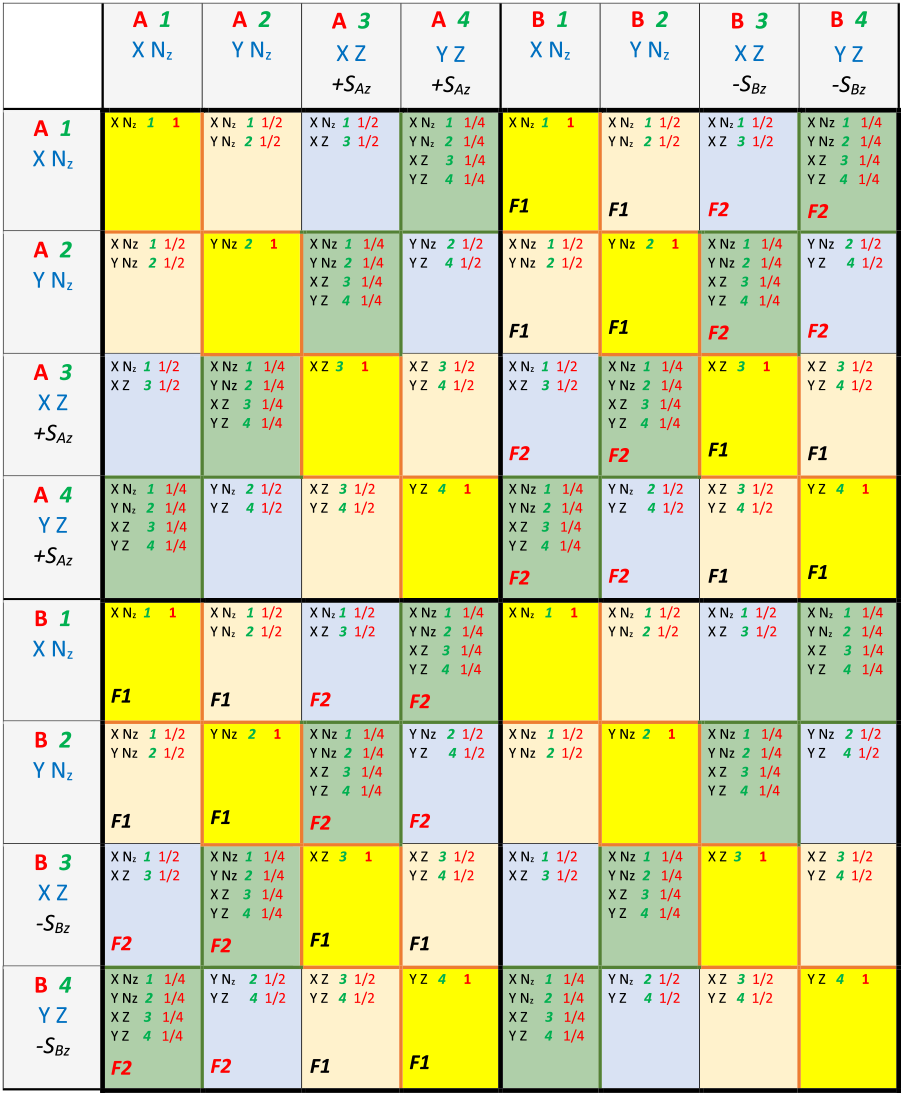
Unit offspring table for coupling to a one-gene-locus BDM barrier mechanism. In inter-niche mating, when one parent carries the *Z* allele and the other parent does not, their unit offspring ratios are determined by the offspring return ratio *F*2, which is conferred by the *Z* allele due to BDM hybrid incompatibility. *F*1 = *f* is the original offspring return ratio for the native ecotypes lacking the *Z* allele.

Mating-bias incompatibility also tends to accumulate as different ecotypes diverge and adapt to their distinct environments. In Fig 22a, this is modeled by introducing a new mating-bias value, *α*1x, which represents the evolved mating incompatibility between *Ax* individuals and the native niche-*A* and niche-*B* ecotypes. The value of *α*1x ranges from 0 to 1. *α*1 = *α* is the original mating-bias value for genotypes without the *Z* mutation. When an *Ax* genotype encounters a native niche-*A* or niche-*B* ecotype, their mating success is determined by multiplying the value of their mating bias by *α*1x. For instance, when an *Ax* genotype encounters a native ecotype carrying the same *X* or *Y* mating-bias allele, their matching success is determined by the mating-bias value 1 × *α*1x. When they carry different mating-bias alleles, their matching success is determined by the mating-bias value *α* × *α*1x. (See the corresponding mating compatibility table, Table 11a, in the Appendix.) An *α*1x value less than one indicates that mating incompatibility has evolved between the diverging ecotypes.

In Fig 22a, *Coff* represents the non-hybrid offspring ratio produced by inter-niche mating, while *Hr* represents the hybrid offspring ratio. Therefore, *Coff* + *Hr* is the total offspring ratio resulting from inter-niche mating. Because our model assumes no viable hybrids, *Coff* can be used as a measure of RI between the niche populations.

Fig 22a shows the emergence of a new, more adaptive mutant allele *Z* in niche *A*, with associated antagonistic pleiotropy values *Sax* = 1.1 and *Sbx* = 0.9, an evolved mating bias *α*1x = 0.8, and increased hybrid incompatibility signified by *F*2 = 0.1. With an initial population ratio of *Ax* = 0.01, such an allele can invade, stabilize at a fixed-point polymorphism (*Ax*/*Bx* = 0.88977/0.18984), and shift the *Ax*/*Bx* fixed point toward the lower right corner in the *Ax*/*Bx* phase portrait (from *Ax*/*Bx* = 0.87368/0.22295 to *Ax*/*Bx* = 0.94506/0.08755), while *Coff* decreases from 0.14690 to 0.073, resulting in stronger overall RI.

Fig 22b uses the same parametric values as in Fig 22a, except that the number of matching rounds, *n*, is decreased from 10 to 4. The *Ax*/*Bx* phase portrait, plotted against the initial *Ax*/*Bx* fixed point, shows an invasion resistant pattern. This occurs because the new mating bias, *α*1x, associated with the *Z* allele gives the *Ax* genotypes a fitness disadvantage in intra-niche mating with the native niche-*A* ecotypes when sexual selection is strong and when the starting population ratio, *Ax*, is small. This invasion resistance pattern can be mitigated by reducing intra-niche sexual selection against the *Z* allele, either by increasing the number of matching rounds, *n*, or by increasing the value of *α*1x. For instance, if *n* = 4, the *Ax*/*Bx* phase portrait becomes globally convert if *α*1x ≥ 0.92, allowing a mutant *Z* allele to invade. If *α*1x = 0.8, then *n* ≥ 7 is necessary for the *Z* allele to invade. Since the invasion-resistant pattern only restricts small invading mutant populations, once the *Z* allele achieves a sufficiently large population ratio within niche *A*, invasion resistance ceases to be a limiting factor. Thus, it may be more feasible for the *Z* allele to invade initially based on its ecological fitness advantage in niche *A*, enabling it to overcome invasion resistance from intra-niche sexual selection before developing enhanced mating-bias incompatibilities.

In general, as long as there is gene flow between the niches, the *Ax*/*Bx* phase portrait will stabilize at a fixed-point polymorphism (see Fig 23a). Increasing the premating barrier by decreasing *α*1x will increase *Ax* and decrease *Bx* at the fixed point and flatten the vector trajectory curve toward the *x*-axis (see Fig 23b). However, if the value of *α*1x in Fig 23b is decreased below 0.75, then a small mutant population with *Ax* = 0.01 cannot invade due to intra-niche sexual selection. In Fig 23a and 23b, decreasing the value of *α*1 also flattens the *Ax*/*Bx* vector trajectory toward the *x*-axis and moves the *Ax*/*Bx* fixed point closer to the bottom right corner of the phase portrait. In the limit, when there is no gene flow between the niches (i.e., when *α*1 = 0 or when *F*1 = *F*2 = 0), the ecological advantage of the *Z* allele in niche *A* (*Sax* = 1.1) propels the *Ax*/*Bx* vector trajectory to move along the *x*-axis and become fixed at *Ax* = 1 and *Bx* = 0.

**Fig 23a.**
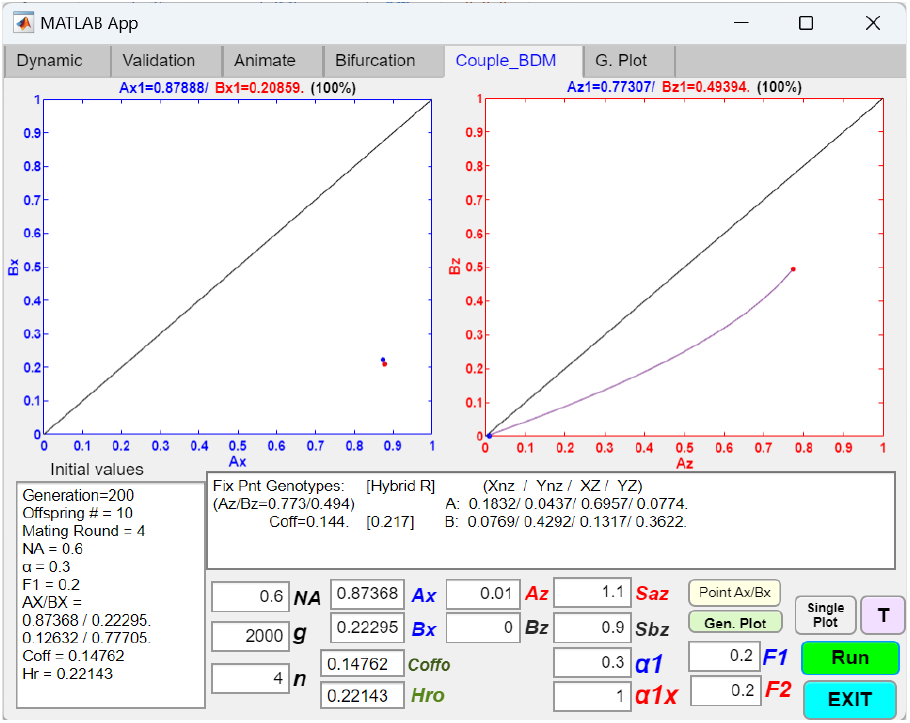
Coupling an initial two-allele premating barrier with a one-gene-locus BDM barrier mechanism, showing a niche-A adaptive mutant Z allele invading and reaching a fixed-point polymorphism. Single-line plots of *Ax*/*Bx* and *Ax*/*Bx* phase portraits show the population vector trajectories after 2000 generations, starting at an initial *Ax*/*Bx* fixed point (*Ax*/*Bx* = 0.87366/0.22295) and an initial mutant population ratio of *Ax* = 0.01. The results show that a mutant *Z* allele with ecological fitness advantage in the niche-*A* environment (*Sax* = 1.1) and an initial population ratio of *Ax* = 0.01 can invade and reach a fixed-point polymorphism in the *Ax*/*Bx* phase portrait. Here, *α*1x = 1 and *F*1 = *F*2 = 0.2 indicate that BDM mating-bias and hybrid incompatibilities have not evolved.

**Fig 23b.**
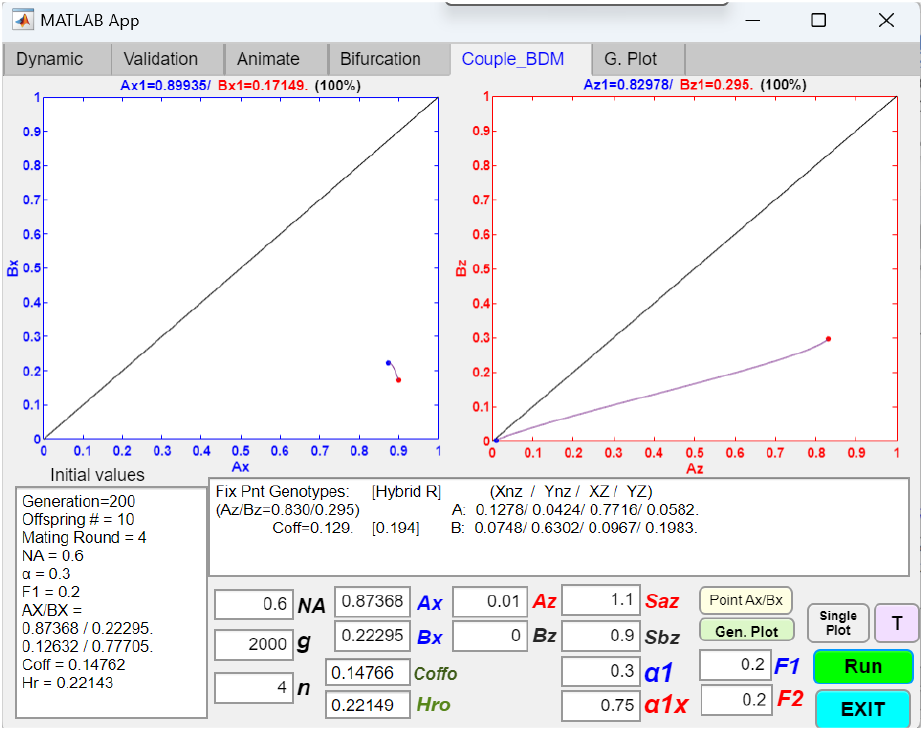
Coupling an initial two-allele premating barrier with a one-gene-locus BDM barrier mechanism, showing increased RI when the mating-bias value, A1x, is decreased. Reducing the value of *α*1x in Fig 23a from 1 to 0.75 lowers the final attained *Coff* and *Hr* values, signifying increased overall RI through coupling. However, for values of *α*1x below 0.75, the *Z* allele cannot invade due to intra-niche sexual selection.

**Fig 23c.**
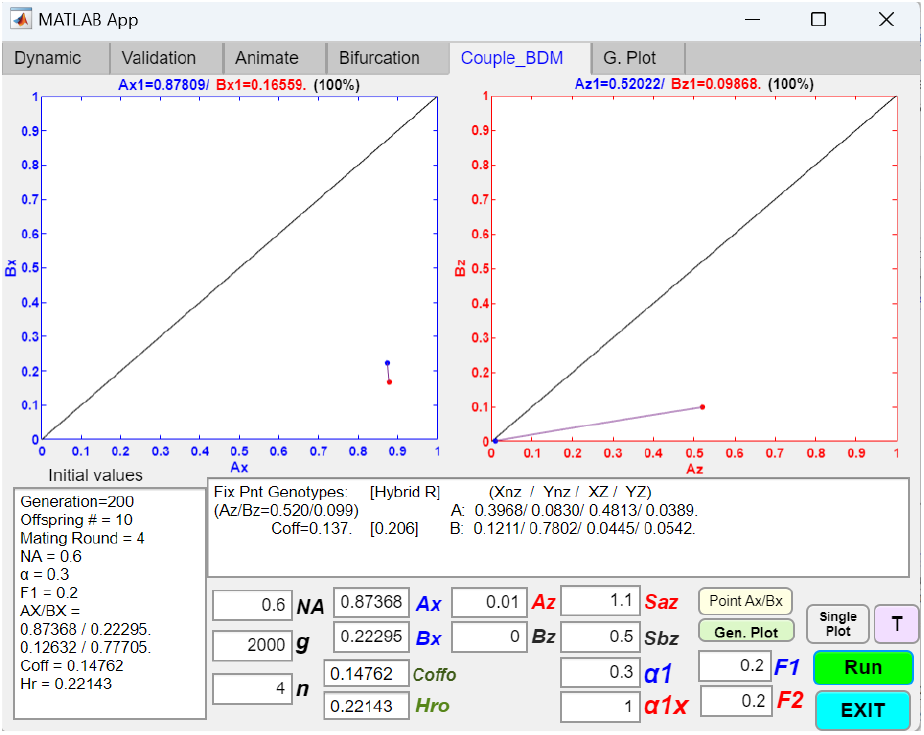
Coupling an initial two-allele premating barrier with a one-gene-locus BDM barrier mechanism, showing decreased *Ax* and *Bx* values at the fixed point when the value of *Sbx* is decreased. When the value of *Sbx* in Fig 23a is decreased from 0.9 to 0.5, indicating increased fitness disadvantage of the *Z* allele in the niche B environment, the ratios of the *Z* allele in both *A* and *B* niches decrease. Even though the *Z* allele has not evolved mating-bias and hybrid incompatibilities, widening the difference between *Sax* and *Sbx* can, by itself, increase the premating RI in the *Ax*/*Bx* phase portrait, resulting in stronger overall RI through coupling.

**Fig 23d.**
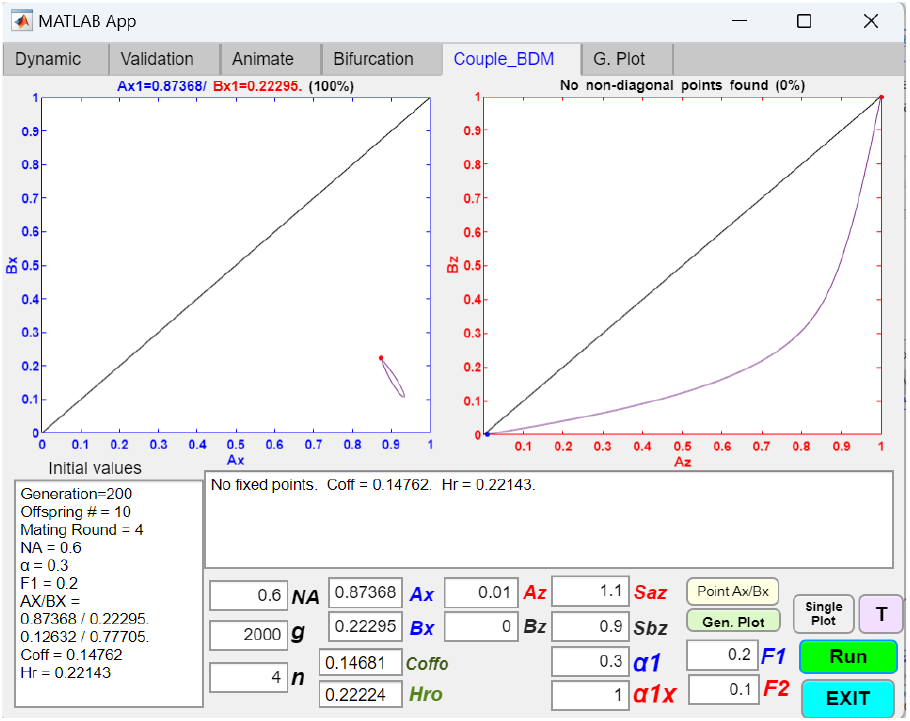
Coupling an initial two-allele premating barrier with a one-gene-locus BDM barrier mechanism, showing that decreasing the value of *F*2 may cause the *Z* allele to become fixed in both niches, leaving RI unchanged. When the value of *F*2 in Fig 23a is decreased from 0.2 to 0.1, the *Z* allele becomes fixed in both niche *A* and niche *B*, leaving the preexisting RI unchanged. However, if *F*2 ≤ 0.06, the Z allele can not invade.

**Fig 23e.**
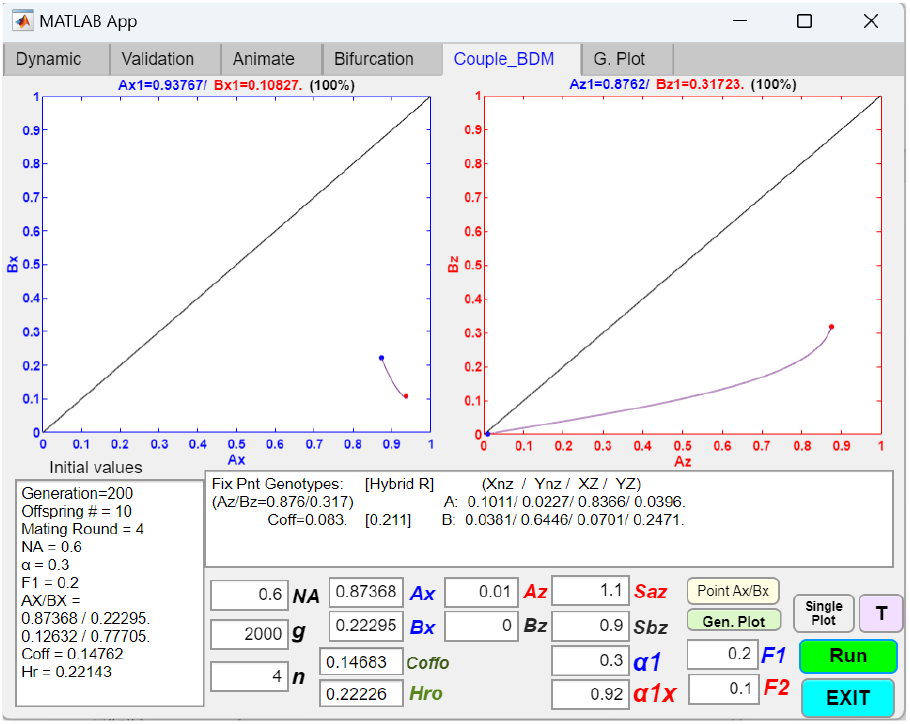
Coupling an initial two-allele premating barrier with a one-gene-locus BDM barrier mechanism, showing that adding mating-bias incompatibility to hybrid incompatibility can strengthen RI. When the value of *α*1x in Fig 23d is decreased from 1 to 0.92, indicating evolved mating-bias incompatibility, the invasion of the *Z* allele reaches a fixed-point polymorphism at *Ax*/*Bx* = 0.8762/0.31723 and shifts the *Ax*/*Bx* fixed point toward the lower right corner of the phase portrait, resulting in stronger overall RI. However, if the value of *α*1x is less than 0.92, the *Z* allele cannot invade.

**Fig 23f.**
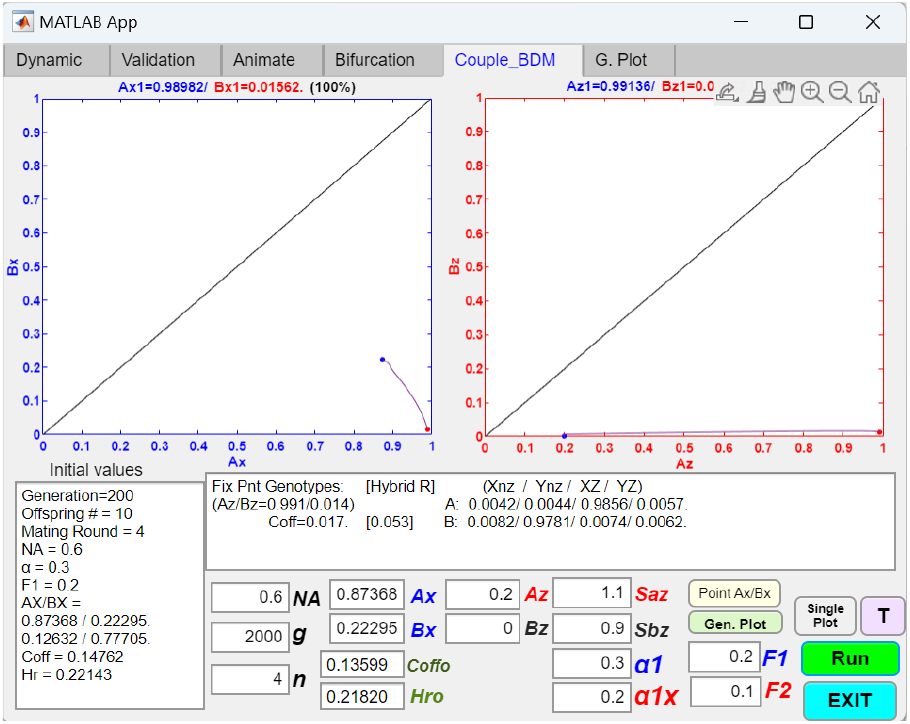
Coupling an initial two-allele premating barrier with a one-gene-locus BDM barrier mechanism, showing that a larger initial population ratio of *Az* can overcome invasion resistance due to intra-niche sexual selection. When the population ratio of *Az* in Fig 23e is increased from 0.01 to 0.2, the *Z* allele can invade even when the value of *α*1x is decreased to zero. In this example, the value of *α*1x is 0.2. Both the *Ax*/*Bx* and *Ax*/*Bx* fixed points are driven close to the lower right corners of the phase portraits, resulting in very strong overall RI.

**Fig 23g.**
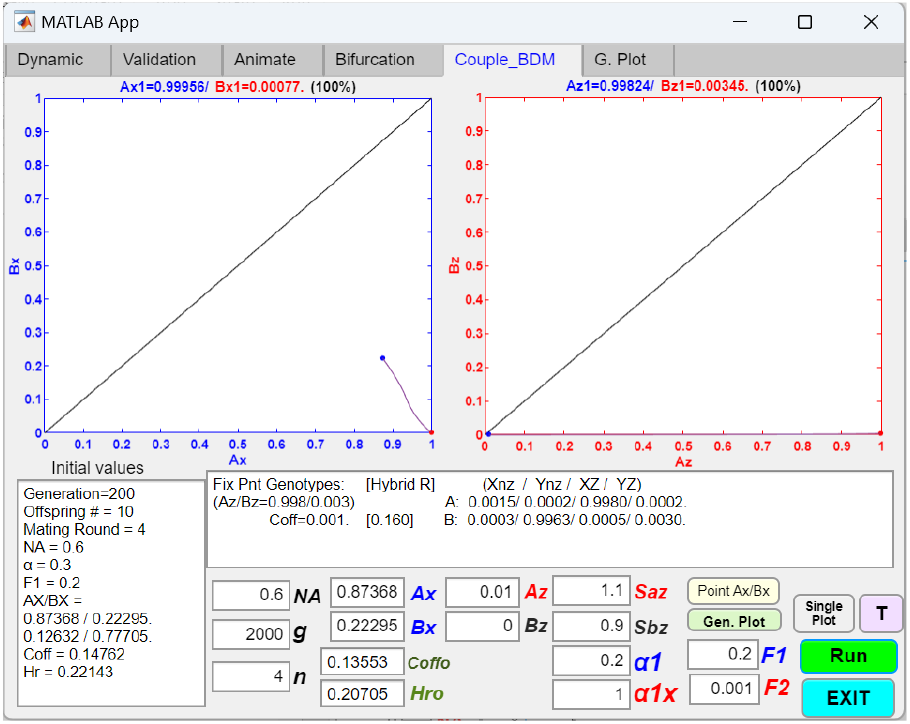
Coupling an initial two-allele premating barrier with a one-gene-locus BDM barrier mechanism, showing that increasing premating RI by decreasing the value of *α*1 facilitates the invasion of a *Z* allele with high hybrid incompatibility. When the value of *α*1 in Fig 23d is decreased from 0.3 to 0.2, the *Z* allele can invade and reach a fixed-point polymorphism. This is possible even when the value of *F*2 is lowered to zero. When *F*2 is zero, the *Ax*/*Bx* phase portrait has a fixed point at *Ax*/*Bx* = 1/0, the *Ax*/*Bx* fixed point shifts to *Ax*/*Bx* = 1/0, *Coff* = 0, and there is no gene flow between the niche populations.

Decreasing the value of *Sbx* decreases the viability of the *Z* allele in niche *B*. This tends to decrease both *Ax* and *Bx* at the fixed point and flatten the curve (Fig 23c). *Bx* is decreased because the *Bx* genotypes have decreased viability in niche *B. Ax* is also decreased because niche *B* now acts as a sink that draws and kills the *Z* alleles in niche *A* through inter-niche mating. Increasing the value of *Sax* produces the opposite effect; i.e., it increases both *Ax* and *Bx* at the fixed point. Nonetheless, increasing *Sax* or decreasing *Sbx* result in stronger premating RI. This is shown in the *Ax*/*Bx* phase portrait in Fig 23c. By widening the difference between *Sax* and *Sbx*, the *Ax*/*Bx* fixed point shifts toward the lower right corner. Thus, antagonistic divergence in adaptive ecological fitness can, by itself, create stronger RI independent of other system variables.

Fig 23d shows that if the value of *F*2 in Fig 23a is reduced from 0.2 to 0.1, the *Z* allele is able to invade and become fixed in both niche *A* and niche *B*, leaving the preexisting RI unchanged. This occurs because the increased fitness of the *Z* allele in niche *A* (*Sax* = 1.1) allows it to invade and increase its population ratio in niche *A*. This gives the *Bx* genotype in niche *B* a fitness advantage over the native niche-*B* ecotypes lacking the *Z* allele, as the offspring return ratio is *F*1 = 0.2 between *Ax* and *Bx* parents versus a ratio of *F*2 = 0.1 between *Ax* and the native niche-*B* parents. This fitness advantage from reduced hybrid loss is able to overcome the ecological disadvantage of the *Z* allele in niche *B* (*Sbx* = 0.9), allowing the *Z* allele to rise to fixation in niche *B*. Decreasing *NA* to less than 0.54 stops the invasion of the *Z* allele, while increasing *Sax* makes the fixation of the *Z* allele in both niches more likely.

When the *F*2 value in Fig 23d is decreased to 0.06 or less, the *Z* allele can no longer invade, as the fitness disadvantage of increased hybrid incompatibility outweighs the ecological advantage of the *Z* allele in niche *A*. Increasing the starting *Ax* population ratio or increasing the ecological advantage *Sax* can overcome this invasion resistance. With a population ratio of *Ax* = 0.2, the *Z* allele is able to invade even with an *F*2 value as low as zero. With this higher *Ax* population ratio, *Ax* = 0.2, as *F*2 tends closer to zero, fixed-point polymorphism, rather than fixation, begins to emerge in the *Ax*/*Bx* phase portrait, resulting in stronger RI.

In Fig 23e, reducing the value of *α*1x in Fig 23d from 1 to 0.92 couples mating-bias incompatibility with hybrid incompatibility in the invading *Z* allele and results in increased RI. Here, the fitness disadvantage of a lower *F*2 is offset by the fitness advantage of a reduced *α*1x, giving the *Z* allele a net fitness advantage to invade and establish RI that surpasses what could be achieved with the same *α*1x value alone. However, in the Fig 23e example, the *Z* allele cannot invade for an *α*1x value less than 0.92 due to intra-niche sexual selection. This invasion resistance can be overcome if the starting *Ax* population ratio is increased.

This is shown in Fig 23f. When the *Ax* population ratio in Fig 23e is changed from 0.01 to 0.2, the *Z* allele is able to invade with an *α*1x value of 0.2 and produce strong RI. In fact, the value of *α*1x in Fig 23f can be as low as zero and still allow the *Z* allele to invade. When *α*1x = 0, the *Z* allele becomes fixed at *Ax* = 1 and *Bx* = 0. These results imply that once the *Z* allele is able to invade and establish a sizable population in niche *A*, invasion resistance due to intra-niche sexual selection is no longer a consideration, and the *Z* allele is free to evolve ever stronger mating-bias incompatibility and strengthen the overall RI.

In summary, Figs 23b and 23d–23f illustrate that in BDM divergence, as hybrid incompatibility increases and *F*2 drops below *F*1, the *Z* allele initially becomes fixed in both niches without altering RI. At this stage, it can only increase RI by coupling with relevant mating-bias incompatibilities (Fig 23e). Further decreases in *F*2 lead to fixed-point polymorphisms in the *Ax*/*Bx* phase portrait, increasing RI. However, below a critical *F*2 threshold, hybrid loss becomes too severe, and the *Z* allele can no longer invade. Similarly, as mating-bias incompatibility evolves and *α*1x decreases below one, RI strengthens until a lower *α*1x limit is reached, beyond which intra-niche sexual selection prevents *Z* allele invasion. Near these critical *F*2 and *α*1x limits, an invasion-resistant pattern emerges in the *Ax*/*Bx* phase portrait, which can only be overcome by a higher initial *Ax* population ratio.

Fig 23g shows that if we increase the premating RI in Fig 23d by decreasing the value of *α*1 from 0.3 to 0.2, an *Ax*/*Bx* fixed-point polymorphism is produced. This shifts the original *Ax*/*Bx* fixed point toward the lower right corner of the phase portrait, creating a stronger overall RI. If, in Fig 23d, the value of *F*2 is kept at 0.2, then for values of *α*1 below 0.29, i.e., 0 ≤ *α*1 ≤ 0.29, both *Ax*/*Bx* and *Ax*/*Bx* polymorphisms appear in the phase portraits, and RI is strengthened. The lower the value of *α*1, the stronger the resultant RI. For values of *α*1 between 0.29 and 0.33, 0.29 < *α*1 ≤ 0.33, the *Z* allele becomes fixed in both niches, and the existing *Ax*/*Bx* fixed point remains unchanged. For *α*1 > 0.33, the *Z* allele fails to invade. Similarly, if the value of *α*1 in Fig 23d is kept constant at 0.3, for 0 ≤ *F*1 < 0.19, *Ax*/*Bx* and *Ax*/*Bx* polymorphisms are present, and RI is strengthened. The lower the value of *F*1, the stronger the resultant RI. If 0.19 < *F*1 ≤ 0.21, the *Z* allele becomes fixed in both niches, and if *F*1 > 0.21, the *Z* allele fails to invade. Therefore, lower values of *α*1, lower values of *F*1, and a greater difference between *Sax* and *Sbx* can facilitate the invasion of the *Z* allele and lead to the emergence of fixed-point polymorphisms and stronger RI. These findings seem to confirm the premise that the BDM mechanism of incompatibility works best when there is already reduced gene flow between niches.

Fig 24a shows the computer interface of a GUI application that models the coupling of an initial mating-bias barrier with a two-gene-locus, two-allele model of the BDM barrier. In the model, two mutant alleles, *Z* and *W*, reside at two separate gene loci. The presence of the *Z* allele in an individual‘s genotype indicates that local adaptations to the niche-*A* environment, through the BDM mechanism, have occurred; similarly, the presence of the *W* allele indicates adaptations to the niche-*B* environment. When a locally adaptive mutant genotype *Ax* emerges in niche *A* and a locally adaptive mutant genotype *Bw* emerges in niche *B, Sax, Sbx, Saw*, and *Sbw* represent the antagonistic pleiotropic advantages and disadvantages of the *Z* and *W* alleles in the different niches. *F*1 = *f* is the original offspring return ratio for the native niche ecotypes that do not carry the *Z* or *W* alleles. *F*2 is the new inter-niche hybrid incompatibility that arises when parents from different niches, carrying different *Z* or *W* alleles, mate to produce offspring (see the unit-offspring table, Table 12b, in the Appendix for details). In this GUI application, mating-bias incompatibilities due to the presence of the *Z* or *W* alleles are not modeled.

**Table 12a.** Matching table for coupling to a two-gene-locus, two-allele model of BDM barrier mechanism. Two alleles, *Z* and *W*, reside at two separate gene loci. The presence of a *Z* allele in an individual’s genotype indicates that adaptive changes to the niche-*A* environment due to the BDM mechanism have occurred; similarly, the presence of a *W* allele indicates that adaptive changes to the niche-*B* environment have occurred. *Nx* indicates the absence of the *Z* allele, and *Nw* indicates the absence of the *W* allele. The *X* and *Y* alleles determine the mating-bias (1 or *α*) between genotype encounters. Hybrid incompatibilities due to the presence of the *Z* or *W* alleles are not modeled.

|  | 1 | 2 | 3 | 4 | 5 | 6 | 7 | 8 |
| --- | --- | --- | --- | --- | --- | --- | --- | --- |
|  | X N<sub>Z</sub> N<sub>w</sub> | Y N<sub>Z</sub> N<sub>w</sub> | X Z N<sub>w</sub> | Y Z N<sub>w</sub> | X N<sub>Z</sub> W | Y N<sub>Z</sub> W | X Z W | Y Z W |
| 1 | 1 | $\alpha$ | 1 | $\alpha$ | 1 | $\alpha$ | 1 | $\alpha$ |
| 2 | $\alpha$ | 1 | $\alpha$ | 1 | $\alpha$ | 1 | $\alpha$ | 1 |
| 3 | 1 | $\alpha$ | 1 | $\alpha$ | 1 | $\alpha$ | 1 | $\alpha$ |
| 4 | $\alpha$ | 1 | $\alpha$ | 1 | $\alpha$ | 1 | $\alpha$ | 1 |
| 5 | 1 | $\alpha$ | 1 | $\alpha$ | 1 | $\alpha$ | 1 | $\alpha$ |
| 6 | $\alpha$ | 1 | $\alpha$ | 1 | $\alpha$ | 1 | $\alpha$ | 1 |
| 7 | 1 | $\alpha$ | 1 | $\alpha$ | 1 | $\alpha$ | 1 | $\alpha$ |
| 8 | $\alpha$ | 1 | $\alpha$ | 1 | $\alpha$ | 1 | $\alpha$ | 1 |

**Table 12b.**
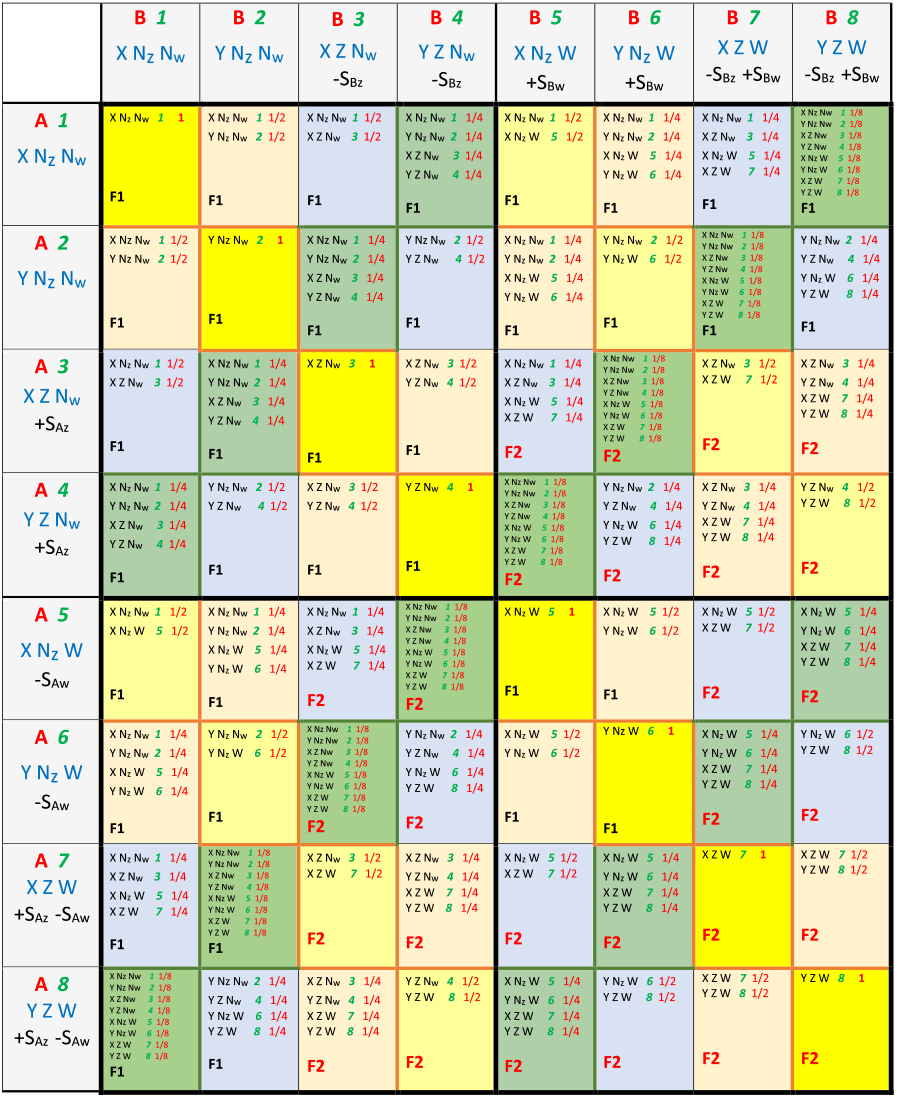
Unit offspring table for coupling to a two-gene-locus, two-allele model of BDM barrier mechanism. In inter-niche mating, the unit offspring return ratio *F*1 changes to *F*2 whenever parents from different niches possess different Z or W alleles.

**Fig 24a.**
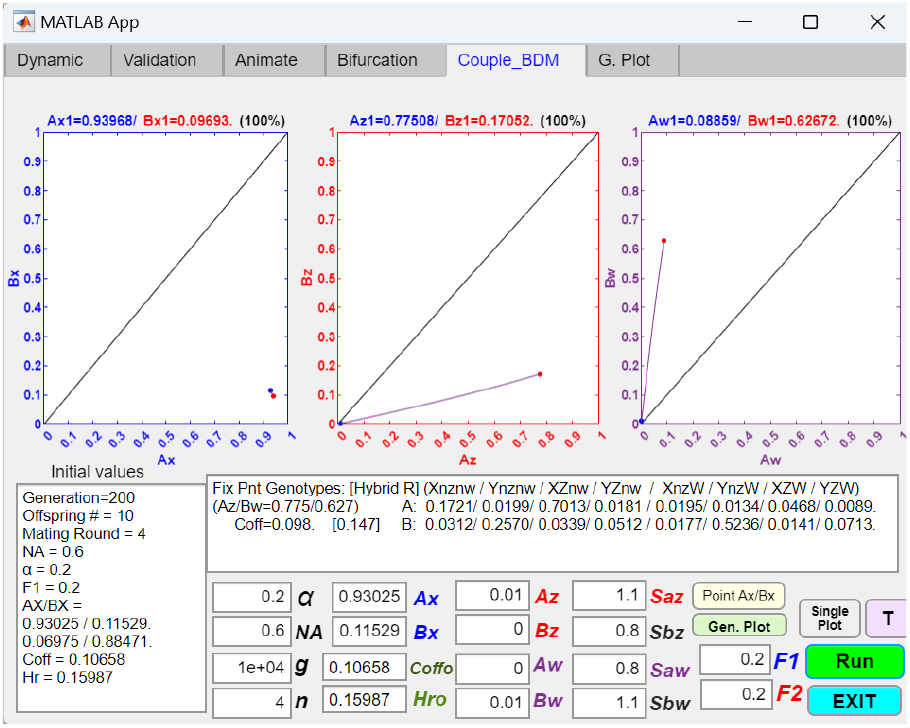
Coupling an initial two-allele premating barrier with a two-gene-locus, two-allele model of the BDM barrier mechanism results in increased RI. Two alleles, *Z* and *W*, reside at two gene loci. The presence of the *Z* allele in an individual‘s genotype indicates that local adaptations to the niche-*A* environment, through the BDM mechanism, have occurred; similarly, the presence of the *W* allele indicates adaptations to the niche-*B* environment. *Sax* represents the fitness advantage of an *Ax* genotype over the native niche-*A* ecotypes, and *Sbx* represents the fitness disadvantage experienced by a niche-*B* ecotype if it acquires the *Z* allele. *Saw* and *Sbw* are similarly defined for the *Bw* genotype. *F*1 is the same as *f*, the offspring return ratio for the original ecotypes. *F*2 is the new offspring return ratio due to hybrid incompatibilities in inter-niche mating when parents from different niches carry different *Z* or *W* alleles. Mating-bias incompatibilities due to divergence of the *Z* and *W* alleles (i.e., *α*1x in the Fig 22a example) are not modeled. In the figure, the phase portrait solutions show that antagonistic ecological divergence, represented by the differences *Sax* − *Sbx* and *Saw* − *Sbw*, can independently drive the invasion of locally adaptive mutant *Z* and *W* alleles, shift the preexisting *Ax*/*Bx* fixed point toward the lower right corner of the *Ax*/*Bx* phase portrait, and increase overall RI. The matching and unit-offspring tables used in the GUI application are included in the Appendix (Tables 12a and 12b).

**Fig 24b.**
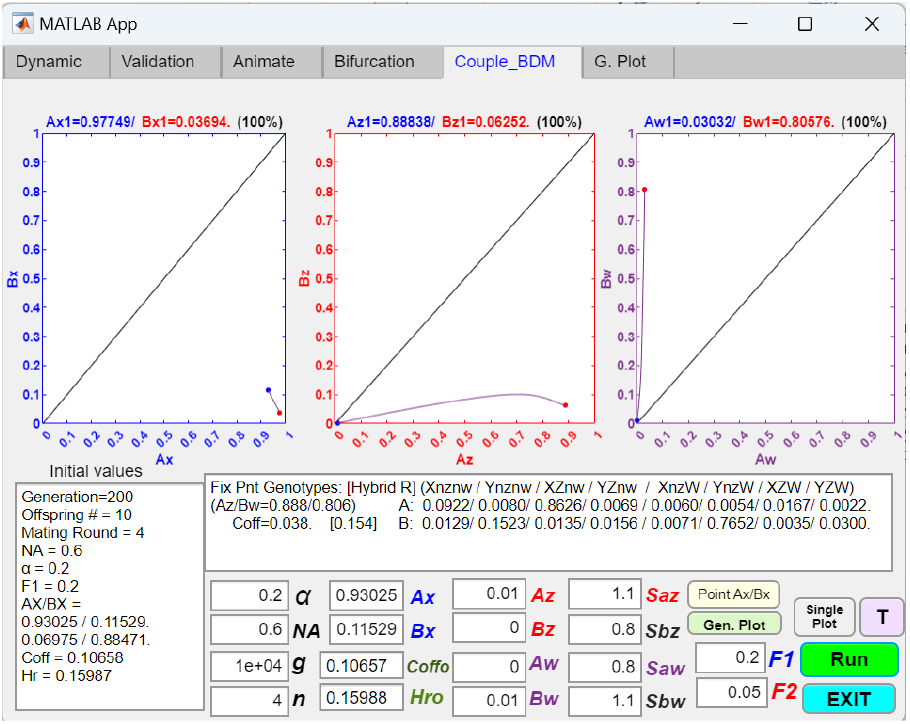
Coupling an initial two-allele premating barrier with a two-gene-locus, two-allele model of the BDM barrier mechanism, showing that increasing hybrid incompatibility between the ecologically divergent *Z* and *W* alleles increases overall RI. Reducing the value of *F*2 in Fig 24a from 0.2 to 0.05 moves the *Ax*/*Bx* fixed point closer to the lower right corner of the phase portrait, resulting in greater RI.

**Fig 24c.**
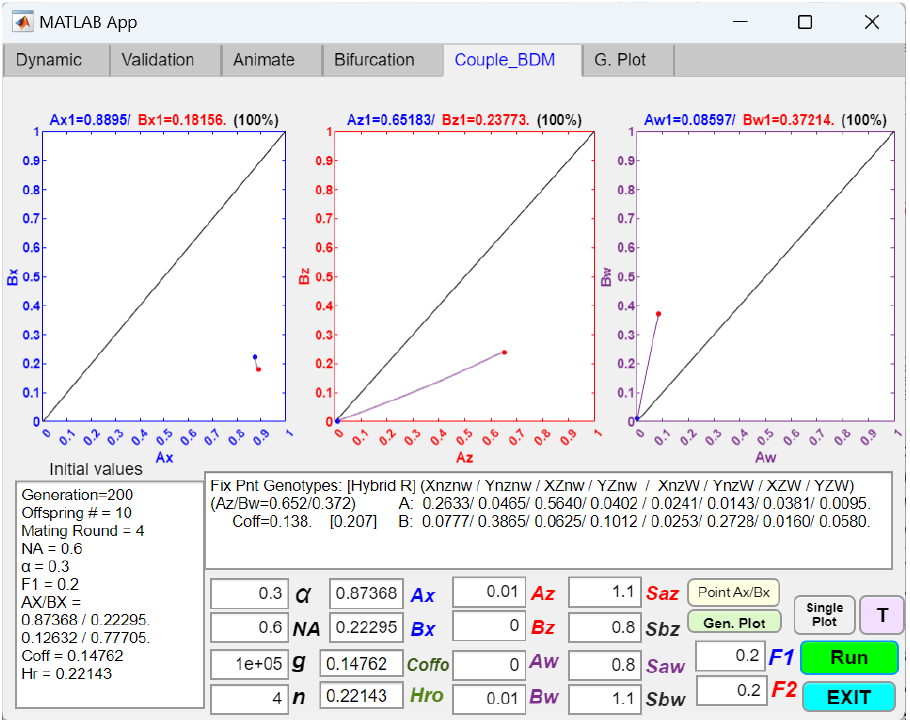
Coupling an initial two-allele premating barrier with a two-gene-locus, two-allele model of the BDM barrier mechanism, showing that decreased pre-invasion RI reduces the differential assortment of *Z* and *W* alleles between the niches. The initial *α* value in Fig 24a is increased from 0.2 to 0.3, creating increased gene flow before the invasion of the *Z* and *W* alleles. This causes the resultant vector trajectories in the *Ax*/*Bx* and *Aw*/*Bw* phase portraits to shorten and move closer to the diagonal lines. Differential assortment of the *Z* and *W* alleles between the niches is reduced at the fixed points.

**Fig 24d.**
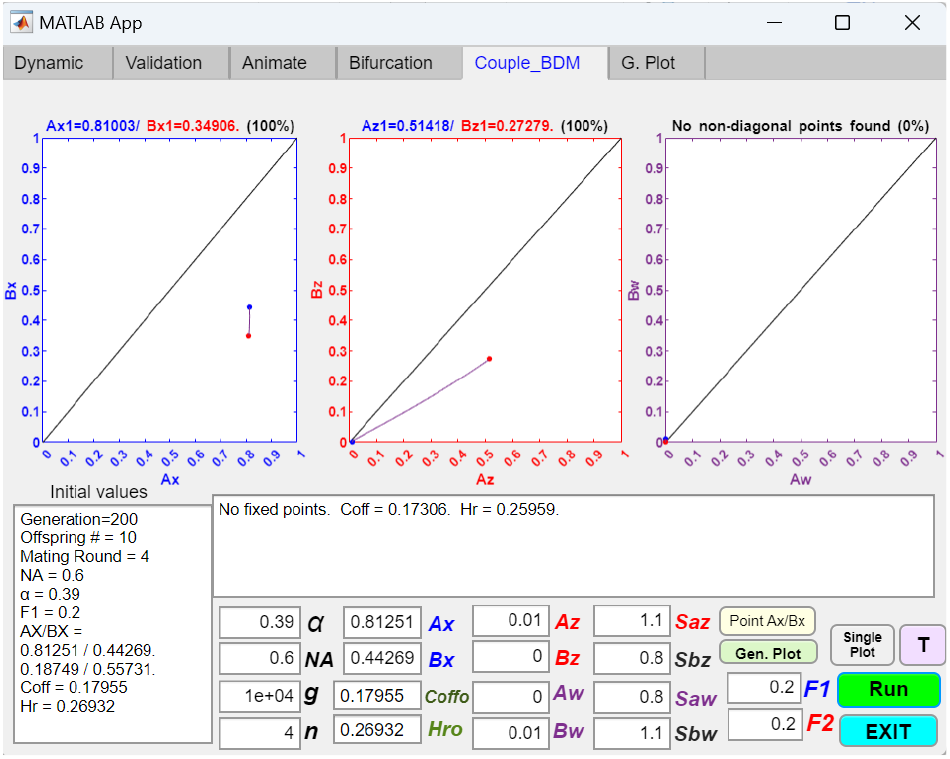
Coupling an initial two-allele premating barrier with a two-gene-locus, two-allele model of the BDM barrier mechanism, showing that when gene flow is large, the *W* allele fails to invade. Here, the value of *α* in Fig 24c is further increased from 0.3 to 0.39. This stops the invasion of the *W* allele. Consequently, the value of *F*2, representing hybrid incompatibility due to ecological divergence, has no effect on system dynamics.

Fig 24a shows that when premating RI is strong enough, *Ax* and *Bw* can invade and establish fixed-point polymorphism in their respective phase portraits. Similar to the example shown in Fig 23c, the differences between *Sax* and *Sbx* and between *Saw* and *Sbw* alone can increase premating RI by moving the pre-invasion *Ax*/*Bx* fixed point toward the lower right corner of the *Ax*/*Bx* phase portrait. Fig 24b shows that when there is sufficient differential assortment of the *Z* and *W* alleles between the two niches, reducing inter-niche hybrid viability (e.g., by decreasing *F*2 in Fig 24a from 0.2 to 0.05) can further increase the overall RI. In the Fig 24c phase portraits, reducing the pre-invasion RI by increasing the value of *α* tends to shorten and flatten the *Ax*/*Bx* and *Aw*/*Bw* trajectories toward the diagonal line, reducing the differential assortment of the *Z* and *W* alleles between the niches. In Fig 24d, further increasing the value of *α* eventually stops the invasion of the *W* allele, and the value of *F*2 no longer has any effect. Overall, the results of the two-niche adaptation model, shown in Fig 24a to 24d, are similar to those of the one-niche model, shown in Fig 23a to 23g. The dynamics of both models confirm that the BDM mechanism works best in an environment with already low gene flow and that it can be coupled with an initial premating barrier to strengthen overall RI.

In Fig 25a, an example is presented illustrating how a mating-bias barrier system that induces initial premating RI can couple with later-stage BDM barrier mechanisms to reinforce RI. During advanced stages of sympatric speciation, niche ecotypes, released from the homogenizing effect of gene flow, can diverge through the BDM mechanism of incompatibilities as they adapt to their respective niche environments. Over time, this divergence strengthens, as reflected by the parametric values depicted in Fig 25a: strong mating bias (*α* = 0.1), increasing ecological differentiation (*Sax* = 1.3, *Sbx* = 0.8, *Saw* = 0.8, and *Sbw* = 1.3), and high levels of hybrid incompatibility (*F*2 = 0.1), ultimately leading to robust and nearly complete RI.

**Fig 25a.**
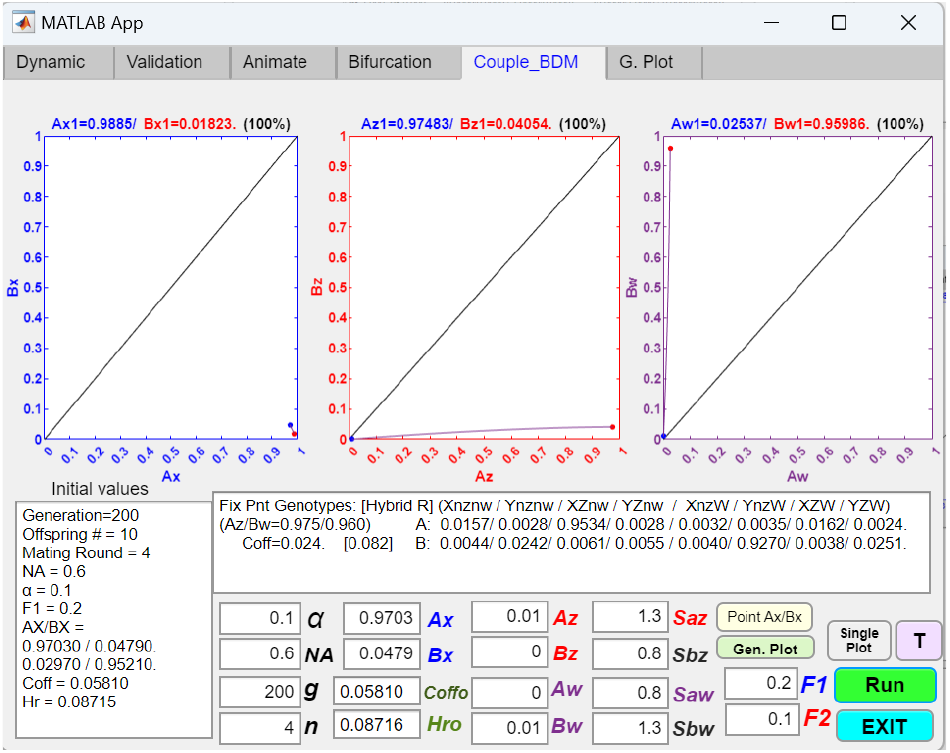
Coupling an initial two-allele premating barrier with a two-gene-locus, two-allele model of the BDM barrier mechanism, showing that, in advanced stages of sympatric speciation, BDM divergence and incompatibilities can generate strong RI. The parametric values reflect advanced stages of BDM divergence and incompatibilities, resulting in robust and nearly complete RI

**Fig 25b.**
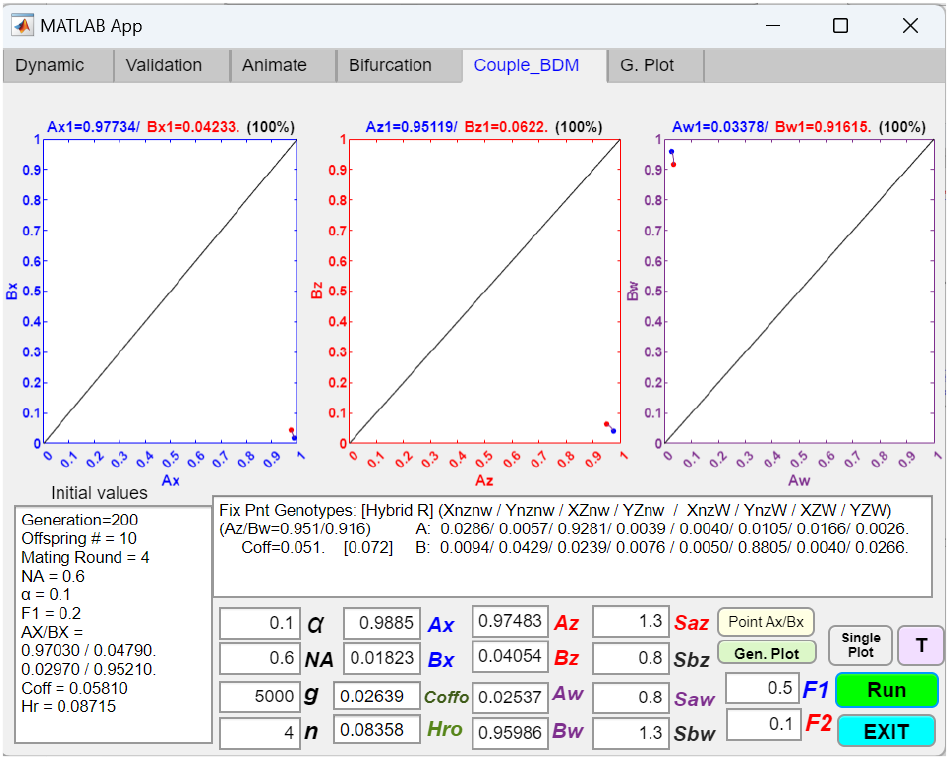
Coupling an initial two-allele premating barrier with a two-gene-locus, two-allele model of the BDM barrier mechanism, showing that the established RI is resistant to reversal even when external disruptive ecological selection is removed. Given the established *Ax*/*Bx, Ax*/*Bx, Aw*/*Bw* fixed-point values in Fig 25a, removing external disruptive ecological selection by changing the value of *F*1 from 0.2 to 0.5 does little to alter the positions of the fixed points or reduce the existing RI. This occurs because, when there is a high degree of differential assortment of the *Z* and *W* alleles across the niches, intrinsic hybrid incompatibility—represented by *F*2 = 0.1—can take over the role of *F*1 and ensure continued hybrid loss.

**Fig 26.**
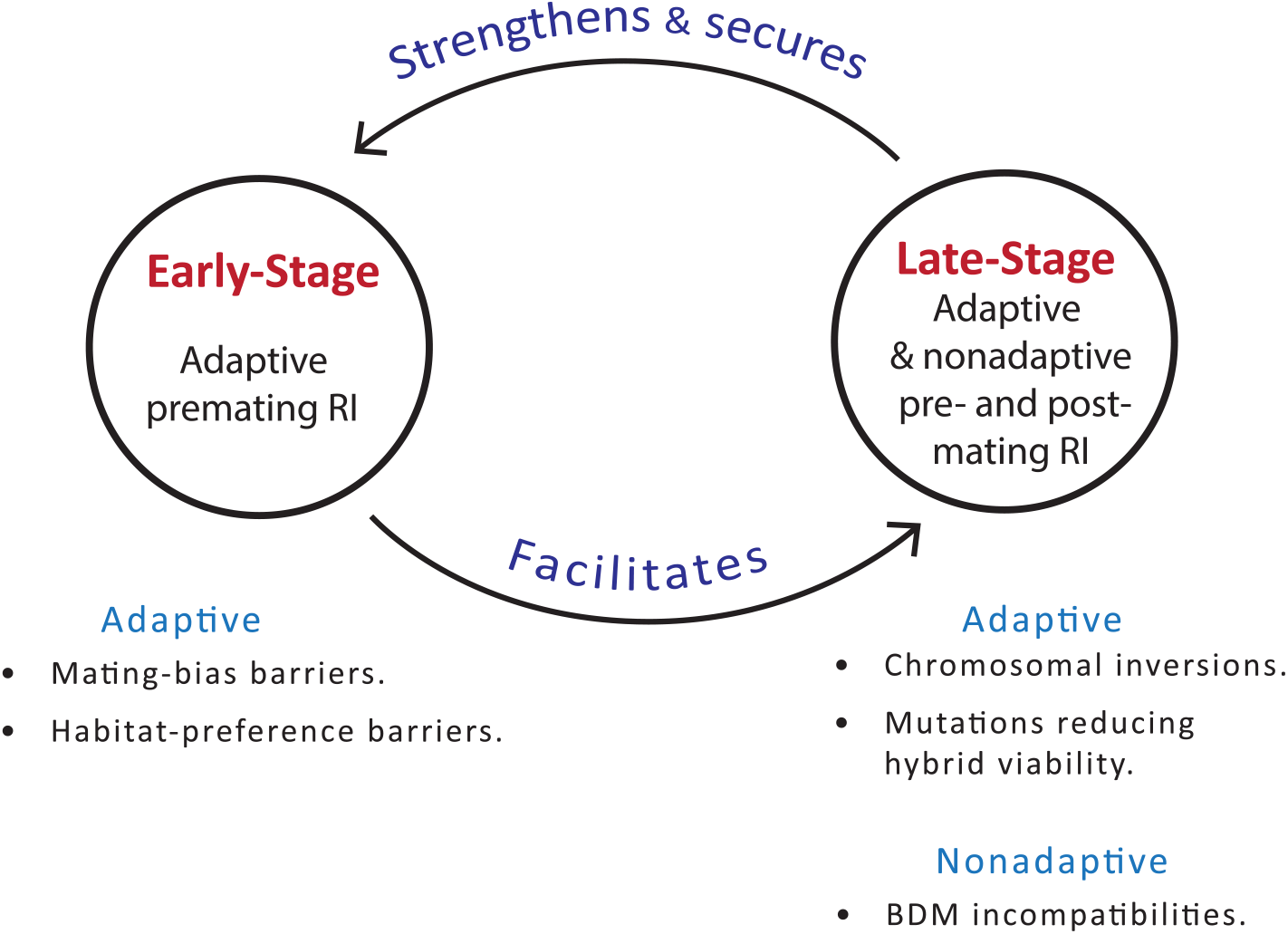
Early-stage and late-stage mechanisms interact synergistically in a positive feedback loop to strengthen and reinforce each other to complete the speciation process. Mating bias and habitat preference create premating barriers, while chromosomal inversions and BDM incompatibilities can generate both premating and post-mating barriers. Mutations that reduce hybrid viability can only produce post-mating barriers.

Additionally, Fig 25b demonstrates that once BDM barriers are firmly established, even the removal of disruptive ecological pressure from Fig 25a—by increasing *F*1 from 0.2 to 0.5—has minimal impact on the entrenched BDM divergence and RI. This underscores the largely irreversible nature of BDM incompatibility mechanisms.

## IV. DISCUSSION

The theory of evolution forms the bedrock of modern biology, and understanding the process of speciation lies at the heart of evolutionary theory. Since Ernst Mayr introduced the biological species concept in 1942 [46], it has become clear that speciation is not a discrete event but a continuum [9]. At one end of this continuum, incipient species exhibit weak reproductive isolation (RI), characterized by substantial gene flow, frequent introgression, and hybridization between sister species. At the opposite end, species display nearly complete RI, with little to no gene flow, introgression, or hybridization. Between these extremes, species undergo a progressive reduction in gene flow as they move toward completion of speciation. Notably, species nearing the end of the speciation continuum often possess multiple reproductive barriers distributed across their genomes, which interact synergistically to reinforce strong overall RI [15-19].

In current evolutionary thinking, allopatric speciation is widely regarded as the default mechanism for explaining the emergence of species in nature [25]. Even in cases where allopatric speciation struggles to account for speciation events that seemingly occurred without geographical barriers, the model of transient allopatric divergence followed by secondary contact is often invoked to resolve these anomalies. In contrast, sympatric speciation is frequently dismissed as rare and unlikely, due to both empirical evidence and theoretical concerns. The major obstacle to the broader acceptance of sympatric speciation appears to be theoretical: namely, how reproductive isolation could evolve in the presence of gene flow, which typically acts to homogenize the genomes of diverging sister species [1, 3].

In a previous study, we used individual-based computer simulations and mathematical modeling to propose a mechanism that could establish initial premating RI in sympatric speciation [28]. A simple mathematical model was developed to demonstrate how the interplay between ecological selection and sexual selection, driven by nonlinear population dynamics, can lead to the formation of an initial premating reproductive barrier. Based on our results, we proposed a five-stage process of sympatric speciation:

1. A single species subsists on a single habitat. Sexual selection favors the predominant type of mating-bias alleles (low mating-bias alleles).
2. Disruptive ecological selection favors divergent adaptive niche ecotypes and selects against the less-fit hybrids.
3. Under favorable conditions, selection against high-mating-bias alleles is reversed. They are now positively selected to invade and establish fixed-point polymorphism, resulting in premating RI.
4. This initial premating RI reduces gene flow between niches and facilitates the recruitment of other late-stage pre-zygotic and post-zygotic RI mechanisms, such as adaptive coupling, chromosomal inversions, and BDM incompatibilities, to complete the speciation process.
5. Sympatric speciation is complete. Intraspecific sexual selection again favors low-mating-bias alleles and selects against high-mating-bias alleles in the reproductively isolated sister species. The speciation process returns to stage one.

It has been postulated that the mechanisms responsible for reproductive barriers during the early stages of speciation differ from those that create and reinforce reproductive isolation in the later stages [20, 21]. An initial barrier mechanism creates an environment of reduced gene flow that is conducive to coupling with other barrier mechanisms to complete the speciation process. For instance, adaptive coupling with other one- and two-allele models of premating barriers becomes easier when the initial mating-bias barrier system is convergent—i.e. when it has established mating-bias-allele polymorphism and a certain degree of premating RI (Figs 2a and 7). Similarly, chromosomal inversions and the BDM mechanism operate most effectively when there is already reduced gene flow between niche ecotypes (Figs 19 and 23).

The aim of our current study is to investigate the coupling of additional reproductive barriers during the later stages of sympatric speciation, after an initial premating barrier has been established in an earlier stage by our proposed mathematical model (stages 1 to 3 of our proposed speciation process). For the initial barrier mechanism, we used a mathematical model of two mating-bias alleles without viable hybrids, as described in our previous study [28]. We then developed MATLAB GUI applications to explore the coupling of this initial barrier with other premating and post-mating barrier mechanisms to complete the speciation process (stages 4 and 5 of the process). Our results demonstrate that coupling in late-stage sympatric speciation can produce the multiple distinct, coupled barriers characteristic of completed species, as observed in empirical studies [15-19].

Fig M3 shows a schematic overview of our mathematical model, highlighting key intervention points where system dynamics and reproductive isolation can be modified. Mechanisms that influence the parameter *α* modulate mating biases and affect premating RI between niche ecotypes, while those altering the parameter *f* impact hybrid viability and post-mating RI. The number of matching rounds (*n*), which contributes to assortative mating costs, and the relative niche carrying capacities (*NA* and *NB*) also play a role in shaping system outcomes. The presence of multi-locus ecological and mating-bias genotypes may produce viable hybrids that hinder system convergence and speciation. This effect can be mitigated by chromosomal inversions that suppress hybrid formation [30]. Lastly, encounter probabilities between genotype groups are subject to modification by factors such as habitat preference mutations, movement barriers, and search costs [28, 29].

Our analysis of coupling is structured in two sections: coupling in the early stages and coupling in the later stages of sympatric speciation. Early-stage coupling examines the coupling of our initial premating barrier with additional one- and two-allele models of premating barriers. Late-stage coupling focuses on barriers that reduce hybrid viability, chromosomal inversions, and the BDM mechanism of divergence and genetic incompatibilities. The influences of habitat preference barriers and the presence of multi-locus ecological and mating-bias genotypes are explored in our separate papers [29, 30]. Table I shows a summary of all the barrier mechanisms analyzed in this study.

### 1. Adaptive coupling with different early-stage premating barriers can progressively increase premating reproductive isolation

Our findings demonstrate that once an initial premating barrier is established, its coupling with additional one-allele and two-allele premating barrier mechanisms to further strengthen RI becomes easier. This coupling is adaptive and driven by the loss of maladaptive hybrids under disruptive ecological selection. As long as hybrid loss persists, selection pressures will continue to favor the coupling of additional premating barriers to stem this loss.

Moreover, our results indicate that additional barrier mechanisms are more likely to couple with a convergent mating-bias system that has already achieved fixed-point polymorphism and premating RI (Figs 1a and 1b). In contrast, converting a divergent mating-bias system without fixed points into a convergent one with fixed points is much more difficult. Typically, larger starting populations, stronger mating biases, and lower assortative mating costs are required (Figs 2a, 2b, and 7).

In general, coupling with a one-allele mechanism of premating barrier is easier than with a two-allele mechanism. This is because the one-allele barrier mechanism is not subject to the homogenizing effects of gene flow (Fig 6).

Because the driving force behind adaptive coupling is maladaptive hybrid loss, coupling additional barrier mechanisms becomes more difficult as overall RI becomes stronger. Consequently, the order in which barriers are coupled can affect the final level of RI achieved. Coupling strong barriers early in the process can generate significant RI that potentially depletes the driving force to couple additional weak barriers (Fig 5).

When attempting to couple multiple premating barrier mechanisms and when hybrid loss is already low due to strong overall RI, an “invasion-resistant” pattern frequently emerges in the phase portrait solutions (Fig 2a, 3a). In such systems, small mutant populations near the origin of the phase portrait are unable to invade and subsequently go extinct. Larger initial populations are required to successfully invade and converge toward the system’s fixed points to establish RI. Theoretically, larger invading allele populations can arise in scenarios of secondary contact, when a population that has undergone geographical separation and allopatric divergence returns with a substantial ratio of foreign alleles to encounter the original population in sympatry.

The invasion-resistant pattern typically emerges in the phase portrait solution of a two-allele mating-bias barrier when inter-niche gene flow and hybrid loss are already significantly reduced prior to coupling. To understand the underlying dynamics, consider the invasion of a mutant allele in a model with two mating-bias alleles, *X* and *Y*. In a sympatric population where only the *Y* mating-bias allele exists, the invasion fitness of a higher mating-bias mutant allele, *X*, arising as a small *Ax* genotype population (*Ax* = 0.01) in niche *A*, is determined by the balance between its fitness advantage in inter-niche mating and its fitness disadvantage in intra-niche mating.

In inter-niche mating, the higher mating bias of the *Ax* genotype reduces mating with the *By* genotype, resulting in lower hybrid offspring loss and conferring a fitness advantage to the *Ax* genotype over the *Ay* genotype in niche *A*. Conversely, the higher mating bias of the *X* allele imposes a fitness disadvantage on the *Ax* genotype during intra-niche mating with the predominant *Ay* genotype. This disadvantage arises because intra-niche sexual selection favors the predominant low-mating-bias genotype in the population while selecting against minority genotypes with higher mating biases. Consequently, the *Ax* genotype can only gain a net fitness advantage to invade niche *A* if its fitness advantage in inter-niche mating outweighs its fitness disadvantage in intra-niche mating.

Reducing inter-niche gene flow prior to coupling decreases the frequency of inter-niche mating, thereby reducing the fitness advantage of the *Ax* genotype in inter-niche mating. At the same time, the frequency of intra-niche mating increases, resulting in a greater fitness disadvantage for the *Ax* genotype due to intra-niche sexual selection. These dynamics can contribute to the emergence of an invasion-resistant pattern in the *Ax*/*Bx* phase portrait. In our simulations, this invasion-resistant pattern is generally—but not always—eliminated by increasing the number of mating rounds, *n*, which reduces the cost of assortative mating and mitigates intra-niche sexual selection against the smaller *Ax* population (Fig 3b).

Similarly, decreasing the niche size *NA* may also eliminate the invasion-resistant pattern in the *Ax*/*Bx* phase portrait. Decreasing the *NA* population ratio shifts most niche-*A* mating encounters to inter-niche rather than intra-niche, resulting in greater invasion fitness for the mutant *Ax* genotype due to increased inter-niche matings. Consequently, the mutant allele is under greater selection pressure to invade and proliferate. However, if the *X* allele is also present in niche *B* (i.e., *Bx* > 0), inter-niche mating between *Ax* and *Bx* genotypes can lead to greater hybrid offspring loss for the mutant *Ax* genotype and hinder its invasion.

In general, decreasing the niche size *NA* increases the fitness advantage of the *Ax* mutant due to increased inter-niche mating. This allows the *Ax* mutant to invade more quickly (i.e., requiring fewer generations to reach a fixed-point polymorphism), but it tends to result in weaker RI. Conversely, when niches *A* and *B* are roughly equal in size, an invading *Ax* mutant takes longer to reach a fixed-point polymorphism, but the fixed point tends to lie closer to the bottom-right corner of the *Ax*/*Bx* phase portrait, resulting in stronger RI.

The observation that a smaller niche size, *NA*, enhances the invasion fitness of a mutant *Ax* genotype—by increasing inter-niche mating and reducing intra-niche sexual selection—may be significant for the evolution of RI during adaptive radiation. In the early stages of adaptive radiation, a diverging population that migrates into a new niche often remains small, either due to niche constraints or incumbent selection imposed by a larger sympatric population [28]. This limited population size facilitates the invasion of high-mating-bias mutant alleles, promoting the establishment of premating RI and enabling the incipient species to escape incumbent selection from the dominant group.

Overall, our simulations demonstrate that preexisting reproductive barriers that reduce gene flow can facilitate the coupling of subsequent two-allele mating-bias barriers. This occurs because gene flow tends to homogenize allele ratios across niches and prevent their differential assortment. However, when a prior reproductive barrier substantially reduces gene flow and hybrid loss, it may produce an invasion-resistant pattern in the phase-portrait solutions of the mating-bias barrier. Increasing the number of mating rounds, *n*, or decreasing the carrying capacity of the niche where an invading mutant allele arises (e.g., *NA*) can mitigate or eliminate this invasion-resistant pattern, with minimal impact on the increased RI resulting from reduced gene flow caused by the first barrier. Similar results were also observed in a related study examining the coupling of a two-allele mating-bias barrier following the establishment of an initial habitat preference barrier [29].

A first premating barrier system that reduces gene flow also decreases hybrid loss in inter-niche matings. This diminishes the fitness advantage of an invading mutant mating-bias allele in inter-niche matings compared to its same-niche counterparts and may lead to the emergence of an invasion-resistant pattern in the phase portrait of the mating-bias barrier. In contrast, when initial gene flow is reduced solely by external disruptive ecological selection (i.e., when *f* < 0.5), gene flow between niches decreases, but hybrid loss increases as disruptive selection intensifies. Since the fitness advantage of a mutant mating-bias allele in inter-niche matings remains unaffected, an invasion-resistant pattern is less likely to develop.

Our simulation results confirm that multiple weak barriers can couple to produce strong overall RI (Fig 4). These findings support the hypothesized mechanisms of adaptive coupling [22, 47, 48]: First, genotypes with coupled barriers gain a selective advantage from their combined barrier effects. As shown in Figs 3c and 4, genotypes with coupled barriers benefit from a stronger overall mating bias, which leads to reduced inter-niche mating, lower hybrid offspring loss, and an enhanced fitness advantage, enabling them to invade and multiply. Second, the increase in overall barrier strength through coupling results in a greater selective advantage for the entire group of coupled barriers. Consequently, individual barriers within the group experience stronger selection than they would in isolation. This allows genes associated with weak reproductive barriers to acquire a disproportionate selective advantage that enables them to invade and proliferate by hitchhiking with other adaptive barrier genes.

Lastly, our results show that both one-allele and two-allele mechanisms of adaptive premating barriers can couple to produce strong reproductive isolation. However, this RI, driven by population dynamics, can be easily reversed when external ecological selection is reduced or removed (Fig 8). This underscores the nonpermanent and reversible nature of adaptive coupling [31]. While premating RI is favored over post-mating RI in speciation because it avoids the costly production of nonviable offspring, our findings suggest that more permanent and irreversible barrier mechanisms, such as chromosomal rearrangements or mutations leading to BDM incompatibilities, are necessary to establish secure RI between species.

### 2. A mutation f ^_^ that reduces hybrid viability faces a fitness disadvantage and cannot invade or increase RI without coupling with a barrier mechanism that provides a fitness advantage

Next, we investigated how an initial mating-bias barrier may couple with another barrier mechanism that reduces post-zygotic hybrid viability. In this second mechanism, the presence of a mutation, *f*_**(−)**_ or *f*_**(+)**_, in an individual’s genotype alters the offspring return ratio, *f*, during inter-niche mating. In other words, the mutation affects hybrid offspring compatibility between parents from different niches.

Both the one- and two-allele models of the barrier mechanism demonstrate that genotypes carrying a mutation, *f*_**(−)**_, that reduces inter-niche offspring viability suffer a fitness disadvantage compared to their same-niche cohorts without the mutation. This disadvantage arises due to increased offspring loss during inter-niche mating. Therefore, the *f*_**(−)**_ mutation, on its own, cannot invade or increase post-mating RI (Fig 9a). In contrast, a mutation, *f*_**(+)**_, that increases hybrid viability confers a fitness advantage and can readily invade, resulting in an increased effective value of *f* and a reduction in overall RI (Figs 9b, 9c, and 10).

These results imply that when two niche ecotypes exist in fixed-point mating-bias-allele polymorphism, their premating RI can be reduced or eliminated by the invasion of mutant alleles that create a higher effective value of *f*, i.e., by increasing hybrid viability. It all depends on whether the disruptive ecological selection is environmental (extrinsic) or genetic (intrinsic). If the environment does not provide any food resources for the hybrid ecotypes, then no adaptive changes in the genomes of the hybrids can increase their viability. Conversely, environmental food resources may exist for the hybrids, but the problem is that the hybrids do not have the optimal genotypes to utilize those resources. In this scenario, an adaptive mutation may increase the ability of the hybrids to extract the food resources and increase their viability, destroying the existing premating RI in the process. Therefore, to achieve sympatric speciation, it is necessary to have environmental disruptive ecological selection that limits the number of hybrids that environmental resources can support.

A negative-fitness mutation, *f*_**(−)**_, that reduces hybrid viability can only invade if it is linked to a positive-fitness barrier that offsets its disadvantage [2]. For instance, a super-allele that induces strong mating bias (low *α*) and reduced hybrid viability (low *f*) may still confer a net fitness advantage, enabling it to invade and create stronger overall premating RI (Fig 11a). In this case, the benefit of strong mating bias compensates for the disadvantage of reduced hybrid viability [49]. Such a dual-action super-allele could arise through chromosomal inversions, close physical linkage of genes controlling mating bias and hybrid viability, relocation of coupled alleles to low-recombination zones, pleiotropic actions of recombination suppressors, or the BDM mechanism of coevolving mating-bias and hybrid incompatibilities.

If a mutation *f*_**(−)**_ that reduces hybrid viability is able to rise to near fixation, the intrinsic hybrid incompatibilities it creates between niche populations could potentially take over to ensure continued hybrid loss and maintain RI when external disruptive ecological selection is reduced (Fig 11b). However, this resilience against reversibility is not absolute, as reproductive isolation typically collapses when external disruptive ecological selection is completely removed.

### 3. Chromosomal inversions can play an important role in sympatric speciation

Chromosomal inversions have long been speculated to play a key role in sympatric speciation. In studies of *Drosophila*, sympatric species pairs differ by a greater number of chromosomal inversions compared to allopatric species pairs. Moreover, the divergent gene loci that mediate RI between sympatric species pairs are located within these inversions. This contrasts with closely related allopatric species pairs, where the barrier gene loci mediating RI are scattered throughout the genome [33, 50, 51].

Chromosomal inversions can facilitate speciation by capturing favorable combinations of locally adaptive ecological alleles, mating-bias alleles, and alleles mediating genetic incompatibilities within their inverted regions, thereby preventing their breakdown during heterokaryotype mating [33, 36, 37, 52]. Coadaptation (epistasis) between the captured loci is not needed [34]. Acting as recombination suppressors, CIs preserve linkage disequilibrium among captured alleles from the homogenizing effects of gene flow, effectively transforming them into super-alleles with advantageous properties [35, 39, 40]. Several weakly adaptive alleles could be combined into a super-allele with a large effect. The group of linked functional genetic elements then segregates as a single Mendelian locus. Through such mechanisms, CIs can produce both premating and post-mating RI in sympatric speciation.

Chromosomal inversions are regarded as a late-stage barrier mechanism, primarily because they are more likely to capture locally adaptive alleles once differential assortment of favorable ecological and mating-bias alleles has already occurred between niches due to earlier barrier mechanisms [42]. The fitness advantage of a CI arises from the beneficial combination of alleles it captures, while CIs that capture deleterious alleles tend to experience a fitness disadvantage. Over time, more fit CIs may emerge to invade and replace less fit CIs [53-55].

Chromosomal inversions are considered an adaptive barrier mechanism: the invasion of an advantageous CI is driven by selection pressure arising from hybrid loss under disruptive ecological selection [35, 42]. As RI between niche populations becomes stronger, it becomes more difficult to couple additional CIs to further reduce hybrid loss and increase RI. Therefore, an environment with an intermediate level of RI—sufficient to permit the assortment of locally adaptive alleles, yet still subject to hybrid loss—appears most optimal for an adaptive CI to invade and exert its effects [42].

Chromosomal inversions exhibit specific properties in heterokaryotype mating that can influence their invasion dynamics. During heterokaryotype mating, single crossover events in the inverted regions result in 50% nonviable offspring [38, 43]. The rarer occurrences of double crossovers and gene conversions in the inverted regions generate “gene flux,” which can restore recombination and weaken the linkage disequilibrium between captured alleles [44, 45]. In our models, we primarily focus on paracentric CIs, considering the effects of single crossovers only when necessary and disregarding the much rarer impacts of double crossovers and gene conversions.

The successful invasion of a CI mutation depends on the net fitness advantage that it experiences. In inter-niche mating, a mutant CI that captures locally adaptive ecological alleles acquires a fitness advantage over its noninverted same-niche cohorts because it experiences less hybrid offspring loss in between-niche heterokaryotype mating. This fitness advantage is amplified in environments with high gene flow and low RI. Conversely, in intra-niche mating, due to single crossover events in inverted regions, the mutant CI incurs a fitness disadvantage from offspring loss in same-niche heterokaryotype mating. This disadvantage is exacerbated in low gene flow environments, where most mating occurs within niches rather than between niches.

As expected, decreasing the niche size *NA* makes the invasion of a mutant CI capturing locally adaptive niche-*A* alleles more likely. This is because the majority of the CI‘s mating encounters will be inter-niche rather than intra-niche, which can help it gain a net fitness advantage to invade. Once the CI rises to near fixation in niche *A*, the invasion dynamics are reversed. It then becomes more difficult for the noninverted native ecotype to invade due to offspring loss from single crossover events in intra-niche heterokaryotype mating (Fig 14a).

In addition to fitness considerations in inter-niche and intra-niche matings, the invasion fitness of a mutant CI is also influenced by the capture of deleterious or maladaptive alleles, as well as mutations that alter hybrid and mating-bias incompatibilities. When multi-locus mating-bias genotypes are present, CIs can combine high-mating-bias alleles into super-allele complexes that reduce the production of mating-bias hybrids and provide the CIs with a fitness advantage to invade and increase overall RI [30].

In our models, once an adaptive CI gains a net fitness advantage, it invariably rises to fixation. This occurs because our models assume there are no viable hybrids. Producing CI polymorphism requires the presence of viable hybrids in the models, which entails creating hybrid niche food resources to support hybrid genotypes. This approach is explored in a separate study modeling the effects of viable ecological and mating-bias hybrids in systems with multi-locus ecological and mating-bias genotypes [30]. The results demonstrate that introducing CI polymorphism does not alter the fundamental processes driving the invasion dynamics of a mutant CI.

### 4. Chromosomal inversions capturing locally adaptive ecological alleles can create negative coupling that reduces premating RI

A mutant CI that captures all of the locally adaptive ecological alleles in a niche is at a fitness advantage to invade because it suffers no hybrid offspring loss in inter-niche mating, compared to the native noninverted ecotypes in the same niche (Fig 12a and 12b). The effective offspring return ratio, *Fk*2, is 0.5 for the mutant CI (discounting the rare event of single crossovers in its inverted region during heterokaryotype mating). When the CI becomes fixed, it destroys inter-niche disruptive ecological selection and premating RI. This creates a situation where there is no gene flow between the ecological loci in the inverted regions, but there is free gene flow in the rest of the genome between the niche populations. Because only the locally adaptive ecological genes are reproductively isolated, this mechanism could potentially explain the emergence of genomic islands of divergence—regions of gene differentiation and reduced recombination—when comparing the genomes of closely related species.

The implication is that a sympatric population could evolve into two freely interbreeding yet ecologically distinct ecomorphs that no longer produce intermediate ecological hybrids. This raises the question of why such discrete ecomorphs in panmictic populations are not more commonly observed in nature. One possible explanation is the difficulties of capturing all locally adaptive alleles within a single chromosomal inversion and achieving an *Fk*2 value of 0.5.

If a mutant CI captures only a partial set of the locally adaptive alleles in a niche, the offspring return ratio, *Fk*2, decreases below 0.5. In this case, hybrid offspring are produced during the CI‘s inter-niche mating. The value of *Fk*2 can be further reduced if the CI captures a *f*_**(−)**_ mutation that decreases hybrid viability. Hybrid incompatibilities mediated by the BDM mechanism, as well as the effects of crossovers and gene conversions in the inverted regions, can also decrease the value of *Fk*2. There is evidence that, in diploid organisms, some heterokaryotype hybrids may exhibit reduced fitness [56-59].

The results of our simulations show that a mutant CI capturing a partial set of ecological alleles in a niche can only invade if its associated offspring return ratio, *Fk*2, exceeds the original offspring return ratio, *f*. When *Fk*2 is 0.5, the system is divergent, and no premating RI exists (Figs 12a and 12b). As the value of *Fk*2 gradually decreases toward *f*, the system can converge, but with a resultant *Ax*/*Bx* fixed point that is shifted further away from the bottom right corner of the *Ax*/*Bx* phase portrait, indicating weaker premating RI compared to the initial premating RI before coupling (Figs 13a and 13b). When *Fk*2 = *f*, the *Ax*/*Bx* fixed point is the same as it was before the appearance of the CI. If *Fk*2 drops below *f*, the CI fails to invade, and the *Ax*/*Bx* phase portrait remains unchanged (Fig 13c). When *Fk*2 > *f*, the outcome exemplifies “negative coupling,” where coupling with an additional barrier mechanism leads to the weakening or loss of preexisting RI [32].

The invasion success of a mutant CI capturing locally adaptive alleles is the combined net effect of its fitness advantage in inter-niche mating and its fitness disadvantage in intra-niche mating. When RI between niches is strong, most matings occur within niches rather than between niches. A small population of locally adaptive CIs is at a fitness disadvantage to invade due to hybrid loss in intra-niche heterokaryotype mating, caused by single crossover events in the inverted regions (Fig 14a). However, once the CI becomes the predominant genotype in the niche, the invasion dynamics are reversed. The original, noninverted genotypes then become the mutant CIs that are trying to invade despite their fitness disadvantage in intra-niche heterokaryotype mating due to single crossover events in the inverted regions.

These intra-niche selection dynamics of chromosomal inversions resemble those seen in sexual selection: the predominant variant is positively selected to eliminate the less prevalent variants. These observations suggest that in late-stage sympatric speciation, once an advantageous CI has become fixed, it is difficult to reverse, as the original, noninverted variant is treated like a new mutant CI that is at a fitness disadvantage to invade the established CI population.

### 5. Chromosomal inversions that capture locally adaptive ecological alleles and mating-bias alleles can generate RI that extends across entire genomes

A CI that captures locally adaptive ecological alleles along with the prevalent mating-bias allele in a niche can invade and establish RI across the entire genome [60-62]. First, consider the case where a mutant CI is able to capture all of the locally adaptive ecological alleles in niche *A*, as well as its most prevalent mating-bias allele, *X*, after an earlier mating-bias barrier has initiated limited nonrandom assortment of these favorable alleles. Based on our previous analyses of CIs that capture all locally adaptive ecological alleles, such a mutant CI has an *Fk*2 value of 0.5, allowing it to invade and become fixed within niche *A*, thereby sweeping the mating-bias allele *X* to fixation due to their linkage association. In this scenario, if the niche ratio *NA* is greater than *NB*, the fixed CI population in niche *A* serves as a source that continuously supplies the population in niche *B* with *X* alleles through inter-niche matings, ultimately driving the *Y* allele in niche *B* to extinction and leaving only the *X* allele in the system (Fig 15). Conversely, if the niche ratio *NB* is greater than *NA*, the more numerous *Y* allele in niche *B* would typically drive the less abundant *X* allele in niche *A* to extinction. However, because the *X* alleles in niche *A* cannot be invaded due to their linkage association with the fixed CI population, a mating-bias polymorphism, *Ax* = 1 and *Bx* = 0, emerges in the phase portrait, resulting in strong premating RI between the niche populations (Figs 16a and 16b). Therefore, when a CI captures all of a niche’s adaptive ecological alleles along with the prevalent mating-bias allele, the emergence of premating RI is contingent on the relative niche ratios of *NA* and *NB*.

It should be noted that even when *NB* is greater than *NA*, large values of *α* and a value of *NA* close to 0.5 can still destroy the *Ax*/*Bx* polymorphism, leaving only the *X* allele in the system (Fig 17a). This occurs because large values of *α* and *NA* increase inter-niche matings for niche-*B* individuals. Consequently, sexual selection against the *X* alleles in niche *B* is reduced, while sexual selection against the *Y* alleles in niche *B* is increased, which could result in the elimination of the *Y* alleles. Eliminating sexual selection by increasing the number of mating rounds, *n*, causes a stable polymorphism of *X* and *Y* alleles to exist in niche *B* (Fig 17b). Therefore, in addition to the relative niche ratios, the creation of premating RI by a CI that captures all adaptive ecological alleles and a favorable mating-bias allele could also depend on the extent of inter-niche matings and the strength of sexual selection.

In contrast to a CI that captures all adaptive ecological alleles, a mutant CI that captures only a partial set of locally adaptive ecological alleles and a favorable mating-bias allele can invade and establish mating-bias-allele polymorphism and premating RI, regardless of relative niche sizes (Figs 18a and 18b). When a CI captures only a subset of adaptive alleles within a niche, the effective *Fk*2 associated with the CI falls below 0.5. Hybrid loss and disruptive ecological selection are restored in the system, which again enables the differential assortment of mating-bias alleles between niches to develop premating RI and stem this hybrid loss. This appears to be the most effective way for a mutant CI to invade and couple with a preexisting barrier to strengthen overall RI. These findings are consistent with previous research showing that pseudomagic traits may be more effective than magic traits in promoting RI and speciation in the presence of gene flow [63, 64].

Our simulation results show that when a mutant CI captures a subset of locally adaptive alleles in niche *A*, along with a prevalent mating-bias allele *X*, and has an associated niche return ratio *Fk*2 > *f*, the CI gains a fitness advantage over its noninverted same-niche cohorts due to reduced hybrid offspring loss from inter-niche mating. Consequently, the CI can successfully invade and become fixed in niche *A*, carrying with it the linked *X* allele to produce premating RI across the entire genome. This can occur even if the initial mating-bias barrier system is divergent, i.e., without an initial *Ax*/*Bx* fixed-point polymorphism (Fig 19). The challenge lies in capturing favorable combinations of ecological and mating-bias alleles without also capturing maladaptive alleles in a divergent system. Conversely, in a convergent system, a high degree of differential assortment of favorable alleles between niches already exists, making their capture easier.

In contrast to the scenario where a mutant CI capturing only adaptive ecological alleles cannot invade if *Fk*2 is less than *f*, a mutant CI capturing a partial set of locally adaptive alleles (*Fk*2 < 0.5) and the predominant mating-bias allele in a niche can invade even when *Fk*2 is less than *f*, albeit to a limited extent. This is because, in late-stage sympatric speciation, when a high degree of differential assortment of mating-bias alleles between niches is already established, the CI’s linkage with the predominant mating-bias allele reduces inter-niche mating relative to its noninverted same-niche counterparts that still carry a polymorphism of mating-bias alleles. This decrease in inter-niche offspring loss, due to linkage with the predominant mating-bias allele, can provide the CI with a fitness advantage to counteract the fitness disadvantage of increased hybrid offspring loss when *Fk*2 < *f*. This represents a case where a CI can function like a supergene, enabling an underdominant *f*_**(−)**_ mutation that reduces *Fk*2 to hitchhike alongside advantageous ecological and mating-bias alleles, synergistically enhancing RI. However, as the value of *Fk*2 decreases further below *f*, hybrid loss becomes substantial enough that the CI suffers a net fitness disadvantage compared to its noninverted same-niche cohorts, and it can no longer invade.

We can call the lower limit of *Fk*2 that still enables the CI to invade and produce premating RI the “lower invasion threshold,” and the difference between *f* and this lower invasion threshold, *f* − *Fk*2, the *“*invasion buffer range.*”* In general, reducing the value of *α* (increasing the mating bias) decreases the lower invasion threshold for *Fk*2 and increases the invasion buffer range. However, as *α* approaches zero, invasion becomes more difficult as the strong premating bias does not provide enough hybrid loss to drive the invasion of the mutant CI. Conversely, increasing the value of *f* decreases the invasion buffer range because the noninverted genotypes suffer less hybrid loss from inter-niche mating. At the limit, when *f* = 0.5, the invasion buffer range is zero because there is no hybrid loss from inter-niche mating. These results imply that the invasion of an adaptive CI is most favored in late-stage sympatric speciation when the values of *α* and *f* are low, indicating strong mating-bias (premating RI) and significant hybrid loss (post-mating RI).

Having established the lower limit of *Fk*2 below *f* at which a mutant CI capturing adaptive ecological and mating-bias alleles can invade, we can show that there is also an upper limit of *Fk*2, above which the mutant CI can invade but cannot establish mating-bias-allele polymorphism and premating RI. This observation follows logically from our previous finding that a CI capturing all of the locally adaptive ecological alleles (*Fk*2 = 0.5) destroys any preexisting premating RI. Therefore, this upper limit of *Fk*2, above which the mating-bias system is divergent, is found close to *Fk*2 = 0.5. Increasing the value of *α* slightly raises this upper limit of *Fk*2, while decreasing the value of *α* slightly lowers it. In sum, it appears that for any given value of *f*, there exists an optimal range of *Fk*2 within which the CI can invade and achieve fixed-point mating-bias-allele polymorphism and premating RI. When *f* = 0.5, this range decreases to zero.

Another variable that can affect the invasion dynamics of the mutant CI is *Fk*1, the offspring return ratio of intra-niche heterokaryotype mating. An *Fk*1 value less than 0.5, due to rare single crossover events in the inverted regions, can create a fitness disadvantage for a mutant CI and hinder its invasion (Fig 14a).

In summary, our results demonstrate that the most favorable conditions for a CI to couple and increase RI occur when the invading CI captures a partial set of locally adaptive ecological alleles and the prevalent mating-bias allele in a niche during late-stage sympatric speciation, when strong mating bias (low *α*) and significant hybrid loss (low *f*) are present. For any given value of *f*, there exists an optimal range of *Fk*2—bounded by *Fk*2 close to 0.5 and a *“*lower invasion threshold” where *Fk*2 ≤ *f*—within which the CI can successfully invade and enhance RI.

### 6. The presence of viable hybrids facilitates the emergence of chromosomal inversion polymorphism

In our model, an invading mutant CI either goes extinct or reaches fixation (see Figs 12–19) without the possibility of existing in a state of fixed-point polymorphism. This is because, in our model, even though the ecological selection factor *f* can be used to compute the effects of having both viable and nonviable hybrids under disruptive ecological selection, the model does not actually allow for the existence of viable hybrid populations in the system because no ecological niche resources are allocated to support their existence.

As it stands, a mutant CI that captures locally adaptive ecological alleles will always have a fitness advantage over the native, noninverted genotypes in its niche. Consequently, the mutant CI is bound to multiply and eliminate the noninverted genotypes in the niche. The only way to achieve CI polymorphism is to have viable hybrid populations in the system that can continuously regenerate and resupply the niche with noninverted genotypes. A balance between selection and immigration can then maintain CI polymorphism. However, this is only possible if hybrids can survive in pre-allocated hybrid niches, so they are not eliminated due to a lack of food resources. In the extreme case of a large hybrid swamp, the viable hybrids in the system may even regenerate enough noninverted genotypes in the niche to overwhelm and eliminate the CI. In the real world, CI polymorphism is likely more common than complete fixation due to the presence of viable hybrids, balancing selection, or frequency-dependent selection [38, 54, 65-68].

In a separate study, we developed MATLAB GUI programs for models that can incorporate any number of gene loci for ecological and mating-bias genotypes. Our results confirmed that CI polymorphisms are prevalent when viable ecological and mating-bias hybrid populations are permitted to exist in the model [30].

When a chromosomal inversion coexists in polymorphism with noninverted genotypes within a niche, the genes within its inverted region can evolve and diverge independently, without the hindrance of recombination. Essentially, these genes are isolated from the rest of the noninverted genome, free to follow their own evolutionary trajectories. These protected genes can evolve beneficial adaptations faster than their noninverted counterparts in the genome, where beneficial genetic changes are susceptible to disruption by recombination [41]. Consequently, fitter CI variants may emerge, potentially invading and replacing less adaptive CI variants and noninverted genotypes. Conversely, without recombination, CIs that acquire deleterious or maladaptive mutations cannot purge these mutations through purifying selection. Consequently, Muller’s ratchet can lead to the accumulation of maladaptive mutations, giving these CIs a fitness disadvantage and making them prone to elimination [55, 66]. These adaptive advantages and fitness disadvantages are most pronounced when the CI population ratio is low within the polymorphism and disappear once the CI becomes fixed.

### 7. Adaptive chromosomal inversions can protect the established reproductive isolation from reversal in late-stage sympatric speciation

After an advantageous CI capturing a partial set of locally adaptive ecological alleles in a niche, along with the predominant mating-bias allele, becomes fixed during late-stage sympatric speciation, the CI population exhibits a degree of resistance to reversal. This is evident in the post-invasion phase portraits, after a high ratio of CI individuals in the niche and strong premating RI between niches have already been established, resulting in stable CI and mating-bias fixed points. These fixed points are surrounded by basins of attraction, which protect the CI population from invasion by native, noninverted genotypes. Any small deviations from the fixed points caused by invading noninverted populations are drawn back to the fixed points, ultimately leading to the extinction of the invaders. These protective basins of attraction around the fixed points persist even if the value of *f* is increased, i.e., when disruptive ecological selection is reduced (Fig 20). In general, decreasing the value of *α* (increasing the mating bias) and increasing the value of *n* (decreasing the assortative mating cost) can expand the invasion-resistant regions surrounding the fixed points. Since a noninverted mutant is treated as a new CI by the existing, established CI population, a lower value of *Fk*1 hinders its invasion (Fig 14a).

Just as the invasion of a mutant CI that captures locally adaptive ecological and mating-bias alleles is limited by a lower threshold of *Fk*2, below which the CI cannot invade, the invasion of a noninverted mutant into a fixed CI population with high premating RI is similarly restricted by a lower threshold of *Fk*2, above which the noninverted mutant cannot invade. If *Fk*2 is below this threshold, a noninverted mutant can invade, become fixed, and eliminate the CI population along with its associated RI. In this way, the invasion dynamics of a noninverted mutant mirror those of a CI mutant invading a noninverted population, though the invasions are constrained in opposite directions with respect to the *Fk*2 value.

In a population consisting entirely of a fixed CI, a noninverted mutant attempting to invade can be viewed as a new CI trying to invade a noninverted population (from the mutant’s perspective), and the value of *f* can be treated as the *Fk*2 of this new invading mutant CI. However, in this case, the established CI population is linked to the predominant mating-bias allele in the niche, while the invading noninverted genotype is not linked to any mating-bias allele. This linkage association can give the existing CI population a fitness advantage over the unlinked noninverted mutant genotype, making its invasion more difficult.

Even when disruptive ecological selection is completely removed (*f* = 0.5), similar protective basins of attraction around the fixed points are observed in the phase portraits following the fixation of a CI that captures a partial set of locally adaptive ecological alleles and the predominant mating-bias allele in late-stage sympatric speciation (Figs 21a and 21b). This confers resistance to invasion by noninverted genotypes and prevents reversal of the established CI. In this scenario, since *f* = 0.5, the noninverted genotypes experience no hybrid loss during inter-niche mating, while the established CI population incurs a fitness disadvantage due to offspring loss in inter-niche mating from an *Fk*2 less than 0.5. However, despite this fitness disadvantage, the established CI population can forestall the invasion of noninverted genotypes through intra-niche sexual selection, which favors the predominant mating-bias alleles in a niche. Accordingly, low values of *α* and *n* increase the invasion difficulty of noninverted, unlinked genotypes. A low value of *α*, reflecting strong premating RI, also reduces inter-niche mating and limits inter-niche offspring loss for the CI population, offsetting its fitness disadvantage. Since the CI population is the predominant genotype in the niche, an *Fk*1 value less than 0.5 also impedes invasion by noninverted genotypes.

Similar to the invasion dynamics of an adaptive CI into a noninverted population, for any given value of *f*, there exists a lower threshold of *Fk*2 above which the invasion of a noninverted genotype into an established CI population becomes impossible (Fig 21c). The difference between *f* and *Fk*2, *f* − *Fk*2, which we call the *“*invasion resistance range,*”* acts as a protective buffer that shields the fixed CI from invasion. Decreasing the value of *α* increases the invasion resistance range (Fig 21d). In the limiting case, when *α* is zero, corresponding to complete premating RI, no invasion by a noninverted mutant is possible, regardless of the value of *Fk*2. These findings support our hypothesis that in late-stage sympatric speciation, when RI is well-established, a fixed CI linking locally adaptive ecological and mating-bias alleles becomes resistant to invasion and reversal.

If *Fk*2 is increased above a certain limit, close to a value of 0.5, the fixed CI population remains resistant to invasion by noninverted mutants; however, the *Ax*/*Bx* phase portrait becomes divergent, and the premating RI disappears (Fig 21e). This likely occurs because the elevated *Fk*2 value significantly weakens disruptive ecological selection, resulting in insufficient hybrid loss to drive the *Ax*/*Bx* system to convergence. Therefore, there appears to be a lower limit of *Fk*2, below which noninverted mutants can successfully invade and destroy existing RI. Conversely, an upper limit of *Fk*2 exists, beyond which the system remains resistant to invasion but cannot sustain premating RI. Between these extremes, an intermediate range of *Fk*2 maintains fixed polymorphisms and RI, while remaining resistant to invasion and reversal.

Theoretically, these CI invasion dynamics imply that if premating RI is not complete, the system could be repeatedly invaded by mutant CIs that capture increasingly higher numbers of locally adaptive ecological alleles. Conversely, if a CI captures deleterious or locally maladaptive alleles, it suffers a fitness disadvantage compared to its same-niche cohorts and is likely to be eliminated. Having a higher number of locally-adaptive ecological alleles in a CI usually means a higher value of *Fk*2 associated with the CI. This could translate into a fitness advantage for the mutant CI to invade due to its reduced inter-niche hybrid offspring loss relative to its same-niche cohorts with a lower value of *Fk*2.

The invasion of such CIs with high values of *Fk*2 could potentially destroy the existing premating RI (i.e., negative coupling [32]), especially when *NA* > *NB*, based on our analyses of CIs that capture all adaptive ecological alleles along with a favorable mating bias allele (Fig 15). When a CI captures all of the locally adaptive alleles in a niche, the effective value of *f* is 0.5, and there is no longer any disruptive ecological selection in the system.

Hopefully, in late-stage sympatric speciation, the strong RI between niches produced by the existing CI and other coupled barriers, combined with the difficulty of capturing all locally adaptive alleles— especially when these alleles are located on different chromosomes—makes the emergence and invasion of such CIs with high *Fk*2 values unlikely. Additionally, rare events such as single and double crossovers and gene conversions in inverted regions during heterokaryotype mating—so far largely ignored in our models—as well as decreased viability of heterokaryotype offspring, may act to ensure that hybrid loss is not reduced to zero [38, 43-45, 56, 57]. When CIs exist in a polymorphic state rather than fixation, hybrid loss also tends to persist.

### 8. Bateson–Dobzhansky–Muller (BDM) incompatibilities are a late-stage, nonadaptive barrier mechanism that can create irreversible premating and post-mating RI

Of all the barrier mechanisms that we have studied so far, the BDM mechanism of divergence and incompatibilities appears to be the slowest to evolve and least reversible because it involves novel adaptive mutational changes. It is considered a late-stage barrier mechanism in sympatric speciation because it operates most effectively in environments with low gene flow [25, 26]. It makes no difference whether the initial reproductive isolation is caused by geographical barriers, as in allopatric speciation, or by mating-bias premating RI, as in sympatric speciation. Absent the homogenizing effect of gene flow, a niche ecotype is free to adapt, diverge, and develop hybrid and mating-bias incompatibilities as chance byproducts. Our study focuses on BDM divergence driven by differential adaptations to distinct niche environments rather than neutral drift, which typically takes longer to evolve. This approach is reasonable given that disruptive ecological selection creates distinct niche environments for the incipient sister species to adapt.

The BDM mechanism is considered nonadaptive because it is not driven by maladaptive hybrid loss. Instead, it develops incidental hybrid and mating-bias incompatibilities as niche ecotypes evolve new mutations, either through drift or differential adaptations to their distinct environments. Consequently, the BDM mechanism can evolve both premating and post-mating RI in late-stage sympatric speciation to further strengthen and cement overall RI.

In our study, we model the BDM process by introducing a mutant allele *Z* in niche *A* that exhibits properties typically associated with BDM-mediated barriers. We assume that the *Z* allele arises due to better mutational adaptations to the niche-*A* environment. As a result, in each generation, niche-*A* ecotypes carrying the *Z* allele gain an ecological fitness advantage over their same-niche cohorts, and their population ratio is boosted by a factor *Sax*. Conversely, niche-*B* ecotypes carrying the *Z* allele become maladapted to the niche-*B* environment, leading to a reduction in their population ratio by a factor *Sbx*. In BDM divergence, the values of *Sax* and *Sbx* should reflect antagonistic pleiotropy, with *Sax* > 1 to indicate an ecological fitness advantage in niche *A*, and *Sbx* < 1 to reflect maladaptive effects in niche *B*.

We use the variable *F*2 to model the hybrid incompatibility and the variable *α*1x to model the mating-bias incompatibility that arise as chance byproducts of the *Z* allele‘s better ecological adaptations in niche *A* through the BDM process. In inter-niche mating, when one parent carries the *Z* allele and the other does not, their offspring return ratio is governed by *F*2. Similarly, when an individual with the *Z* allele encounters another individual without it, their mating success is determined by the product of the mating-bias value *α* × *α*1x if they carry different mating-bias alleles (*X* or *Y*) and by *α*1x if they carry the same mating-bias allele (*X* or *Y*). Therefore, an *F*2 value less than *f* and an *α*1x value less than 1 indicate the presence of BDM incompatibilities.

Next, we investigate the invasion dynamics of the *Z* allele to determine whether it can couple with an initial premating barrier to strengthen and solidify overall RI. The invasion fitness of the *Z* allele is governed by its properties, derived from the BDM mechanism and represented by the variables *Sax, Sbx, α*1x, and *F*2. It is the net result of the fitness advantages and disadvantages contributed by each of these property variables.

In general, coupling occurs with typical variable values expected from the BDM barrier mechanism in late-stage speciation: antagonistic ecological fitness in different niches (*Sax* > 1 and *Sbx* < 1), mating-bias incompatibility (*α*1*x* < 1), and hybrid incompatibility (*F*2 < *f*). In such instances, the *Z* allele is able to invade successfully and establish fixed-point polymorphism in the *Ax*/*Bx* phase portrait, shift the preexisting *Ax*/*Bx* fixed point toward the bottom right corner in the *Ax*/*Bx* phase portrait, and create stronger overall RI (Fig 22a). However, if the variable values are suboptimal, the *Z* allele either becomes fixed in both niches and leaves the preexisting *Ax*/*Bx* fixed point unchanged, or it fails to invade (Figs 22b and 23d). To understand the factors that determine the invasion outcome requires an understanding of the interaction dynamics among the different system variables.

To maximize RI, we want the *Z* allele to invade and establish a fixed-point polymorphism as close to the lower right corner of the *Ax*/*Bx* phase portrait as possible. To this end, a high value of *Sax*, signifying enhanced mutational adaptations to the niche-*A* environment, gives the *Ax* genotypes a fitness advantage to invade and proliferate in niche *A*. Conversely, a low *Sbx* value, signifying maladaptation caused by the same niche-*A* mutational changes in the niche-*B* environment, suppresses the *Bx* population in niche *B*. Thus, the antagonistic effects of these two variables act to drive the *Ax*/*Bx* fixed point toward the lower right corner, where *Ax* = 1 and *Bx* = 0, in the phase portrait. Overall, more effective mutational adaptation to a niche environment appears to be the primary driving force behind the invasion fitness of a mutant genotype produced by the BDM mechanism, with mating-bias and hybrid incompatibilities arising as secondary byproducts.

As long as there is gene flow between the niches, the *Z* allele will stabilize at a fixed-point *Ax*/*Bx* polymorphism after successful invasion. Reducing the value of *Sbx* decreases *Bx* at the fixed point because the *Z* allele is maladaptive in niche *B*. It also decreases *Ax* at the fixed point because niche *B* acts as a sink that draws in and eliminates the *Z* allele from niche *A* (Fig 23c). Conversely, increasing the value of *Sax* has the opposite effect, as it increases both *Ax* and *Bx* at the fixed point.

Nonetheless, the net effect of widening the difference between *Sax* and *Sbx* is to shift the *Ax*/*Bx* fixed point toward the lower right corner of the *Ax*/*Bx* phase portrait, resulting in stronger premating RI (Fig 23c). This occurs because the *Bx* offspring from interniche mating are less viable in niche *B*, which reduces the gene flow from niche *A*. Since offspring from niche-*A* parents tend to carry the *X* allele, this creates a net loss of *X* alleles in niche *B*. Similarly, inter-niche mating produces niche-*A* offspring genotypes without *Z* alleles. These offspring are less adaptive than the *Ax* genotypes in niche *A*, so they tend to be eliminated, reducing the gene flow from niche *B* as well. Since offspring from niche-*B* parents tend to carry the *Y* allele, this creates a net loss of *Y* alleles in niche *A*. As a result, the *Ax*/*Bx* fixed point shifts toward the lower right corner of the phase portrait as the *Z* allele invades. As expected, this process is most effective in a convergent mating-bias system that already exhibits *Ax*/*Bx* fixed-point polymorphism, i.e., *Ax* ≠ *Bx*. Thus, antagonistic divergence in ecological fitness, represented by *Sax* − *Sbx* and arising from the BDM mechanism of differential adaptations, can independently strengthen RI without changes in any other system variables.

Next, we can examine how the value of *α*1x affects the invasion dynamics of the *Z* allele. If there is mating-bias incompatibility due to BDM divergence, the value of *α*1x will be less than 1, and the *Z* allele can acquire a fitness advantage in inter-niche mating due to decreased hybrid offspring loss and invade, resulting in stronger overall RI. However, as long as *α*1x is not zero—that is, there is gene flow between the niches—the *Z* allele will exist in a state of fixed-point polymorphism in the *Ax*/*Bx* phase portrait (Fig 23a). In general, decreasing the value of *α*1x increases the value of *Ax* and decreases the value of *Bx* at the fixed point, and flattens the vector trajectory curve toward the *x*-axis (Fig 23b). Only when there is no gene flow between the niches (i.e., when *α* = 0 or *α*1x = 0, or *F*2 = *f* = 0) will the ecological advantage of the *Z* allele in niche *A* propel the *Ax*/*Bx* vector trajectory along the *x*-axis and become fixed at *Ax* = 1 and *Bx* = 0.

At times, when the value of *α*1x is reduced below a certain threshold, a small population of mutant *Ax* genotypes cannot invade due to the appearance of an invasion-resistant pattern in the *Ax*/*Bx* phase portrait (Fig 22b). This occurs because the high mating bias created by *α*1x gives the *Ax* genotype a fitness disadvantage in intra-niche mating with the native niche-*A* ecotypes when sexual selection is strong and the initial population of *Ax* is small. This invasion-resistant pattern can be mitigated by reducing intra-niche sexual selection against the *Z* allele, either by increasing the number of mating rounds, *n*, or by increasing the value of *α*1x.

Because the invasion-resistant pattern only acts against small invading mutant populations, it is no longer a concern once the *Z* allele has successfully invaded and become the dominant genotype in niche *A*. This allows the mating bias, *α*1x, to further decrease as the ecotypes diverge over time through the BDM mechanism. Therefore, it appears advantageous for strong mating-bias incompatibility to evolve later rather than earlier in the BDM mechanism, so that the invasion of ecologically adaptive mutants is not hindered by intra-niche sexual selection.

We can also examine how the value of *F*2 affects the invasion dynamics of the *Z* allele. Once hybrid incompatibility has developed as a result of BDM divergence, the value of *F*2 associated with the *Z* allele will be less than *f*. Initially, lowering the value of *F*2 will cause the *Z* alleles to become fixed in both niche *A* and niche *B*, leaving the preexisting *Ax*/*Bx* fixed point and premating RI unchanged (Fig 23d). The *Ax*/*Bx* fixed point remains unchanged because hybrid offspring incompatibility only exists between parents with and without the *Z* allele. If the *Z* allele is fixed in both niches, then the offspring return ratio between niche ecotypes reverts to *f*, instead of *F*2, and the *Ax*/*Bx* system parameters remain the same.

The ability of the *Z* allele to invade and become fixed in both niches can be explained by the following invasion dynamics. First, the better niche-*A* adaptation of the *Z* allele (*Sax* > 1) gives the *Ax* genotype a fitness advantage to invade and increase in population size in niche *A*. This confers a fitness advantage to the *Bx* genotype in niche *B* over the native niche-*B* ecotypes lacking the *Z* allele, as the offspring return ratio is *f* between *Ax* and *Bx* parents versus a ratio of *F*2 between *Ax* and the native niche-*B* parents without the *Z* allele. Because *F*2 is less than *f*, the fitness advantage from reduced hybrid loss is able to overcome the ecological disadvantage of the *Z* allele in niche *B* (*Sbx* < 1) and allow the *Z* allele to rise to fixation in niche *B*. Decreasing the value of *NA* can stop the fixation of the *Z* allele, while increasing *Sax* makes the fixation of the *Z* allele in both niches more likely.

If the value of *F*2 is further decreased, a lower limit is reached below which the mutant *Z* allele can no longer invade, as the fitness disadvantage from increased hybrid incompatibility outweighs the ecological advantage of the *Z* allele in niche *A*. An invasion resistance pattern appears in the *Ax*/*Bx* phase portrait, which only allows a high *Ax* population ratio to invade and become fixed in both niches.

If *F*2 continues to decrease below this lower limit and approaches zero, the invasion-resistant pattern persists, but fixed points begin to appear in the *Ax*/*Bx* phase portrait. This enables a higher initial *Ax* population ratio to invade, achieve *Ax*/*Bx* fixed-point polymorphism, shift the preexisting *Ax*/*Bx* fixed point toward the bottom right corner of the *Ax*/*Bx* phase portrait, and produce stronger overall RI.

Thus far, our results indicate that the fitness disadvantage posed by increased hybrid incompatibility (*F*2 < *f*) can be offset by the ecological fitness advantage (*Sax* > 1) in an invading *Z* allele. Similarly, the mating-bias incompatibility (*α*1x < 1) can compensate for the fitness disadvantage from hybrid incompatibility (*F*2 < *f*), enabling the *Z* allele to invade and enhance overall RI (Fig 23e). However, as *α*1x decreases further, the *Z* allele‘s strong mating bias could generate an invasion-resistant pattern in the *Ax*/*Bx* phase portrait due to intra-niche sexual selection. Overcoming this resistance pattern would require a larger invading mutant population ratio (*Ax*) or an increased number of mating rounds (*n*). A larger invading population of *Ax* may be achieved if a mutant *Z* allele first increases its population size through evolved ecological adaptations before evolving incompatibilities later. Alternatively, a large initial *Ax* population may arise in parapatry or secondary contact when another population introduces a substantial number of *Z* alleles.

In sum, as hybrid and mating-bias incompatibilities increase in BDM divergence, decreasing values of *F*2 and *α*1x lead to fixed-point polymorphisms in the *Ax*/*Bx* phase portraits and progressively stronger RI. However, there are lower limits of *F*2 and *α*1x below which the *Z* allele cannot invade. At these limits, hybrid loss becomes excessive due to low *F*2, and intra-niche sexual selection becomes too intense due to low *α*1x to permit invasion. As *F*2 and *α*1x approach these thresholds, invasion-resistant patterns emerge in the *Ax*/*Bx* phase portraits, which can only be overcome by a higher initial *Ax* population ratio (Figs 23b and 23d–f). Thus, it may be more feasible for a mutant *Z* allele to invade initially based on its evolved ecological fitness in niche *A* (*Sax* > 1) and only later evolve stronger mating-bias and hybrid incompatibilities after establishing a sizable population in niche *A*. This progression aligns with the expected sequence of BDM barrier evolution: initial ecological adaptation drives genomic divergence, which subsequently gives rise to hybrid and mating-bias incompatibilities as incidental byproducts.

The BDM mechanism of incompatibility predicts that reducing gene flow between two populations of a species facilitates their genomic divergence, either through drift or differential adaptations. Such a process produces mating and hybrid incompatibilities as the two populations diverge. In the incipient stage of sympatric speciation, premating RI between niche populations reduces the homogenizing effect of gene flow and facilitates the emergence of novel, more adaptive ecotypes in their unique niche environments. As these more adaptive ecotypes invade and outcompete the original ecotypes, they bring with them new inter-niche mating-bias and hybrid incompatibilities. This is modeled in Fig 22a and Fig 23e. An invading new niche-*A* ecotype, *Ax*, with antagonistic pleiotropic fitness values, *Sax* and *Sbx*, develops associated new inter-niche mating bias, *α*1x, and hybrid incompatibility, *F*2, as chance byproducts of its widening divergence from the niche-*B* ecotype.

These new incompatibilities, represented by the values of *α*1x and *F*2, are most effective at increasing the overall RI between niches when the fixed point *Ax*/*Bx* is closest to the lower right corner of the phase portrait (where *Ax* = 1 and *Bx* = 0)—in other words, when niche *A* has mostly the *A*_*x*_ ecotype and niche *B* has mostly the less compatible original *B* ecotype. When premating RI is not strong enough and there is significant gene flow, the *Z* allele either fails to invade or becomes fixed in both niche *A* and niche *B* (Fig 23d). When the *Z* allele is fixed in both niches, the values of *α*1x and *F*2 have no effect on the system or on the existing *Ax*/*Bx* fixed point, because there is no differential assortment of the *Z* allele between the niches and because BDM incompatibilities only apply to mating encounters between genotypes with and without the *Z* allele.

In our models, increasing the initial *Ax*/*Bx* premating RI facilitates the invasion of the *Z* allele to reach *Ax*/*Bx* fixed-point polymorphism and produce stronger overall RI. Holding all other parameters constant, a high value of *α* prevents the *Z* allele from invading. As the value of *α* decreases, the system enters an intermediate range where the *Z* allele can invade and become fixed in both niches without altering the existing *Ax*/*Bx* fixed point. Further reduction of *α* toward zero allows both *Ax*/*Bx* and *Ax*/*Bx* fixed-point polymorphisms to emerge in the phase portraits, leading to stronger RI (Fig 23g). The lower the value of *α*, the stronger the resultant RI.

Similar dynamics are observed when varying *f* within the initial *Ax*/*Bx* barrier system. At high values of *f*, the *Z* allele cannot invade; however, as *f* decreases, an intermediate range is reached where the *Z* allele successfully invades and becomes fixed in both niches, again leaving the *Ax*/*Bx* fixed point unchanged. With further reductions of *f* toward zero, *Ax*/*Bx* and *Ax*/*Bx* polymorphisms appear, and RI is strengthened. Here, too, lower values of *f* correlate with stronger resultant RI.

Thus, lower values of *α* and *f*, along with increased antagonistic pleiotropy *Sax* − *Sbx*, facilitate the invasion of the *Z* allele and the emergence of fixed-point polymorphisms and enhanced RI through coupling. These findings support the notion that the BDM barrier mechanism operates most effectively in an environment with preexisting RI and reduced gene flow.

So far, coupling with BDM barriers appears to generate complex invasion and population dynamics, driven by nonlinear interactions among numerous variables. To simplify and clarify our analyses, we can turn our focus to the fundamental processes driving the system’s dynamics as a guide.

The invasion dynamics of the BDM model can be characterized by the interactions of the following key processes: (1) mutations increasing mating bias, *α*-, or decreasing mating bias, *α*+ (2) mutations decreasing hybrid viability *f*- or increasing hybrid viability *f*+ (3) mutations affecting *Sax* and (4) mutations affecting *Sbx*. A *Z* mutation that causes *Sax* > 1 (*Sax*+) can be described as having a conditionally-positive effect for the local species and a conditionally-neutral effect for the foreign species. Conversely, a *Z* mutation that causes *Sbx* < 1 (*Sbx*-) can be described as having a conditionally-negative effect for the foreign species and a conditionally-neutral effect for the local species [69]. A conditionally-beneficial mutation, *Sax*+, increases niche-*A* fitness due to better adaptation to niche *A*. Meanwhile, a conditionally-deleterious mutation, *Sbx*-, can serve as a defense mechanism for niche-*A* species against the invasion of niche-*B* species. Mutations that cause antagonistic pleiotropy result from the coupling of *Sax*+ and *Sbx*-effects. Mutations producing *f*- and *Sbx*-, while capable of creating stronger RI between niche ecotypes, cannot invade on their own because they suffer a fitness disadvantage compared to their same-niche cohorts. Consequently, to invade and increase in number, these negatively-selected mutations must hitchhike or become coupled with other positively-selected mutations that can successfully invade, such as *α*- and *Sax*+, thereby strengthening RI in the system. This coupling can occur through chromosomal inversions [34, 56, 62, 70-72], close physical linkage [35, 61], pleiotropy [73], or the actions of recombination suppressors [62, 74]. These results demonstrate that a positively-selected barrier mechanism can be coupled with a negatively-selected barrier mechanism to produce an overall RI that is stronger than what either uncoupled barrier could achieve independently. The overall invasion fitness of a mutant *Z* allele is determined by the net additive effects of all its associated positively- and negatively-selected barrier processes.

In our BDM model, mating-bias incompatibility (represented by the value of *α*1x) is coupled with hybrid incompatibility (represented by the value of *F*2) in a mutant *Z* allele. The model assumes that the same mating-bias value, *α*1x, applies to both intra-niche and inter-niche encounters of the *Z* allele with native ecotypes. Therefore, even though a small mutant *Ax* population may enjoy a fitness advantage in inter-niche mating with native niche-*B* ecotypes, the same mating-bias incompatibility in intra-niche mating with native niche-*A* cohorts can hinder its invasion. This invasion resistance due to intra-niche sexual selection is mitigated if we assume that the *Z* allele has less mating-bias incompatibility in intra-niche encounters with native niche-*A* cohorts than in inter-niche encounters with native niche-*B* ecotypes. Such an assumption may be justified, as a mutant *Ax* genotype is likely more similar to native niche-*A* ecotypes adapted to the same niche environment than to niche-*B* ecotypes from a different niche environment. Once a mutant *Z* allele invades and establishes a sizable population in niche *A*, this invasion resistance due to intra-niche sexual selection becomes irrelevant, and the *Z* allele is free to evolve increasingly stronger mating-bias and hybrid incompatibilities through the BDM mechanism of divergence.

Lastly, we developed computer applications to simulate the coupling between an initial mating-bias barrier and a two-gene-locus model of the BDM barrier. In this model, two alleles, *Z* and *W*, located at two separate gene loci, exhibit properties characteristic of the BDM mechanism of divergence and incompatibilities and are locally adaptive to niche *A* and niche *B*, respectively. Antagonistic ecological divergence, represented by *Sax* − *Sbx* and *Saw* − *Sbw*, results in increased *Ax*/*Bx* premating RI. Limited gene flow caused by the initial *Ax*/*Bx* premating RI facilitates differential assortment of the *Z* and *W* alleles between the two niches and leads to stronger RI through coupling. The simulation results of the two-gene-locus model closely align with those of the one-gene-locus model, revealing similar invasion and system dynamics as well as resistance to reversal (Figs 24–25).

In advanced stages of sympatric speciation, firmly established BDM barriers—characterized by large antagonistic pleiotropic ecological divergence, strong mating-bias incompatibility, and a high level of hybrid incompatibility—produce robust and nearly complete premating and post-mating RI between niche populations (Fig 25a). Under these conditions, our simulation results confirm that the entrenched BDM reproductive barriers cannot be reversed even if extrinsic disruptive ecological selection is completely removed (i.e., by setting *f* = 0.5). Instead, intrinsic hybrid incompatibility generated by the BDM mechanism is able to take over to ensure continued selection against hybrids and sustain the established RI (Fig 25b). These findings attest to the largely irreversible nature of RI generated through the BDM mechanism of divergence and incompatibilities.

### 9. Among barrier mechanisms, adaptive barriers are the most reversible, chromosomal inversions have limited reversibility, and BDM incompatibilities are nearly irreversible

Our study has demonstrated that after an initial premating RI is established by a two-allele mechanism of mating-bias-allele polymorphism, it is relatively easy to couple other one-allele and two-allele mechanisms of premating barriers to produce ever stronger overall premating RI between niche ecotypes. However, because hybrid loss is the driving force behind all these adaptive premating barriers, reducing or eliminating disruptive ecological selection (by increasing the value of *f*) can cause all the premating barriers to collapse (Fig 8). This outcome reveals the nonpermanent and reversible nature of premating barriers, and we must look elsewhere to find mechanisms that can make the premating barriers irreversible.

An allele *f*_**(−)**_ that increases hybrid incompatibility and decreases hybrid viability, by itself, cannot invade because individuals carrying the allele are at a fitness disadvantage compared to their same-niche cohorts, which do not experience as much reduced offspring return in inter-niche matings. However, when coupled with an allele that has an invasion advantage—e.g., through physical linkage or chromosomal inversion— the *f*_**(−)**_ allele can hitchhike with the advantageous allele and invade.

In our simulations, a mutant Z allele that combines the properties of an *f* _**(−)**_ mutation (low hybrid viability) with a strong mating-bias value, *β*, can invade and couple with an initial *Ax*/*Bx* premating barrier to increase overall RI (Fig 11a). In this scenario, the fitness advantage of a strong mating bias compensates for the fitness disadvantage due to reduced hybrid viability, giving the Z allele a net fitness advantage to invade. After coupling, the lower overall *f* value conveyed by the Z allele can further augment the premating RI due to the stronger mating bias in a synergistic manner, resulting in even stronger overall RI.

After the Z allele is able to invade and rise to near fixation in niche *A*, even if disruptive ecological selection is reduced, the intrinsic hybrid incompatibility conveyed by the *Z* alleles in niche *A* can still prevail to ensure continued high hybrid loss and maintain the established fixed-point polymorphism and premating RI (Fig 11b). However, if disruptive ecological selection is completely removed by setting *f* = 0.5, both the *Ax*/*Bx* and *Ax*/*Bx* fixed points disappear, the system becomes divergent, and no premating RI exists. Thus, coupling an advantageous allele with an *f*_**(−)**_ allele that decreases hybrid viability can secure a certain degree of RI irreversibility, but the irreversibility is not absolute.

Similarly, chromosomal inversions can capture an *f*_**(−)**_ allele along with adaptive ecological alleles. The *f*_**(−)**_ allele can then invade as part of the CI and alter the effective *f* value in the system. The effect of capturing an *f* _**(−)**_ allele that further decreases hybrid viability can be modeled by decreasing the value of *Fk*2 in Fig 12 through Fig 21. The lower value of *Fk*2 can then help to increase the irreversibility of the CI. Compared to adaptive premating barriers, chromosomal inversions can create RI that is more difficult to reverse. After a CI that links favorable ecological alleles with a mating-bias allele has become fixed in a niche, a noninverted mutant that wants to invade must have a higher fitness advantage over the established CIs. In the same niche, a noninverted mutant that has the same ecological fitness as the predominant CIs suffers a fitness disadvantage because of offspring loss in intra-niche heterokaryotype mating (*Fk*1 < 0.5) and because of increased hybrid loss in inter-niche mating. The reason for the latter is that, unlike genotypes with CIs, the favorable allele combinations of the noninverted genotypes can be broken up by recombination in inter-niche mating. Therefore, to gain a selective advantage in intra-niche competition, the noninverted mutant must either evolve a more adaptive ecological genotype than the native CIs or gain a larger supply of noninverted offspring from inter-niche mating.

The noninverted mutant can receive a larger return of noninverted offspring from inter-niche mating if the strength of disruptive ecological selection is reduced, i.e., by increasing *f* so that *f* > *Fk*2 (Fig 20). However, this is difficult to accomplish, because *Fk*2 usually rises with *f* as well. Additionally, when disruptive ecological selection is weakened, the existence of the *“*invasion resistant range,*”* which can be extended by decreasing the value of *α* (Figs 20, 21a, and 21d), provides an additional buffer zone against invasion, even when taking into account the small offspring loss of the CI genotypes from heterokaryotype mating due to single crossovers in inverted regions.

In late-stage sympatric speciation, when RI is already strong, coupling with a fixed CI barrier can make the RI difficult to reverse, even if disruptive ecological selection is completely removed by setting *f* = 0.5 (Figs 21a-21b and 21d-21e). This outcome is possible if *Fk*2 ≤ *Fk*1 and *Fk*2 stays in the *“*invasion resistant range,*”* the *f* − *Fk*2 difference that protects the fixed CI from invasion. To keep *Fk*2 ≤ *Fk*1 when *f* = 0.5 typically requires the CI to capture an *f* ^_^ allele that decreases hybrid viability or for inter-niche heterokaryotype mating to produce less viable hybrid offspring. The inherent hybrid loss (*Fk*2) provided by the CI can then take over to maintain RI when extrinsic disruptive ecological selection is completely removed. In general, decreasing the values of *α* and *Fk*1 increases the invasion resistant range for a fixed CI.

The reason is that for a noninverted mutant to invade, it must overcome the decreased fitness due to intra-niche heterokaryotype mating (as defined by *Fk*1). Thus, a lower value of *Fk*1 makes it more difficult for a small population of noninverted mutant genotypes to invade (Fig 14b). To gain an advantage in inter-niche mating, the noninverted mutant needs to receive more noninverted offspring that are produced by a higher value of *f* (less disruptive ecological selection). However, an increased mating bias (caused by a lower value of *α*) reduces that offspring supply. These findings affirm the expectation that in late-stage sympatric speciation, when premating RI is already strong, a fixed CI linking favorable ecological alleles and mating-bias alleles becomes more resistant to invasion and reversal.

If *f* can be increased to more than 0.5 (while *Fk*2 cannot exceed 0.5 in our model), the fixed CI population can be invaded by a noninverted mutant, resulting in the collapse of the RI. This can occur during a hybrid swamp, when heterosis or the emergence of abundant hybrid niche resources causes a large hybrid population to form that can continuously supply the noninverted mutant with a large quantity of noninverted offspring to overwhelm the existing CI population. Therefore, chromosomal inversions can create a more stable and lasting overall barrier to gene flow when coupled with other barriers in late-stage sympatric speciation, but they are not invincible.

In summary, a CI can create a supergene comprising locally adaptive and linked alleles that is resistant to being broken up or reverted to a noninverted state [35, 39, 40]. Hybrid loss due to heterokaryotype mating within the same niche (caused by *Fk*1 < 0.5) protects predominant CI genotypes from invasion. For a noninverted genotype to successfully invade, it must carry alleles that are even more adaptive than those within the existing CI. However, the advantageous alleles in the invading noninverted genotype, being unlinked, are susceptible to disruption by recombination in the presence of gene flow. In environments with low gene flow, while noninverted genotypes may evolve to become more adaptive than the current CI through the BDM mechanism, their favorable allele combinations remain vulnerable to recombination and the incursion of maladaptive alleles via gene flow. Ultimately, a balance between gene flow and selection can establish a polymorphism that allows a more adaptive noninverted genotype to coexist with the existing CI.

The BDM mechanism of incompatibility appears to provide the most enduring and permanent solution to ensure irreversible RI. The example in Fig 25b shows that in late-stage sympatric speciation, when RI is already very strong, eliminating disruptive ecological selection by setting *f* = 0.5 does little to reverse the RI that has already been established by the BDM mechanism. The BDM incompatibilities develop when genomes of isolated populations diverge over time, either through drift or differential adaptations to local environments, producing mating-bias and hybrid incompatibilities as chance byproducts in the process. Here, we analyze divergence due only to differential local adaptations, as divergence due to drift usually takes a long time, even though the mechanisms causing invasion and reversibility of both are the same.

Because in the BDM process, mating-bias and hybrid incompatibilities are caused by new genetic adaptations to a niche environment (i.e., *Sax* >1 and *Sbx* < 1 in Figs 22-25), they tend to involve fundamental changes in the genome that are very difficult to reverse. It is difficult for diverged genotypes to retrace the same mutational steps that caused the new adaptations and revert back to the original ancestral genotypes. Even when the niche environment is restored to the original state, at best, those diverged genotypes have to take new mutational paths to adapt their phenotypes to the restored ancestral environment through parallel evolution. In the end, the resulting genotypes are still different from the ancestral genotypes. In a sense, well-diverged genomes produced by the BDM mechanism can be treated as those of a new species that has evolved its own unique ecological adaptations as well as mating-bias and hybrid incompatibilities against foreign ecotypes.

For a mutant to invade a niche already occupied by well-adapted and incompatible genotypes formed through the BDM mechanism, it must evolve a higher adaptive fitness than the incumbent population, a requirement that is inherently challenging. After strong BDM incompatibilities have evolved in both niche populations, reducing disruptive ecological selection (by increasing the value of *f*) may provide a small population of ancestral genotypes with a fitness advantage in inter-niche mating by increasing their offspring return ratio. However, to maximize the benefits of this increased inter-niche offspring return requires that the ancestral genotypes have low inter-niche mating bias (so that more inter-niche mating can occur to increase the offspring return). At the same time, intra-niche mating bias between ancestral and BDM-diverged genotypes must also be low to prevent the elimination of ancestral genotypes via intra-niche sexual selection.

These conditions are difficult to meet because the high mating biases between ancestral and diverged genotypes result from fundamental genomic changes, making it unlikely that ancestral genotypes can reduce these incompatibilities without additional mutational changes. Even if this limitation is overcome, ancestral genotypes lacking a definitive niche ecological advantage are vulnerable to competitive niche exclusion by the established divergent genotypes. At most, a polymorphism is achieved when a weakened disruptive sexual selection allows viable hybrid niches to act as a source that continuously supplies ancestral offspring to a niche that acts as a sink to eliminate them.

These challenges attest to the generally irreversible nature of RI produced by the BDM mechanism. In essence, what secures the irreversibility of the BDM process is the high adaptive fitness of the diverged genotypes in their niche environments. The genotypes produced by the BDM process of incompatibility cannot be reversed; they can only be replaced or eliminated by a foreign species of genotypes with higher ecological fitness within the same niche.

Our findings on the reversibility of established RI appear to introduce an additional layer of complexity to the biological species concept [46]. Specifically, the coupling of multiple adaptive premating reproductive barriers (Figs 1–8) can result in strong RI between niche ecotypes under disruptive ecological selection, aligning with the definition of distinct species according to the biological species concept. However, this RI readily collapses when external disruptive selection is removed—for example, when individuals are forced to mate and reproduce under artificial laboratory conditions that may not accurately reflect RI in natural environments. In contrast, RI produced by the BDM mechanism of mutational adaptations, divergence, and incompatibilities is more permanent; species formed through this process remain reproductively isolated even when external disruptive selection is removed. Therefore, merely meeting the biological species definition does not distinguish how stable or reversible the diverging species are. It may be more appropriate to refer to the former as context-dependent or “extrinsically reversible” species, sustained by external selective pressures, and the latter as “intrinsically irreversible” species, characterized by more permanent RI that is independent of environmental conditions.

### 10. Chromosomal inversions can reduce the production of hybrids by multi-locus ecological and mating-bias genotypes

In a separate study, we developed computer applications to investigate how the population dynamics of our models are influenced by multi-locus ecological genotypes and multi-locus mating-bias genotypes [30]. These applications allow us to establish hybrid niche resources that make the existence of viable ecological hybrids possible, given any specified number of ecological gene loci. Similarly, the applications let us specify any arbitrary number of gene loci for mating-bias genotypes to explore their impact on system dynamics.

Consistent with the results of prior studies, the existence of viable hybrids in ecological or sexual selection tends to weaken the strength of premating RI and increase the thresholds for its emergence [28]. In our model (Fig M3), viable ecological hybrids increase the effective values of *f* and decrease the strength of ecological selection. Similarly, multi-locus mating traits produce intermediate mating-bias hybrid genotypes that are ecologically viable in the same niche. These viable mating-bias hybrids tend to decrease sexual selection and hinder the development of premating RI. Therefore, to maximize the chance of premating RI in the presence of disruptive ecological selection, it is desirable to have as many ecological gene loci as possible (so the effective value of *f* is kept small) and to have as few mating-bias gene loci as possible (so fewer mating-bias hybrids are produced, and the effective value of *α* is kept low) [5, 28, 75].

Next, we used the computer applications to investigate the invasion dynamics of chromosomal inversions (CIs) that capture various combinations of mating-bias alleles in a model of multi-locus mating-bias genotypes. The results show that, in convergent systems that already have fixed-point polymorphisms, CIs that capture the predominant high-mating-bias alleles in a niche (e.g., *XXX* in niche *A*) can readily invade and increase the premating RI. Because the captured mating-bias alleles within CIs are resistant to recombination, these alleles cannot be broken up to produce hybrid offspring when CI genotypes mate with noninverted genotypes. As a result, the CIs behave like discrete super-alleles that segregate as a single Mendelian locus, resulting in decreased production of mating-bias hybrids in the system. This reduces the effective number of interacting mating-bias gene loci and promotes the development of premating RI. The more favorable high-mating-bias alleles that a mutant CI can capture (e.g., *XXXX* versus *XX*0*X* in a 4-gene-locus system, where “0” represents the gene locus that is not captured), the greater its fitness advantage to invade, and the stronger the resultant premating RI. When a CI captures unfavorable alleles (e.g., *XXYX* in niche *A*), it suffers a fitness disadvantage that can cause it to be eliminated. Conversely, if an invading CI can capture all the favorable high-mating-bias alleles in a niche (e.g., *XXX*), it can eliminate all intermediate mating-bias hybrids.

Because chromosomal inversion is a late-stage barrier mechanism, it is more likely to capture favorable, locally adaptive alleles when there is already nonrandom, differential assortment of those alleles in different niches [42]. Nonetheless, in a divergent system that does not permit fixed-point polymorphism, a fortuitous CI mutant that captures the prevalent high-mating-bias allele combination within a niche could convert the divergent system into a convergent one. This is more likely to occur when there is already some degree of differential assortment of the mating-bias alleles in the two niches, as might happen, for instance, during secondary contact of two allopatrically diverged populations in a hybrid zone [72, 76, 77].

Our study also investigated the invasion dynamics of CIs that capture locally adaptive ecological alleles in a model with viable ecological hybrids [30]. In the computer applications we developed, the ecological genotypes can have any specified number of gene loci. The population ratios of the various hybrid genotypes are determined by the carrying capacities of their corresponding hybrid niche resources in the system.

In general, the results from models with viable ecological hybrids closely correspond to the results from models without viable ecological hybrids used in this study. The major difference is that in models without viable ecological hybrids, CIs capturing locally adaptive ecological alleles always go to fixation after invasion, whereas CI polymorphism is more likely in models with viable ecological hybrids. This is because viable hybrids can continuously supply noninverted genotypes to the major ecological niches (i.e., niche *A* and niche *B* in the model), even though those noninverted genotypes cannot compete with the fitter, locally adaptive CI genotypes in those niches. A balance between immigration and selection could then lead to a polymorphism of CI genotypes and noninverted genotypes within the niches.

For instance, in our model without viable ecological hybrids, a CI capturing all of the locally adaptive ecological alleles with the predominant mating-bias allele in niche *A* (Fig 15) cannot produce fixed-point polymorphism and premating RI if *NA* is greater than *NB*. This is because, as the CI becomes fixed after invasion, the effective value of *f* becomes 0.5 in the model, and no ecological disruptive selection exists in the system to facilitate convergence. However, if there are viable ecological hybrids in the system that can regenerate the noninverted genotypes in niche *A*, then the CI is no longer fixed in niche *A* but exists in polymorphism with the noninverted genotype. This can restore disruptive ecological selection in the system and lead to premating RI. When invading CIs that capture adaptive ecological and mating-bias alleles exist in polymorphism rather than fixation, their linked mating-bias alleles may also fail to reach fixation, which can result in weaker premating RI.

Even though adaptive CIs that capture favorable ecological and/or mating-bias alleles have a fitness advantage to invade a system with multiple ecological and mating-bias gene loci, CI polymorphisms appear to be the rule rather than the exception when viable hybrids are present. Our simulation results also confirmed that a hybrid swamp of ecological or mating-bias hybrid genotypes, such as in a secondary contact scenario [72, 77], can suppress or eliminate adaptive CI populations even when they carry favorable ecological and mating-bias alleles.

### 11. Habitat preference is another independent barrier mechanism that can produce initial premating RI to jump-start the speciation process

In a sympatric environment, if niche resources exist at distinct sites separated by geographical distance, habitat preference adaptations can be a powerful means to induce initial premating RI between niche ecotypes [78-81]. A classical example is the host-shift speciation observed in maggot apple flies, which diverged from hawthorn tree flies following the introduction of apple trees to North America in the mid-19th century [82, 83]. Under disruptive ecological selection, a mutation that predisposes niche ecotypes to remain in their adaptive habitats and mate with similarly adapted ecotypes gains a fitness advantage to invade and produce premating RI. This fitness advantage arises from reducing inter-niche mating encounters and minimizing the production of maladaptive hybrids.

In a separate study, we investigate the one- and two-allele models of habitat-preference barrier mechanisms [29]. Our findings demonstrate that habitat preference functions as an independent mechanism capable of inducing initial premating RI between geographically separated niche ecotypes under disruptive ecological selection. Similar to our two-allele mating-bias barrier mechanism, once a habitat-preference allele establishes initial premating RI, it can be reinvaded by alleles that confer stronger habitat preferences. Additionally, it may couple with mating-bias barriers or other pre- and post-mating barriers to further strengthen overall RI and complete the speciation process [83].

Habitat preference is considered an early-stage, adaptive, premating barrier mechanism. It is driven by selection pressures caused by maladaptive hybrid loss in disruptive ecological selection. In the absence of hybrid loss, a mutant allele conferring beneficial habitat preference cannot successfully invade. Moreover, if disruptive ecological selection is removed, the premating RI established by habitat-preference barriers readily collapses, similar to other types of adaptive barrier mechanisms.

Two key variables in the habitat-preference models are the offspring return ratio, *f*, and the habitat-preference bias, *θ*. The variable *f* represents the strength of disruptive ecological selection against hybrid genotypes and reflects the degree of hybrid loss. The variable *θ*, which ranges from 0 to 1, quantifies the strength of habitat preference. For instance, a *K* allele with *θ* = 0.3 causes a genotype carrying the allele to remain within its preferred habitat 70% of the time and venture outside (free-roaming) 30% of the time.

Our results revealed that it is much easier for habitat-preference barriers to invade and couple than for the two-allele mating-bias barrier. As long as hybrid loss persists in the system, a mutant allele associated with habitat preference can always invade to reduce hybrid loss to the extent allowed by its habitat-preference bias, *θ*. This holds true for both the one-allele and two-allele models. However, the one-allele model tends to generate stronger habitat-preference RI because it is not affected by the homogenizing effects of gene flow. In the one-allele model, invading mutant alleles either stop at a fixed-point polymorphism when hybrid loss is exhausted or proceed to fixation across both niches. In the two-allele model, unless *θ* = 0, invading mutant alleles tend to stabilize at fixed-point polymorphisms within the niches.

In contrast to the two-allele mating-bias system, which fails to converge when *f* and *α* exceed certain thresholds, habitat-preference barriers do not appear to be subject to such constraints. As long as *f* < 0.5 and *θ* < 1, a habitat-preference mutation can always invade and reduce any remaining hybrid loss to the extent permitted by its habitat-preference bias. This occurs because reduced hybrid offspring loss from inter-niche mating provides a genotype carrying the habitat-preference mutation with a consistent fitness advantage over same-niche genotypes lacking the mutation. Moreover, unlike the two-allele mating-bias system, the habitat-preference mutation does not experience a fitness disadvantage during intra-niche mating with native ecotypes lacking the mutation. Consequently, as long as maladaptive hybrid loss persists in the system, a habitat-preference mutant allele remains under positive selection to invade and proliferate.

By contrast, in the two-allele model of mating-bias barriers, a mutant allele with a higher mating bias than its same-niche counterparts does not always achieve a net fitness advantage to invade, despite reducing hybrid offspring loss from inter-niche mating. This is because its higher mating bias imposes a fitness disadvantage during intra-niche mating with same-niche ecotypes that have lower mating biases, due to the effects of intra-niche sexual selection.

Additionally, our results indicate that establishing an initial habitat-preference barrier can facilitate the subsequent invasion and coupling of a mating-bias barrier, as long as hybrid loss persists [29, 78]. By reducing gene flow, the habitat-preference barrier mitigates the homogenizing effects of recombination and facilitates the differential assortment of mating-bias alleles between niche ecotypes within the mating-bias barrier system. The caveat is that if gene flow is very low, an invasion-resistant pattern may appear in the phase portrait of the mating-bias barrier due to increased intra-niche mating and sexual selection [29]. This invasion resistance pattern may be eliminated by increasing the number of matching rounds (decreasing the assortative mating cost) or reducing the niche carrying capacity (*NA*).

The relative ease and effectiveness of the habitat-preference barrier mechanism make it a strong candidate for initiating premating RI in sympatric speciation. Moreover, this initial barrier can facilitate the subsequent coupling of mating-bias barriers and further strengthen premating RI. Given the frequent criticism that the conditions required for the two-allele mating-bias barrier to establish initial premating RI are too stringent, the “habitat-preference-first, mating-bias-second” pathway offers a compelling alternative to address those concerns.

Even though habitat preference is an adaptive barrier mechanism, it is more difficult to reverse than other adaptive premating barriers when disruptive ecological selection is relaxed. This is because, as long as there is maladaptive hybrid loss remaining, a habitat preference barrier is always under positive selection pressure to eliminate the hybrid loss, to the extent allowable by its associated habitat bias, *θ*.

For instance, in a one-allele mechanism, an invading habitat-preference mutant allele *K* either goes to fixation in both niches after it has exhausted its capability to eliminate maladaptive hybrids, or it will stop at a fixed-point polymorphism in the *Ak*/*Bk* phase portrait after it has eliminated all the inter-niche hybrid loss and there is no more driving force for it to progress [29]. In the latter case, altering the disruptive ecological selection will cause compensatory changes in the *Ak*/*Bk* ratio at the fixed point to eliminate the resultant increase in hybrid loss.

In a two-allele mechanism, fixed-point polymorphism typically occurs in the *Ak*/*Bk* phase portrait, driven by selection pressure arising from hybrid loss [29]. Stronger habitat preference tends to move the *Ak*/*Bk* fixed point toward the bottom right corner of the phase portrait where *Ak* = 1 and *Bk* = 0. Similar to the one-allele mechanism, the *Ak*/*Bk* ratios at the fixed point automatically adjust to accommodate varying levels of maladaptive hybrids produced by changes in the intensity of disruptive ecological selection.

Once habitat-preference mutant alleles successfully invade and reach significant population frequencies within the niches, removing disruptive ecological selection by setting *f* = 0.5 does not eliminate the habitat-preference alleles or the premating RI they create. When *f* = 0.5, genotypes carrying habitat-preference alleles no longer have a fitness advantage in inter-niche mating over their same-niche ecotypes lacking these alleles; however, they do not suffer any fitness disadvantage either. Consequently, they have the same fitness as other genotypes within the same niche, which allows them to persist and sustain premating isolation between niche populations.

In real-world scenarios, however, habitat preference likely incurs costs when disruptive ecological selection and maladaptive hybrid loss no longer exist. Such costs could arise from increased local competition or when the advantages of dispersal to explore new resources outweigh the benefits of remaining confined to the same niche. These costs could impose a fitness disadvantage on habitat-preference genotypes compared to same-niche genotypes that are free to disperse, leading to the elimination of habitat-preference alleles from the population.

In sum, even though the invasion of habitat-preference alleles requires disruptive ecological selection and maladaptive hybrid loss, once these alleles become widespread and establish premating RI, their persistence can only be reversed by costs associated with habitat preference when disruptive ecological selection is removed.

Theoretically, distinct niche resources do not need to be geographically separated for the habitat preference barrier mechanism to work. Consider a lake where two types of algae bloom at different times. In this environment, a sympatric species may evolve two distinct ecotypes, each adapted to feed on one type of algae. Disruptive ecological selection acting on temporal resource use then favors ecotypes that synchronize their feeding and mating cycles with the algae bloom periods. A habitat preference mutation that increases the likelihood of ecotypes remaining in their adaptive niches—by aligning their cycles with the algae blooms—confers a fitness advantage by reducing maladaptive hybrid loss. This can create a magic trait, where ecological adaptation simultaneously drives divergence in mating encounters [84-86]. Thus, the habitat preference barrier mechanism can produce temporal RI between the niche ecotypes that are caused by temporal separation of their resources rather than by geographical separation [81].

Therefore, the habitat preference barrier mechanism can operate across a wide range of premating reproductive barriers in sympatric populations, including spatial, temporal, behavioral, and mechanical isolation, by reducing opportunities for mating encounters. This general mechanism should be applicable as long as separate habitats exist along any of these dimensions. In contrast to the mating-bias barrier mechanism, which establishes premating RI through mating-trait incompatibilities during mating encounters, the habitat preference barrier mechanism achieves isolation by limiting the likelihood of such encounters. In effect, it functions analogously to a geographical barrier by limiting contact between the two niche populations.

### 12. Early-stage adaptive barriers can couple with late-stage adaptive and nonadaptive barriers in a positive feedback loop to complete the speciation process

Our study investigates how an initial premating reproductive barrier may recruit additional adaptive and nonadaptive reproductive barriers to produce sympatric speciation. We can classify the barrier mechanisms in sympatric speciation as early-stage or late-stage and as adaptive or nonadaptive (see Table I). Fig 26 shows how the early-stage mechanisms and the late-stage mechanisms can interact to complete the speciation process. The early-stage mechanisms produce premating RI, and they are adaptive in nature, driven by the selection pressures to minimize hybrid loss in disruptive ecological selection. They include the one- and two-allele models of mating-bias assortment as well as the one- and two-allele models of habitat preference [29]. An additional one-allele model of mating-bias barriers, not included in our study, involves a mutant allele that causes individuals to prefer mating with ecotypes similar to their own [87, 88]. As RI becomes strong and maladaptive hybrid loss is reduced, the selection pressure driving adaptive barriers to invade and to couple also diminishes. However, as long as there is ample hybrid loss, these early-stage barrier mechanisms can readily couple together to produce progressively stronger premating RI (Figs 1-8).

All early-stage premating barrier mechanisms can establish an initial barrier in sympatric speciation and reduce gene flow. Our simulations show that a prior barrier, by reducing the homogenizing effect of gene flow, facilitates the coupling of a subsequent two-allele mating-bias barrier. Such facilitation is seen in our study of habitat-preference barriers, where an initial habitat-preference barrier facilitates the coupling of a subsequent two-allele model of mating-bias barrier [29, 78]. Similarly, when coupling two mating-bias barriers, it is easier for a second barrier to couple with a convergent first barrier that has already established premating RI than with a divergent first barrier that has not reduced gene flow. Nonetheless, a strong first barrier that creates very low gene flow can cause an invasion-resistant pattern to appear in the phase portrait of a second mating-bias barrier. This resistance may be eliminated by increasing the number of mating rounds, *n*, (Fig 3a, 3b) or by reducing the niche carrying capacity, e.g., *NA*.

Early-stage barrier mechanisms produce prezygotic, premating RI, but they do not affect post-zygotic, post-mating RI. In contrast, late-stage mechanisms—such as mutations that reduce hybrid viability, chromosomal inversions, and the BDM mechanism—can generate post-zygotic barriers and post-mating RI by increasing hybrid incompatibility and reducing hybrid viability.

The strength of premating RI for mating-bias barriers depends on both ecological selection (i.e., the value of *f*) and sexual selection (i.e., the values of *α* and *n*). Consequently, any coupled mechanism that affects hybrid viability (i.e., by altering the effective value of *f*) will also affect the degree of premating RI in other coupled adaptive premating barriers. For instance, our modeling has demonstrated that a mutant allele *f*_**(−)**_ that decreases hybrid viability and reduces the effective value of *f*, on its own, cannot invade because it suffers greater offspring loss in inter-niche mating compared to its same-niche cohorts. However, if a chromosomal inversion can capture such a negatively-selected *f*_**(−)**_ ) allele with other positively-selected, advantageous ecological or mating-bias alleles, the *f*_**(−)**_ allele can invade as a part of the CI and establish a lower effective *f* value (stronger effective ecological selection) in the system (Fig 11a). The increased hybrid loss created by the invasion of the *f*_**(−)**_ allele can then act as positive feedback to synergistically increase the premating RI in other coupled premating barriers. Once the invading *f*_**(−)**_ allele becomes prevalent, it can also protect, to a certain extent, the established premating RI from being reversed by a reduction in environmental disruptive ecological selection (Fig 11b).

Chromosomal inversion is considered a late-stage mechanism in sympatric speciation. This is because a CI mutant is more likely to capture locally-advantageous ecological and mating-bias alleles if other barriers have caused differential assortment of these alleles across different niches [42]. A CI can protect linked alleles within its inverted region from being broken up by recombination during heterokaryotype mating. Consequently, a CI can combine many adaptive small-effect alleles to create a single supergene of large effect. CIs offer little advantage when traits are already controlled by tightly linked or large-effect genes [35, 89]. Prior research analyses have confirmed that advantageous CIs are most likely to emerge and invade when highly polygenic traits exist without large-effect genes and when there is high gene flow between niches [35].

In our model, CIs are considered an adaptive barrier mechanism. The invasion of CIs that capture locally adaptive alleles is driven by selection pressures created by ecological hybrid loss. Therefore, the invasion of a locally adaptive CI is most likely after early-stage barriers have created incomplete RI that permits moderate gene flow but before strong RI completely halts gene flow. If RI is weak, it is difficult for CIs to capture locally adaptive alleles. However, if RI is strong, it is difficult for CIs to invade [42].

Because CIs can act like supergenes, when they capture high-mating-bias alleles in a multi-locus genotype system, they can gain a fitness advantage to invade and reduce the production of mating-bias hybrids that impede system convergence. The end result is that the parametric thresholds for system convergence are lowered, and stronger premating RI is achieved through the invasion of high-mating-bias CI supergenes [30].

Similarly, when CIs capture locally adaptive ecological alleles, they gain a selective advantage over their noninverted cohorts in the same niche, as the ensembles of adaptive alleles in the noninverted genotypes can be broken up by recombination. Our model results show that such adaptive CIs can invade even if they capture only a partial set of locally adaptive alleles (Figs 13a and 13b).

However, CIs that capture locally adaptive ecological alleles reduce the production of ecological hybrids that are subject to disruptive ecological selection. As a result, their invasion decreases hybrid loss and weakens ecological selection. This can reduce premating RI and create “negative coupling” [32], which decreases overall RI rather than enhancing it (Figs 13a and 13b). Consequently, while gene flow within the CIs is reduced, gene flow in the rest of the noninverted genome is increased (see the “interchromosomal effect” in [44]). In the extreme case where an invading CI captures all the locally adaptive alleles, no hybrid offspring are produced, the value of *f* in the Fig M3 model increases to 0.5, and the system becomes divergent (Fig 12a). Although there is 100% RI for the alleles within the CI, the rest of the genome outside the CI experiences no RI.

On the other hand, if an invading CI can capture locally adaptive ecological alleles along with the prevalent mating-bias allele in the niche, it can extend premating RI to the entire genome (Figs 16a, 16b, 18a, and 18b), consistent with findings from prior studies [60-62]. For this to occur effectively, the CI should only capture a partial number of the adaptive ecological alleles, so hybrid loss can be restored (*f* <0.5). The CI-linked mating-bias allele can then hitchhike with the positively-selected, locally-adaptive CI to invade and create premating RI between entire genomes of the two niches. Similarly, when a CI captures locally adaptive alleles with a mutation *f*_**(−)**_ that reduces hybrid viability, the *f*_**(−)**_ mutant can hitchhike with the CI and increase ecological hybrid loss (decrease *f*), resulting in stronger premating RI.

A CI that captures locally adaptive ecological alleles along with mating-bias alleles, in effect, creates a “magic trait” that links ecological traits with sexual-discrimination traits [84-86]. According to our results, CIs that capture mating-bias alleles with only a partial number of the locally adaptive ecological alleles (i.e., “pseudomagic traits” [63, 64]) tend to facilitate the development of premating RI better than CIs that capture all of the locally adaptive alleles (i.e., bona fide “magic traits”). This seems to support the findings of a prior study that greater linkage association of mating-bias alleles with locally adaptive alleles may not always lead to stronger RI [63].

Because a CI protects its inverted region from recombination, it can be treated as an isolated enclave that is free to evolve and adapt separately from the rest of the genome. Compared to its noninverted cohorts, a CI that has already captured locally adaptive alleles has the advantage of building upon its already favorable allele combination to further diverge and evolve more adaptive CI variants without the hindrance of recombination (the “capture and gain” mechanism [35, 42, 90-92]).

Adaptive mutations arising within a chromosomal inversion can gain a greater selective advantage than if they were to occur in noninverted genotypes. This advantage is due to the mutations being linked with other beneficial alleles, and the CI prevents them from being separated through recombination. In contrast, the same adaptive mutations arising in noninverted genotypes are liable to be broken up by recombination in inter-niche mating. Consequently, when a CI coexists in polymorphism with noninverted genotypes within a niche, it can evolve and accumulate advantageous adaptations faster than the noninverted genotypes [41]. This gives the CI a fitness advantage to invade and proliferate.

Genes within the inverted region of a CI are shielded from recombination during inter-niche mating. As a result, the CI can function as “an asexual gene unit” [37] and becomes a hotspot for adaptation and divergence. The reduction in gene flow creates RI for the captured genes, which facilitates their adaptive divergence and the evolution of incompatibilities through the BDM mechanism [92].

In effect, CIs can be conceptualized as adaptive alleles of a newly formed supergene that segregates as a polymorphic single Mendelian locus [39, 40, 55], while the noninverted genotypes represent less adaptive allelic variants of this supergene. Through the BDM mechanism, the alleles associated with the CIs can subsequently evolve to produce variants that are more ecologically adaptive and possess greater hybrid and mating-bias incompatibilities [41]. Polymorphisms of these CI super-alleles may be maintained by mechanisms similar to those that preserve polymorphisms of single-locus alleles [54]. Such polymorphisms can expand the range and standing variations of adaptive alleles within the population, enhancing the species’ adaptive fitness across varying environmental conditions [52, 54, 93-96].

In systems with multi-locus mating-bias genotypes, CIs capturing locally prevalent mating-bias alleles evolve stronger incompatibilities faster than noninverted genotypes due to suppressed recombination. This adaptive process is accelerated when the system is under selection pressure to evolve greater mating biases to further reduce gene flow and hybrid loss. As a result, major reproductive barrier genes may become concentrated within the CIs.

When locally adaptive CIs evolve mating-bias incompatibilities through the BDM mechanism, they effectively function like CIs that capture both locally adaptive ecological alleles and mating-bias alleles. This is particularly likely when ecological traits also act as mating-bias traits (i.e., a magic trait [84-86]). This process provides a plausible mechanism for linking adaptive ecological traits to reproductive barrier traits within CIs.

Once a CI rises to near fixation within a niche, driven by the fitness advantage conferred by its captured locally adaptive ecological alleles, it effectively establishes a protected genomic island of inter-niche (post-zygotic) RI for genes within its inverted region. This restricted gene flow environment within the inversion facilitates the evolution of the BDM mechanism and coupling with other late-stage barrier mechanisms for the captured genes [90-92]. Notably, even if disruptive ecological selection and adaptive premating RI are lost, the inverted regions remain reproductively isolated due to suppressed recombination during inter-niche heterokaryotype matings. Furthermore, if the BDM mechanism enables the CI to evolve ecological and mating-bias incompatibilities alongside its ecological adaptations, these incompatibilities can sustain RI across entire genomes between niche populations, even without disruptive ecological selection, thereby rendering the established RI irreversible [41].

Over time, a more adaptive CI is likely to replace a less adaptive CI [53, 55]. One potential mechanism to aid in this process is through purifying selection [39, 66]. Suppressed recombination often accelerates the accumulation of deleterious mutations and reduces the efficacy of purifying selection. When CIs exist in polymorphism, the more locally adaptive CIs are more likely to gain in frequency than the less adaptive CIs. As the effective population sizes of these more adaptive CIs grow, recombination in their inverted regions also increases, which helps purge deleterious mutations that tend to accumulate over time [5]. Meanwhile, less populous and less adaptive CIs are eliminated by their mutation loads [39, 66]. This creates a positive feedback loop that allows more adaptive CI variants to invade and replace less adaptive CI populations [97].

The accumulation of mutational loads within CIs has also been proposed as a mechanism to explain the maintenance of CI polymorphism [55, 66]. Suppressed recombination in CIs causes the accumulation of deleterious mutations. In diploid organisms, these deleterious mutations increase the likelihood of expressing harmful recessive traits in offspring that are homozygous for the CIs. In contrast, heterozygous offspring resulting from matings between CIs and noninverted genotypes are relatively unaffected and exhibit higher fitness. This dynamic creates negative frequency-dependent selection, where the deleterious effects of homozygosity prevent the CIs from rising to fixation in the population [55, 98].

When CIs are minority populations in a polymorphism, they evolve as “asexual gene units” within the genome [37], accompanied by the advantages and disadvantages of asexual reproduction. In contrast, their noninverted counterparts are subject to recombination and evolve under the dynamics of sexual reproduction. In the noninverted colinear regions, recombination facilitates adaptive evolution but breaks up associations among coadapted genes and hinders divergence and speciation (Felsenstein’s dilemma [27, 54]). In the asexual mode, CIs can more rapidly accumulate adaptive mutations, which enables them to outcompete less adaptive variants and increase in frequency. However, without the purifying selection inherent to sexual reproduction, CIs are more vulnerable to deleterious mutations, which can accelerate their elimination. Consequently, the proportion of CIs within a polymorphism determines the relative contributions of these asexual and sexual modes of evolution to future adaptations.

In our models, CI polymorphisms of inverted and noninverted ecological genotypes in niche *A* or niche *B* are maintained by the presence of viable hybrids. In ecological selection, the presence of hybrid niche resources can sustain viable hybrid populations that continuously supply noninverted ecological genotypes to the two major parental *A* and *B* niches. In sexual selection, when mating traits are polygenic, all the mating-bias hybrids existing in the same niche are ecologically viable. If mating bias is not complete (i.e., *α* > 0), CI polymorphisms of mating-bias alleles in the niches tend to be the rule rather than the exception [30]. However, when mating bias is complete (i.e., *α* = 0), there is no selection pressure for a CI capturing mating-bias alleles to invade. CIs of mating-bias alleles create super-alleles that can reduce the production of mating-bias hybrid offspring. If a CI is able to capture all the extreme mating bias alleles (e.g., *XXX* or *YYY*), then no mating-bias hybrids are produced at the fixed points. If a CI only captures a portion of the extreme mating-bias alleles (e.g., *XX*0 or *Y*0*Y*, where *“*0*”* indicates the gene locus that is not captured), then mating-bias hybrids can exist at the fixed points [30].

When disruptive ecological selection is reduced or when the maximum mating bias between extreme mating-bias genotypes is weak, a hybrid swamp can significantly lower the CI ratio in a polymorphism. This decrease can make the CI vulnerable to elimination through genetic drift.

Our simulation results indicate that when a CI captures locally adaptive ecological alleles along with a favorable mating-bias allele, it can lead to genome-wide premating RI. Strong RI can protect the successful invasion of such a CI from being reversed (Figs 21d and 21e), particularly when inter-niche heterokaryotype mating produces less viable hybrid offspring (e.g., when *F*2*k* in Fig 21d is low). This can occur if the CI captures an *f*_**(−)**_ allele that decreases the ecological viability of hybrids, or if it results from evolved BDM hybrid incompatibility within the CI. Consequently, when environmental disruptive ecological selection relaxes, the CI’s intrinsic selection against hybrids can take over and sustain the existing RI.

In summary, CIs are a late-stage barrier mechanism that can combine locally adaptive ecological alleles with favorable mating-bias alleles and with alleles that reduce hybrid viability to strengthen premating and post-mating RI. It is less easily reversed, especially in an environment of strong RI, when compared to other adaptive barrier mechanisms. It is fast and adaptive but still subject to reversal. It is an important late-stage mechanism that can buy time and prepare the way for BDM incompatibilities to develop and complete the speciation process.

Because all the adaptive barriers, arising from shifts in population dynamics, are easily reversed when environmental disruptive ecological disruption is removed, it appears that new mutational changes in BDM incompatibilities and chromosomal rearrangements offer the best prospects for achieving permanent and irreversible RI. Among the various types of chromosomal rearrangements, those that do not result in the loss of essential genes—such as chromosomal inversions, duplications, and translocations—are the most promising candidates [99].

In our study, BDM incompatibilities appear to be the only mechanism capable of ensuring lasting and permanent reproductive barriers between sympatric species. The primary issue with the BDM mechanism is that it requires significant time to develop. BDM incompatibilities arise either through drift or differential adaptation to distinct environments. We have ignored the drift mechanism in our study, because it takes too long. In contrast, differential adaptations can occur more rapidly and are facilitated in sympatric speciation by disruptive ecological selection, which forces incipient sister species to adapt to very different niche environments.

The BDM process is a late-stage mechanism. Only in an environment of low gene flow can different niche ecotypes be free to diverge and adapt to their different niche environments without interference from the homogenizing effects of recombination. Subsequently, this divergence leads to the evolution of ecological hybrid incompatibilities and mating-bias incompatibilities as incidental byproducts. The process works because the low gene flow environment created by premating RI in sympatric speciation is indistinguishable from that created by geographical barriers in allopatric speciation [28].

In a low-gene-flow environment, as mutant ecotypes in a niche evolve improved ecological adaptations to a particular niche environment, they gain a fitness advantage to invade and replace the native same-niche ecotypes, creating, as chance byproducts, increased hybrid and mating incompatibilities via the BDM mechanism. These incompatibilities reduce inter-niche gene flow and strengthen RI. Antagonistic pleiotropy in ecological fitness across niche environments—i.e., *Sax* > 1 and *Sbx* < 1 in Figs 22 and 23—ensures that the evolved ecological adaptations and their associated incompatibilities remain localized within the niche where they originated. This prevents their introgression into the opposite niche, which can weaken the BDM incompatibilities between the niche populations (e.g., there are no BDM incompatibilities between the *Ax* and *Bx* genotypes in Fig 23d).

In the BDM barrier mechanism, as sympatric sister species evolve locally adaptive mutations and diverge ecologically, they also develop mutual hybrid and mating-bias incompatibilities as a result of their genomic divergences. These locally adaptive mutation steps, along with the resulting incompatibilities, are very difficult to reverse, as retracing and undoing all the mutation steps is nearly impossible. Furthermore, the new mating-bias incompatibility is tied to the new ecological hybrid incompatibility through the evolved genotypes, and it is difficult to reverse one without affecting the other. As a result, they can function like “magic traits” [84-86].

In effect, the two niche ecotypes have evolved into distinct species. Even if disruptive ecological selection is removed and the environment reverts to its original sympatric state, the evolved species can only readapt to their ancestral niches through parallel evolution. They will not possess the same genomic makeup as their ancestors. These sympatric sister species are also immune to the effects of hybrid swamping by ancestral genotypes. The only way they could be displaced from their adapted niches is through replacement by other fitter, more adaptive species, in accordance with the competitive exclusion principle.

Our study has revealed a two-stage process of sympatric speciation (see Fig 26). The early-stage mechanisms are adaptive and premating in nature. They include the emergence and coupling of the one- and two-allele models of mating-bias barriers, as well as the one- and two-allele models of habitat-preference barriers. These mechanisms establish the initial reproductive barriers in sympatric speciation, reduce gene flow between niche ecotypes, and facilitate the coupling of late-stage barriers.

The late-stage mechanisms operate most effectively in an environment of reduced gene flow. They can be adaptive or nonadaptive, and they can create premating or post-mating RI. The late-stage adaptive mechanisms include chromosomal inversions that can couple locally adaptive ecological alleles with favorable mating-bias alleles and with alleles that reduce hybrid viability to strengthen and secure the overall RI. The late-stage nonadaptive mechanisms involve the development of both premating and post-mating BDM incompatibilities. Although these incompatibilities take longer to evolve, they tend to be more irreversible and permanent. Both adaptive and nonadaptive late-stage barrier mechanisms can develop their own intrinsic post-zygotic hybrid incompatibilities.

Because adaptive premating RI is affected by hybrid loss, intrinsic hybrid incompatibilities produced by the late-stage mechanisms can lead to a further increase in early-stage premating RI. When environmental disruptive ecological selection is reduced, these intrinsic hybrid incompatibilities can take over to ensure continued hybrid loss and maintain the existing RI. Over time, as BDM mating-bias incompatibilities mature and restrict inter-niche gene flow, intra-niche sexual selection drives the predominant mating-bias alleles in each niche to fixation (e.g., *X* allele in niche *A* and *Y* allele in niche *B*).

As illustrated in Fig 26, premating barriers facilitate the development of post-mating barriers, which, in turn, strengthen and secure existing premating RI, leading to even stronger post-mating RI. Such interactions create a positive feedback loop, in which early-stage and late-stage mechanisms interact synergistically to produce progressively stronger and more irreversible RI over time, ultimately completing the speciation process.

### 13. Chromosomal inversions can weaken preexisting RI via negative coupling while facilitating linkage between locally adaptive ecological alleles and incompatibility alleles

The positive feedback loop in Fig 26 assumes that late-stage CIs can capture locally adaptive ecological alleles along with the locally prevalent mating-bias allele or a mutation reducing hybrid viability, *f*_**(−)**_, to further strengthen and secure the early-stage premating barriers. When such a CI captures the locally prevalent mating-bias allele, it extends premating RI across the entire genome, and when the CI captures a mutation that reduces hybrid viability, it creates post-zygotic RI. However, if the CIs capture only locally adaptive ecological alleles, they can produce negative coupling that weakens the preexisting premating RI by reducing hybrid loss and the effective strength of disruptive ecological selection. This can potentially disrupt or break down the positive feedback loop.

We can examine factors that may make such negative feedback more or less likely. As early-stage premating RI increases, late-stage CIs become more likely to capture locally adaptive ecological alleles, thus acquiring a fitness advantage that allows them to invade. The resulting negative coupling reduces recombination within adaptive CIs but increases gene flow across the rest of the genome by weakening preexisting premating RI (“interchromosomal effect” [44]). Subsequently, this reduced premating RI makes the capture of locally adaptive ecological alleles by additional CIs less likely. Nevertheless, existing CIs that have successfully invaded become difficult to eliminate due to their acquired fitness advantage in their adaptive niches.

The properties of locally adaptive ecological alleles and their distribution across the genome also appear to play a role. Large-effect alleles offer less fitness advantage for CIs to capture. Adaptive alleles are easier to capture when they are clustered together in close physical proximity on the same chromosomes than when they are scattered throughout the genome. Many small-effect, locally adaptive alleles that are scattered across the genome and located on different chromosomes are difficult to capture by a single or a few CIs, and when they are captured, the effect of negative coupling they exert tends to be limited.

Nonetheless, in late-stage sympatric speciation, when existing premating RI has created nonrandom assortment of locally adaptive ecological alleles across different niches, the probability of CIs capturing these coadapted alleles greatly increases. These adaptive CIs invariably acquire a fitness advantage that enables them to invade and proliferate within the genomes of niche ecotypes, driven by their ability to reduce inter-niche hybrid offspring loss compared to their same-niche, noninverted counterparts. This could explain the observation that in closely related sympatric sister species, most of the locally adaptive genes (represented by regions of high *FST*) are located within CIs. Collectively, these locally adaptive CIs reduce the effective number of ecological loci mediating disruptive ecological selection and maladaptive hybrid loss, resulting in weaker premating RI.

If mating-bias incompatibilities arise within CIs that capture only locally adaptive ecological alleles, they can restore premating RI across the entire genome. A mating-bias mutation arising within a CI that captures locally adaptive ecological alleles provides the CI with a fitness advantage over its same-niche, noninverted counterparts due to decreased inter-niche mating and reduced hybrid offspring loss.

The manner in which a mating-bias mutation arises in a locally adaptive CI appears to influence its invasion dynamics. When a CI captures locally adaptive ecological alleles along with the locally prevalent mating-bias allele at its inception, it immediately gains an augmented fitness advantage to invade. In this case, the fitness advantage of having the locally favored mating-bias allele is augmented by the fitness advantage of having locally coadapted ecological alleles combined within the same CI.

A mating-bias mutation may arise in a locally adaptive CI that has already successfully invaded a niche and reached a stable polymorphism with the noninverted genotypes. Because CIs act as protected enclaves of evolution that are isolated from the rest of the genome, such a mutation can arise de novo within a CI due to further adaption and divergence of the genes in its inverted region or through the BDM mechanism. Because genes within their inverted regions are shielded from recombination, CIs can rapidly build upon the fitness advantages of their captured, locally coadapted ecological alleles and accelerate their adaptive evolution relative to their same-niche, noninverted counterparts. Alternatively, the mating-bias mutation may come from outside of the CI, such as through the actions of transposons, pleiotropy, or other recombination suppressors.

When existing in a polymorphism with non-inverted genotypes, a locally adaptive CI with mating-bias incompatibility retains augmented ecological and mating-bias fitness advantages over the native, noninverted genotypes in the same niche. However, its fitness advantage over other adaptive CIs without mating-bias properties in the polymorphism is solely due to the benefit of possessing the mating bias. Thus, the frequency of the locally adaptive CI within the polymorphism determines the overall invasion fitness of a new CI mutant that acquires mating-bias incompatibility.

Because chromosomal inversion is an adaptive barrier mechanism, its invasion is driven by selection pressure arising from hybrid loss. When premating RI reduces hybrid loss to a sufficiently low level, it can inhibit the invasion of additional adaptive CIs. Reducing inter-niche gene flow decreases the fitness advantage that additional CIs capturing locally adaptive alleles can gain from inter-niche mating. Meanwhile, as inter-niche mating becomes rare, hybrid loss from intra-niche heterokaryotype mating (represented by *Fk*1 < 0.5) becomes more important in hindering the invasion of new adaptive CIs. Our simulation results have shown that when RI is very strong, it becomes very difficult for additional highly adaptive CI mutants to invade, even with an associated *f* value of 0.5, making the established CIs irreversible (Fig 19-21).

In sum, locally adaptive CIs facilitate the emergence of mating-bias incompatibilities in their inverted regions. By capturing locally advantageous ecological and mating-bias alleles, a locally adaptive CI with favorable mating-bias incompatibility gains an augmented fitness advantage from their combination. This fitness advantage allows it to invade and strengthen preexisting early-stage premating barriers (Fig 26) and, in certain cases, convert a divergent mating-bias barrier into a convergent one to facilitate its coupling (Fig 19).

In contrast to mating-bias incompatibilities, the emergence of post-zygotic hybrid incompatibility in the presence of gene flow remains difficult to explain. This challenge arises because a mutation reducing hybrid viability, *f*_**(−)**_, incurs greater hybrid offspring loss in inter-niche matings than its same-niche counterparts, rendering it underdominant and unable to invade on its own. To spread within a population, it must hitchhike with other processes that provide a sufficient fitness advantage to offset its inherent disadvantage. Consequently, a mutation reducing hybrid viability is likely to arise only as an unavoidable or unintended byproduct of other processes that generate a net fitness benefit, such as the BDM mechanism of adaptation and divergence.

When hybrid incompatibility arises in a locally adaptive CI, its fitness cost relative to same-niche, non-inverted genotypes is offset by the advantage of its linked ecological alleles. However, if the CI already exists in a stable polymorphism, a mutant CI with newly acquired hybrid incompatibility faces a greater disadvantage against other CIs than against non-inverted genotypes. This is because all CIs share the same adaptive ecological alleles, so the mutant CI gains no additional ecological advantage. Consequently, the frequency of the CI within the polymorphism determines the overall invasion fitness disadvantage of a new CI mutant that acquires hybrid incompatibility.

For a locally adaptive CI to capture an *f*_**(−)**_ mutation at its inception, the mutation must persist in the genome despite the homogenizing effects of gene flow. Several natural mechanisms can enable mildly deleterious or underdominant mutations to escape purging by purifying selection. These include pleiotropy, balancing selection, genetic drift [100], recessive genetic load [101], homozygous advantage [86], meiotic drive [102], stepwise mutation [103], ecological epistaxis [104], or linkage with beneficial alleles [34, 54, 86, 92, 101]. For instance, in small niche populations, genetic drift may allow an underdominant *f*_**(−)**_ mutation to rise in frequency—or even fix—despite its fitness cost. Once established, such a mutation could further limit gene flow between the small niche and larger niche populations, thereby reinforcing RI. Alternatively, if hybrid incompatibilities inevitably arise as byproducts of divergent ecological adaptation [105], baseline levels of postzygotic incompatibility may be expected among ecotypes within sympatric populations. In such cases, multiple locally adaptive CIs may coexist in a stable polymorphism maintained by balancing selection, with their relative frequencies fluctuating in response to environmental variation [52, 106]. Because these CIs encapsulate distinct “adaptive cassettes,” hybrid incompatibilities could naturally emerge among them as a consequence of their divergent adaptations [107].

Because locally adaptive CIs promote the accumulation of novel ecological and mating-bias adaptations within their inverted regions, they create fertile ground for the evolution of hybrid incompatibilities through continued divergence [33, 54, 92, 108, 109]. If an *f*_**(−)**_ mutation becomes embedded within a CI—whether through de novo mutation, introgression, or the BDM mechanism—its success within an existing CI polymorphism depends on the relative frequencies of CIs in the niche population. In a polymorphic context where multiple CIs already confer similar ecological advantages, a new CI variant carrying hybrid incompatibility gains no inherent fitness advantage unless it also evolves stronger adaptive traits. Only when hybrid incompatibility arises as an unavoidable byproduct of further beneficial ecological or mating-bias adaptations—and the resulting CI still achieves a net fitness gain over its competitors despite the associated cost—can it successfully invade and persist within the polymorphism.

Our study uses the “gene flow during speciation” model to explain the relative increase in divergence and hybrid incompatibility alleles within CIs compared to colinear regions [108]. According to this model, incompatible alleles that are captured or evolve alongside locally adaptive alleles in CIs persist because the fitness advantage of linked adaptive alleles compensates for the fitness cost of the incompatibilities. In contrast, colinear regions of diverging genomes remain subject to homogenization by gene flow between sympatric populations, which purges incompatible alleles outside of CIs. However, alternative theoretical models also exist to explain hybrid incompatibility within locally adaptive CIs in sympatric populations. Most of these alternative theories suggest that hybrid incompatibilities initially evolved during periods of allopatric isolation and were later maintained following secondary contact.

In the “gene flow after speciation” model [108], allopatrically isolated populations evolve hybrid incompatibility genes uniformly across both inverted and colinear regions of the genome. Upon secondary contact, incompatible alleles linked with other beneficial alleles in CIs tend to be preserved when sister species re-hybridize, while incompatible alleles in unlinked colinear regions are eliminated because they produce unfit hybrid progeny [33, 59, 108]. The end result is that after secondary contact and gene flow, incompatibility loci exist only within CIs.

Another alternative model assumes the presence of CI polymorphism in the ancestor of two species. Through incomplete lineage sorting, different types of CIs are inherited by the two descendant species, generating immediate divergence between their populations. This divergence then increases the likelihood of evolving further divergence and hybrid incompatibilities in their inverted regions compared to colinear regions. This model does not require gene flow to homogenize colinear regions, as they are inherited from the same ancestral species [108].

These alternative models highlight the critical role of gene flow in shaping both the emergence and maintenance of hybrid incompatibilities during sympatric speciation between diverging sister species. An underdominant *f*_**(−)**_ mutation that reduces hybrid viability faces selective disadvantages because individuals carrying it produce fewer viable offspring through inter-niche mating than their same-niche counterparts without the mutation. Consequently, stronger premating RI, by significantly reducing inter-niche mating, mitigates the fitness cost associated with underdominance, thus facilitating the mutation’s initial emergence and long-term persistence.

In contrast, allopatric speciation completely halts inter-niche mating due to geographic barriers, eliminating the fitness penalties associated with underdominant *f*_**(−)**_ mutations. This geographic isolation enables hybrid incompatibilities to accumulate via the BDM mechanism without immediate negative consequences. In general, gene flow acts to eliminate or reconcile incompatibilities between populations [110]. Prolonged geographic separation provides intra-niche genotypes sufficient time to reconcile or eliminate internal incompatibilities, while inter-niche incompatibilities remain untested. Consequently, upon secondary contact, inter-niche hybrid incompatibility substantially exceeds intra-niche incompatibility, enabling stronger *f*_**(−)**_ mutations to persist via mechanisms akin to the Templeton effect [111, 112].

Robust premating RI—whether arising from coupled premating barriers in sympatric populations or geographic isolation in allopatric populations— alleviates negative selection pressure against underdominant *f*_**(−)**_ mutations responsible for hybrid incompatibility between emerging sister species. Similarly, underdominant *f* _**(−)**_ mutations situated within genomic regions of reduced recombination, such as chromosomal inversions, are shielded from purifying selection because suppression of recombination decreases the production of inviable hybrid offspring. This protection promotes the accumulation and maintenance of such mutations.

Conversely, weak premating RI between divergent ecotypes, occurring either during sympatric divergence or following secondary contact, exacerbates the fitness disadvantage of *f*_**(−)**_ mutations. Increased inter-niche mating under these conditions generates higher numbers of inviable hybrid offspring, thereby continually selecting against *f*_**(−)**_ mutations conferring hybrid incompatibility. As long as appreciable inter-niche gene flow persists, these incompatible *f*_**(−)**_ mutations are under constant selection pressure to be eliminated or replaced by variants lacking the incompatibility. Thus, the strength of premating RI profoundly influences the likelihood of emergence, establishment, and long-term persistence of post-zygotic incompatibilities between incipient sister species.

Thus far, we have considered that an *f*_**(−)**_ mutation reducing hybrid viability can arise only as an unintended and unavoidable byproduct of advantageous adaptations. It is unintended because, being underdominant, such a mutation confers no direct fitness advantage and thus cannot be favored by selection on its own. It is unavoidable because if the mutation could be decoupled from the beneficial adaptation that produced it, it would be purged by selection and replaced by alternative variants lacking this detrimental effect.

However, a scenario exists in which an *f*_**(−)**_ mutation may be positively selected to invade and spread within a niche population. This can occur during late-stage sympatric speciation, when strong premating RI and nonrandom assortment of locally adaptive alleles into distinct niches already exist. Under these conditions, a mutation reducing hybrid viability can further restrict gene flow and prevent introgression of maladaptive alleles from foreign ecotypes into the locally adapted population [110]. By preserving the integrity of parental genotypes against foreign introgression, the *f*_**(−)**_ mutation confers a direct fitness advantage to locally adapted genotypes carrying it relative to same-niche genotypes lacking the mutation. Consequently, the *f*_**(−)**_ mutation could gain a positive fitness advantage to invade and proliferate within the niche.

According to the biological species concept, reproductive isolation results from reduced gene flow between species. However, gene flow can only occur if viable hybrids are produced through inter-specific mating. Therefore, stopping gene flow requires stopping the production of viable hybrids. In natural systems, premating RI prevents inter-specific mating, thereby reducing the number of viable hybrid offspring produced. Subsequently, post-mating RI— e.g., in the form of hybrid incompatibility—acts to further eliminate any surviving hybrid offspring. Together, these two isolating mechanisms produce nearly complete RI between species.

Our simulation results have demonstrated that a low-gene-flow environment facilitates the emergence, establishment, and persistence of hybrid incompatibilities. The post-zygotic RI produced by the hybrid incompatibilities reinforces existing premating RI (Fig 26) and further restricts gene flow through a positive feedback mechanism, making RI in late-stage sympatric speciation robust and difficult to reverse.

Because the BDM mechanism of divergence and incompatibility is nonadaptive (i.e., not driven by hybrid loss), it can freely generate additional hybrid incompatibilities in environments with very low gene flow. Under these conditions, hybrid incompatibilities can accumulate through neutral drift or as incidental byproducts of ecological adaptations to distinct niches. Since the BDM mechanism is nonadaptive, in the absence of purifying selection, there is no inherent limit to the number of incompatibilities that can independently arise, regardless of the strength of existing RI. Consequently, multiple types of nonadaptive hybrid incompatibilities can accumulate within genomes over time, producing robust postmating RI. Eventually, reversing the RI produced by all these independently evolved incompatibilities through individual mutational changes becomes virtually impossible.

Chromosomal inversions are not static barriers but can act as dynamic elements in the evolution of reproductive isolation. In our speciation model, large CIs can serve as important early-stage barriers by maintaining linkage between locally adaptive alleles, mating-bias alleles, and hybrid incompatibility mutations. These large inversions effectively suppress recombination, preserving advantageous allele combinations and facilitating the establishment of RI. However, empirical observations suggest that while ancestral CIs tend to be large, inversions that appear later in evolution are often smaller, indicating that large CIs may not remain stable over time [35, 38, 113, 114]. One possible explanation is that large inversions undergo fission, breaking into smaller inversions that retain key divergence-associated loci while restoring recombination in other genomic regions. This process reduces the genetic load associated with recombination suppression and allows for fine-tuned adaptation. Additionally, new inversions may arise within previously inverted regions, reinforcing linkage between divergence-associated alleles and maintaining reproductive isolation even after some recombination is restored. These patterns suggest that while large inversions play a crucial role in facilitating early-stage divergence, post-speciation genomic architecture may evolve toward smaller, more refined inversions that balance recombination suppression with adaptive flexibility.

To conclude, in speciation theories, devising a plausible mechanism to explain the emergence and persistence of post-zygotic hybrid incompatibility in sympatric speciation remains challenging. This is because an *f*_**(−)**_ mutation that reduces hybrid viability is underdominant and subject to purging in the presence of gene flow. Although such a mutation may promote the speciation of niche-adapted ecotypes by producing post-mating RI and reinforcing existing premating RI, it suffers a fitness disadvantage relative to same-niche individuals lacking the mutation and is thus prone to elimination.

Strong premating RI, by reducing the production of unfit hybrids, can mitigate the underdominance disadvantage of an *f*_**(−)**_ mutation and facilitate its emergence and persistence. Likewise, greatly reduced hybrid viability—driven by strong disruptive ecological selection or pre-existing hybrid incompatibilities— can diminish the relative disadvantage faced by a new *f*_**(−)**_ mutation compared to same-niche genotypes lacking it. In such scenarios, all individuals within a niche already experience similarly low hybrid viability during inter-niche mating, so the accumulation of additional incompatibilities does not lead to further substantial increase in hybrid loss.

An *f*_**(−)**_ mutation can be considered a late-stage barrier mechanism because its emergence, invasion, and persistence are facilitated by strong premating RI in a low-gene-flow environment. It is adaptive because hybrid loss influences its underdominance and invasion fitness.

In our models, it appears the only way for such an *f*_**(−)**_ mutation to persist is for it to be linked with other beneficial alleles that confer a positive fitness advantage sufficient to offset its fitness disadvantage. This could occur if the *f*_**(−)**_ incompatibility is an integral and unavoidable part of another adaptive process. For instance, under the BDM barrier mechanism, hybrid incompatibility may evolve as an accidental and unintended byproduct of local adaptation and genomic divergence. In this case, hybrid incompatibility becomes linked to adaptive mutational changes within a niche environment.

The BDM mechanism operates most effectively under conditions of low gene flow. In such environments, an underdominant *f*_**(−)**_ mutation avoids the fitness cost typically associated with hybrid offspring loss from inter-niche mating, relative to same-niche counterparts. Therefore, the emergence and persistence of hybrid incompatibility are facilitated by strong RI between niche ecotypes. For this reason, transient allopatric divergence followed by secondary contact appears to be the most plausible mechanism to explain the emergence of hybrid incompatibility with advantageous ecological and mating-bias traits. However, at least in theory, if strong RI can also be produced by premating barriers in sympatric speciation, it should be able to facilitate the same BDM incompatibilities as effectively as geographical isolation in allopatric speciation.

Another way for an *f*_**(−)**_ mutation to exist in the face of gene flow is for it to be linked with other advantageous alleles in chromosomal inversions [34, 92]. Because a CI protects the genes in its inverted region from recombination, it can combine many favorable properties into a supergene that is transmitted as a single Mendelian locus. In our models, we can simulate the capture of an *f*_**(−)**_ mutation by decreasing the values of *FK*2 associated with CIs in Figs 19–21. By suppressing recombination, adaptive CIs can facilitate the emergence, evolution, and coupling of further mating-bias and hybrid incompatibilities within their inverted regions.

Incompatibilities arising within adaptive CIs can derive fitness benefits from their linkage with other locally advantageous alleles. Locally adaptive CIs in the genome reduce the effective number of ecological loci, which decreases the production of hybrids and mitigates the cost of incompatibilities. In late-stage speciation, when there is already strong assortment of locally adaptive alleles within niches, incompatibilities are positively selected to emerge and preserve the integrity of parental genotypes from the introgression of deleterious alleles.

In general, gene flow acts to purge and eliminate incompatibilities between divergent populations. However, because CIs suppress recombination, incompatibilities become more difficult to eliminate when they reside within CIs and are linked with locally adaptive alleles.

Thus, adaptive premating barriers and adaptive CIs with their evolved incompatibilities can mutually reinforce one another in a positive feedback loop to create progressively stronger RI. Eventually, in a low gene flow environment, the nonadaptive BDM mechanism becomes operative and generates—either through neutral drift or differential adaptations—many additional premating and post-mating incompatibilities as byproducts of irreversible mutational changes, completing and cementing the speciation process.

## V. LIMITATIONS OF THE STUDY

In our study, genetic inheritance from parents to offspring occurs through the random assortment of alleles at each gene locus. This approach is biologically realistic when the relevant genes are located on different chromosomes. However, if the genes reside on the same chromosome or within regions of varying recombination rates, specific recombination ratios for allele inheritance must be incorporated into the model.

Similarly, in our models, we assume that chromosomal inversions can capture adaptive alleles and gain a fitness advantage to invade and proliferate. However, this is only feasible if these alleles are located in close physical proximity on a chromosome to allow for their collective capture. In the genomes of closely related sister species, genes mediating ecological adaptation and reproductive isolation often cluster within islands of divergence [36, 61, 115-119]. The reasons for the existence of such genomic islands remain uncertain, even though several hypotheses have been proposed [120, 121]. Transposon activity, gene duplication, or adaptive divergence within CIs may position related genes in close proximity [71]. Perhaps, through processes of accretion or elimination, such clustering of related genes is favored by evolution, as they are less likely to be broken up by recombination [120]. In computer simulations, such “genomic clustering” often appears because of its adaptive advantage as a recombination suppressor [35, 61]. Selection may also favor the emergence of genes within CIs that can epistatically or pleiotropically affect adaptation or mating-bias genes located outside the CIs [62, 74]. In CIs that capture locally adaptive ecological alleles, further BDM divergence can lead to the evolution of hybrid and mating-bias incompatibilities within the CIs, thereby linking ecological traits with mating-bias traits. In addition, CIs that capture optimal allele combinations can transmit these sets en bloc, akin to a supergene [40, 107]. Nevertheless, the precise mechanisms underlying the physical clustering of adaptive alleles remain a topic of ongoing debate [61].

Multi-locus genotypes for ecological and mating traits appear to be the norm rather than the exception in nature, as is the existence of viable hybrids [115, 117, 122-124]. Polymorphisms due to balanced selection or frequency-dependent selection also abound [38, 53, 54, 125, 126]. Their true impact on our simplified toy model remains to be fully explored.

In our study, we have kept the number of variables in our models to a minimum for analytical tractability, while still ensuring they are sufficient to illuminate the fundamental principles of coupling between different barrier mechanisms. Additional important considerations—such as diploidy, gonochorism, dominance hierarchies, variable mating strategies, complex genotype-phenotype maps, and competition—would inevitably introduce more variables and increase model complexity. These factors could be integrated into our skeletal model in future work to better understand their effects and implications.

## VI. CONCLUSION

Our study uses computer simulations and analyses to investigate coupling among different reproductive barriers in late-stage sympatric speciation. In nature, closely related species often possess multiple different barrier mechanisms in their genomes that interact to reduce gene flow and make RI irreversible [15-19]. In a previous study, we developed a mathematical model to demonstrate how ecological selection and sexual selection can interact to produce an initial premating reproductive barrier based on population dynamics [28]. Here, we extend the model to demonstrate that an initial barrier can facilitate the coupling of additional barrier mechanisms to further strengthen and secure overall RI, ultimately completing the speciation process.

The barrier mechanisms analyzed in this study are summarized in Fig 26 and Table I. Broadly, they can be categorized as early-stage or late-stage, and as adaptive or nonadaptive. The early-stage barriers are all adaptive, meaning their invasion and coupling are driven by selection pressures arising from hybrid loss under disruptive ecological selection. These include the one- and two-allele models of mating-bias barriers, detailed in our previous study [28], as well as the one- and two-allele models of habitat-preference barriers examined in a separate study [29]. In sympatric populations under disruptive ecological selection, these early-stage barriers establish initial premating RI to reduce gene flow. The late-stage barriers include mutations that reduce hybrid viability, which can generate post-zygotic RI, as well as chromosomal inversions and BDM incompatibilities, both of which can produce pre-zygotic and post-zygotic RIs. The invasion fitness of an adaptive CI is at its most optimal in an environment with intermediate gene flow, whereas the BDM barrier mechanism is most effective in an environment of low gene flow.

Our general modeling approach is to create a mutant allele with properties characteristic of different barrier mechanisms. We then developed computer applications to investigate the invasion fitness of this mutant allele as it attempts to couple with existing barrier systems and to observe the resultant population dynamics. This allows us to examine how the coupling of additional barriers affects the strength and reversibility of overall RI.

The invasion fitness of a mutant allele is the sum of the fitness contributions of its component properties in both intra-niche and inter-niche matings. For example, in environments with reduced gene flow, an allele with a high mating bias may gain a fitness advantage in inter-niche mating by reducing hybrid offspring loss but suffer reduced fitness in intra-niche mating due to intra-niche sexual selection. Changes in parametric variables—such as the strength of disruptive ecological selection, assortative mating costs, and niche carrying capacities—alter the invasion fitness of the mutant allele by shifting the balance between its fitness advantage in inter-niche mating and its fitness disadvantage in intra-niche mating. Similarly, a mutation that reduces hybrid viability is underdominant in inter-niche matings and, on its own, cannot successfully invade. It needs to hitchhike with other positively-selected properties to gain a net fitness advantage to invade and exert its synergistic effects on overall RI.

Positive-effect and negative-effect properties can be combined through various mechanisms that suppress their dissociation by recombination, such as close physical proximity on a chromosome, relocation to low-recombination genomic regions, pleiotropy, or the action of recombination suppressors. In our models, these properties are combined within the same mutant allele because they are integral to the barrier mechanism the allele embodies. For instance, chromosomal inversions can link alleles with both positive and negative invasion fitness into a super-allele with combined properties. By integrating both premating and post-mating barrier properties, a CI with net invasion fitness can achieve a synergistic reduction in gene flow and enhanced RI. Additionally, in systems with multi-locus mating-bias genotypes, CIs can acquire invasion fitness by capturing advantageous high-mating-bias alleles to reduce mating-bias hybrids and promote speciation [30]. While our models use CIs as a framework for combining these properties, equivalent effects can be achieved when alleles are linked by physical proximity or other recombination suppressors. Likewise, the BDM mechanism facilitates the evolution of locally adaptive ecological fitness along with hybrid and mating-bias incompatibilities, thereby achieving strong and irreversible RI through their combined effects. Evolving ecological adaptations before incompatibilities prevents invasion resistance arising from hybrid loss and intra-niche sexual selection. Antagonistic pleiotropy in ecological fitness across niches ensures that evolved BDM traits remain within their adapted niche and prevents their introgression into the opposite niche, where they could weaken BDM-mediated inter-niche incompatibilities and RI.

Our simulation results demonstrate that after an initial mating-bias barrier reduces gene flow, it can facilitate coupling with various other barrier mechanisms to produce progressively stronger and irreversible RI. For early-stage barriers, a first barrier that reduces gene flow can facilitate the coupling of a subsequent two-allele mating-bias barrier. However, if the initial barrier creates very low gene flow, it can result in an invasion-resistant pattern in the phase portrait of the second mating-bias barrier. This invasion resistance can be eliminated by increasing the number of mating rounds (*n*) or by reducing the niche carrying capacity (e.g., *NA*). In contrast, habitat-preference barriers can always couple with existing barriers as long as maladaptive hybrid loss remains [29].

Both mating-bias and habitat-preference barriers can independently establish initial barriers, which can then couple with other similar adaptive premating barrier mechanisms to progressively increase premating RI and further reduce gene flow. Because habitat-preference barriers are more readily established than the two-allele mating-bias barrier, establishing habitat preference as the initial barrier and subsequently coupling it with a mating-bias barrier presents a compelling pathway for initiating premating RI in sympatric speciation [29].

Late-stage barrier mechanisms, such as chromosomal inversions and the BDM mechanism, operate most effectively in a low gene flow environment. Their coupling with adaptive premating barriers could create both premating and post-mating RI, further strengthening and securing the existing RI.

Adaptive barriers, arising from changes in population dynamics and driven by selection pressure due to hybrid loss, are readily reversed when disruptive ecological selection is reduced or eliminated. Consequently, adaptive mutational changes, such as those involving chromosomal rearrangements and the BDM mechanism of divergent adaptations and incompatibilities, appear necessary to secure more permanent and lasting RI [56, 99]. Therefore, the role of adaptive barriers is to quickly secure a low gene flow environment for these late-stage barrier mechanisms to take hold and mature, making the RI irreversible.

A positive feedback loop between early-stage and late-stage barrier mechanisms is created (Fig 26). Initially, early-stage adaptive premating barriers reduce gene flow and facilitate the coupling of late-stage barriers, which create pre-zygotic and post-zygotic RIs that further strengthen and secure the early-stage premating barriers. This leads to a further reduction in gene flow, which in turn facilitates the late-stage mechanisms, continuing the cycle. The end result is that, over time, the overall RI becomes progressively stronger and irreversible, ultimately leading to the completion of the speciation process.

However, once initial premating RI is established, the invasion of adaptive CIs capturing only locally adaptive ecological alleles can weaken existing barriers through negative coupling, disrupting the feedback loop. Nonetheless, these CIs may also facilitate linkage among ecological alleles, mating preferences, and hybrid incompatibilities, thereby restoring the positive feedback loop and the trajectory toward complete speciation.

In general, mutations that reduce hybrid viability are underdominant and must hitchhike with beneficial processes—such as linkage to advantageous alleles within adaptive CIs—to invade and spread. By suppressing recombination within their inverted regions, CIs can promote the accumulation of hybrid incompatibilities through ongoing local adaptation and BDM divergence [33, 41, 54, 109, 127]. In late-stage speciation, when adaptive alleles are already assorted across niches, a mutation causing hybrid incompatibility may also gain a direct selective advantage by blocking the introgression of deleterious foreign alleles and preserving the integrity of parental genotypes [110].

Strong premating RI prevents the formation of maladaptive hybrids and reduces the fitness cost of mutations that cause hybrid incompatibility, making their invasion and persistence more likely. Eventually, the nonadaptive BDM mechanism can take over and evolve multiple independent incompatibilities as byproducts of continued adaptation and divergence. These incompatibilities arise from mutational changes and are difficult to reverse. Their cumulative effects produce strong, irreversible RI that completes the speciation process.

In our previous study, we proposed a theoretical model for establishing an initial premating reproductive barrier in a sympatric population under disruptive ecological selection [28]. Here, we have demonstrated that this initial barrier can couple with additional premating and post-mating barrier mechanisms to strengthen overall RI and render the speciation process irreversible. Together, the results of our two studies outline a pathway forward for explaining the full progression of sympatric speciation. While our proposed mechanism still requires empirical validation and theoretical refinement, we hope it offers a valuable framework that can be improved and extended through future research and discussions to advance our understanding of sympatric speciation.

## VII. ACKNOWLEDGMENTS

We would like to express our grateful appreciation to Ashley Carter, David Hernandez and Karyn How for reviewing the manuscript and offering valuable comments.

## VIII. APPENDIX

